# Controlled Delivery and Light-Induced Release of Magic Spot Nucleotides in *Escherichia coli*

**DOI:** 10.1101/2025.10.22.683773

**Authors:** Christoph Popp, Patrick Moser, Xinwei Liu, Johannes Freitag, Isabel Prucker, Robert Zscherp, Xuan Wang, Philipp Klahn, Hans-Georg Koch, Gert Bange, Henning J. Jessen

## Abstract

The “magic spot nucleotides” (MSNs) ppGpp and pppGpp (also: (p)ppGpp) are bacterial alarmones central to the conserved stringent response, a stress adaptation mechanism that helps bacteria adapt to stress conditions and hostile environments. Current strategies to manipulate MSN levels rely mainly on genetic or environmental approaches, which are slow and lack temporal control. Chemical tools such as photocaged MSN analogues could provide such temporal control over MSN levels. However, the high negative charge of MSNs prevents spontaneous passage through the complex bacterial cell envelope. Here, we report the synthesis of photocaged, clickable, and isotope-labeled MSN analogues and their delivery into *Escherichia coli* comparing different approaches. A cyclodextrin-based synthetic nucleotide transporter provides particular advantages. Upon 400 nm irradiation, these probes were photo-released inside living cells, where we tracked their conversion from pppGpp to ppGpp by capillary electrophoresis mass spectrometry and studied their ability to alter growth in a (p)ppGpp^0^ mutant. This work provides the first demonstration that highly charged, photocaged MSNs can traverse the bacterial envelope, be photo-released intracellularly, and be metabolically tracked in real time. These probes lay the foundation for spatially and temporally controlled studies of MSN function and of other highly negatively charged metabolites in bacteria.

## Introduction

When bacteria are exposed to stress conditions like temperature change, pH change, starvation, or antibiotics, alarmones like the magic spot nucleotides (MSNs) guanosine tetraphosphate (ppGpp) and guanosine pentaphosphate (pppGpp) accumulate and induce the stringent response (Figure 1A).^[1–5]^ MSNs help bacteria survive by inducing a dormant-like state until conditions improve. The extent of stringency is a function of MSN concentration. Beyond stress adaptation, MSNs also act as second messengers during normal growth, where basal levels are essential for regulating transcription, translation, replication, and metabolism through a wide network of protein interactions.^[4,6]^

**Figure 1.**
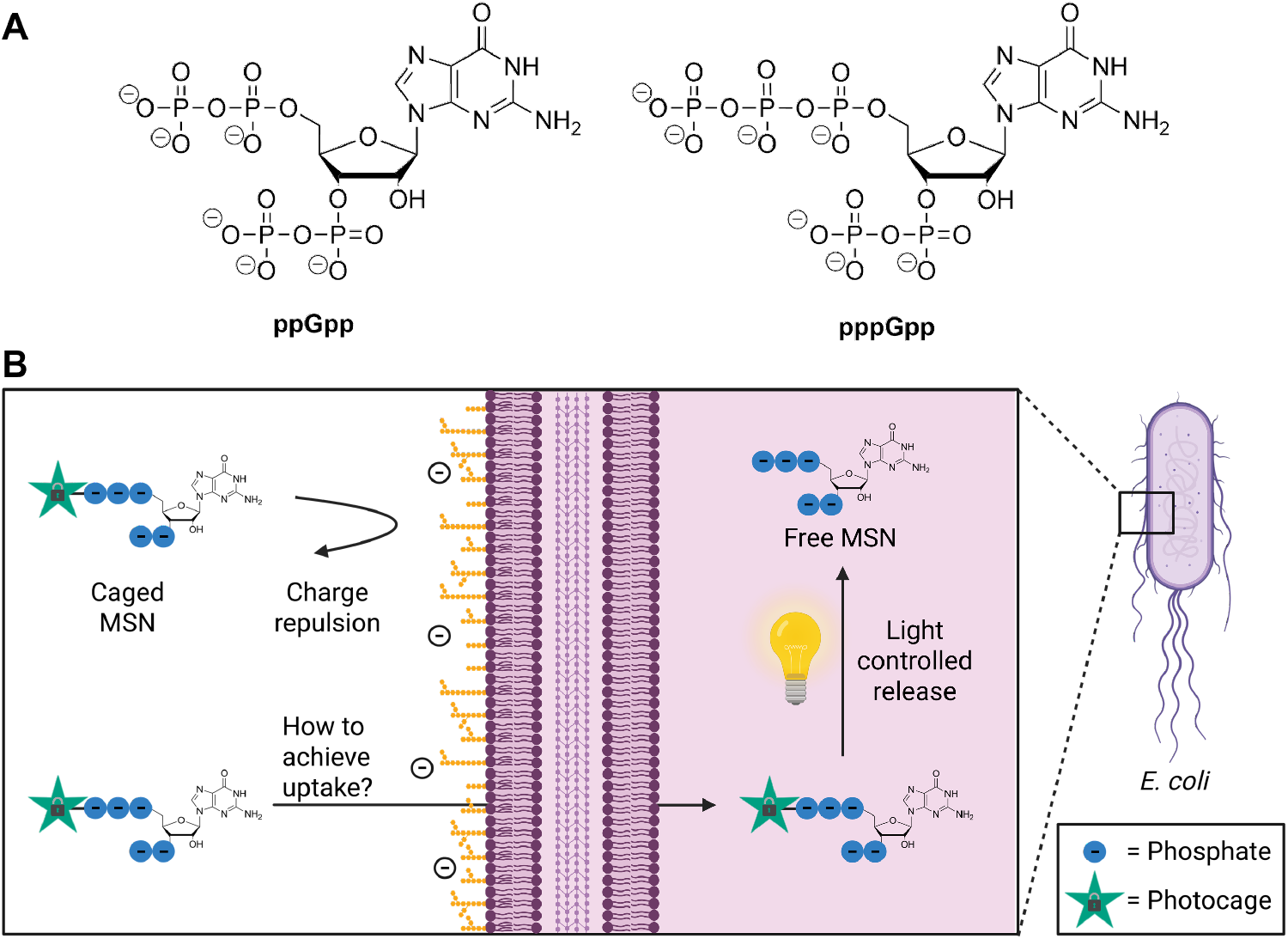
(A) Chemical structure of ppGpp and pppGpp. (B) Overview of our strategy to modulate MSN concentrations inside living cells. Due to charge repulsion from the negatively charged lipopolysaccharides and lipids, MSNs cannot readily pass through the complex bacterial cell envelope. We tested various strategies to deliver a photocaged MSN analogue into the bacterial cell and release the MSNs by light irradiation. Created in BioRender. Popp, C. (2025) https://BioRender.com/wnqejr2

To study the concentration-dependent role of MSNs in cellular processes it is useful to be able to manipulate their abundance. Currently this can be achieved by genetic modifications, treatment with small molecule drugs or exposure to stress conditions.^[7–9]^ Additionally, it would be highly desirable to study effects of local, constrained pools of these molecules. Genetically modifying organisms is time consuming and often causes secondary effects. In addition, MSN-producing enzymes can synthesize both pppGpp and ppGpp, making it difficult to distinguish between different MSNs. To influence concentration levels of single MSNs further genetic modifications are needed.^[10]^ An alternative chemical biology approach for modulation of MSN levels inside bacteria would provide new avenues for complementary experiments and thus enable further investigations into the effects of MSNs on bacterial physiology. The aim of this study is to deliver photocaged MSNs into bacterial cells thus enabling their controlled release by light (Figure 1B). Since *in vitro* data of *Escherichia coli* and *Bacillus subtilis* MSN synthesizing enzymes RelA and SAS1 show that MSNs can stimulate their own synthesis, the introduction of small amounts of MSNs into the bacteria might be sufficient to induce high cellular concentrations.^[8,11–15]^

The main obstacle for this approach is the bacterial cell envelope. In contrast to the lipid bilayer of mammalian cells, bacteria possess a much more challenging barrier. In Gram-negative bacteria the cell wall is composed of several peptidoglycan layers surrounded by an outer membrane containing phospholipids and lipopolysaccharides. MSNs can carry up to seven negative charges that impair translocation through the negatively charged cell envelope.^[16,17]^ Several methods have been developed to transport highly negatively charged small molecules like nucleotides into mammalian cells. For example, prodrugs with masked negative charges and/or lipophilic groups,^[18–21]^ positively charged polymers,^[22,23]^ covalently linked cationic groups,^[24,25]^ permeabilization of the plasma membrane with surfactants^[26]^ or by physical and mechanical means^[27–30]^. In bacteria, however, delivery studies of charged metabolites are very limited and only few examples are known. It is possible to transport non-natural nucleotides into bacterial cells by heterologously expressing importers from other species.^[31,32]^ However, this approach is limited to several non-natural nucleotides and needs genetic modification of the organism. Carlson and co-workers reported the development of a cationic polymer, promoting internalization of an adenosine triphosphate derived chemical probe into *E. coli* and *B. subtilis*.^[33]^ They proposed that the cationic polymers act as general permeabilization reagents that promote the entry of various molecules into the cytosol. Kraus and coworkers used a cyclodextrin based synthetic nucleotide transporter (SNT).^[34]^ It consists of a per-6-amino-β-cyclodextrin coupled to a peptide consisting of eight aminocaproic acid/arginine pairs. The cyclodextrin binds the nucleotide while the peptide disrupts the membrane to facilitate cellular delivery. They postulate that the bound nucleotide is released in the cytosol due to replacement by endogenous nucleotides. Successful delivery of a fluorescently modified deoxyuridine triphosphate into live *Mycobacterium smegmatis* and *E. coli* was shown with this approach, yet the study was mainly focused on delivery into mammalian cells.^[34]^

Other highly negatively charged molecule that are routinely transported into bacterial cells are nucleic acids. Transformation is usually done by electroporation or by heat shock of chemically competent cells.^[35–38]^ However, it is also possible to use other methods such as cationic polymers and peptides.^[39,40]^ For mammalian cells several advanced methods have been developed, allowing targeted delivery of nucleic acids *in vivo*.^[41]^ It has to be noted though that for many applications of nucleic acid delivery, it is sufficient to deliver very small copy numbers, as these will be amplified or are catalytically active. However, in case of nucleotide transport, high concentrations are desirable.

For less polar or cationic substances several further methods have been developed to facilitate delivery.^[42,43]^ One example that showed promising results for cargo delivery into various gram-negative bacteria are siderophores.^[44,45]^ Coupling these Fe(III)-chelating molecules to a cargo allows hijacking of the siderophore-mediated iron uptake pathway. This approach was extensively studied for antibiotic conjugates but was also shown to deliver larger molecules like nucleic acid therapeutics into bacteria.^[45–50]^ Carbohydrate conjugates were used to deliver imaging probes into bacteria through the bacteria-specific maltodextrin transport pathway. ^[51,52]^ Other approaches involve the use of nanoparticles^[41,53]^, cell-penetrating peptides^[54–56]^, boron clusters^[57]^, or disulfides.^[58,59]^

An established method to control the release of biologically active molecules inside cells or tissue is the introduction of photolabile protecting groups, so called photocages.^[60–64]^ These structures are covalently bound to the bioactive compound rendering them biologically inert and can be cleaved by light irradiation. The use of light allows temporal and spatial control over the release of the biomolecule. One commonly used caging group is based on the coumarin scaffold. The coumarin photocage stands out through high biocompatibility, easy synthesis, flexibility of structural modifications and a well-studied mechanism of photocleavage.^[65,66]^

Here, we demonstrate that coumarin caged MSN can be delivered into living *E. coli* cells using a modified cyclodextrin as additive and that they can be released on demand by irradiation with light.

## Results and Discussion

### Synthesis of Caged MSNs

The synthesis of caged MSNs followed our previously reported synthetic methods employing chemoselective phosphorylation with phosphoramidites and regioselective RNase T2 hydrolysis (Scheme 1).^[67–71]^ The synthesis started with guanosine or ^15^N–labeled guanosine, which was treated with pyrophosphoryl chloride. Controlled hydrolysis leads to the 2′-3′ cyclophosphate which was regioselectively opened with RNase T2 to form pGp (**3**) or ^15^N– pGp (**4**).^[69,70]^ Unlabeled and ^15^N–labeled pGp was phosphorylated with bisfluorenmethyl phosphoramidite (**5**), oxidized and Fm-deprotected to form ppGpp (**6**) and ^15^N–ppGpp (**7**). ^15^N–ppGpp was purified, while unlabeled ppGpp was used without purification. Treatment with RNase T2 and purification by strong anion-exchange chromatography yielded ppGp (**8**) and ^15^N–ppGp (**9**).^[69,70]^

For photocaging, we adopted the 7-(diethylamino)-4-(but-3-yn-1-yl)coumarin (DEACBY) chromophore introduced by Seyfried *et al*.^[72]^. We chose to use this photocage because of its easy access and further because the alkyne group would enable late-stage functionalization of the caged MSNs by copper-catalyzed azide-alkyne cycloadditions.^[73,74]^

Treatment of ppGp (**8**) and ^15^N–ppGp (**9**) with mixed P–amidite **10** containing the photocage DEACBY followed by oxidation with *m*CPBA led to the 3′-diphosphate 5′-triphosphate nucleotides with triesters at the terminal phosphates. The triester at the 3′-end acts as a leaving group when the molecules are dissolved in MeOH allowing fast and efficient cyclization to the corresponding 2′-3′ cyclophosphates. The fluorenmethyl (Fm) group was removed with piperidine and the cyclophosphate was regioselectively opened with RNase T2 leading to 5′-DEACBY protected pppGp (**11)** and ^15^N – pppGp (**12)** in 58 % (97 % considering recovered starting material) and 52 % yield, respectively. Bisfluorenmethyl phosphoramidite reacted chemoselectively with the monoester phosphate. Oxidation and Fm deprotection with 1,8-diazabicyclo[5.4.0]undec-7-en led to the caged MSNs DpppGpp (**13**) and ^15^N-DpppGpp (**14**) in 65 % and 41 % yield respectively. The efficiency of this synthesis enabled the production of 13 mg ^15^N labeled caged pppGpp (**14**) starting from 70 mg ^15^N-labeled ppGpp (**7**).

**Scheme 1.**
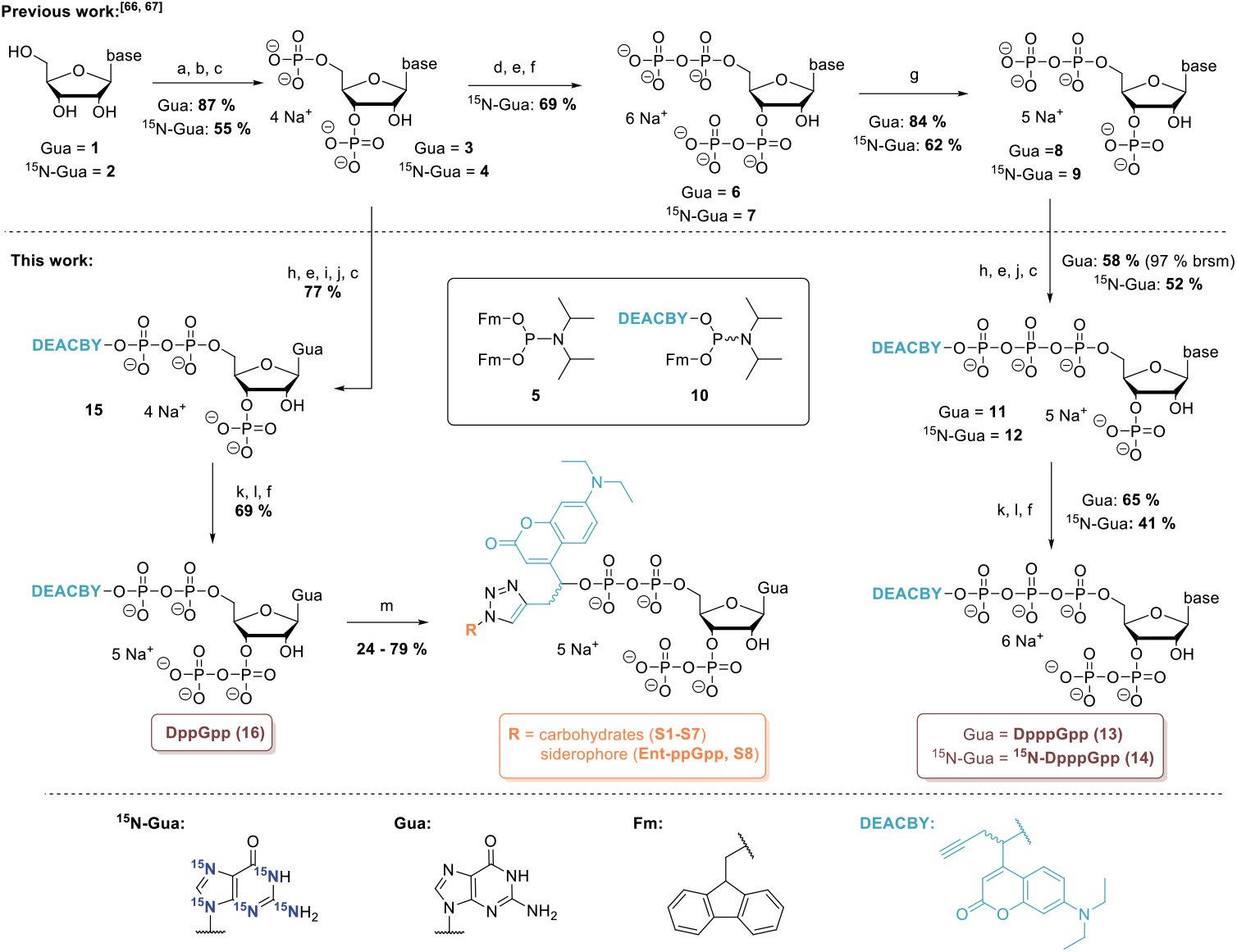
Synthesis of caged and isotope labeled MSN derivatives.^[69,70]^ a) P_2_Cl_4_O_3_, (11 eq), 0 °C, 3 h. b) NaHCO_3_ (1 M), 0 °C. c) RNase T2, pH 7.5, 37 °C, 12 h. d) **5** (3.0 eq), ETT (5.0 eq), DMF, rt, 15 min. e) mCPBA (3.0 eq), – 20 °C, 10 min. f) DBU, 0 °C→rt, 30 min. g) RNase T2, aq. HCl, pH 5.5, 37 °C. h) **9** (2.3 – 2.4 eq), ETT (5.0 eq), DMF, rt, 1 h. i) MeOH, rt, 5 min j) Piperidine, DMF, rt, 5 min. k) **5** (1.5 – 1.7 eq), ETT (3.0 – 3.5 eq), DMF, rt, 15 min. l) mCPBA (1.5 – 2.1 eq), -20 °C, 10 min. m) Azide (1.5 eq), Na-ascorbate (5.0 eq), CuSO_4_ (0.1 eq),THPTA (0.5 eq), H_2_O, rt, 3 h. Abbreviations: brsm: based on recovered starting material, DBU: 1,8-diazabicyclo(5.4.0)undec-7-ene, ETT: 5-(ethylthio)-1H-tetrazole, Fm: fluorenmethyl, Gua: guanine, mCPBA: meta-chloroperbenzoic acid, rt: room temperature. THPTA: tris(3-hydroxypropyltriazolylmethyl)amine.

To synthesize the caged magic spot nucleotide DppGpp, pGp was treated with mixed amidite **9**, oxidized, cyclized in methanol and deprotected. Regioselective ring-opening with RNase T2 led to DppGp (**14**) in 77 % yield on a 700 mg scale. Another cycle of P-amidite coupling with bisfluorenmethyl phosphoramidite, that selectively only reacts with terminal phosphates,^[75]^ oxidation and Fm deprotection with DBU led to the caged guanosine tetraphosphate DppGpp (**15**) in 69 % yield.

The alkyne handle on the photocage now offered the possibility to easily introduce structures that could facilitate uptake into bacterial cells, while photocleavage would eventually remove them again. Various carbohydrates (glucose, galactose, mannose, maltose, lactose, trehalose) and a biomimetic enterobactin analogue^[76]^ were readily coupled to DppGpp via copper-catalyzed azide-alkyne cycloadditions^[73,74]^ with yields of 24–79 % (**S1– 8**).

For control experiments, caged adenosine triphosphate (D-ATP, **S9**) and caged guanosine diphosphate (D-GDP, **S10**) were synthesized by treating commercial ADP and GMP with mixed amidite **9**. Oxidation, and Fm deprotection, yielded D-ATP and D-GDP.

### Photolysis Behavior

To study the photolysis behavior of the coumarin photocages, the caged MSNs DppGpp (**16**) and DpppGpp (**13**) were irradiated with a 400 nm LED in a small glass vial (25 nmol, 50 µM, 14 mW), and the uncaging was monitored by HPLC (Figure 2). After 2 min irradiation, only trace amounts of caged MSNs remained and the corresponding free MSNs were formed without detectable nucleotide byproducts indicating a clean and efficient photocleavage.

**Figure 2.**
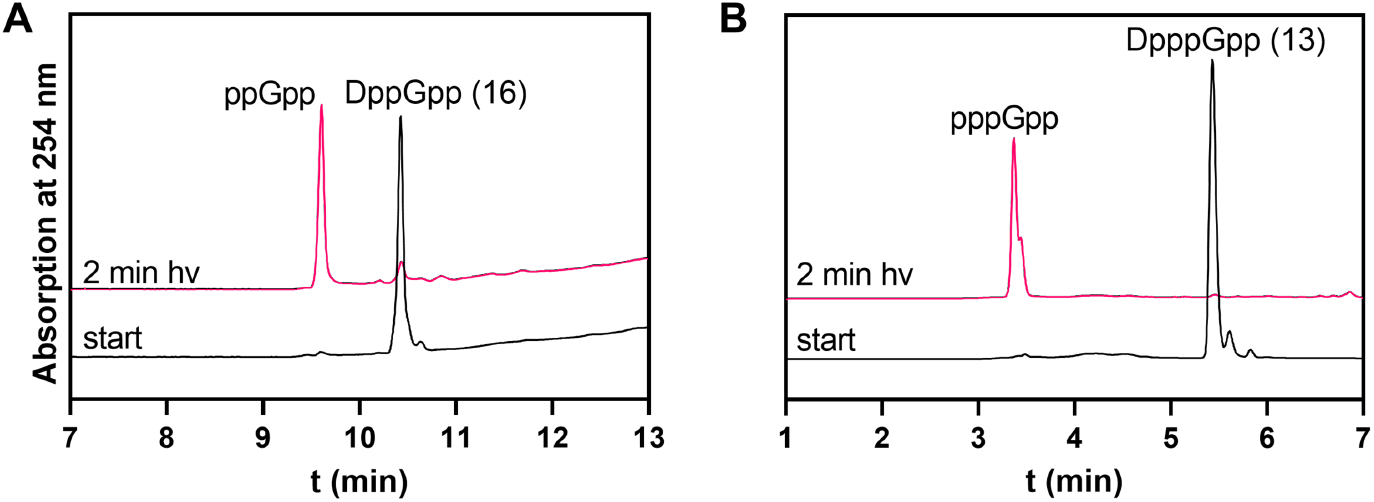
HPLC chromatograms (254 nm) of photolysis of DppGpp (A) and DpppGpp (B). Chromatograms were measured before and after irradiation with a 400 nm LED (25 nmol, 50 µM, 14 mW).

### Uptake of MSN Probes into *E. coli*

Next, uptake into bacteria was assessed by incubating *E. coli* cell suspensions with solutions of the caged nucleotides. After incubation, the cells were separated by centrifugation, washed, resuspended and fluorescence of the cell suspension was measured. The intrinsic fluorescence of the DEACBY caging group enabled this rapid initial study of potential delivery. MSN derivatives coupled to carbohydrates and siderophore (**S1-8**) did not produce fluorescence above the autofluorescence of *E. coli* cells (Figure 3A, S2) under the experimental conditions and thus further optimization was required.

**Figure 3.**
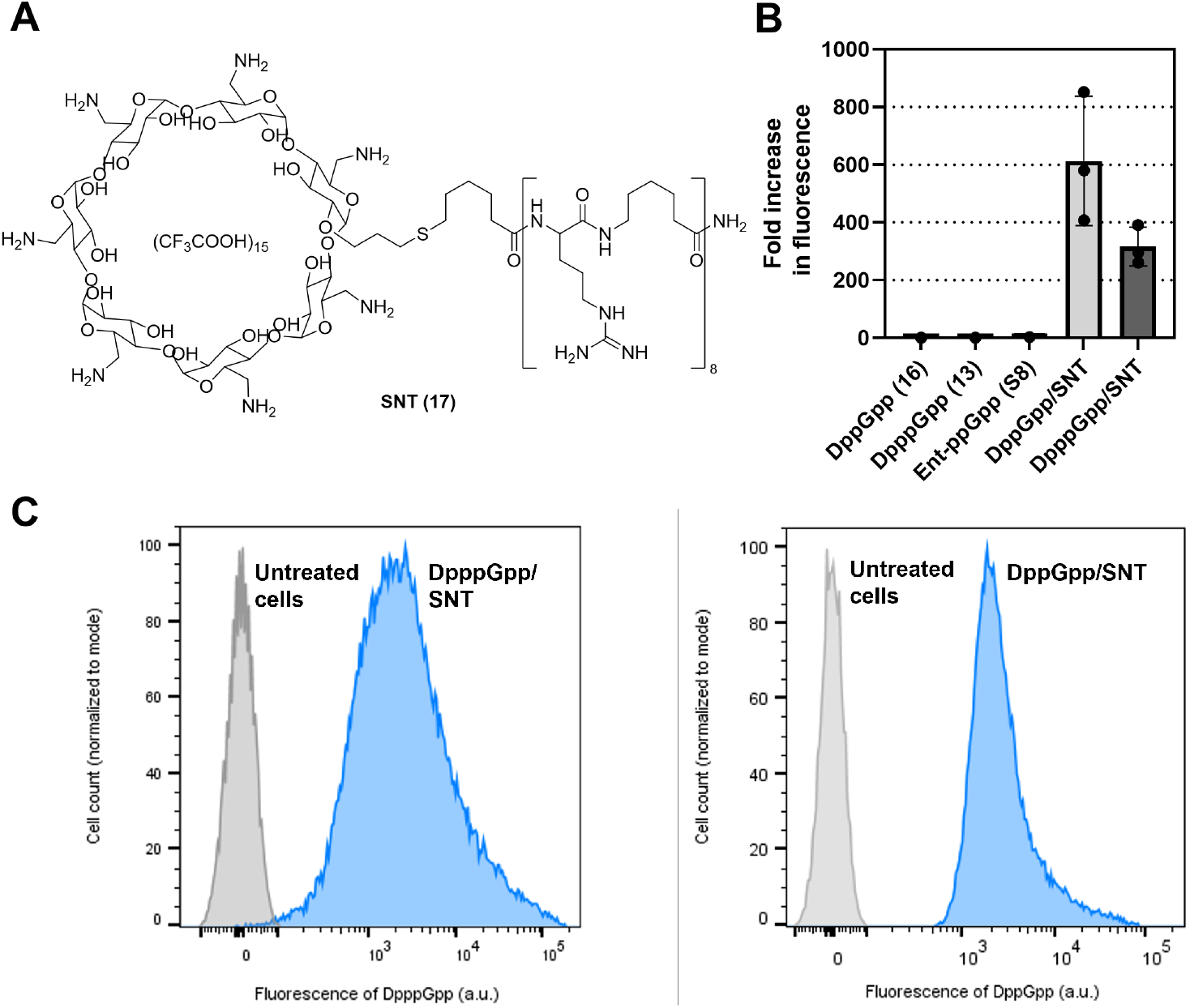
Uptake measurements of MSN in E. coli. (A) Structure of the synthetic nucleotide transporter (SNT). (B) Cellular uptake determined by measuring fluorescence of cell suspensions. Values are normalized to the autofluorescence of cells treated with DppGpp or DpppGpp alone. Mean ± standard deviation is shown. (C) Histogram plots of flow cytometry measurements indicate MSN cage fluorescence in nearly every cell.

Therefore, we studied the synthetic nucleotide transporter (SNT, **17**, Figure 3A) developed by Kraus.^[34]^ This commercially available compound was mixed with DppGpp (**16**) and DpppGpp (**13)** and then incubated with an *E. coli* suspension. After washing, the cell suspension exhibited a substantial increase in fluorescence, a first indication of efficient delivery (Figure 3B). Flow cytometry confirmed these results (Figure 3C), demonstrating largely increased fluorescence of nearly all bacterial cells.

### Confocal Microscopy Analysis of Cellular Distribution

The previously described experiments cannot distinguish whether the MSNs are just bound to the exterior of the *E. coli* membrane or if they are internalized into the periplasm or cytoplasm. Thus, uptake was additionally evaluated with confocal microscopy under optimized conditions using the high throughput methods described above (Figure 4A+B). The cellular membranes were stained with Nile red. The intensity profiles of Nile red fluorescence compared to DEACBY fluorescence show the distribution of the caged MSNs inside the bacterial cells (Figure 4C+D). While Nile red fluorescence is maximal at the start and at the end of the intensity profiles representing the cellular membrane, DEACBY fluorescence is maximal between the Nile red maxima. This clearly demonstrates that the caged MSNs enter the cytoplasm of *E. coli* cells. Additionally, confocal fluorescence images revealed heterogeneous caged MSN uptake into different bacterial cells. Some cells exhibit high intracellular fluorescence, while others show low or even no DEACBY fluorescence, contrasting with the flow cytometry measurements in Figure 3C.

**Figure 4.**
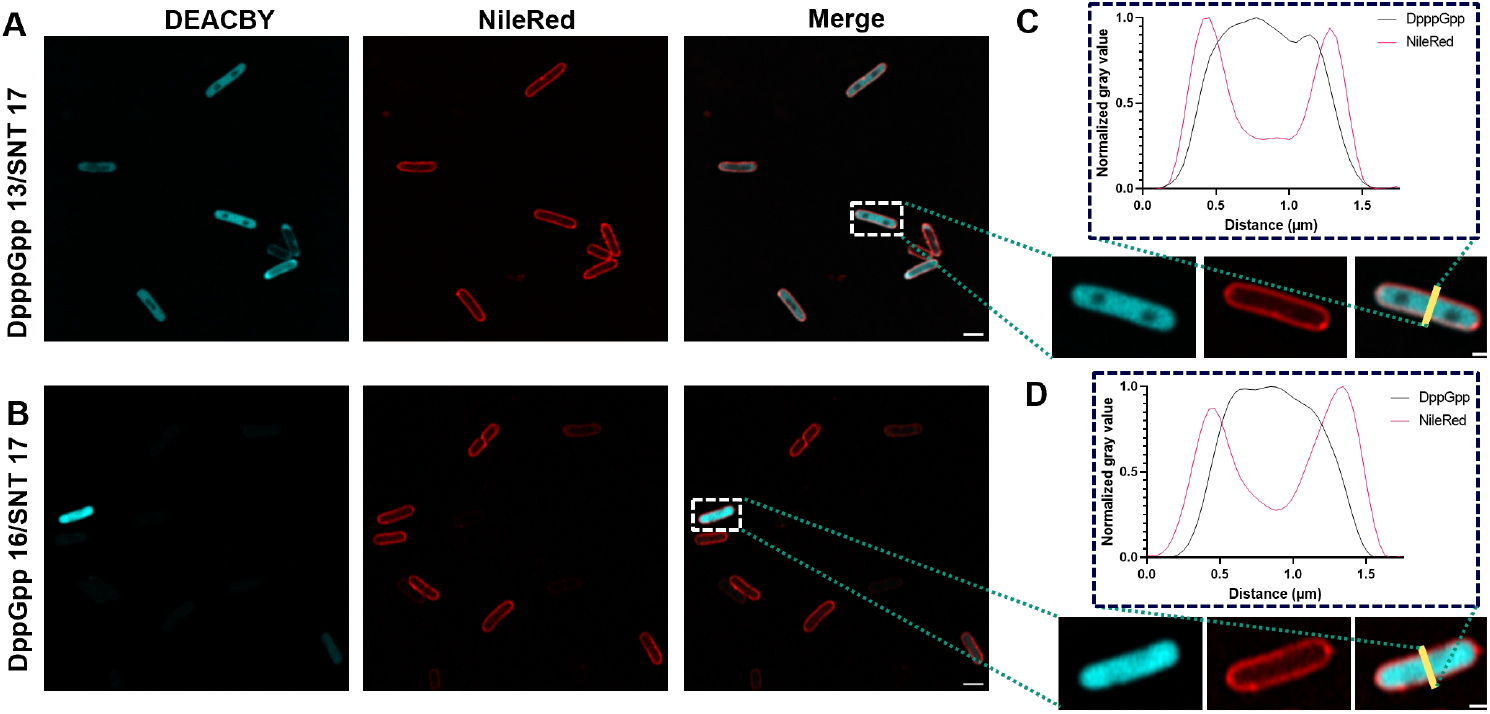
Microscopy images of E. coli treated with a mixture of DppGpp (16, B + D) or DpppGpp (13, A + C) with SNT (17) and the membrane stain Nile Red (red fluorescence). The fluorescence of the photolabile protecting group DEACBY is shown in cyan. C + D) Intensity profiles through a single cell normalized to the highest and lowest grey values. Scale bar = 2 µm in large images and 1 µm in magnified images.

Cell viability was checked next. It was found that using 50 µl/OD_600_/ml incubation led to no reduced viability, while 200 µl/OD_600_/ml led to a decrease of viability to 50% (Figure S6).

### Metabolic Tracking of MSNs

In order to confirm that uncaging inside *E. coli* is possible and to track the fate of the released MSNs, we applied an extraction and quantification assay using capillary electrophoresis mass spectrometry (CE-MS).^[69,77]^ After incubation with DpppGpp (**13**) and SNT (**17**) and successive washing, the *E. coli* cells were resuspended in medium the first sample for MSN extraction was taken and the cell suspension was irradiated by a 400 nm LED (14 mW, 3 min) to release the free nucleotides (Figure 5A). The culture was then incubated at 37 °C for 1 h and samples were collected at predetermined time points before and after irradiation. The samples were lysed, spiked with heavy isotope-labeled standards and extracted using weak anion exchange. Quantification was subsequently performed by CE-MS.

**Figure 5.**
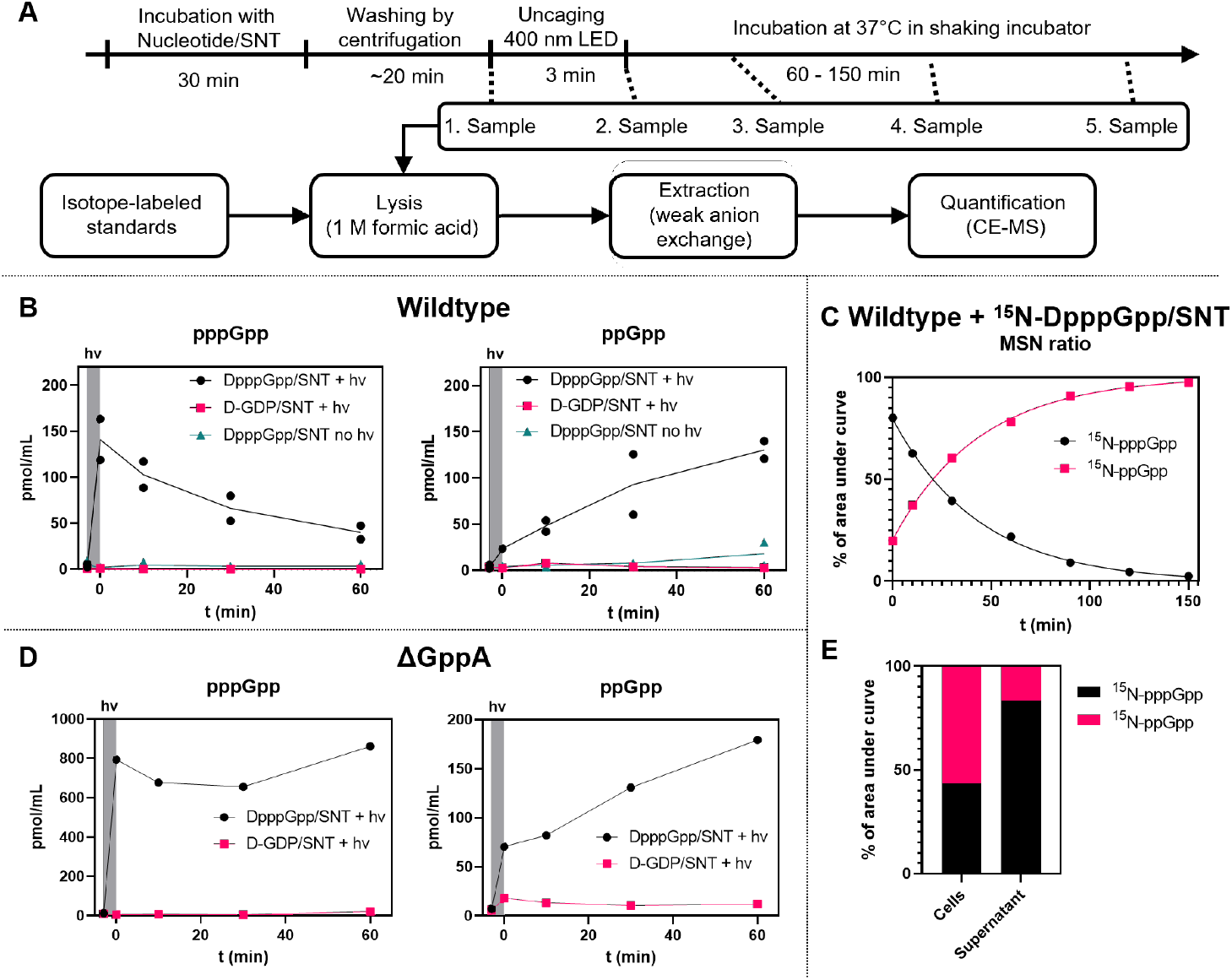
MSNs extracted from E. coli after incubation with caged MSNs. (A) Experimental timeline of extraction experiments. (B) pppGpp and ppGpp concentrations extracted from cell suspension of E. coli cells incubated with DpppGpp/SNT or D-GDP/SNT. Cell suspension was irradiated with 400 nm light before t = 0 min. (C) Ratio of ^15^N labeled pppGpp and ppGpp extracted from E. coli cell suspension incubated with ^15^N-DpppGpp/SNT and irradiated with 400 nm light. Before irradiation, no ^15^N-labeled nucleotides were detected. (D) pppGpp and ppGpp concentrations extracted from cell suspension of E. coli ΔgppA mutant incubated with DpppGpp/SNT or D-GDP/SNT. Cell suspension was irradiated with 400 nm light before t = 0 min. (E) Ratio of ^15^N labeled pppGpp and ppGpp extracted from E. coli cell suspension that was incubated with ^15^N-DpppGpp/SNT and irradiated with 400 nm light. After uncaging, cells were separated by centrifugation (5 min, 5 kG) and then immediately lysed in 1 M formic acid.

After irradiation with 400 nm light, we observed a direct and sharp increase in pppGpp levels, confirming successful photolysis (Figure 5B). Over time, pppGpp levels decreased while ppGpp increased, consistent with phosphatase-mediated hydrolysis. In *E. coli* GppA is the phosphatase responsible for the conversion of pppGpp to ppGpp. In fact, ppGpp ist the main alarmone in *E. coli* due to GppA activity. ^[78,79]^ This was confirmed using ^15^N-DpppGpp (**14**): we detected a corresponding increase in [^15^N]_5_-ppGpp as [^15^N]_5_-pppGpp declined (Figure 5C), indicating that the ppGpp originates mostly from the uncaged ^15^N-labeled pppGpp and not through indirect effects of the uncaging process. Bacteria treated with caged GDP or bacteria that were not irradiated with 400 nm light did not show this change in MSN levels (Figure 5B).

To further validate this assumption, we also performed the extraction experiment with a *ΔgppA* mutant^[80]^ (Figure 5D). In this strain, pppGpp levels remained stable due to the absence of GppA, however, ppGpp still accumulated, consistent with previous reports that pppGpp allosterically activates RelA, thereby enhancing ppGpp synthesis.^[12,15]^

We performed another control: Bacteria were treated with DpppGpp (**13**) and SNT (**17**) and [^15^N]_5_-pppGpp was spiked into the medium directly after uncaging. The conversion of the heavy-isotope labeled [^15^N]_5_–pppGpp to [^15^N]_5_–ppGpp resembled the conversion of pppGpp to ppGpp (Figure S7), even though [^15^N]_5_–pppGpp should only be present in the medium, while pppGpp should be located inside the bacterial cells. These results could imply that periplasmic or extracellular phosphatases, such as PhoA^[81]^ and/or phosphatases released from damaged cells can also process MSNs. Furthermore, damaged cells may release MSNs into the culture medium or take up ^15^N-labelled MSNs from the medium.

To better understand which portion of the quantified nucleotides were truly intracellular, we separated the cell pellets and the culture supernatants after photolysis by centrifugation and analyzed both fractions. We could detect MSNs in both fractions, but surprisingly, the supernatant contained more pppGpp but less ppGpp than the pellet fraction (Figure S8). This shows that MSNs leak into the medium, potentially through the action of the SNT. However, rapid intracellular metabolism could reduce detectable MSN levels inside the cells, exaggerating the relative abundance measured in the supernatant. When we repeated these experiments with ^15^N-DpppGpp (**14**), we found that pppGpp-to-ppGpp conversion was indeed markedly faster in the pellet fraction, whereas extracellular conversion was slower (Figure 5E).

Next, we investigated whether a drug efflux system could be responsible for MSN leakage into the medium.^[82–84]^ Therefore, we compared knockout strains lacking the major drug efflux components AcrA, AcrB and TolC with the wild type.^[80]^ In a simple experiment, we incubated the cell suspensions with DpppGpp/SNT and performed four washing steps where we centrifuged the cell suspension, removed the supernatant and resuspended the pellet. We then analyzed the washing solution for DEACBY fluorescence to measure DpppGpp leakage. Even after four washing steps, significant amounts of DpppGpp could still be detected in the washing solution of all the strains investigated (Figure S9). This finding suggests that SNT-induced membrane perturbation, as opposed to active export of internalized caged MSN, serve as the predominant contributors to the observed phenomenon.

Taken together, these data suggest that the extracellular MSN pool arises largely from membrane damage, which releases (p)ppGpp into the medium. At the same time, rapid intracellular turnover depletes the internal MSN pool, leading to lower intracellular levels relative to extracellular ones. Consequently, while CE-MS provides robust tracking of MSN photo-release and metabolism, careful interpretation is required, as a portion of the detected signal may not reflect true intracellular pools. Notwithstanding, confocal microscopy unambiguously showed internalization of the probes into a fraction of cells. Taken together, our current interpretation is that part of the probe is released inside of the cells, while another fraction is leaking from cells due to SNT action.

### Bacterial Growth Assay

The bacterial growth rate is apparently regulated by MSN.^[85–87]^ As simple read-out we envisioned that photo-release of caged MSN should slow down bacterial growth. Therefore, the growth of *E. coli* was monitored after incubation with the D(p)ppGpp/SNT mixture with and without following photo-release.

For wild type *E. coli*, we could not detect this projected difference in growth rate. As shown in the confocal microscopy experiments described above, uptake of the caged MSNs was heterogeneous. We therefore attribute the absence of a measurable population-level effect to this cell-to-cell variability: while some cells internalized high amounts of D(p)ppGpp, others likely contained only minimal levels, resulting in no apparent change in overall growth.

To overcome this limitation, we switched to a mutant strain incapable of synthesizing MSNs ((p)ppGpp^0^) due to deletion of the relA and spoT genes. As basal levels of MSNs are important to maintain cellular component homeostasis, this mutant has multiple disabilities in cell division, transcription and translation and can have difficulties to adapt to environmental changes.^[4,6]^ Due to the absence of MSNs in this mutant, the influence of artificially added MSNs on the growth rate should be pronounced, but in this case potentially increase the growth rate due to compensation of above-mentioned defects. We observed that treatment with SNT alone decreases the bacterial growth rate, likely due to its toxicity by damaging the bacterial envelope. To evaluate the influence of DppGpp and DpppGpp mixed with SNT on the growth rate of the *E. coli* (p)ppGpp^0^ strain, we used D-ATP (**S9**) mixed with SNT as a control. Interestingly, incubation of (p)ppGpp^0^ with SNT and caged (p)ppGpp already recovered the loss in growth rate caused by SNT. Caged ATP did not show this effect (Figure 6A). One possible conclusion is that caged MSNs, due to their versatile binding patterns,^[88]^ can already interact with some of the binding partners of MSN and thus restore growth. In line with this interpretation, uncaging of (p)ppGpp did not further affect the recovery of the growth rate (Figure 6B). This interpretation is in line with previous findings for caged molecules.^[63,89,90]^ Future studies will now address the increase of bulkiness in the cage through click chemistry on the alkyne handle to abrogate unwanted binding prior to uncaging.^[91]^

**Figure 6.**
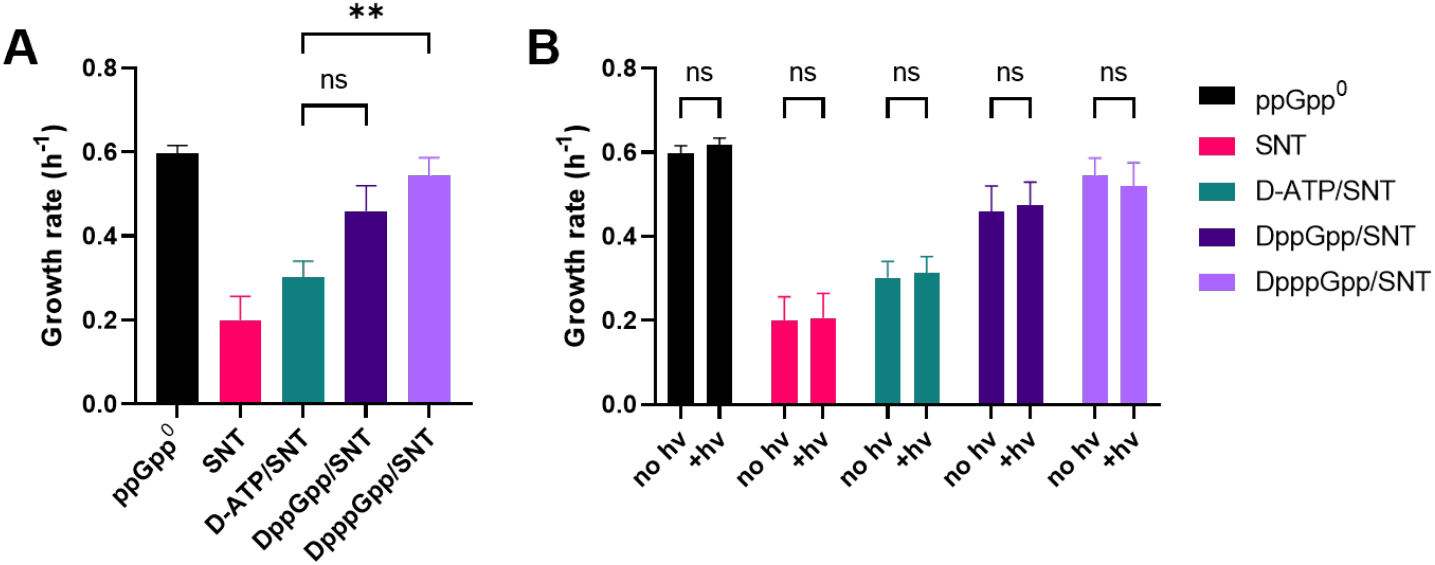
Growth rate of the E. coli (p)ppGpp^0^ mutant treated with SNT and different nucleotides. (A) Comparison of the different treatment conditions without light irradiation. Analyzed by Brown-Forsythe and Welch ANOVA test with Dunnett′s T3 multiple comparisons test; individual variances computed for each comparison. ns: p =0.125; **: p = 0.0054. (B) Comparison of samples irradiated or not irradiated with 400 nm light. Analyzed with two-way ANOVA with Šídák multiple comparisons test, with a single pooled variance. ns: p > 0.99. Mean and standard error of mean of at least four separate experiments is shown.

### Summary and Conclusion

In this study, we present the first photocaged derivatives of the MSNs pppGpp and ppGpp. DppGpp (**16**), DpppGpp (**13**), and the heavy isotope labeled ^15^N-DpppGpp (**14**) were efficiently synthesized in high yields using a chemoenzymatic synthesis approach. By comparing different delivery strategies, we achieve the delivery of highly negatively charged, caged MSNs through the complex cell envelope of living *E. coli*. Several analytical approaches combined provide evidence for intracellular delivery into a subpopulation of cells. Upon 400 nm irradiation, these probes release free nucleotides, enabling tracking of pppGpp-to-ppGpp conversion by CE-MS. Moreover, initial growth curve studies in wild type and (p)ppGpp^0^ strains demonstrate the potential of these probes for *in vivo* studies after further optimization: While photolysis and metabolic conversion were clearly demonstrated, extraction experiments revealed substantial leakage of MSNs into the extracellular medium, complicating accurate quantification of intracellular levels. Growth studies with a (p)ppGpp0 strain confirmed that caged MSNs can modulate bacterial physiology, but no light-dependent growth effect was yet observed, likely due to residual activity of the caged species.

Future work should prioritize improved transporter strategies and the development of bulkier, biologically inert cages. Single-cell studies of cells that have received large amounts of caged nucleotide could help attribute heterogeneous uptake to specific cellular states. Despite its current limitations, this study provides first significant strategies how to deliver highly charged signaling molecules into living bacteria and control cellular concentrations on a minute time-scale through photo-uncaging.

## Acknowledgement

We thank Dr. Guizhen Liu, Dr. Mengsi Lu and Dr. Anuj Shukla for CE-MS measurements. Furthermore, we thank the staff of the Life Imaging Center (LIC) of the University of Freiburg, in particular I. Bierschenk, A. Neumann and R. Nitschke for help with their microscopy resources, and the excellent support in image recording and analysis. The following instrument funded by the Deutsche Forschungsgemeinschaft (DFG, German Research Foundation) was used: ZEISS LSM 880 Observer / Fast Airyscan with inverted microscope (Instrument Grant Number 317784314). In addition, we thank D. Herchenbach and M. Follo of the Lighthouse Core Facility, funded in part by the Medical Faculty, University of Freiburg (Project Number 2023/B3-Fol). Hans-Georg Koch acknowledges funding by the SFB1381 (Project-ID 403222702). This project has received funding from the European Research Council (ERC) under the European Union’s Horizon 2020 research and innovation program (grant agreement no. 864246, to H.J.J.). This study was supported by the Deutsche Forschungsgemeinschaft (DFG) under Germany’s excellence strategy (CIBSS, EXC-2189, Project ID 390939984, to H. J. J)

## Author Contributions

Christoph Popp: Synthesis, Investigation, Formal analysis, Visualization, Writing; Patrick Moser: Synthesis of ^15^N-DpppGpp (14); Xinwei Liu: Supervision and discussion of *in vivo* experiments; Johannes Freitag: Supervision and discussion of *in vivo* experiments; Isabel Prucker: Discussion of CE-MS results; Robert Zscherp: Synthesis of S11; Xuan Wang: Synthesis of S13; Philipp Klahn: Supervision and discussion ; Hans-Georg Koch: Supervision and discussion; Gert Bange: Supervision and discussion of *in vivo* experiments; Henning J. Jessen: Conceptualization, Supervision, Funding acquisition, Writing – review & editing

## Bibliography

[1] G. Bange, D. E. Brodersen, A. Liuzzi, W. Steinchen, “Two P or Not Two P: Understanding Regulation by the Bacterial Second Messengers (p)ppGpp” Annu Rev Microbiol 2021, 75, 383–406.

[2] O. Pacios, L. Blasco, I. Bleriot, L. Fernandez-Garcia, A. Ambroa, M. López, G. Bou, R. Cantón, R. Garcia-Contreras, T. K. Wood, M. Tomás, “(p)ppGpp and Its Role in Bacterial Persistence: New Challenges” Antimicrob Agents Chemother 2020, 64, 1–14.

[3] S. E. Irving, N. R. Choudhury, R. M. Corrigan, “The stringent response and physiological roles of (pp)pGpp in bacteria” Nat Rev Microbiol 2021, 19, 256–271.

[4] W. Steinchen, V. Zegarra, G. Bange, “(p)ppGpp: Magic Modulators of Bacterial Physiology and Metabolism” Front Microbiol 2020, 11, 2072.

[5] M. Cashel, J. Gallant, “Two Compounds implicated in the Function of the RC Gene of Escherichia coli” Nature 1969, 221, 838–841.

[6] L. Fernández-Coll, M. Cashel, “Possible Roles for Basal Levels of (p)ppGpp: Growth Efficiency Vs. Surviving Stress” Front Microbiol 2020, 11, 592718.

[7] S. Han, J. Min, Y. Park, W. Park, “Fine-tuning regulation of (p)ppGpp-driven outer membrane vesicle formation in Acinetobacter baumannii” FEBS J 2025, 292, 3696– 3717.

[8] D. K. Fung, J. T. Barra, J. Yang, J. W. Schroeder, F. She, M. Young, D. Ying, D. M. Stevenson, D. Amador-Noguez, J. D. Wang, “A shared alarmone–GTP switch controls persister formation in bacteria” Nat Microbiol 2025, 10, 1617–1629.

[9] M. Zhu, H. Mu, X. Dai, “Integrated control of bacterial growth and stress response by (p)ppGpp in Escherichia coli: A seesaw fashion” iScience 2024, 27, 108818.

[10] U. Mechold, K. Potrykus, H. Murphy, K. S. Murakami, M. Cashel, “Differential regulation by ppGpp versus pppGpp in Escherichia coli” Nucleic Acids Res 2013, 41, 6175–6189.

[11] V. Shyp, S. Tankov, A. Ermakov, P. Kudrin, B. P. English, M. Ehrenberg, T. Tenson, J. Elf, V. Hauryliuk, “Positive allosteric feedback regulation of the stringent response enzyme RelA by its product” EMBO Rep 2012, 13, 835–839.

[12] P. Kudrin, I. Dzhygyr, K. Ishiguro, J. Beljantseva, E. Maksimova, S. R. A. Oliveira, V. Varik, R. Payoe, A. L. Konevega, T. Tenson, T. Suzuki, V. Hauryliuk, “The ribosomal A-site finger is crucial for binding and activation of the stringent factor RelA” Nucleic Acids Res 2018, 46, 1973–1983.

[13] W. Steinchen, J. S. Schuhmacher, F. Altegoer, C. D. Fage, V. Srinivasan, U. Linne, M. A. Marahiel, G. Bange, “Catalytic mechanism and allosteric regulation of an oligomeric (p)ppGpp synthetase by an alarmone” Proc Natl Acad Sci USA 2015, 112, 13348–13353.

[14] B. W. Anderson, D. K. Fung, J. D. Wang, “Regulatory Themes and Variations by the Stress-Signaling Nucleotide Alarmones (p)ppGpp in Bacteria” Annu Rev Genet 2021, 55, 115–133.

[15] M. Roghanian, K. Van Nerom, H. Takada, J. Caballero-Montes, H. Tamman, P. Kudrin, A. Talavera, I. Dzhygyr, S. Ekström, G. C. Atkinson, A. Garcia-Pino, V. Hauryliuk, “(p)ppGpp controls stringent factors by exploiting antagonistic allosteric coupling between catalytic domains” Mol Cell 2021, 81, 3310-3322.e6.

[16] T. J. Silhavy, D. Kahne, S. Walker, “The Bacterial Cell Envelope” Cold Spring Harb Perspect Biol 2010, 2, a000414.

[17] G. K. Auer, D. B. Weibel, “Bacterial Cell Mechanics” Biochemistry 2017, 56, 3710– 3724.

[18] V. T. Sterrenberg, D. Stalling, J. I. H. Knaack, T. K. Soh, J. B. Bosse, C. Meier, “A Tri PPP ro-Nucleotide Reporter with Optimized Cell-Permeable Dyes for Metabolic Labeling of Cellular and Viral DNA in Living Cells” Angew Chem Int Ed 2023, 62, e202308271.

[19] T. Gollnest, T. D. de Oliveira, D. Schols, J. Balzarini, C. Meier, “Lipophilic prodrugs of nucleoside triphosphates as biochemical probes and potential antivirals” Nat Commun 2015, 6, 8716.

[20] T. Bittner, C. Wittwer, S. Hauke, D. Wohlwend, S. Mundinger, A. K. Dutta, D. Bezold, T. Dürr, T. Friedrich, C. Schultz, H. J. Jessen, “Photolysis of Caged Inositol Pyrophosphate InsP 8 Directly Modulates Intracellular Ca 2+ Oscillations and Controls C2AB Domain Localization” J Am Chem Soc 2020, 142, 10606–10611.

[21] C. McGuigan, M. Derudas, B. Gonczy, K. Hinsinger, S. Kandil, F. Pertusati, M. Serpi, R. Snoeck, G. Andrei, J. Balzarini, T. D. McHugh, A. Maitra, E. Akorli, D. Evangelopoulos, S. Bhakta, “ProTides of N-(3-(5-(2′-deoxyuridine))prop-2-ynyl)octanamide as potential anti-tubercular and anti-viral agents” Bioorg Med Chem 2014, 22, 2816–2824.

[22] S. V Vinogradov, A. D. Zeman, E. V Batrakova, A. V Kabanov, “Polyplex Nanogel formulations for drug delivery of cytotoxic nucleoside analogs.” J Control Release 2005, 107, 143–57.

[23] I. Pavlovic, D. T. Thakor, J. R. Vargas, C. J. McKinlay, S. Hauke, P. Anstaett, R. C. Camuña, L. Bigler, G. Gasser, C. Schultz, P. A. Wender, H. J. Jessen, “Cellular delivery and photochemical release of a caged inositol-pyrophosphate induces PH-domain translocation in cellulo” Nat Commun 2016, 7, 10622.

[24] J. Ma, J. Wehrle, D. Frank, L. Lorenzen, C. Popp, W. Driever, R. Grosse, H. J. Jessen, “Intracellular delivery and deep tissue penetration of nucleoside triphosphates using photocleavable covalently bound dendritic polycations” Chem Sci 2024, 15, 6478– 6487.

[25] A. E. Fouda, M. K. H. Pflum, “A Cell-Permeable ATP Analogue for Kinase-Catalyzed Biotinylation” Angew Chem Int Ed 2015, 54, 9618–9621.

[26] A. Maya-Mendoza, P. Olivares-Chauvet, F. Kohlmeier, D. A. Jackson, “Visualising chromosomal replication sites and replicons in mammalian cells” Methods 2012, 57, 140–148.

[27] K. Koberna, D. Staněk, J. Malínský, M. Eltsov, A. Pliss, V. Čtrnáctá, Š. Cermanová, I. Raška, “Nuclear organization studied with the help of a hypotonic shift: Its use permits hydrophilic molecules to enter into living cells” Chromosoma 1999, 108, 325–335.

[28] S. M. Hacker, A. Buntz, A. Zumbusch, A. Marx, “Direct Monitoring of Nucleotide Turnover in Human Cell Extracts and Cells by Fluorogenic ATP Analogs” ACS Chem Biol 2015, 10, 2544–2552.

[29] E. M. M. Manders, H. Kimura, P. R. Cook, “Direct Imaging of DNA in Living Cells Reveals the Dynamics of Chromosome Formation” J Cell Biol 1999, 144, 813–822.

[30] D. Zink, T. Cremer, R. Saffrich, R. Fischer, M. F. Trendelenburg, W. Ansorge, E. H. K. Stelzer, “Structure and dynamics of human interphase chromosome territories in vivo” Hum Genet 1998, 102, 241–251.

[31] D. A. Malyshev, K. Dhami, T. Lavergne, T. Chen, N. Dai, J. M. Foster, I. R. Corrêa, F. E. Romesberg, “A semi-synthetic organism with an expanded genetic alphabet” Nature 2014, 509, 385–388.

[32] Y. Zhang, B. M. Lamb, A. W. Feldman, A. X. Zhou, T. Lavergne, L. Li, F. E. Romesberg, “A semisynthetic organism engineered for the stable expansion of the genetic alphabet” Proc Natl Acad Sci USA 2017, 114, 1317–1322.

[33] H. K. Lembke, A. Espinasse, M. G. Hanson, C. J. Grimme, Z. Tan, T. M. Reineke, E. E. Carlson, “Cationic Polymers Enable Internalization of Negatively Charged Chemical Probes into Bacteria” ACS Chem Biol 2023, 18, 2063–2072.

[34] Z. Zawada, A. Tatar, P. Mocilac, M. Buděšínský, T. Kraus, “Transport of Nucleoside Triphosphates into Cells by Artificial Molecular Transporters” Angew Chem Int Ed 2018, 57, 9891–9895.

[35] J. L. Young, D. A. Dean in Adv Genet, Academic Press, 2015, pp. 49–88.

[36] S. N. Cohen, A. C. Y. Chang, L. Hsu, “Nonchromosomal Antibiotic Resistance in Bacteria: Genetic Transformation of Escherichia coli by R-Factor DNA” Proc Natl Acad Sci USA 1972, 69, 2110–2114.

[37] M. Mandel, A. Higa, “Calcium-dependent bacteriophage DNA infection” J Mol Biol 1970, 53, 159–162.

[38] D. Hanahan, “Studies on transformation of Escherichia coli with plasmids” J Mol Biol 1983, 166, 557–580.

[39] M. M. Islam, M. Odahara, T. Yoshizumi, K. Oikawa, M. Kimura, M. Su’etsugu, K. Numata, “Cell-Penetrating Peptide-Mediated Transformation of Large Plasmid DNA into Escherichia coli” ACS Synth Biol 2019, 8, 1215–1218.

[40] V. V. de Souza, P. A. M. Vitale, F. H. Florenzano, R. K. Salinas, I. M. Cuccovia, “A novel method for DNA delivery into bacteria using cationic copolymers” Braz J Med Biol Res 2021, 54, e10743.

[41] R. van der Meel, P. A. Wender, O. M. Merkel, I. Lostalé-Seijo, J. Montenegro, A. Miserez, Q. Laurent, H. Sleiman, P. Luciani, “Next-generation materials for nucleic acid delivery” Nat Rev Mater 2025, 10, 490–499.

[42] A. Nazli, D. L. He, D. Liao, M. Z. I. Khan, C. Huang, Y. He, “Strategies and progresses for enhancing targeted antibiotic delivery” Adv Drug Deliv Rev 2022, 189, 114502.

[43] A. V. Cheng, W. M. Wuest, “Signed, Sealed, Delivered: Conjugate and Prodrug Strategies as Targeted Delivery Vectors for Antibiotics” ACS Infect Dis 2019, 5, 816– 828.

[44] J. Kramer, Ö. Özkaya, R. Kümmerli, “Bacterial siderophores in community and host interactions” Nat Rev Microbiol 2020, 18, 152–163.

[45] P. Klahn, M. Brönstrup, “Bifunctional antimicrobial conjugates and hybrid antimicrobials” Nat Prod Rep 2017, 34, 832–885.

[46] M. J. Pals, L. Wijnberg, Ç. Yildiz, W. A. Velema, “Catechol-Siderophore Mimics Convey Nucleic Acid Therapeutics into Bacteria” Angew Chem Int Ed 2024, 63, e202402405.

[47] R. Zscherp, J. Coetzee, J. Vornweg, J. Grunenberg, J. Herrmann, R. Müller, P. Klahn, “Biomimetic enterobactin analogue mediates iron-uptake and cargo transport into E. coli and P. aeruginosa” Chem Sci 2021, 12, 10179–10190.

[48] J. H. Boyce, B. Dang, B. Ary, Q. Edmondson, C. S. Craik, W. F. DeGrado, I. B. Seiple, “Platform to Discover Protease-Activated Antibiotics and Application to Siderophore– Antibiotic Conjugates” J Am Chem Soc 2020, 142, 21310–21321.

[49] M. Madaoui, O. Vidal, A. Meyer, M. Noël, J. Lacroix, J. Vasseur, A. Marra, F. Morvan, “Modified Galacto-or Fuco-Clusters Exploiting the Siderophore Pathway to Inhibit the LecA-or LecB-Associated Virulence of Pseudomonas aeruginosa” ChemBioChem 2020, 21, 3433–3448.

[50] T. Zheng, J. L. Bullock, E. M. Nolan, “Siderophore-Mediated Cargo Delivery to the Cytoplasm of Escherichia coli and Pseudomonas aeruginosa: Syntheses of Monofunctionalized Enterobactin Scaffolds and Evaluation of Enterobactin–Cargo Conjugate Uptake” J Am Chem Soc 2012, 134, 18388–18400.

[51] X. Ning, S. Lee, Z. Wang, D. Kim, B. Stubblefield, E. Gilbert, N. Murthy, “Maltodextrin-based imaging probes detect bacteria in vivo with high sensitivity and specificity” Nat Mater 2011, 10, 602–607.

[52] A. Zlitni, G. Gowrishankar, I. Steinberg, T. Haywood, S. Sam Gambhir, “Maltotriose-based probes for fluorescence and photoacoustic imaging of bacterial infections” Nat Commun 2020, 11, 1250.

[53] R. Guo, K. Li, J. Qin, S. Niu, W. Hong, “Development of polycationic micelles as an efficient delivery system of antibiotics for overcoming the biological barriers to reverse multidrug resistance in Escherichia coli” Nanoscale 2020, 12, 11251–11266.

[54] H.-M. Lee, J. Ren, K. M. Tran, B.-M. Jeon, W.-U. Park, H. Kim, K. E. Lee, Y. Oh, M. Choi, D.-S. Kim, D. Na, “Identification of efficient prokaryotic cell-penetrating peptides with applications in bacterial biotechnology” Commun Biol 2021, 4, 205.

[55] G. Inoue, D. Toyohara, T. Mori, T. Muraoka, “Critical Side Chain Effects of Cell-Penetrating Peptides for Transporting Oligo Peptide Nucleic Acids in Bacteria” ACS Appl Bio Mater 2021, 4, 3462–3468.

[56] A. Schmitt, H. Wennemers, “Amphipathic Proline-Rich Cell Penetrating Peptides for Targeting Mitochondria” ACS Chem Biol 2025, 20, 2298–2307.

[57] A. Barba-Bon, G. Salluce, I. Lostalé-Seijo, K. I. Assaf, A. Hennig, J. Montenegro, W. M. Nau, “Boron clusters as broadband membrane carriers” Nature 2022, 603, 637–642.

[58] Q. Laurent, R. Martinent, B. Lim, A.-T. Pham, T. Kato, J. López-Andarias, N. Sakai, S. Matile, “Thiol-Mediated Uptake” JACS Au 2021, 1, 710–728.

[59] I. S. Shchelik, K. Gademann, “Thiol- and Disulfide-Containing Vancomycin Derivatives Against Bacterial Resistance and Biofilm Formation” ACS Med Chem Lett 2021, 12, 1898–1904.

[60] G. C. R. Ellis-Davies, “Caged compounds: photorelease technology for control of cellular chemistry and physiology” Nat Methods 2007, 4, 619–628.

[61] Y. Li, M. Wang, F. Wang, S. Lu, X. Chen, “Recent progress in studies of photocages” Smart Molecules 2023, 1, e20220003.

[62] R. Weinstain, T. Slanina, D. Kand, P. Klán, “Visible-to-NIR-Light Activated Release: From Small Molecules to Nanomaterials” Chem Rev 2020, 120, 13135–13272.

[63] G. C. R. Ellis-Davies, “Reverse Engineering Caged Compounds: Design Principles for their Application in Biology” Angew Chem Int Ed 2023, 62, e202206083.

[64] J. H. Kaplan, B. Forbush, J. F. Hoffman, “Rapid photolytic release of adenosine 5’-triphosphate from a protected analog: utilization by the sodium:potassium pump of human red blood cell ghosts” Biochemistry 1978, 17, 1929–1935.

[65] M. J. Hansen, W. A. Velema, M. M. Lerch, W. Szymanski, B. L. Feringa, “Wavelength-selective cleavage of photoprotecting groups: strategies and applications in dynamic systems” Chem Soc Rev 2015, 44, 3358–3377.

[66] P. Klán, T. Šolomek, C. G. Bochet, A. Blanc, R. Givens, M. Rubina, V. Popik, A. Kostikov, J. Wirz, “Photoremovable protecting groups in chemistry and biology: Reaction mechanisms and efficacy” Chem Rev 2013, 113, 119–191.

[67] T. M. Haas, D. Qiu, M. Häner, L. Angebauer, A. Ripp, J. Singh, H. Koch, C. Jessen-Trefzer, H. J. Jessen, “Four Phosphates at One Blow: Access to Pentaphosphorylated Magic Spot Nucleotides and Their Analysis by Capillary Electrophoresis” J Org Chem 2020, 85, 14496–14506.

[68] T. M. Haas, B. Laventie, S. Lagies, C. Harter, I. Prucker, D. Ritz, R. Saleem-Batcha, D. Qiu, W. Hüttel, J. Andexer, B. Kammerer, U. Jenal, H. J. Jessen, “Photoaffinity Capture Compounds to Profile the Magic Spot Nucleotide Interactomes**” Angew Chem Int Ed 2022, 61, e202201731.

[69] D. Qiu, E. Lange, T. M. Haas, I. Prucker, S. Masuda, Y. L. Wang, G. Felix, G. Schaaf, H. J. Jessen, “Bacterial Pathogen Infection Triggers Magic Spot Nucleotide Signaling in Arabidopsis thaliana Chloroplasts through Specific RelA/SpoT Homologues” J Am Chem Soc 2023, 145, 16081–16089.

[70] T. M. Haas, P. Ebensperger, V. B. Eisenbeis, C. Nopper, T. Dürr, N. Jork, N. Steck, C. Jessen-Trefzer, H. J. Jessen, “Magic spot nucleotides: tunable target-specific chemoenzymatic synthesis” Chem Commun 2019, 55, 5339–5342.

[71] G. S. Cremosnik, A. Hofer, H. J. Jessen, “Iterative Synthesis of Nucleoside Oligophosphates with Phosphoramidites” Angew Chem Int Ed 2014, 53, 286–289.

[72] P. Seyfried, L. Eiden, N. Grebenovsky, G. Mayer, A. Heckel, “Photo-Tethers for the (Multi-)Cyclic, Conformational Caging of Long Oligonucleotides” Angew Chem Int Ed 2017, 56, 359–363.

[73] V. V Rostovtsev, L. G. Green, V. V Fokin, K. B. Sharpless, “A Stepwise Huisgen Cycloaddition Process: Copper(I)-Catalyzed Regioselective ‘Ligation’ of Azides and Terminal Alkynes” Angew Chem Int Ed 2002, 41, 2596–2599.

[74] C. W. Tornøe, C. Christensen, M. Meldal, “Peptidotriazoles on Solid Phase: [1,2,3]-Triazoles by Regiospecific Copper(I)-Catalyzed 1,3-Dipolar Cycloadditions of Terminal Alkynes to Azides” J Org Chem 2002, 67, 3057–3064.

[75] H. J. Jessen, T. Dürr-Mayer, T. M. Haas, A. Ripp, C. C. Cummins, “Lost in Condensation: Poly-, Cyclo-, and Ultraphosphates” Acc Chem Res 2021, 54, 4036– 4050.

[76] R. Zscherp, J. Coetzee, J. Vornweg, J. Grunenberg, J. Herrmann, R. Müller, P. Klahn, “Biomimetic enterobactin analogue mediates iron-uptake and cargo transport into E. coli and P. aeruginosa” Chem Sci 2021, 12, 10179–10190.

[77] D. Qiu, M. S. Wilson, V. B. Eisenbeis, R. K. Harmel, E. Riemer, T. M. Haas, C. Wittwer, N. Jork, C. Gu, S. B. Shears, G. Schaaf, B. Kammerer, D. Fiedler, A. Saiardi, H. J. Jessen, “Analysis of inositol phosphate metabolism by capillary electrophoresis electrospray ionization mass spectrometry” Nat Commun 2020, 11, 6035.

[78] A. Hara, J. Sy, “Guanosine 5’-triphosphate, 3’-diphosphate 5’-phosphohydrolase. Purification and substrate specificity.” J Biol Chem 1983, 258, 1678–1683.

[79] C. R. Somerville, A. Ahmed, “Mutants of Escherichia coli defective in the degradation of guanosine 5’-triphosphate, 3’-diphosphate (pppGpp)” Mol Gen Genet 1979, 169, 315–323.

[80] T. Baba, T. Ara, M. Hasegawa, Y. Takai, Y. Okumura, M. Baba, K. A. Datsenko, M. Tomita, B. L. Wanner, H. Mori, “Construction of Escherichia coli K-12 in-frame, single-gene knockout mutants: the Keio collection” Mol Syst Biol 2006, 2, DOI 10.1038/msb4100050.

[81] Chung Nan Chang, K. Wun-Jing, E. Y. Chen, “Nucleotide sequence of the alkaline phosphatase gene of Escherichia coli” Gene 1986, 44, 121–125.

[82] D. Du, X. Wang-Kan, A. Neuberger, H. W. van Veen, K. M. Pos, L. J. V. Piddock, B. F. Luisi, “Multidrug efflux pumps: structure, function and regulation” Nat Rev Microbiol 2018, 16, 523–539.

[83] K. Nishino, S. Yamasaki, R. Nakashima, M. Zwama, M. Hayashi-Nishino, “Function and Inhibitory Mechanisms of Multidrug Efflux Pumps” Front Microbiol 2021, 12, DOI 10.3389/fmicb.2021.737288.

[84] K. M. Pos, “Drug transport mechanism of the AcrB efflux pump” Biochim Biophys Acta 2009, 1794, 782–793.

[85] K. Potrykus, H. Murphy, N. Philippe, M. Cashel, “ppGpp is the major source of growth rate control in E. coli” Environ Microbiol 2011, 13, 563–575.

[86] N. C. E. Imholz, M. J. Noga, N. J. F. van den Broek, G. Bokinsky, “Calibrating the Bacterial Growth Rate Speedometer: A Re-evaluation of the Relationship Between Basal ppGpp, Growth, and RNA Synthesis in Escherichia coli” Front Microbiol 2020, 11, 574872.

[87] M. Zhu, X. Dai, “Growth suppression by altered (p)ppGpp levels results from non-optimal resource allocation in Escherichia coli” Nucleic Acids Res 2019, 47, 4684– 4693.

[88] G. S. Kushwaha, A. Patra, N. S. Bhavesh, “Structural Analysis of (p)ppGpp Reveals Its Versatile Binding Pattern for Diverse Types of Target Proteins” Front Microbiol 2020, 11, 575041.

[89] H. Thirlwell, J. A. Sleep, M. A. Ferenczi, “Inhibition of unloaded shortening velocity in permeabilized muscle fibres by caged ATP compounds” J Muscle Res Cell Motil 1995, 16, 131–137.

[90] S. Geibel, A. Barth, S. Amslinger, A. H. Jung, C. Burzik, R. J. Clarke, R. S. Givens, K. Fendler, “P3-[2-(4-hydroxyphenyl)-2-oxo]ethyl ATP for the Rapid Activation of the Na+,K+-ATPase” Biophys J 2000, 79, 1346–1357.

[91] R. Durand-de Cuttoli, P. S. Chauhan, A. Pétriz Reyes, P. Faure, A. Mourot, G. C. R. Ellis-Davies, “Optofluidic control of rodent learning using cloaked caged glutamate” Proc Natl Acad Sci USA 2020, 117, 6831–6835.

