## supplementary information for "Controlled Delivery and Light-Induced Release of Magic Spot Nucleotides in *Escherichia coli*"

#### Abbreviations

| | |
|---|---|
| DBU | 1,8-Diazabicyclo[5.4.0]undec-7-ene |
| DMF | Dimethylformamide |
| DMSO | Dimethyl sulfoxide |
| eq | Equivalents |
| ETT | 5-(Ethylthio)-1*H*-tetrazole |
| HPLC | High-performance liquid chromatography |
| HRMS | High resolution mass spectrometry |
| *m*CPBA | meta-Chloroperoxybenzoic acid |
| MSN | Magic spot nucleotide |
| NMR | Nuclear magnetic resonance |
| P-amidite | Phosphoramidite |
| pGp | Guanosine-3’,5’-bisphosphate |
| ppGp | Guanosine-3’-phosphate-5’-diphosphate |
| ppGpp | Guanosine-3’,5’-bisdiphosphate |
| pppGpp | Guanosine-3’-diphosphate-5’-triphosphate |
| SAX | Strong anion exchange |
| TBA | Tetrabutylammonium |
| TEAA | Triethylammonium acetate |
| TLC | Thin Layer Chromatography |

### 1 Materials and General Information

**General:** Chemicals were purchased from Carl Roth GmBH & Co. KG, Merck KGaA, TCI Chemicals GmbH, VWR International and Acros Organics or obtained from previously synthesized stocks available in our laboratory and were used without further purification. Carbohydrate azides (except trehalose) were obtained from Synthose Inc. Ultrapure water from a Mili-Q Reference water purification system (Merck KGaA) was always used for experiments. Dry solvents were always used under argon atmosphere in flame-dried glassware. Phosphoramidites and nucleotides were stored at -20°C. Concentration of photocaged MSN stock solutions was determined by absorption and purity was regularly verified by analytical HPLC.

**Thin Layer Chromatography (TLC)** was performed on aluminum plates pre-coated with silica gel $60F_{254}$ (*Merck*), which were visualized by UV fluorescence (λ = 254 nm, 366 nm). TLC was used to verify purification results and to monitor reactions. (TLC, Silica gel 60G F254 on aluminum foil, Merck KGaA).

**Nuclear Magnetic Resonance (NMR)** spectra were acquired on a *Bruker Avance 400* (1H-NMR: 400 MHz, $^{13}$C-NMR: 101 MHz, $^{31}$P-NMR: 162 MHz). Data for $^{1}$H and $^{31}$P NMR spectra is reported as follows: chemical shift (δ ppm), multiplicity (s = singlet, d = doublet, t = triplet, q = quartet, m = multiplet), coupling constant (Hz), integration. All $^{13}$C and $^{31}$P NMR spectra were recorded with proton decoupling. Data for $^{13}$C NMR spectra is reported by chemical shift (δ ppm). Where coupling with phosphor or nitrogen nuclei was observed, multiplicity and coupling constants are reported. Peak shifts were reported relative to solvent peaks according to Fulmer *et al.*[1]

**HRMS** (High resolution mass spectrometry) was performed by the analytical department of the Institute of Organic Chemistry at the University of Freiburg on a *Thermo Scientific Exactive* and a *Agilent 6546 LC/Q-TOF* mass spectrometer.

**Strong ion-exchange chromatography** was performed using *ÄKTA pure*™ system and a *HiTrap*™ *Q HP, Capto*™ *Q ImpRes* or *Q FF* column from Cytiva. Crude products were loaded as aqueous solutions and eluted by increasing the concentration of $NaClO_4$ (anion exchange, 1 M). UV-absorption at λ = 400 nm was used for fraction detection.

**Analytical HPLC** was performed with a *Dionex UltiMate 3000* system of Thermo Fisher using a *ProntoSIL C18 AQ* column (120 Å, 3 µm, 150 x 3.0 mm). We used a gradient of acetonitrile in water with a constant triethylamine acetate (TEAA) buffer concentration (10 mM, pH 7.0) Analytes were detected at $\lambda$ =254 and 400 nm.

**Photolysis LEDs:** High power LED collimator source LCS-0400-03-48 from Mightex Systems.

- Nominal Wavelength: 400 nm
- Aperture: 48 mm
- Half diverging angle: 0.75°
- Typical output power 265 mW

Measured data:

- At 4 cm distance in center: ~18.0 mW $cm^{-2}$
- $\lambda_{max}$= 396 nm
- FWHM = 13 nm

For the growth assay we used a Darkbeam 10W 395 nm LED ($\lambda_{max}$= 396 nm, FWHM = 13 nm).

Instruments and material related to cell culture experiments are described in section 4.1.

### 2 Chemical Synthesis

#### 2.1 Precursor

##### Ent-$N_3$ (S11)

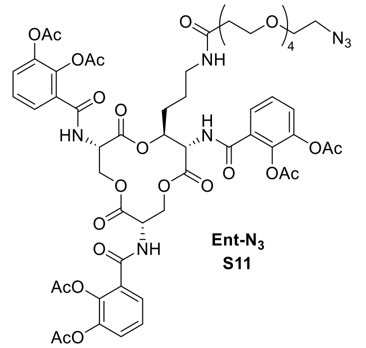

The biomimetic enterobactin analogue Ent-$N_3$ (S11) was synthesized by Robert Zscherp as previously published.[2]

##### DEACBY-OH (S12)

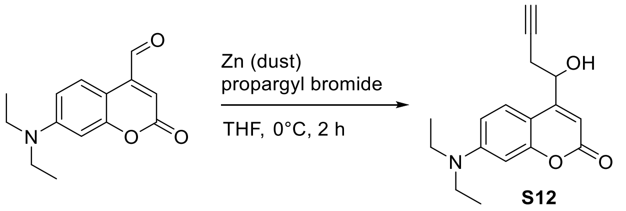

7-(diethylamino)-4-(1-hydroxybut-3-yn-1-yl)-coumarin (**DEACBY-OH, S12**) was synthesized according to a modified procedure adapted from Seyfried *et al.*[3]

In a 50 mL Schlenk flask zinc dust (<10 µm, 8.00 g, 122 mmol, 6 eq) was heated under vacuum using a "heat gun" for several minutes, suspended in dry THF (60 mL) and propargyl bromide (6.06 g, 51.0 mmol, 2.5 eq) was added at 0 °C. The reaction mixture was stirred at 0 °C for 1 h and then 7-(diethylamino)-coumarin-4-carbaldehyde (5.00 g, 20.4 mmol) was added in one portion. After 1.5 h the reaction was stopped by addition of 15 mL saturated $NH_4Cl$ solution. The mixture was stirred for 5 min, excess zinc solid was filtrated washed with ethyl acetate and the solution was treated with 10% aqueous HCl solution until pH < 3. After 5 min sat. $NaHCO_3$ was added until pH 7. After extraction with ethyl acetate (3 x 20 mL) the combined organic layers were dried over $Na_2SO_4$, filtrated and the solvent was removed under reduced pressure. Purification by flash chromatography (10% acetone in dichloromethane with 1% triethylamine) yielded **S12** (5.0 g, 18 mmol, 86%) as an orange foam.

**$^1$H NMR** (300 MHz, $CDCl_3$) δ 7.39 (d, *J* = 9.0 Hz, 1H), 6.57 (dd, *J* = 9.0, 2.7 Hz, 1H), 6.52 (d, *J* = 2.6 Hz, 1H), 6.30 (d, *J* = 1.0 Hz, 1H), 5.17 – 5.09 (m, 1H), 3.41 (q, *J* = 7.1 Hz, 4H), 2.84 (ddd, *J* = 17.0, 4.2, 2.6 Hz, 1H), 2.64 (ddd, *J* = 17.0, 7.7, 2.6 Hz, 1H), 2.50 (d, *J* = 4.2 Hz, 1H), 2.17 (t, *J* = 2.6 Hz, 1H), 1.21 (t, *J* = 7.1 Hz, 6H).
**$^{13}$C NMR** (101 MHz, $CDCl_3$) δ 162.4, 156.7, 155.4, 150.6, 124.8, 108.7, 106.0, 106.0, 98.2, 79.4, 72.4, 68.1, 44.9, 27.9, 12.6.
**HRMS** ($ESI^+$) m/z for $[C_{17}H_{20}NO_3]^+$ ($M+H^+$): calcd. 286.1438, found 286.1436.

##### 6-azido-trehalose (S13)

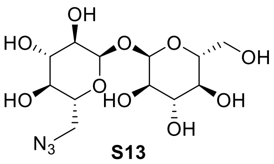

**S13**

6-azido-trehalose was synthesized by Xuan Wang as previously published.[4]

#### 2.2 Phosphor amidite synthesis

##### Synthesis of FmO-P-[N(*i*Pr)$_2$]$_2$ (S14)

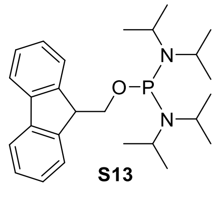

**S13**

The P-diamidite **S14** was synthesized as previously described. Analytical data in accordance with literature. Recrystallization from pentane yielded the crystalline solid in high purity.[5]

##### Synthesis of (FmO)$_2$-P-N(*i*Pr)$_2$ (5)

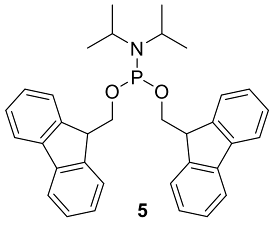

**5**

The P-amidite **5** was synthesized as previously reported in two steps from $PCl_3$. Analytical data in accordance with literature.[6]

#### Synthesis of (DEACBY)(FmO)-P-N(*i*Pr)$_2$ (10)

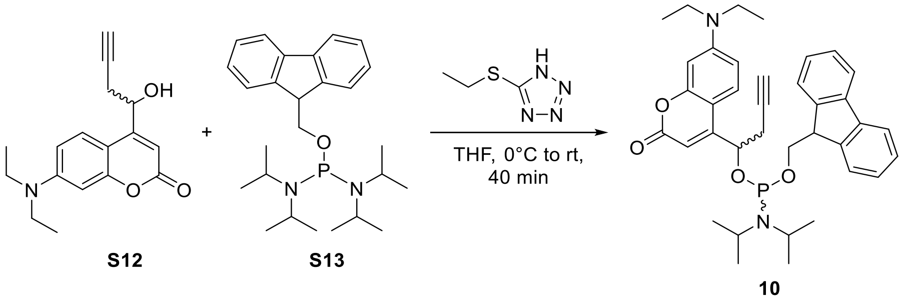

Compound **S12** (2.00 g, 7.01 mmol) and P-diamidite **S14** (3.29 g, 7.71 mmol, 1.1 eq) were coevaporated with dry acetonitrile (3 x 3 mL). The mixture was dissolved in dry THF (8 mL) and cooled to 0°C. 5-Ethylthio-1H-tetrazole stock solution (1 M in acetonitrile, 7.01 mL, 7.01 mmol, 1 eq) was added slowly over 5 min. After stirring for 10 min at room temperature dry diethyl ether was added (20 mL), formed precipitate was filtered off, 2 drops of triethylamine were added and the solvent was evaporated to dryness. The crude product was purified by flash column chromatography with deactivated silica (silica pretreated with 10% triehylamine in cyclohexane, 1 - 25% ethyl acetate in cyclohexane). **10** was isolated as an orange solid containing a 1:1 mixture of both diastereomers (3.89 g, 6.37 mmol, 91 %)

**$^{1}$H NMR** (400 MHz, $CDCl_3$, *mixture of diastereomers*) δ 7.82 – 7.78 (m, 2H), 7.78 – 7.66 (m, 4H), 7.53 – 7.30 (m, 10H), 7.24 – 7.14 (m, 2H), 6.58 – 6.51 (m, 4H), 6.28 (d, J = 0.8 Hz, 1H), 6.26 (d, J = 0.8 Hz, 1H), 5.10 – 4.96 (m, 2H), 4.26 (t, J = 6.4 Hz, 1H), 4.24 – 4.17 (m, 1H), 4.04 (t, J = 6.8 Hz, 1H), 3.99 – 3.90 (m, 2H), 3.78 – 3.55 (m, 5H), 3.48 – 3.39 (m, 8H), 2.82 – 2.61 (m, 4H), 2.04 (t, J = 2.6 Hz, 1H), 1.99 (t, J = 2.6 Hz, 1H), 1.28 (d, J = 6.8 Hz, 6H), 1.25 – 1.20 (m, 12H), 1.16 (d, J = 4.5 Hz, 6H), 1.14 (d, J = 4.6 Hz, 6H), 1.08 (d, J = 6.8 Hz, 6H).

**$^{31}$P NMR** (162 MHz, $CDCl_3$, *mixture of diastereomers*) δ 148.56 (s, 1P), 148.25 (s, 1P).

**$^{13}$C NMR** (101 MHz, $CDCl_3$, *mixture of diastereomers*) δ162.3, 156.8, 156.7, 155.0, 154.9 (d, *J* = 2.6 Hz), 150.5, 145.1, 144.7, 144.7, 144.5, 141.6, 141.5, 141.4, 141.3, 128.9, 127.6, 127.6, 127.5, 127.5, 127.2, 127.1, 127.0, 127.0, 126.9, 125.6, 125.5, 125.3, 125.2, 125.1, 121.1, 120.0, 120.0, 119.9, 119.9, 119.8, 108.5, 107.9, 107.5, 107.4, 106.2, 106.2, 98.2, 98.1, 79.7, 79.7, 71.6, 71.6, 71.4 (d, *J* = 17.5 Hz), 70.0 (d, *J* = 17.9 Hz), 66.1 (d, *J* = 16.6 Hz), 65.7 (d, *J* = 17.0 Hz), 49.3 (d, *J* = 7.6 Hz), 49.1 (d, *J* = 8.0 Hz), 44.9, 43.5, 43.3, 27.8, 27.7, 24.9, 24.8, 24.8, 24.7, 24.7, 24.6, 24.6, 12.6.

**HRMS** ($ESI^+$) m/z for $[C_{37}H_{44}N_2O_4P]^+$ ($M+H^+$): calcd. 611.3033, found 611.3032.

#### 2.3 Caged (p)ppGpp synthesis

##### Synthesis of guanosine-3′,5′-bisphosphate (pGp, 3)

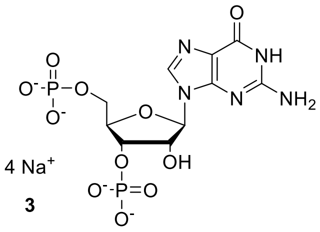

Guanosine 3′,5′ bisphosphate was synthesized as previously reported from guanosine. Analytical data in accordance with literature.[7]

##### Synthesis of guanosine-3′-phosphate-5′-diphosphate (ppGp, 8)

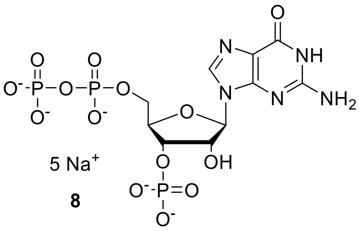

Guanosine-3’-phosphate-5’-diphosphate was synthesized as previously reported from guanosine 3’,5’ bisphosphate. Analytical data in accordance with literature.[7]

##### 5′-DEACBY-ppGp (15)

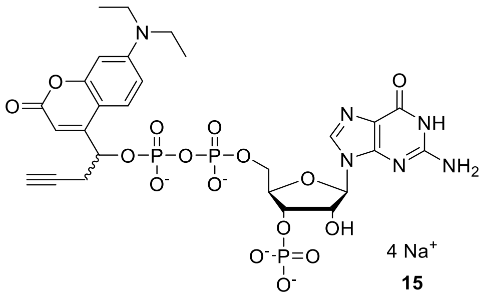

**5** was synthesized according to a modified procedure adapted from Haas *et al*.[8]

Guanosine-3’,5’-bisphosphate (mixed TBA/ammonium salt with 1.54 eq. TBA, 883 mg, 1.03 mmol, 1.0 eq) and P-amidite **10** (1.51 g, 2.47 mmol, 2.4 eq.) were coevaporated together with acetonitrile (2 x 2.5 mL). The mixture was dissolved in dry DMF (10 mL) and ETT stock solution (1 M in dry acetonitrile, 5.15 mL, 5.15 mmol, 5.0 eq.) was added. The resulting mixture was stirred for 20 min. After cooling to -20°C, *m*CPBA (427 mg, 2.47 mmol, 2.4 eq) was added, the mixture was stirred for 10 min at 0°C before precipitation with diethyl ether (300 mL). The Precipitate was separated by centrifugation, dissolved in methanol (50 mL) and after 5 min the solution was concentrated to a volume of 10 mL. After another precipitation with diethyl ether (180 mL), separation by centrifugation and washing with diethyl ether (40 mL) the solid was dried under high vacuum.

The solid was dissolved in DMF (5 mL), piperidine (500 µL) was added and the after 5 min precipitation was induced by the addition of diethyl ether (90 mL). The precipitate was separated by centrifugation, washed with diethyl ether (3 x 40 mL) and dried under high vacuum. Solid was dissolved in water (35 mL), ribonuclease T2 was added and the solution was kept at 37°C for 16 h. Then the product was purified by strong ion-exchange (Q-Sepharose) using a gradient from 0 - 200 mM $NaClO_4$ buffer. Fractions containing product were lyophilized, dissolved in 10 mL water, precipitated with 0.5 M $NaClO_4$ in acetone (80 mL), separated by centrifugation and washed with acetone (3 x 40 mL). After drying under high vacuum, the sodium salt of **15** was isolated as a slightly yellow powder (697 mg, 793 µmol, 77 %, 1:1 mixture of diastereomers)

**15** was converted into its TBA salt using a chelex® 100 column preconditioned with TBA(Br) (500 mM). Fractions containing **15** were combined and lyophilized.

**$^1$H NMR** (400 MHz, $D_2O$, *mixture of diastereomers*) δ 7.89 (s, 1H), 7.71 (s, 1H), 7.34 (d, *J* = 9.2 Hz, 1H), 7.26 (d, *J* = 9.0 Hz, 1H), 6.53 (dd, *J* = 9.2, 2.6 Hz, 1H), 6.45 (dd, *J* = 9.1, 2.6 Hz, 1H), 6.42 (d, *J* = 2.5 Hz, 1H), 6.40 – 6.28 (m, 3H), 5.79 – 5.58 (m, 4H), 4.68 – 4.58 (m, 2H), 4.54 – 4.46 (m, 2H), 4.41 – 4.31 (m, 3H), 4.26 – 4.05 (m, 3H), 3.42 – 3.32 (m, 8H), 3.13 – 3.02 (m, 1H), 3.01 – 2.91 (m, 1H), 2.85 – 2.71 (m, 2H), 2.40 (t, *J* = 2.6 Hz, 1H), 2.35 (t, *J* = 2.6 Hz, 1H), 1.18 – 1.10 (m, 12H). Assignments see appendix.
**$^{31}$P NMR** (162 MHz, $D_2O$, *mixture of diastereomers*) δ 3.18 (s, 1P), 3.02 (s, 1P), -11.04 (d, *J* = 18.1 Hz, 1P), -11.20 (d, *J* = 17.4 Hz, 1P), -12.50 (d, *J* = 18.2 Hz, 1P), -12.70 (d, *J* = 17.3 Hz, 1P).
**$^{13}$C NMR** (101 MHz, $D_2O$) δ 165.8, 165.5, 158.5, 158.4, 155.6 (d, *J* = 5.3 Hz), 155.5, 155.4, 155.3 (d, *J* = 3.3 Hz), 153.4, 153.3, 151.0, 150.8, 150.6, 150.6, 137.0, 136.6, 125.2, 124.9, 116.0, 115.9, 109.3, 104.9, 104.8, 104.7, 104.5, 96.5, 87.0, 86.6, 83.2 (dd, *J* = 9.1, 4.2 Hz), 82.7 (dd, *J* = 10.3, 4.2 Hz), 79.3, 79.2, 74.7 (d, *J* = 3.9 Hz), 74.2 (d, *J* = 4.2 Hz), 73.5 (d, *J* = 5.0 Hz), 73.3 (d, *J* = 5.0 Hz), 72.7, 72.5, 70.9 (d, *J* = 5.2 Hz), 70.8 (d, *J* = 5.4 Hz), 65.4 (d, *J* = 5.4 Hz), 64.9 (d, *J* = 5.5 Hz), 44.5, 44.5, 26.7 (d, *J* = 5.5 Hz), 26.3 (d, *J* = 4.5 Hz), 11.7, 11.7.
**HRMS** ($ESI^-$) m/z for $[C_{27}H_{31}N_6O_{16}P_3]^{2-}$ ($M^{4-}$+$2H^+$): calcd. 394.0510, found 394.0508.

#### 5′-DEACBY-ppGpp (DppGpp, 16)

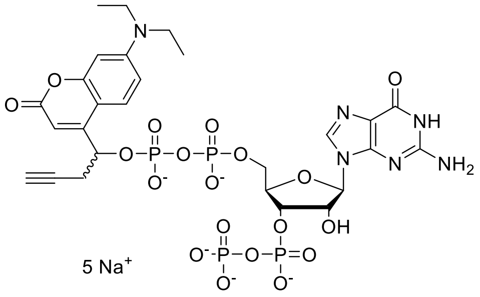

**DppGpp (16)**

**16** was synthesized according to a modified procedure adapted from Haas *et al.*[8]

TBA salt of **15** (91 mg, 52 µmol, 1.0 eq) was coevaporated with acetonitrile (0.5 mL) and dissolved in dry DMF (0.5 mL). ETT stock solution (1 M in dry acetonitrile, 155 µL, 155 µmol, 3.0 eq) and P-amidite **5** (41 mg, 78 µmol, 1.5 eq) were added successively and the solution was stirred for 15 min at room temperature. After cooling to -20°C *m*CPBA (13.4 mg, 78 µmol, 1.5 eq) was added and the mixture was stirred for 10 min before heating to 0°C and stirring for 15 min. Then DBU (100 µL) was added, the solution was stirred 5 min at 0°C and 30 min at room temperature before precipitation was induced by the addition of diethyl ether (20 mL). The precipitate was separated by centrifugation and washed with diethyl ether (3 x 10 mL). The product was purified by strong ion-exchange (Q-Sepharose) using a gradient from 0 - 200 mM $NaClO_4$ buffer. Fractions containing product were precipitated with 0.5 M $NaClO_4$ in acetone (100 mL) separated by centrifugation and washed with acetone (3 x 40 mL). After drying under high vacuum, the sodium salt of **DppGpp (16)** was isolated as a yellow solid (34 mg, 69 %, 1:1 mixture of diastereomers).

**$^1$H NMR** (400 MHz, $D_2O$, *mixture of diastereomers*) δ 7.86 (s, 1H), 7.67 (s, 1H), 7.35 (d, *J* = 9.2 Hz, 1H), 7.25 (d, *J* = 9.8 Hz, 1H), 6.54 (dd, *J* = 9.2, 2.7 Hz, 1H), 6.46 – 6.42 (m, 2H), 6.39 (d, *J* = 1.0 Hz, 1H), 6.36 (d, *J* = 2.6 Hz, 1H), 6.31 (d, *J* = 0.9 Hz, 1H), 5.80 – 5.71 (m, 2H), 5.67 – 5.57 (m, 2H), 4.54 – 4.30 (m, 5H), 4.85 – 4.70 (m, 2H, *under HDO peak*), 4.54 – 4.30 (m, 5H), 4.25 – 4.06 (m, 3H), 3.43 – 3.30 (m, 8H), 3.12 – 3.03 (m, 1H), 3.00 – 2.91 (m, 1H), 2.85 – 2.69 (m, 2H), 2.40 (t, *J* = 2.6 Hz, 1H), 2.35 (t, *J* = 2.6 Hz, 1H), 1.19 – 1.10 (m, 12H). Assignments see appendix.
**$^{31}$P{$^1$H} NMR** (162 MHz, $D_2O$, *mixture of diastereomers*) δ -5.80 (d, J = 22.6 Hz, 2P), -10.79 – -11.39 (m, 4P), -12.42 (d, J = 18.0 Hz, 1P), -12.66 (d, J = 17.2 Hz, 1P).
**$^{13}$C NMR** (101 MHz, $D_2O$, *mixture of diastereomers*) δ 165.9, 165.5, 158.5, 158.4, 155.6 (d, *J* = 5.0 Hz), 155.5, 155.4, 155.3, 153.4, 153.3, 150.9, 150.6, 150.6, 136.8, 136.5, 125.1, 124.9, 116.0, 116.0, 109.3, 104.8, 104.7, 104.7, 104.5, 96.5, 96.4, 87.5, 87.1, 82.8 (dd, *J* = 8.9, 5.1 Hz), 82.4 – 82.0 (m), 79.4, 79.2, 74.5 (d, *J* = 2.9 Hz), 74.3 – 74.0 (m), 73.9 (d, *J* = 5.3 Hz), 72.6, 72.5, 70.9 (d, *J* = 5.3 Hz), 70.7 (d, *J* = 5.2 Hz), 65.1 (d, *J* = 5.5 Hz), 64.6, 44.5, 44.5, 26.7 (d, *J* = 5.3 Hz), 26.3 (d, *J* = 3.2 Hz), 11.8, 11.7.
**UV/Vis** ($H_2O$, pH 7, nm ($mol^{-1}dm^3cm^{-1}$)): $\lambda_{max}$ ($\varepsilon$)=397 (19500)
**HRMS (ESI$^-$)** m/z for $[C_{27}H_{32}N_6O_{19}P_4]^{2-}$ ($M^{5-}+3H^+$): calcd 434.0342, found 434.0341.

#### 5′-DEACBY-pppGp (11)

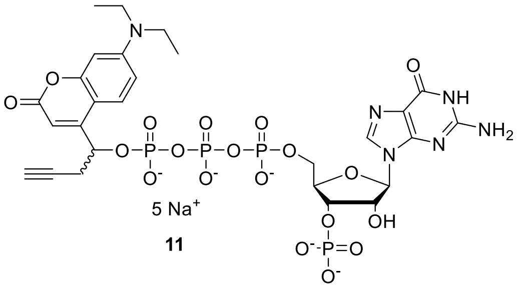

11

**8** (TBA salt with 4.46 eq. TBA, 200 mg, 124 µmol, 1.0 eq) was coevaporated once with acetonitrile (1 mL) and dissolved in DMF (1.5 mL). Then ETT (80.8 mg, 620 µmol, 5 eq) was added in one portion. P-amidite **10** (174 mg, 285 µmol, 2.3 eq.) was dissolved in DMF (1 mL) and added to the reaction mixture over 30 s. The resulting yellow solution was stirred for 30 min at room temperature. After cooling to -20°C, *m*CPBA (70.4 mg, 285 µmol, 2.3 eq) was added, the mixture was stirred for 10 min at 0°C before precipitation with diethyl ether (40 mL). The precipitate was separated by centrifugation, washed two times (30 mL diethyl ether), dissolved in methanol (5 mL) and after 5 min methanol was removed under reduced pressure and the mixture was dissolved in DMF (1 mL) and piperidine (50 µL) was added. After another 5 min precipitation was induced by the addition of diethyl ether (40 mL). The precipitate was separated by centrifugation, washed with diethyl ether (2 x 40 mL) and dried under high vacuum. Solid was dissolved in water (10 mL), precipitated with 0.5 M $NaClO_4$ in acetone and washed with acetone (2 x 40 mL). After dissolving the precipitate again in water (15 mL), ribonuclease T2 was added (332 U) and the solution was kept at 37°C for 8 h. The mixture was purified by strong ion-exchange (Q-Sepharose FF) using a gradient from 50 – 180 mM $NaClO_4$ in water. Fractions containing product or ppGp were lyophilized, dissolved in 10 mL water, precipitated with 0.5 M $NaClO_4$ in acetone (80 mL), separated by centrifugation and washed with acetone (3 x 40 mL). After drying under high vacuum, the sodium salt of **11** was isolated as a slightly yellow powder (71 mg, 72.4 µmol, 58 %, 1:1 mixture of diastereomers) along with starting material **8** (31 mg, 49 µmol, 39 %).

**11** was converted into its TBA salt using a chelex® 100 column preconditioned with TBA(Br) (500 mM). Fractions containing **11** were combined and lyophilized.

**$^1$H NMR** (400 MHz, $D_2O$, *mixture of diastereomers*) δ 7.81 (s, 1H), 7.45 (d, *J* = 9.2 Hz, 1H), 7.35 (d, *J* = 9.2 Hz, 1H), 6.56 (dd, *J* = 9.2, 2.6 Hz, 1H), 6.53 (dd, *J* = 9.2, 2.6 Hz, 1H), 6.47 (d, *J* = 2.5 Hz, 1H), 6.46 (d, *J* = 2.6 Hz, 1H), 6.41 (d, *J* = 0.8 Hz, 1H), 6.35 (d, *J* = 0.7 Hz, 1H), 5.86 – 5.79 (m, 1H), 5.81 (d, *J* = 7.1 Hz, 1H), 5.79 – 5.72 (m, 1H), 5.74 (d, *J* = 7.0 Hz, 1H), 4.74 – 4.67 (m, 2H), 4.57 – 4.49 (m, 2H), 4.44 (p, *J* = 3.0 Hz, 1H), 4.39 – 4.15 (m, 4H), 3.37 (q, *J* = 7.1 Hz, 8H), 3.19 (ddd, *J* = 17.0, 5.6, 2.5 Hz, 1H), 3.12 (ddd, *J* = 17.5, 5.3, 2.7

Hz, 1H), 2.89 (dt, $J$ = 17.4, 3.3 Hz, 1H), 2.81 (ddd, $J$ = 17.4, 4.1, 2.7 Hz, 1H), 2.40 – 2.36 (m, 2H), 1.21 – 1.10 (m, 12H). 1 Proton under HDO peak. Assignments see appendix.
**$^{31}$P{$^{1}$H} NMR** (162 MHz, $D_2O$, *mixture of diastereomers*) δ 2.31 (s, 2P), -11.03 (d, $J$ = 18.8 Hz, 1P), -11.15 (d, $J$ = 18.1 Hz, 1P), -12.66 (d, $J$ = 17.1 Hz, 1P), -12.81 (d, $J$ = 18.2 Hz, 1P), -22.43 (t, $J$ = 17.3 Hz, 1P), -22.65 (t, $J$ = 18.3 Hz, 1P).
**$^{13}$C NMR** (101 MHz, $D_2O$, *mixture of diastereomers*) δ 165.8, 165.6, 158.5, 158.5, 156.1 (d, J = 4.8 Hz), 155.7 (d, J = 5.3 Hz), 155.5, 153.4, 153.2, 151.5, 151.1, 150.8, 150.7, 137.7, 137.1, 125.6, 125.1, 116.3, 116.0, 109.6, 109.5, 105.1, 104.9, 104.5, 104.4, 96.7, 86.9, 86.9, 83.6 (d, J = 4.7 Hz), 83.5 (d, J = 4.4 Hz), 79.5, 79.2, 74.8 (d, J = 3.9 Hz), 74.0 (d, J = 4.8 Hz), 73.8 (d, J = 4.7 Hz), 73.0 (d, J = 3.9 Hz), 72.8, 72.8, 70.8 (d, J = 5.1 Hz), 70.5 (d, J = 4.7 Hz), 65.9 (d, J = 5.3 Hz), 65.7 (d, J = 6.0 Hz), 44.6, 44.5, 26.5 (d, J = 3.6 Hz), 26.1 (d, J = 4.7 Hz), 11.8, 11.7.
**HRMS (ESI$^-$)** m/z for $[C_{27}H_{32}N_6O_{19}P_4]^{2-}$ ($M^{5-}$+ $3H^+$): calcd 434.0342, found 434.0340.

#### 5′-DEACBY-pppGpp (DpppGpp, 13)

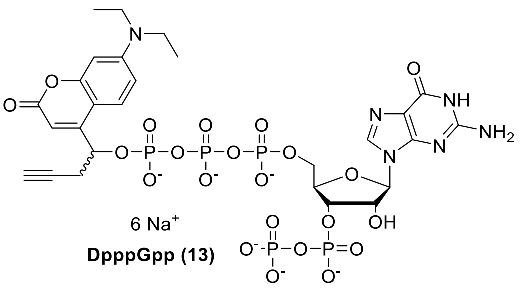

TBA salt of **11** (120 mg, 60 µmol, 1.0 eq) was dissolved in dry DMF (1 mL) under Argon. Then ETT (23 mg, 179 µmol, 3.0 eq) was added in one portion and subsequently a solution of P-amidite **5** (54 mg, 89 µmol, 1.5 eq) in dry DMF (1 mL) was added over 30 s. After stirring for 10 min, the solution was cooled to -20°C and *m*CPBA (22 mg, 89 µmol, 1.5 eq) was added. The mixture was stirred for 10 min at 0°C before DBU (100 µL) was added. After another 5 min at 0°C the solution was stirred for 10 min at room temperature. Precipitation was induced by the addition of diethyl ether (45 mL), the precipitate was separated by centrifugation and washed with diethyl ether (1 x 40 mL).

The crude product was purified by strong ion-exchange (Q-Sepharose) using a gradient from 80 - 200 mM $NaClO_4$ in water. Fractions containing product were precipitated with 0.5 M $NaClO_4$ in acetone separated by centrifugation and washed with acetone (3 x 40 mL). After drying under high vacuum, the sodium salt of **DpppGpp (13)** was isolated as a pale yellow solid (42 mg, 39 µmol, 65 %, 1:1 mixture of diastereomers).

**$^{1}$H NMR** (400 MHz, $D_2O$, *mixture of diastereomers*) δ 7.94 (s, 1H), 7.78 (s, 1H), 7.45 (d, $J$ = 9.2 Hz, 1H), 7.36 (d, $J$ = 9.2 Hz, 1H), 6.59 – 6.52 (m, 2H), 6.49 (d, $J$ = 2.6 Hz, 1H), 6.46 (d, $J$ = 2.5 Hz, 1H), 6.41 (d, $J$ = 0.9 Hz, 1H), 6.35 (d, $J$ = 0.8 Hz, 1H), 5.86 – 5.80 (m, 2H), 5.79 – 5.73 (m, 2H), 4.91 – 4.83 (m, 2H), 4.58 – 4.50 (m, 2H), 4.46 (p, $J$ = 3.0 Hz, 1H), 4.43

– 4.34 (m, 1H), 4.34 – 4.15 (m, 3H), 3.44 – 3.33 (m, 8H), 3.20 (ddd, $J$ = 17.4, 5.7, 2.6 Hz, 1H), 3.11 (ddd, $J$ = 17.4, 5.2, 2.6 Hz, 1H), 2.90 (dt, $J$ = 17.3, 3.3 Hz, 1H), 2.80 (ddd, $J$ = 17.3, 4.1, 2.6 Hz, 1H), 2.39 (t, $J$ = 2.6 Hz, 1H), 2.38 (t, $J$ = 2.6 Hz, 1H), 1.23 – 1.12 (m, 12H). 1 Proton under HDO peak. Assignments see appendix.
**$^{31}$P{$^{1}$H} NMR** (162 MHz, $D_2O$, *mixture of diastereomers*) δ -5.67 (d, $J$ = 22.2 Hz, 2P), -10.51 – -11.42 (m, 4P), -12.65 (d, $J$ = 17.0 Hz, 1P), -12.81 (d, $J$ = 18.2 Hz, 1P), -22.31 (t, $J$ = 17.1 Hz, 1P), -22.56 (t, $J$ = 17.8 Hz, 1P).
**$^{13}$C NMR** (101 MHz, $D_2O$, *mixture of diastereomers*) δ 165.8, 165.7, 158.6, 158.5, 156.1 (d, $J$ = 4.9 Hz), 155.7 (d, $J$ = 6.1 Hz), 155.6, 155.5, 153.4, 153.2, 151.4, 151.0, 150.8, 150.7, 137.6, 137.0, 125.6, 125.2, 116.3, 116.1, 109.6, 109.5, 105.0, 104.9, 104.5, 104.3, 96.8, 96.7, 87.3, 87.2, 83.3 (dd, $J$ = 9.1, 4.1 Hz), 83.1 (dd, $J$ = 8.9, 4.0 Hz), 79.6, 79.3, 79.1, 78.8, 74.9 (d, $J$ = 5.5 Hz), 74.7 (d, $J$ = 5.4 Hz), 74.4 (d, $J$ = 4.1 Hz), 72.8 (d, $J$ = 4.0 Hz), 70.7 (d, $J$ = 5.1 Hz), 70.5 (d, $J$ = 5.0 Hz), 65.8 (d, $J$ = 5.3 Hz), 65.6 (d, $J$ = 5.7 Hz), 44.6, 44.5, 26.4 (d, $J$ = 3.8 Hz), 26.0 (d, $J$ = 4.7 Hz), 11.8, 11.8.
**UV/Vis** ($H_2O$, pH 7, nm ($mol^{-1}dm^{3}cm^{-1}$)): $\lambda_{max}$ ($\varepsilon$)=397 (19600)
**HRMS** (ESI$^-$) m/z for $[C_{27}H_{31}CaN_6O_{22}P_5]^{2-}$ ($M^{6-}$+$Ca^{2+}$+$2H^+$): calcd 492.9908, found 492.9914.

#### 2.4 Synthesis of caged $^{15}$N-DpppGpp

##### [$^{15}$N]$_5$ - ppGpp (7)

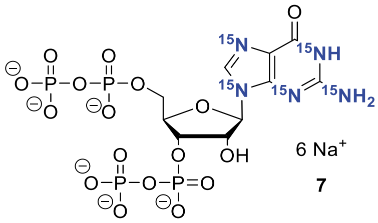

7

[$^{15}$N]$_5$ - ppGpp was synthesized starting from [$^{15}$N]$_5$ – guanosine as previously described by Qiu *et al.*[9] Analytical data were in accordance with the literature.[9]

##### [$^{15}$N]$_5$-5′-DEACBY-pppGp (12)

**Step 1:** synthesis of [$^{15}$N]$_5$ – ppGp (**9**)

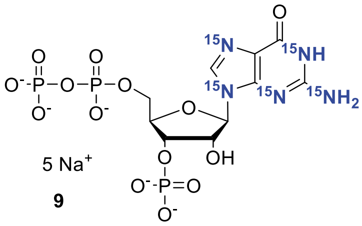

9

[$^{15}$N]$_5$ - ppGpp (70.0 mg, 94.3 μmol) was dissolved in $H_2O$ (10 mL) and RNase T2 (1 ku) was added. The solution was acidified with HCl to pH 5.5 and incubated for 12 h at 37 °C. The crude product was purified by SAX ($NaClO_4$ – buffer). The product containing fractions were precipitated with a fourfold volume of $NaClO_4$ – solution (-20°C, 0.5 M in acetone). The precipitate was washed with acetone (2 x 20 mL) and dried under high vacuum. The intermediate product $[^{15}N]_5$ – ppGp (36.9 mg, 58.5 µmol, 62 %) was isolated as Na-salt and yellow solid. The cations were exchanged to TBA by Dowex $H^+$.

**Step 2:** synthesis of **12**

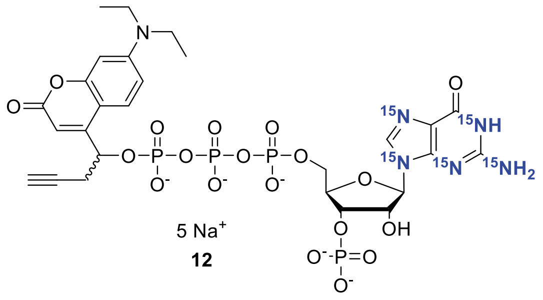

$[^{15}N]_5$ -ppGp (TBA salt with 3.5 eq. TBA, 85.5 mg, 58.5 µmol, 1.0 eq) was coevaporated once with acetonitrile (1 mL) and dissolved in DMF (1.0 mL). Then ETT (52.8 mg, 403 µmol, 5.0 eq) was added in one portion. P-amidite **10** (113 mg, 185 µmol, 2.3 eq.) was dissolved in DMF (1 mL) and added to the reaction mixture over 30 s. The resulting yellow solution was stirred for 30 min at room temperature. After cooling to -20°C, *m*CPBA (45.5 mg, 185 µmol, 2.3 eq) was added, the mixture was stirred for 10 min at 0°C before precipitation with diethyl ether (40 mL). The Precipitate was separated by centrifugation, washed two times (30 mL diethyl ether), dissolved in methanol (5 mL) and sonicated for 5 min. Methanol was removed under reduced pressure and the mixture was dissolved in DMF (1 mL) and piperidine (35 µL) was added. The solution was sonicated for 5 min before precipitation was induced by the addition of diethyl ether (40 mL). The precipitate was separated by centrifugation, washed with diethyl ether (2 x 40 mL) and dried under high vacuum. Solid was dissolved in water (10 mL), precipitated with 0.5 M $NaClO_4$ in acetone and washed with acetone (2 x 40 mL). After dissolving the precipitate again in water (15 mL), ribonuclease T2 was added (332 U) and the solution was kept at 37°C for 8 h. The mixture was purified by strong ion-exchange (Q-Sepharose FF) using a gradient from 50 – 180 mM $NaClO_4$ in water. Fractions containing product were precipitated with 0.5 M $NaClO_4$ in acetone (80 mL), separated by centrifugation and washed with acetone (3 x 40 mL). After drying under high vacuum, the sodium salt of **12** was isolated as a slightly yellow powder (16.0 mg, 30.4 µmol, 52 %, 1:1 mixture of diastereomers). **12** was converted into its TBA salt using a chelex® 100 column preconditioned with TBA(Br) (500 mM). Fractions containing **12** were combined and lyophilized.

**$^1$H NMR** (400 MHz, $D_2O$ *mixture of diastereomers*) δ 7.94 (dd, J = 10.9, 8.1 Hz, 1H), 7.80 (dd, J = 11.0, 7.8 Hz, 1H), 7.44 (d, J = 9.2 Hz, 1H), 7.34 (d, J = 9.1 Hz, 1H), 6.59 – 6.49 (m, 2H), 6.50 – 6.43 (m, 2H), 6.43 – 6.37 (m, 1H), 6.37 – 6.32 (m, 1H), 5.86 – 5.78 (m, 2H), 5.78 – 5.71 (m, 2H), 4.68 (s, 2H), 4.54 – 4.46 (m, 2H), 4.45 – 4.41 (m, 1H), 4.39 – 4.16 (m, 4H), 3.37 (q, J = 7.2 Hz, 8H), 3.27 – 3.06 (m, 2H), 2.96 – 2.73 (m, 2H), 2.38 (t, J = 2.6 Hz, 2H), 1.20 – 1.11 (m, 12H).
**$^{31}$P{$^1$H} NMR** (162 MHz, $D_2O$ *mixture of diastereomers*) δ 4.13 (s, 2P) , -10.94 (d, *J* = 18.9 Hz, 1P), -11.06 (d, *J* = 18.0 Hz, 1P), -12.63 (d, *J* = 16.8 Hz, 1P), -12.80 (d, *J* = 18.1 Hz, 1P), -22.35 (t, *J* = 17.1 Hz, 1P), -22.59 (t, *J* = 18.1 Hz, 1P).
**$^{13}$C NMR** (101 MHz, $D_2O$ *mixture of diastereomers*) δ 165.8, 165.7, 156.1 (d, *J* = 4.7 Hz), 155.7 (d, *J* = 5.1 Hz), 155.6, 155.5, 150.8, 150.7, 137.5 (d, *J* = 11.1 Hz), 136.9 (d, *J* = 11.4 Hz), 125.6, 125.1, 116.7 – 115.8 (m), 109.6, 109.5, 105.0, 104.9, 104.5, 104.4, 96.8, 96.7, 87.1, 86.9, 84.1 – 83.5 (m), 79.1, 78.8, 75.2, 73.7 (d, *J* = 4.5 Hz), 73.4 (d, *J* = 6.9 Hz), 70.8 (d, *J* = 5.3 Hz), 70.5 (d, *J* = 5.0 Hz), 66.1 (d, *J* = 5.5 Hz), 66.0 (d, *J* = 5.4 Hz), 44.6, 44.5, 26.4 (d, *J* = 3.2 Hz), 26.0 (d, *J* = 4.7 Hz), 11.8, 11.7. Some peaks not visible due to $^{15}$N-coupling.j
**HRMS** (ESI$^-$) m/z for $[C_{27}H_{34}N[^{15}N]_5O_{19}P_4]^-$ ($M^{5-}+4H^+$): calcd 874.0608, found 874.0605.

#### $[^{15}N]_5$-5′-DEACBY-pppGpp ($^{15}$N-DpppGpp, 14)

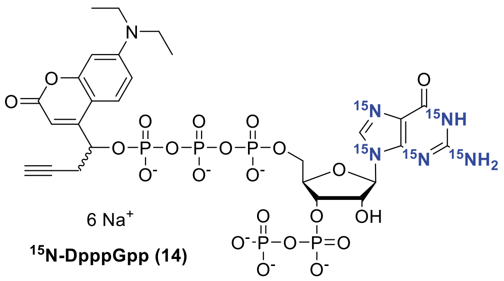

**$^{15}$N-DpppGpp (14)**

**12** (TBA salt with 5.0 eq TBA, 58.2 mg, 29 µmol, 1.0 eq) was dissolved in DMF (3.0 mL). ETT (13.2 mg, 102 µmol, 3.5 equiv) was added, followed by a solution of P-amidite **5** (27.0 mg, 52 µmol, 1.7 equiv) in DMF (1.0 mL). The resulting solution was stirred for 15 min and cooled to 0 °C before *m*CPBA (77%, 14 mg, 61 µmol, 2.1 equiv) was added. The solution was stirred for 10 min at 0 °C, and DBU (0.4 mL, 10 vol%) was added. The mixture was stirred for 30 min at room temperature and precipitated with diethyl ether (4 vol). The precipitate was separated by centrifugation (7700g, 5 min), washed with diethyl ether (2 × 15 mL), and dried under a high vacuum. The crude was purified by strong ion-exchange (Q-Sepharose FF) using a gradient from 50 – 180 mM $NaClO_4$ in water. Fractions containing product were precipitated with 0.5 M $NaClO_4$ in acetone (80 mL), separated by centrifugation and washed with acetone (3 x 40 mL). After drying under high vacuum, the sodium salt of **$^{15}$N-DpppGpp (14)** was isolated as a slightly yellow powder (13.2 mg, 12.1 µmol, 41 %, 1:1 mixture of diastereomers)

**$^1$H NMR** (400 MHz, $D_2O$, *mixture of diastereomers*) δ 7.84 (dd, *J* = 11.0, 8.0 Hz, 1H), 7.68 (dd, *J* = 11.1, 7.8 Hz, 1H), 7.37 (d, *J* = 9.2 Hz, 1H), 7.29 (d, *J* = 9.2 Hz, 1H), 6.47 (dd, *J* = 9.2, 2.6 Hz, 2H), 6.42 (d, *J* = 2.6 Hz, 1H), 6.38 (d, *J* = 2.5 Hz, 1H), 6.32 (d, *J* = 1.0 Hz, 1H), 6.26 (d, *J* = 0.8 Hz, 1H), 5.77 – 5.71 (m, 2H), 5.71 – 5.64 (m, 2H), 4.82 – 4.76 (m, 2H), 4.50

– 4.40 (m, 2H), 4.37 (s, 1H), 4.34 – 4.27 (m, 1H), 4.25 – 4.08 (m, 3H), 3.37 – 3.24 (m, 8H), 3.17 – 3.08 (m, 1H), 3.07 – 2.99 (m, 1H), 2.82 (d, *J* = 17.3 Hz, 1H), 2.71 (dd, *J* = 17.2, 4.1 Hz, 1H), 2.29 (t, *J* = 2.5 Hz, 1H), 2.28 (t, *J* = 2.6 Hz, 1H), 1.17 – 1.02 (m, 12H). 1 Proton under HDO peak.
**$^{31}P\{^1H\}$ NMR** (162 MHz, $D_2O$, *mixture of diastereomers*) δ -5.68 (d, *J* = 22.6 Hz, 2P), -10.73 – -11.31 (m, 4P), -12.75 (d, *J* = 17.6 Hz, 1P), -12.92 (d, *J* = 19.0 Hz, 1P), -22.50 (t, *J* = 17.4 Hz, 1P), -22.77 (t, *J* = 18.1 Hz, 1P).
**$^{13}C$ NMR** (101 MHz, $D_2O$, *mixture of diastereomers*) δ 165.8, 165.7, 158.6, 156.1 (d, J = 4.7 Hz), 155.7 (d, J = 5.0 Hz), 155.6, 155.5, 150.8, 150.7, 137.6, 137.0 (d, J = 10.7 Hz), 125.6, 125.1, 116.1, 109.6, 109.5, 105.0, 104.9, 104.5, 104.3, 96.7 (d, J = 5.6 Hz), 87.3, 87.2, 83.2 (d, J = 4.9 Hz), 79.6, 79.3, 74.9 (d, J = 5.7 Hz), 74.7 (d, J = 5.2 Hz), 74.5 (d, J = 4.5 Hz), 72.8 (d, J = 4.5 Hz), 70.7 (d, J = 5.4 Hz), 70.5 (d, J = 4.9 Hz), 65.8 (d, J = 5.6 Hz), 65.6 (d, J = 5.7 Hz), 44.6, 44.5, 26.6 – 26.3 (m), 26.2 – 25.9 (m), 11.8, 11.8.
**HRMS** ($ESI^-$) m/z for $[C_{27}H_{33}N[^{15}N]_5O_{12}P_5]^{2-}$ ($M^{6-}+4H^+$): calcd 476.5099, found 476.5098

#### 2.5 Copper-Catalyzed Azide-Alkyne Cycloaddition (CuAAC)

##### General Procedure Copper Catalyzed Azide-Alkyne Cycloaddition

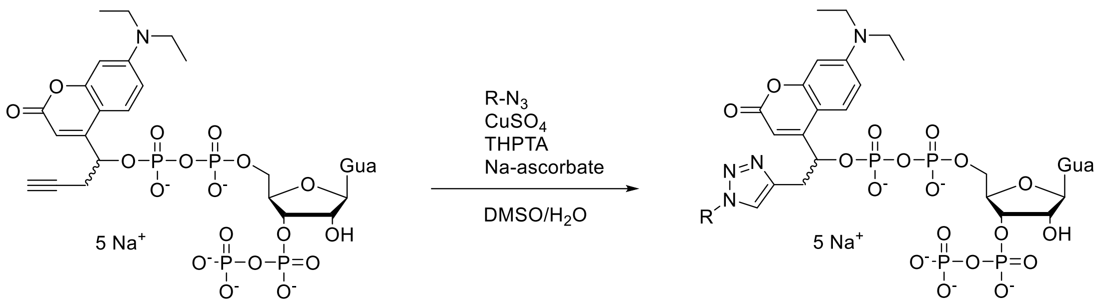

DEACBYppGpp

Azide (1-2 eq), alkyne (1-2 eq) and sodium ascorbate (1-5 eq) were dissolved in TEAA buffer (50 mM) and degassed with argon for 10 min. Stock solutions of copper sulfate (20 mM, 0.1 eq) and Tris((1-hydroxy-propyl-1H-1,2,3-triazol-4-yl)methyl)amine (THPTA, 50 mM, 0.5 eq) were premixed and added to the reaction mixture. After 1 h a small spatula of Chelex®-Na was added. The solution was filtered through a 0.2 µm filter and purified by strong ion exchange (Q-Sepharose) using a gradient from 0–1 M $NaClO_4$ buffer.

Due to the low sample quantities and the resulting long $^{13}C$ NMR acquisition times, partial degradation of the compounds occurred during measurement. To preserve sample purity, $^{13}C$ NMR spectra were therefore not recorded.

#### Glucose-DppGpp (S1)

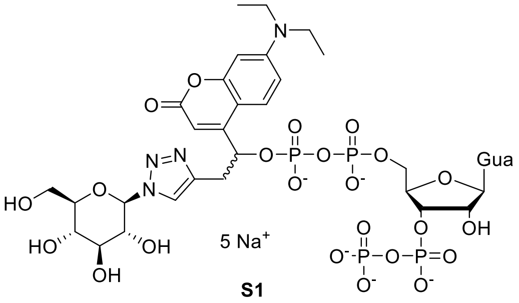

**S1**

Synthesized from **DppGpp (16)** (4.9 mg, 5 µmol, 1 eq) β-D-Glucopyranosyl azide (2.1 mg, 10 µmol, 2 eq) according to the general procedure. **S1** was obtained as a yellow powder (4.0 mg, 3.4 µmol, 67 %).

**$^{1}$H NMR** (400 MHz, $D_2O$, *mixture of diastereomers*) δ 7.94 (s, 1H), 7.92 (s, 1H), 7.69 (s, 1H), 7.50 (s, 1H), 7.20 (d, *J* = 9.2 Hz, 1H), 7.11 (d, *J* = 8.8 Hz, 1H), 6.40 (dd, *J* = 9.2, 2.6 Hz, 1H), 6.29 (d, *J* = 9.8 Hz, 2H), 6.20 (d, *J* = 2.6 Hz, 1H), 5.81 (s, 1H), 5.75 (s, 1H), 5.71 (dt, *J* = 9.2, 4.7 Hz, 1H), 5.63 (d, *J* = 5.8 Hz, 1H), 5.61 – 5.54 (m, 1H), 5.53 – 5.44 (m, 3H), 4.39 – 4.29 (m, 2H), 4.23 (t, *J* = 4.8 Hz, 1H), 4.17 – 4.07 (m, 2H), 4.07 – 3.88 (m, 3H), 3.87 – 3.70 (m, 4H), 3.67 – 3.41 (m, 8H), 3.39 – 3.03 (m, 12H), 1.06 – 0.92 (m, 12H). 2 protons under HDO peak
**$^{31}$P{$^{1}$H} NMR** (162 MHz, $D_2O$, *mixture of diastereomers*) δ -5.67 (d, *J* = 22.5 Hz, 2P), -10.68 – -11.55 (m, 4P), -12.28 (d, *J* = 18.0 Hz, 1P), -12.63 (d, *J* = 17.7 Hz, 1P).
**HRMS** (ESI-) m/z for $[C_{33}H_{43}N_9O_{24}P_4]^{2-}$ ($M^{5-}$+$3H^{+}$): calcd. 536.5691, found 536.5693.

#### Glucose-C2-DppGpp (S2)

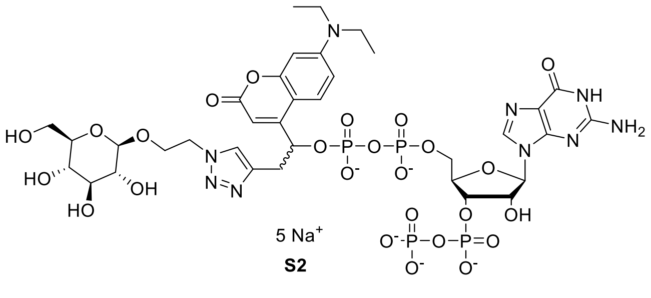

**S2**

Synthesized from **DppGpp (16)** (4.9 mg, 5 µmol, 1 eq) and 2-Azidoethyl β-D-glucopyranoside (2.5 mg, 10 µmol, 2 eq) according to the general procedure. **S2** was obtained as a yellow powder (3.7 mg, 3.0 µmol, 60 %).

**$^{1}$H NMR** (400 MHz, $D_2O$, *mixture of diastereomers*) δ 7.88 (s, 1H), 7.87 (s, 1H), 7.85 (s, 1H), 7.68 (s, 1H), 7.31 (d, *J* = 9.2 Hz, 1H), 7.15 (d, *J* = 9.8 Hz, 1H), 6.56 (dd, *J* = 9.2, 2.6 Hz, 1H), 6.44 – 6.39 (m, 2H), 6.36 (d, *J* = 2.6 Hz, 1H), 6.08 (s, 1H), 5.89 (s, 1H), 5.87 – 5.72 (m, 3H), 5.64 (d, *J* = 4.8 Hz, 1H), 4.59 – 4.49 (m, 6H), 4.43 – 4.29 (m, 5H), 4.25 – 3.96 (m, 6H), 3.90 – 3.82 (m, 3H), 3.68 (dd, *J* = 12.3, 5.4 Hz, 1H), 3.61 (dd, *J* = 12.3, 6.0 Hz, 1H), 3.53 – 3.20 (m, 20H), 1.22 – 1.11 (m, 12H). 2 protons under HDO peak

**$^{31}P\{^{1}H\}$ NMR** (162 MHz, D$_2$O, *mixture of diastereomers*) δ -5.74 (bs, 2P) -10.60 – -11.60 (m, 4P), -12.31 (d, *J* = 18.0 Hz, 1P), -12.65 (d, *J* = 17.4 Hz, 1P).
**HRMS** (ESI-) m/z for $[C_{35}H_{47}N_9O_{25}P_4]^{2-}$ ($M^{5-}+3H^{+}$): calcd. 558.5822, found 558.5824.

##### Galactose-DppGpp (S3)

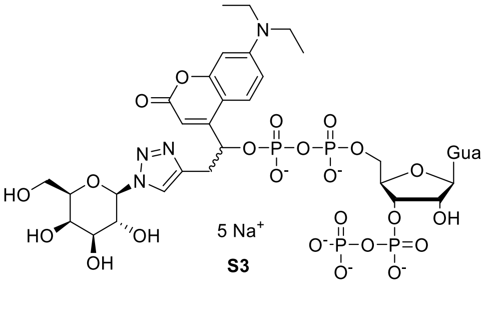

Synthesized from **DppGpp (16)** (4.9 mg, 5 µmol, 1 eq) and β-D-galactopyranosyl azide (2.1 mg, 10 µmol, 2 eq) according to the general procedure. **S3** was obtained as a yellow powder (3.4 mg, 2.9 µmol, 58 %).

**$^{1}H$ NMR** (400 MHz, D$_2$O, *mixture of diastereomers*) δ 8.13 (s, 1H), 8.11 (s, 1H), 7.87 (s, 1H), 7.67 (s, 1H), 7.37 (d, *J* = 9.2 Hz, 1H), 7.29 (d, *J* = 9.2 Hz, 1H), 6.57 (dd, *J* = 9.3, 2.6 Hz, 1H), 6.47 (dd, *J* = 9.2, 2.6 Hz, 1H), 6.44 (d, *J* = 2.6 Hz, 1H), 6.36 (d, *J* = 2.6 Hz, 1H), 5.99 (d, *J* = 0.9 Hz, 1H), 5.94 (d, *J* = 0.9 Hz, 1H), 5.87 (dt, *J* = 9.5, 4.7 Hz, 1H), 5.80 (d, *J* = 5.9 Hz, 1H), 5.73 (dt, *J* = 9.4, 4.9 Hz, 1H), 5.64 (d, *J* = 4.8 Hz, 1H), 5.60 (dd, *J* = 9.2, 0.9 Hz, 2H), 4.55 – 4.45 (m, 2H), 4.39 (t, *J* = 4.7 Hz, 1H), 4.33 – 4.11 (m, 6H), 4.13 – 4.05 (m, 3H), 4.02 – 3.94 (m, 2H), 3.86 (t, *J* = 3.1 Hz, 1H), 3.84 (t, *J* = 3.1 Hz, 1H), 3.83 – 3.73 (m, 4H), 3.55 – 3.20 (m, 12H), 1.21 – 1.11 (m, 12H). *2 Protons under HDO Peak*
**$^{31}P$ NMR** (162 MHz, D$_2$O, *mixture of diastereomers*) δ -5.74 (d, J = 19.3 Hz, 2P), -10.5 – -11.6 (m, 4P), -12.27 (d, J = 17.9 Hz, 1P), -12.65 (d, J = 17.6 Hz).
**HRMS** (ESI-) m/z for $[C_{33}H_{43}N_9O_{24}P_4]^{2-}$ ($M^{5-}+3H^{+}$): calcd. 536.5691, found 536.5693.

##### Maltose-DppGpp (S4)

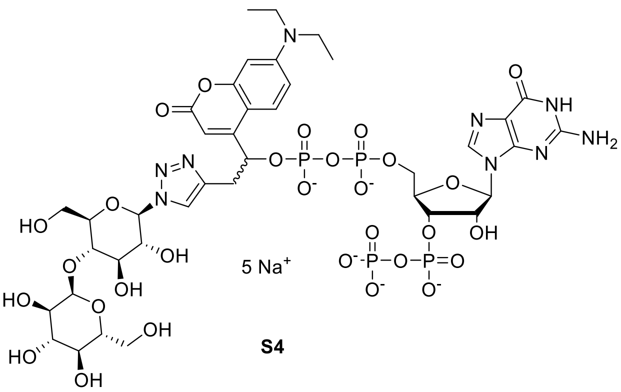

Synthesized from **DppGpp (16)** (4.9 mg, 5 µmol, 1 eq) and β-D-maltosyl azide (3.7 mg, 10 µmol, 2 eq) according to the general procedure. **S4** was obtained as a yellow powder (5.3 mg, 3.9 µmol, 79 %).

**$^1$H NMR** (400 MHz, $D_2O$, *mixture of diastereomers*) δ 7.95 (s, 1H), 7.92 (s, 1H), 7.71 (s, 1H), 7.51 (s, 1H), 7.19 (d, *J* = 9.2 Hz, 1H), 7.12 (d, *J* = 9.3 Hz, 1H), 6.40 (dd, *J* = 9.3, 2.6 Hz, 1H), 6.30 (d, *J* = 9.2 Hz, 1H), 6.27 (d, *J* = 2.4 Hz, 1H), 6.20 (d, *J* = 2.5 Hz, 1H), 5.81 (d, *J* = 1.4 Hz, 1H), 5.77 – 5.74 (m, 1H), 5.74 – 5.67 (m, 1H), 5.63 (d, *J* = 5.8 Hz, 1H), 5.58 (dd, *J* = 11.6, 7.0 Hz, 1H), 5.53 – 5.44 (m, 3H), 5.33 (dd, *J* = 3.9, 1.8 Hz, 2H), 4.39 – 4.29 (m, 2H), 4.23 (t, *J* = 4.8 Hz, 1H), 4.17 – 4.08 (m, 2H), 4.06 – 3.96 (m, 2H), 3.96 – 3.88 (m, 1H), 3.89 – 3.52 (m, 20H), 3.48 – 3.39 (m, 2H), 3.39 – 3.04 (m, 14H), 1.09 – 0.92 (m, 12H). 2 protons under HDO peak
**$^{31}P\{^1H\}$ NMR** (162 MHz, $D_2O$, *mixture of diastereomers*) δ -5.72 (d, *J* = 18.6 Hz, 2P), -10.58 – -11.55 (m, 4P), -12.29 (dd, *J* = 18.2 Hz, 1P), -12.65 (d, *J* = 17.3 Hz, 1P).
**HRMS** (ESI-) m/z for $[C_{39}H_{53}N_9O_{29}P_4]^{2-}$ ($M^{5-}$+$3H^+$): calcd. 617.5955, found 617.5950.

#### Mannose-DppGpp (S5)

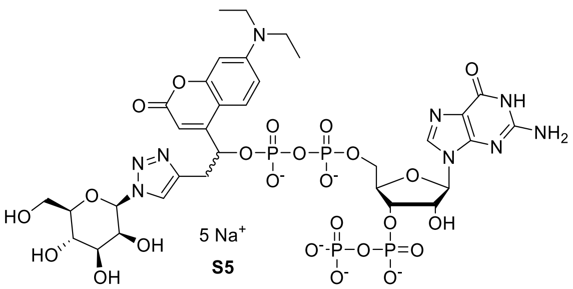

Synthesized from **DppGpp (16)** (4.9 mg, 5 µmol, 1 eq) and β-D-mannopyranosyl azide (2.1 mg, 10 µmol, 2 eq) according to the general procedure. **S5** was obtained as a yellow powder (4.3 mg, 3.6 µmol, 73 %).

**$^1$H NMR** (400 MHz, $D_2O$, *mixture of diastereomers*) δ 7.85 (s, 1H), 7.79 (s, 1H), 7.79 (s, 1H), 7.60 (s, 1H), 7.27 (d, *J* = 9.2 Hz, 1H), 7.19 (d, *J* = 9.1 Hz, 1H), 6.50 (dd, *J* = 9.3, 2.6 Hz, 1H), 6.39 (dd, *J* = 9.0, 2.6 Hz, 1H), 6.35 (d, *J* = 2.6 Hz, 1H), 6.29 (d, *J* = 2.6 Hz, 1H), 5.93 (d, *J* = 2.0 Hz, 1H), 5.92 – 5.89 (m, 2H), 5.81 (d, *J* = 0.8 Hz, 1H), 5.80 – 5.74 (m, 1H), 5.71 (d, *J* = 5.9 Hz, 1H), 5.69 – 5.64 (m, 1H), 5.56 (d, *J* = 4.9 Hz, 1H), 4.63 – 4.60 (m, 1H), 4.48 – 4.40 (m, 2H), 4.32 (t, *J* = 4.8 Hz, 1H), 4.25 – 4.18 (m, 2H), 4.15 – 3.89 (m, 6H), 3.75 – 3.63 (m, 6H), 3.41 – 3.15 (m, 13H), 2.99 (dt, *J* = 9.8, 3.9 Hz, 1H), 2.92 – 2.85 (m, 1H), 1.07 (q, *J* = 7.2 Hz, 12H). *1 Proton under HDO peak*
**$^{31}$P NMR** (162 MHz, $D_2O$, *mixture of diastereomers*) δ -5.7 (d, *J* = 22.1 Hz), 2P), -10.69 – -11.50 (m, 4P), -12.37 (d, *J* = 18.6 Hz, 1P), -12.67 (d, *J* = 17.6 Hz, 1P).
**HRMS** (ESI-) m/z for $[C_{33}H_{43}N_9O_{24}P_4]^{2-}$ ($M^{5-}$+$3H^+$): 536.5691 calcd., 536.5692 found.

#### Lactose-DppGpp (S6)

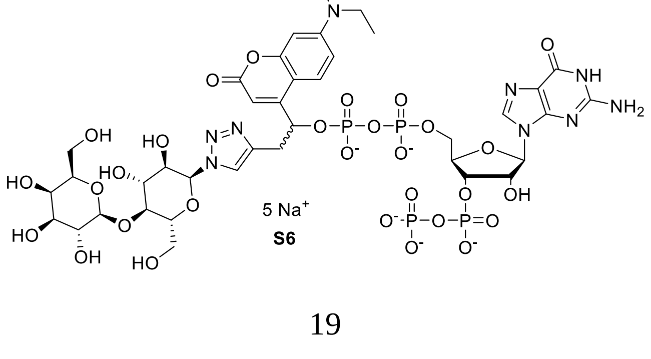

Synthesized from **DppGpp (16)** (4.9 mg, 5 µmol, 1 eq) and lactosyl azide (3.7 mg, 10 µmol, 2 eq) according to the general procedure. **S6** was obtained as a yellow oil (4.0 mg, 3.0 µmol, 59 %).

**$^{1}$H NMR** (400 MHz, $D_2O$, *mixture of diastereomers*) δ 8.11 (s, 1H), 8.09 (s, 1H), 7.87 (s, 1H), 7.68 (s, 1H), 7.35 (d, J = 9.3 Hz, 1H), 7.28 (d, J = 9.2 Hz, 1H), 6.57 (dd, J = 9.3, 2.6 Hz, 1H), 6.47 (dd, J = 9.1, 2.6 Hz, 1H), 6.44 (d, J = 2.5 Hz, 1H), 6.36 (d, J = 2.6 Hz, 1H), 5.98 (s, 1H), 5.90 (s, 1H), 5.90 – 5.83 (m, 1H), 5.79 (d, J = 5.9 Hz, 1H), 5.78 – 5.72 (m, 1H), 5.71 – 5.63 (m, 3H), 4.54 – 4.47 (m, 4H), 4.40 (t, J = 4.8 Hz, 1H), 4.34 – 4.27 (m, 2H), 4.25 – 4.12 (m, 2H), 4.12 – 4.06 (m, 1H), 4.06 – 3.73 (m, 20H), 3.73 – 3.68 (m, 2H), 3.62 – 3.21 (m, 14H), 1.22 – 1.08 (m, 12H). *2 protons under HDO peak*
**$^{31}$P NMR** (162 MHz, $D_2O$, *mixture of diastereomers*) δ -5.75 (d, J = 20.7 Hz, 2P), -10.57 – -11.45 (m, 4P), -12.32 (d, J = 17.5 Hz, 2P), -12.67 (d, J = 17.6 Hz, 2P).
**HRMS** (ESI-) m/z for $[C_{39}H_{53}N_9O_{29}P_4]^{2-}$ ($M^{5-}$+$3H^{+}$): calcd. 617.5955, found 617.5955.

#### Trehalose-DppGpp (S7)

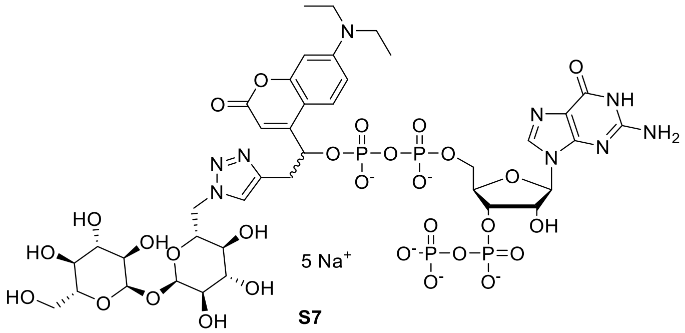

**S7**

Synthesized from **DppGpp (16)** (4.9 mg, 5 µmol, 1 eq) and 6-azido-trehalose (3.7 mg, 10 µmol, 2 eq) according to the general procedure. **S7** was obtained as a yellow powder (4.7 mg, 3.5 µmol, 70 %).

**$^{1}$H NMR** (400 MHz, $D_2O$, *mixture of* diastereomers) δ 7.87 (s, 1H), 7.81 (s, 1H), 7.77 (s, 1H), 7.69 (s, 1H), 7.39 (d, *J* = 9.2 Hz, 1H), 7.30 (d, *J* = 9.2 Hz, 1H), 6.60 (dd, *J* = 9.2, 2.6 Hz, 1H), 6.48 (d, *J* = 9.3 Hz, 1H), 6.43 (d, *J* = 2.5 Hz, 1H), 6.38 (d, *J* = 2.6 Hz, 1H), 6.00 (s, 1H), 5.93 – 5.83 (m, 2H), 5.84 – 5.71 (m, 2H), 5.64 (d, *J* = 4.8 Hz, 1H), 5.11 (d, *J* = 3.8 Hz, 1H), 5.07 (d, *J* = 3.8 Hz, 1H), 4.71 – 4.50 (m, 6H), 4.44 – 4.06 (m, 8H), 3.90 – 3.70 (m, 10H), 3.64 – 3.23 (m, 18H), 3.19 – 3.04 (m, 2H), 1.30 – 1.07 (m, 12H) 4 protons under HDO peak
**$^{31}$P NMR** (162 MHz, $D_2O$, *mixture of diastereomers*) δ -5.72 (d, *J* = 22.1 Hz, 2P), -10.57 – -11.54 (m, 4P), -12.32 (d, *J* = 18.2 Hz, 1P), -12.63 (d, *J* = 17.5 Hz, 1P).
**HRMS** (ESI-) m/z for $[C_{39}H_{53}N_9O_{29}P_4]^{2-}$ ($M^{5-}$+$3H^{+}$): calcd. 617.5955, 617.5960 found.

##### Ent-ppGpp (S8)

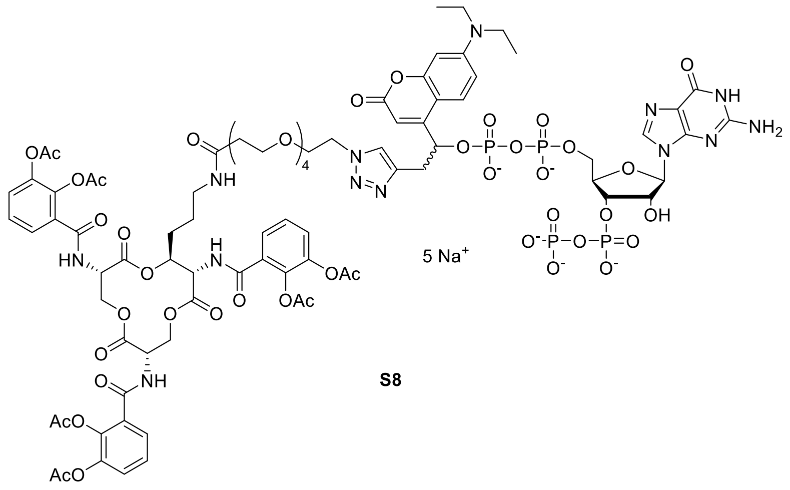

S8

Synthesized from **DEACBYppGpp (16)** (1.4 mg, 1.4 µmol, 2 eq) and Ent-N3 (S11, 0.90 mg, 0.72 µmol, 1 eq) according to the general procedure. The reaction was carried out in 66% DMSO. **S8** was obtained as an orange powder (0.37 mg, 0.17 µmol, 24 %).

**HRMS** (ESI-) m/z for $[C_{83}H_{97}N_{13}O_{45}P_4]^{2-}$ ($M^{5-}+3H^+$): 1059.7331 calcd. 1059.7335 found. Due to low scale only HRMS was performed.

#### 2.6 Synthesis of other caged nucleotides

##### 5'-DEACBY-ATP (D-ATP, S9)

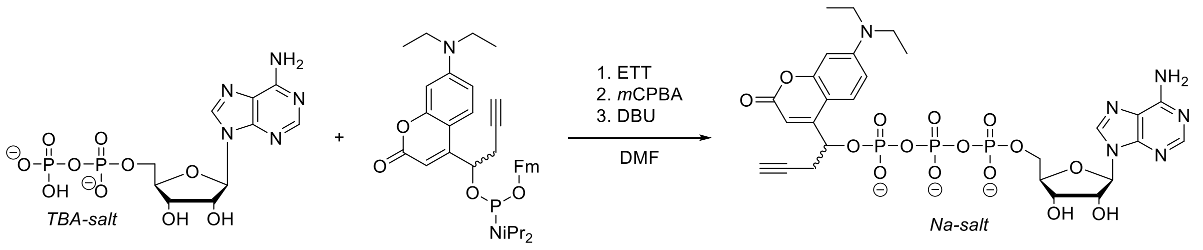

D-ATP (S9)

The sodium salt of adenosine 5′-diphosphate (ADP) was converted from into its TBA salt using a Dowex-H$^+$ column followed by addition of TBA(OH) and lyophilization resulting in a colorless powder with 2.3 TBA equivalents (determined by quantitative $^1$H NMR).

The ADP TBA salt (190 mg, 193 µmol, 1.0 eq) was dissolved in DMF (5 mL). Then ETT stock solution (1 M in dry DMF, 773 µL, 4 eq) and P-amidite **2** (295mg, 290 µmol, 1.5 eq) were added in one portion successively. The reaction mixture was stirred for 10 min at room temperature and cooled to 0°C with an ice bath before *m*CPBA (70%, 71 mg, 290 µmol, 1.5 eq) was added. After stirring for 10 min at 0°C DBU (500 µL) was added and the mixture was stirred for another 20 min at rt. Precipitation was induced by the addition of diethyl ether (40 mL), the precipitated was separated by centrifugation (4500 g, 5 min) and washed with diethyl ether (2 x 20 mL). The crude product was purified by strong ion exchange (Q-Sepharose) using a gradient from 0 to 150 mM $NaClO_4$ in water. Nucleotide precipitation was induced by dilution with 0.5 M $NaClO_4$ in acetone (10:1 v/v). The Precipitate was separated by centrifugation (4500 g, 5 min) and washed with acetone (2 x 40 mL). Drying under high vacuum resulted in the sodium salt of **S9** (**D-ATP**, 60 mg, 71 µmol, 37 %, 1:1 mixture of diastereomers) as a pale-yellow solid.

**$^{1}$H NMR** (400 MHz, $D_2O$, *mixture of diastereomers*) δ 8.28 (s, 1H), 8.15 (s, 1H), 8.15 (s, 1H), 8.12 (s, 1H), 7.40 (d, *J* = 9.2 Hz, 1H), 7.30 (d, *J* = 9.1 Hz, 1H), 6.53 (d, *J* = 8.8 Hz, 1H), 6.45 (dd, *J* = 9.1, 2.5 Hz, 1H), 6.39 (s, 1H), 6.37 – 6.29 (m, 3H), 5.93 (d, *J* = 5.5 Hz, 1H), 5.86 (d, *J* = 5.4 Hz, 1H), 5.82 (d, *J* = 5.0 Hz, 2H), 5.74 – 5.67 (m, 1H), 4.51 (t, *J* = 5.4 Hz, 1H), 4.46 – 4.34 (m, 3H), 4.34 – 4.16 (m, 6H), 3.40 – 3.21 (m, 8H), 3.13 (ddd, *J* = 17.4, 11.8, 5.7 Hz, 2H), 2.92 – 2.74 (m, 2H), 1.17 – 1.04 (m, 12H).
**$^{31}$P NMR** (162 MHz, $D_2O$) δ -11.12 – -11.38 (m, 2P), -12.57 – -12.95 (m ,2P), -22.20 – -23.07 (m, 2P).
**$^{13}$C NMR** (101 MHz, $D_2O$) δ 165.8, 165.5, 155.5, 155.4, 155.1, 155.1, 152.4, 152.3, 150.7, 150.5, 148.4, 148.1, 139.3, 139.1, 125.3, 124.8, 118.4, 118.3, 109.5, 109.3, 104.9, 104.8, 104.6, 104.5, 96.6, 96.4, 87.4, 86.9, 83.6 (d, *J* = 9.9 Hz), 83.5 (d, *J* = 8.9 Hz), 75.5, 74.7, 72.9, 71.2 (d, *J* = 4.0 Hz), 70.5 (d, *J* = 5.0 Hz), 70.4, 70.1, 65.5 – 65.1 (m), 44.4, 44.3, 26.3 (d, *J* = 3.7 Hz), 26.1 (d, *J* = 3.5 Hz), 11.7, 11.7.
**HRMS** ($ESI^-$) m/z for $[C_{27}H_{32}N_6O_{15}P_3]^-$ ($M^{3-}+2H^+$): calcd 773.1144, found 773.1136.

#### 5’-DEACBY-GDP (D-GDP, S10)

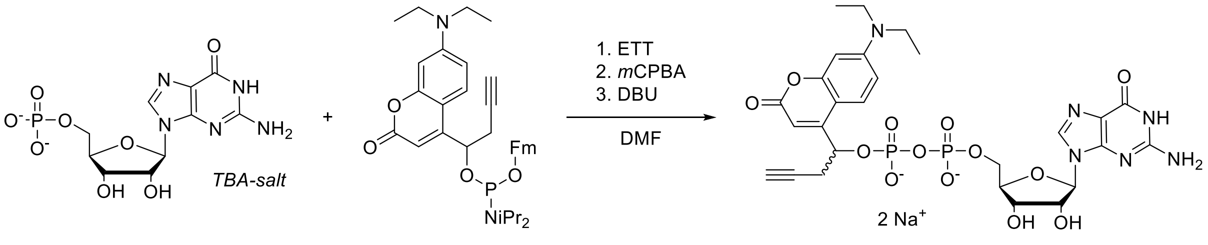

D-GDP (S10)

The sodium salt of 5′-guanylic acid was converted from its TBA salt using a chelex® 100 column preconditioned with TBA(Br) (500 mM). Lyophilization resulted in a colorless powder with 3.1 TBA equivalents (determined by quantitative $^{1}$H NMR, probably salt impurities in starting material).

The TBA salt (200 mg, 179 µmol, 1.0 eq) was dissolved in DMF (5 mL). Then ETT stock solution (1 M in dry DMF, 718 µL, 4.0 eq) and P-amidite **2** (121 mg, 197 µmol, 1.1 eq) were added in one portion successively. The reaction mixture was stirred for 10 min at room temperature and cooled to 0°C with an ice bath before *m*CPBA (70%, 53 mg, 215 µmol, 1.2 eq) was added. After stirring for 10 min at 0°C DBU (500 µL) was added and the mixture was stirred for another 20 min at rt. Precipitation was induced by the addition of diethyl ether (40 mL), the precipitated was separated by centrifugation (4500 g, 5 min) and washed with diethyl ether (2 x 20 mL). The crude product was purified by strong ion exchange (Q-Sepharose) using a gradient from 0 to 200 mM $NaClO_4$ in water. Nucleotide precipitation was induced by dilution with 0.5 M $NaClO_4$ in acetone (5:1 v/v). The Precipitate was separated by centrifugation (4500 g, 5 min) and washed with acetone (2 x 40 mL). Drying under high vacuum resulted in the sodium salt of **S10** (**D-GDP**, 32 mg, 42 µmol, 24 %) as a yellow solid with a diastereomeric ratio of 1:2.8 (diastereomer 1/diastereomer 2).

**$^{1}$H NMR** (400 MHz, $D_2O$)
*Diasteremer 1:*
δ 7.67 (s, 1H), 7.34 (d, *J* = 8.9 Hz, 1H), 6.49 – 6.40 (m, 3H), 5.81 – 5.76 (m, 1H), 5.63 (d, *J* = 3.9 Hz, 1H), 4.35 – 4.02 (m, 5H), 3.39 – 3.30 (m, 4H), 3.11 – 2.96 (m, 1H), 2.93 – 2.76 (m, 1H), 2.41 (t, *J* = 2.5 Hz, 1H), 1.13 (t, *J* = 7.1 Hz, 6H).
*Diasteremer 2:*
δ 7.81 (s, 1H), 7.49 (d, *J* = 9.2 Hz, 1H), 6.56 (dd, *J* = 9.2, 2.6 Hz, 1H), 6.43 (d, *J* = 2.6 Hz, 1H), 6.36 (s, 1H), 5.74 (d, *J* = 4.8 Hz, 1H), 5.73 – 5.68 (m, 1H), 4.35 – 4.02 (m, 5H), 3.35 (q, *J* = 5.8 Hz, 4H), 3.11 – 2.96 (m, 1H), 2.93 – 2.76 (m, 1H), 2.39 (t, *J* = 2.6 Hz, 1H), 1.13 (t, *J* = 7.1 Hz, 6H).
**$^{31}$P NMR** (162 MHz, $D_2O$)
*Diasteremer 1:*
δ -11.47 (d, *J* = 19.0 Hz, 1P), -12.87 (d, *J* = 19.1 Hz, 1P).
*Diasteremer 2:*
δ -11.18 (d, *J* = 19.1 Hz, 1P), -12.66 (d, *J* = 19.1 Hz, 1P).
**$^{13}$C NMR** (101 MHz, $D_2O$, *mixture of diastereomers*) δ 165.9, 165.5, 161.8, 161.7, 156.0, 155.9, 155.7, 155.6, 155.6, 155.5, 155.5, 150.8, 150.7, 150.7, 150.5, 136.2, 135.8, 125.2, 124.9, 116.7, 116.6, 109.4, 104.9, 104.6, 104.5, 96.8, 96.5, 87.6, 87.4, 82.8 (d, *J* = 9.3 Hz), 82.3 (d, *J* = 9.8 Hz), 78.8, 74.9, 74.4, 71.2 (d, *J* = 5.2 Hz), 70.9 (d, *J* = 5.4 Hz), 69.6, 69.2, 64.8 (d, *J* = 5.5 Hz), 64.4 (d, *J* = 5.6 Hz), 44.5, 44.3, 26.7 (d, *J* = 5.0 Hz), 26.4 (d, *J* = 5.0 Hz), 11.7, 11.6.
**HRMS** ($ESI^-$) m/z for $[C_{27}H_{31}N_6O_{13}P_2]^-$ ($M^{2-}+H^+$): calcd 709.1430, found 709.1424.

### 3 Photolysis (Figure S1)

Caged MSN were irradiated for 2 min with the Mightex 400 nm LED ($\lambda_{max}$= 396 nm, FWHM = 13 nm, distance= 4-5 cm, ~14 mW)[10] in a small glass vial under constant stirring (Figure S1). We used a 50 µM solution in 500 µL TEAA buffer (10 mM). The solution was analyzed before and after irradiation by RP-HPLC.

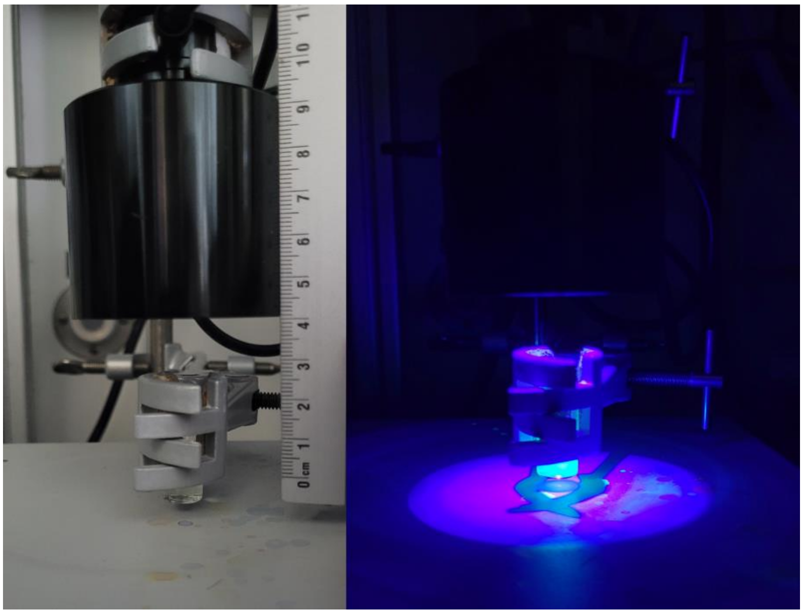

**Figure S1.** Experimental setup of photolysis experiment. Left image with LED turned off, right image with LED turned on. Cyan fluorescence of DEACBY photocage is visible.

### 4 Cell culture experiments

#### 4.1 Instruments and Material

***E. coli* Strain** BW25113 (wildtype) and knockout strains *ΔgppA, ΔacrA, ΔacrB* and *ΔtolC* from the Keio collection were ordered from Horizon Discovery Biosciences Limited.

The Escherichia coli BW25113 Δ*relA ΔspoT* strain (or (p)ppGpp$^0$ strain) was constructed using λ-Red recombination as previously described.[11] Recombinants were selected on Lennox Broth (LB) agar plates supplemented with 25 µg/mL chloramphenicol or 50 µg/mL kanamycin. The antibiotic resistance cassette was subsequently removed by FLP recombinase expressed from the temperature-sensitive plasmid pCP20.

**Media.** LB medium (Lennox) was purchased from Carl Roth. 3-(N-morpholino)propanesulfonic acid (MOPS) rich defined medium (MOPSr) was prepared as described by Neidhardt *et al*.[12] with Neidhardt Supplement mixture (Formedium™) and 0.2 % or 0.4 % glucose. For the growth assay we used MOPS rich medium with 0.2% casamino acids (MOPS+CAA) instead of Neidhardt Supplement mixture. MOPS lowP medium is MOPS minimal medium with only 200 µM phosphate and 0.4% glucose.

**Cell culture.** *E. coli* strains were streaked onto LB agar plates from a frozen glycerol stock and grown overnight at 37°C. Keio collection strains were streaked onto LB agar plates containing 50 µg/mL kanamycin. From these plates, a single colony was inoculated into LB medium (+ 50 µg/mL kanamycin for Keio collection strains) and grown overnight at 37°C while being shaken at 140 rpm. Growth was monitored by absorption at 600 nm ($OD_{600}$) with a UviLine 9400 (Schott) spectrometer. Overnight cultures were diluted by a factor of at least 10 and grown in fresh LB or MOPSr medium until $OD_{600}$ ~ 0.5−0.6.

**Confocal Microscopy** was conducted on a *ZEISS LSM 880 Observer / Fast Airyscan with inverted microscope* in the Life Imaging Center (LIC) of the University of Freiburg. The microscope was equipped with a *Plan-Apochromat 63x/1.40 Oil DIC M27* objective. We used the 405 nm and 561 nm LSM880 LASER lines together with fast Airyscan. Images were processed and analyzed with ImageJ/Fiji[13] and ZEN Software.

**Fluorescence and absorption** measurements were performed with a Tecan Spark 10M plate reader. Fluorescence was analyzed by measuring excitation at 398 nm (20 nm bandwidth) and emission at 488 nm (20 nm bandwidth).

**Flow cytometry** was performed with a *BD LSR Fortessa* in the Lighthouse Core Facility of the Medical Faculty, University of Freiburg. Data was analyzed with FlowJo.

#### 4.2 Uptake Experiments

##### 4.2.1 Conditions

**Carbohydrate conjugates.** *E. coli* cell culture was grown until $OD_{600}$ 0.5 in LB medium and washed two times with PBS. Cells were incubated with carbohydrate-DppGpp conjugates (100 µL, 100 µM in PBS) for 1 h at 37°C, washed two times with PBS resuspended in PBS (150 µL) and fluorescence was measured.

**Siderophore conjugate.** Cells were treated as described by Zscherp *et al.*[2] with compound **S8** (10 µM).

**Synthetic nucleotide transporter (SNT)**. To adjust to the cell concentration, we took samples equal to $\frac{0.5}{OD_{600}}$ mL. Samples were centrifuged (5 min, 5000 G), washed 1x with MOPSr medium and the incubated in a freshly prepared SNT/caged MSN mixture (100 µL, 40 µM) in MOPSr medium for 30 min at room temperature. Then the samples were washed two times and further analyzed by fluorescence spectroscopy, flow cytometry or confocal microscopy.

##### 4.2.2 Analysis

**Fluorescence measurements** were conducted in a black 96 well plates in a Tecan plate reader. Samples were resuspended in 150 µL PBS or MOPSr medium and transferred to the wells. Wells were excited with 398 nm and emission was detected at 488 nm. Fluorescence was normalized to autofluorescence of untreated cells.

**Flow cytometry** was recorded without washing, samples were resuspended in PBS (concentration adjusted to <10 000 events/s, usually 2−5 mL) and flow cytometry measurements were conducted immediately.

**For confocal microscopy experiments**, cells were resuspended in a freshly prepared solution of Nile red (10 µM in PBS). Aliquots of 5−10 µL were immobilized under an agar pad and cells were imaged directly.

##### 4.2.3 Uptake of carbohydrate derivatives (Figure S2)

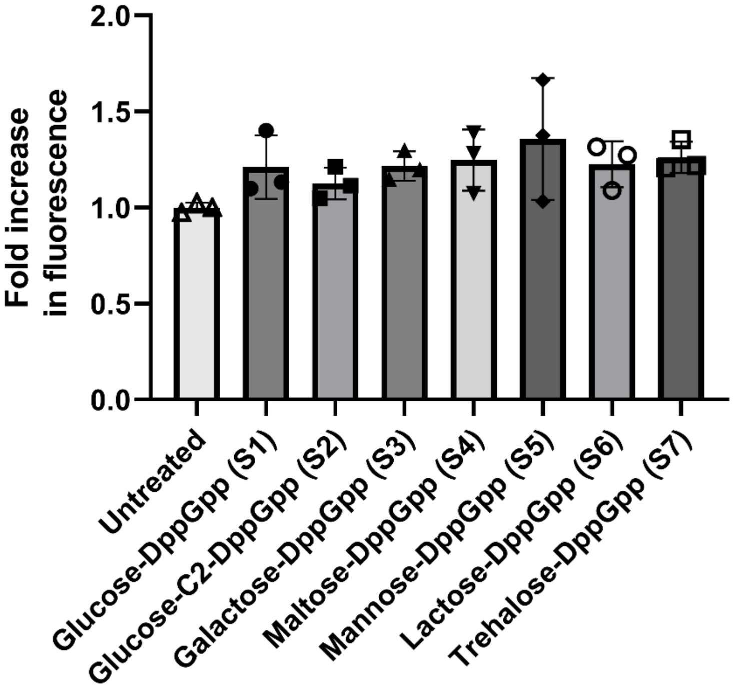

**Figure S2.** Cellular uptake determined by measuring fluorescence of cell suspensions. Values are normalized to the autofluorescence of untreated cells. Mean ± standard deviation is shown.

##### 4.2.4 Supporting microscopy images (Figure S3-S5)

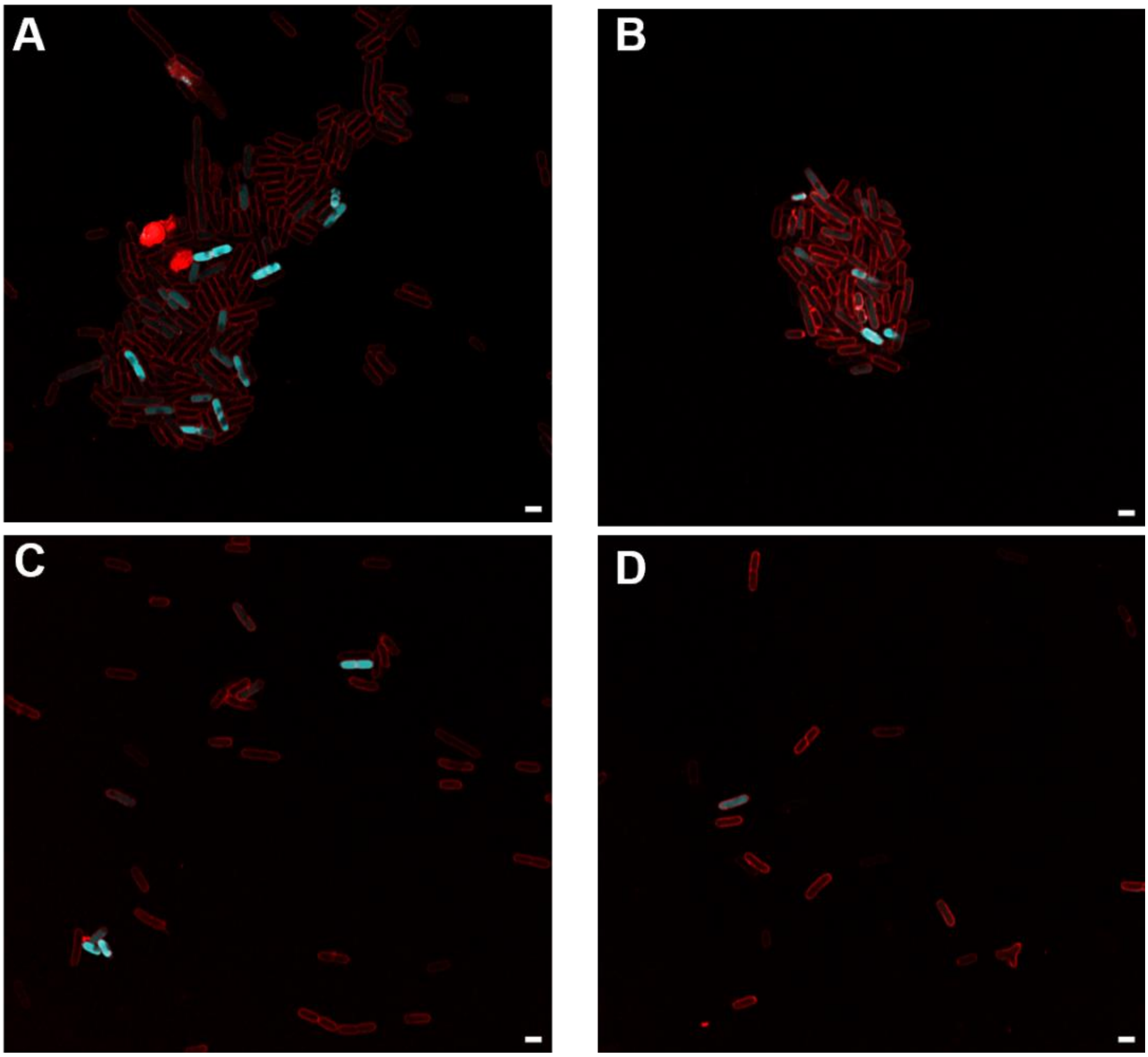

**Figure S3.** Confocal microscopy images of *E. coli* treated with DppGpp/SNT and the membrane stain Nile Red. Cage (DEACBY) fluorescence is depicted in cyan and Nile Red fluorescence in red. Scale bar = 2 µm.

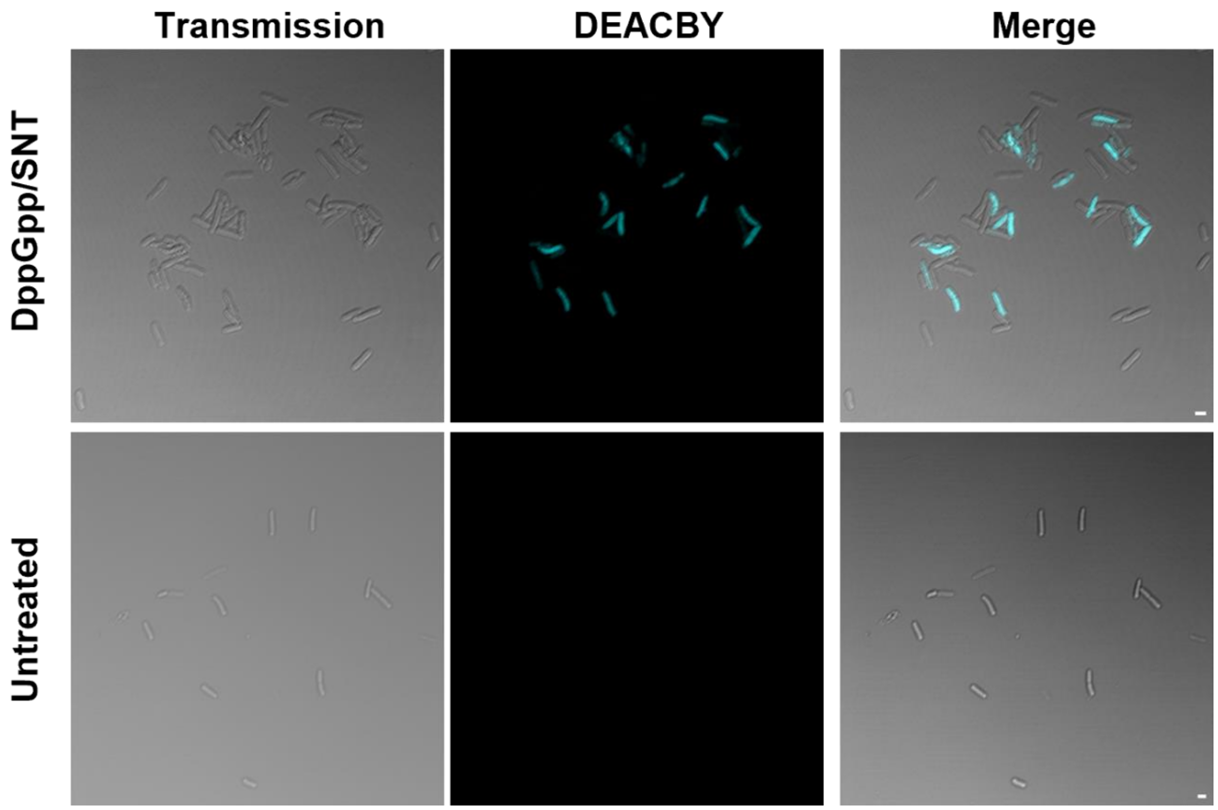

**Figure S4.** Transmission and fluorescence images of *E. coli* cells treated with DppGpp/SNT (upper images) or untreated (lower images). Scale bar = 2 µm.

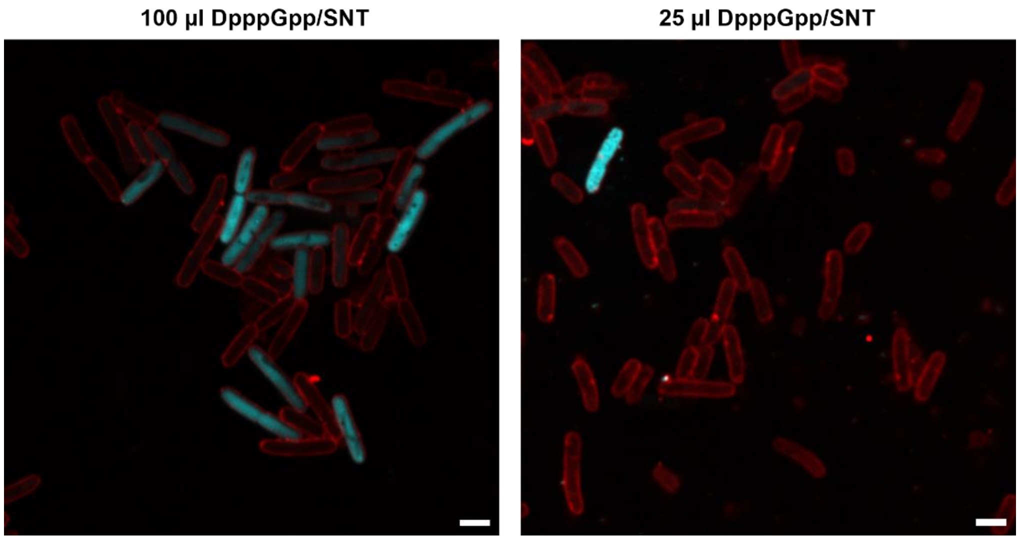

**Figure S5.** Confocal microscopy images of *E. coli* treated with either 100 µL or 25 µL DpppGpp/SNT and the membrane stain Nile Red. Cage (DEACBY) fluorescence is depicted in cyan and Nile Red fluorescence in red. Scale bar = 2 µm.

#### 4.3 Viability

*E. coli* (BW25113) cell culture was grown in MOPSr until $OD_{600}$ 0.89 and we took 8 samples of 0.6 mL. Each sample was centrifuged (10 min, 5000 g) incubated for 30 min at room temperature with the following conditions:

- 2 x only DpppGpp (13) 25 µL, 40 µM
- 3x DpppGpp/SNT 25 µL, 40 µM
- 3x DpppGpp/SNT 100 µL, 40 µM

The cells were centrifuged (3 min, 2000 g), washed with PBS (0.5 mL) and then resuspended in 100 µL MOPSr medium. We diluted 2 µL of this suspension in 150 µL MOPSr medium and, added 20 µL resazurin solution (0.15 mg/mL) and measured viability after 2 h at 37 °C through the absorption at 570 and 600 nm. In previous experiments, we found that treatment with DpppGpp alone had no effect on viability, so we treated this condition as fully viable.

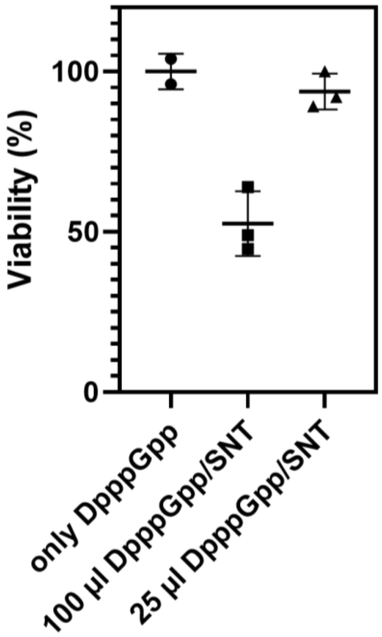

**Figure S6**. Viability of cells treated with 25 or 100 µL DpppGpp/SNT determined using resazurin.

#### 4.4 Nucleotide extraction

##### 4.4.1 General Procedure

*E. coli* (BW25113) cell culture was grown in MOPSr until $OD_{600}$ 0.5–0.6. To adjust to the cell concentration, we took samples equal to $\frac{3}{OD_{600}}$ mL. Cells were separated by centrifugation (10 min, 5000 g) incubated in freshly prepared SNT/Nucleotide mixture in MOPSr medium (150–600 µL, 40 µM) for 30 min. Incubation solution was removed by centrifugation (5 min, 2400 g). Then the samples were washed two times (1 mL PBS, 5 min, 2400 g) and resuspended in 6 mL MOPSr medium. Suspension was put on ice and the first sample (1 mL) was taken. Suspension was irradiated 3 min with the Mightex 400 nm, and the second sample (1 mL) was taken. Then the suspension was incubated at 37 °C (140 rpm) and 3 more samples were taken at 5 min, 30 min, and 60 min. All samples were directly added to 38 µL ice cold formic acid (resulting in a 1 M formic acid solution) then frozen in liquid nitrogen.

Nucleotides were extracted based on a method described by Bartoli *et. al.*[14]. Samples were thawed at 37 °C, isotopic standards were added and the samples were incubated on ice for 30 min with regular vortexing. Then, $NH_4OAc$ (3 mL, 50 mm, pH 5.5, pre-chilled) was added and cell debris was removed by centrifugation (10 min, 7500 rpm, 4 °C). Supernatants were loaded onto EVOLUTE® EXPRESS WAX 100 mg/3 mL columns pretreated with MeOH (1 mL, pre-chilled) and $NH_4OAc$ (1 mL, 50 mm, pH 4.5, pre-chilled). Columns were washed with $NH_4OAc$ (1 mL, 50 mm, pH 4.5, pre-chilled) and MeOH (1 mL, pre-chilled). Nucleotides were eluted with a mixture of MeOH/$H_2O$/$NH_4OH$ (20:70:10, 1.5 mL, pre-chilled), diluted with water (2 mL, pre-chilled), lyophilized and then dissolved in water (45 µL) for CE/MS analysis.

##### 4.4.2 Wildtype (BW25113) + DpppGpp/SNT (Figure 5B)

Cells were incubated with two times 150 µL DpppGpp/SNT and with SNT/D-GDP mixture (40 µM in MOPSr) for 30 min at rt. One DpppGpp/SNT served as control and was not irradiated with 400 nm light. Experiment was done two times.

##### 4.4.3 *ΔgppA* strain + DpppGpp/SNT (Figure 5D)

Cells were incubated with 500 µL DpppGpp/SNT and SNT/D-GDP mixture (40 µM in MOPSr) for 30 min at rt.

##### 4.4.4 Wildtype (BW25113) + $^{15}$N-DpppGpp/SNT (Figure 5C)

Cells (8.33 mL, $OD_{600}$=0.60) were incubated with 1000 µL $^{15}$N-DpppGpp/SNT (40 µM in MOPSr) for 30 min at rt. After washing the pellet was resuspended in 10 mL MOPSr medium. Additional samples were taken after 90, 120 and 150 min.

##### 4.4.5 Wildtype (BW25113) + DpppGpp/SNT + [$^{15}$N]$_5$-pppGpp spiked after uncaging (Figure S7)

Cells (6.36 mL, $OD_{600}$=0.55) were incubated with 200 µL DpppGpp/SNT. After washing the pellet was resuspended in 10 mL MOPSr medium. Cell suspension was spiked with [$^{15}$N]$_5$-pppGpp after irradiation with 400 nm light.

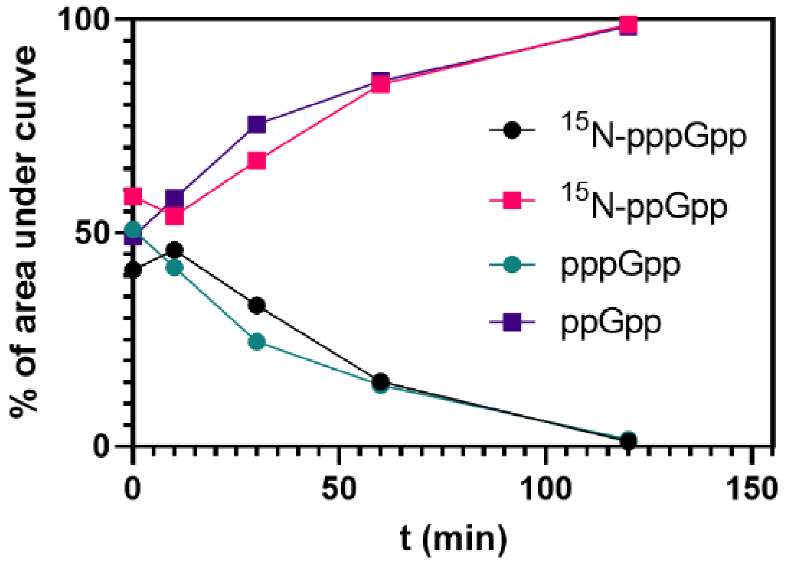

**Figure S7.** Ratio of MSNs and $^{15}$N-MSNs of *E. coli* incubated with DpppGpp/SNT and irradiated with 400 nm light. After irradiation, the suspension was spiked with [$^{15}$N]$_5$-pppGpp.

##### 4.4.6 Wildtype (BW25113) + DpppGpp/SNT + cell separation (Figure S8)

Cells (7.1 mL, $OD_{600}$=0.42) were incubated with 150 µL DpppGpp/SNT. After washing once with MOPSr medium, the pellet was resuspended in 6 mL MOPSr medium. Prior to lysis, samples were centrifuged (5 min 2 kG followed by 5 min 5 kG), supernatant was transferred to a tube with ice cold formic acid (38 µL) and the pellet was lysed in 1 M ice cold formic acid. The samples were directly frozen in liquid nitrogen.

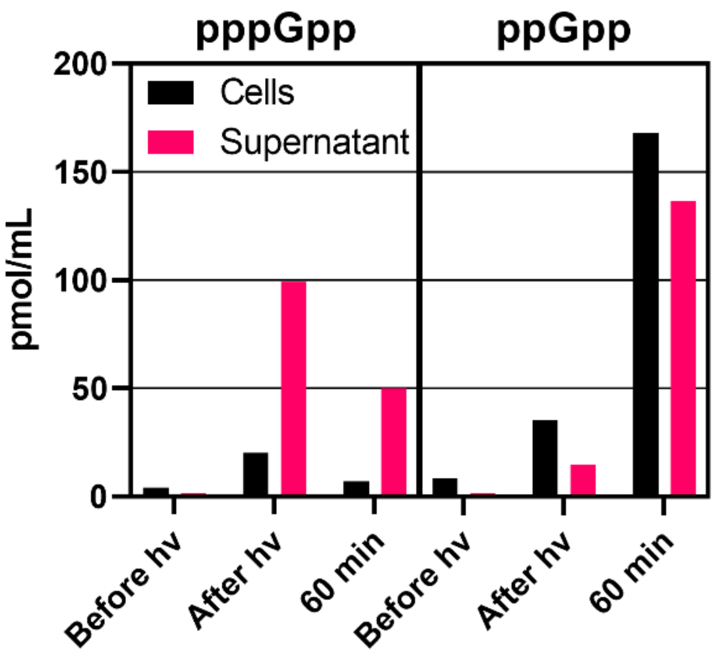

**Figure S8.** Concentration of MSNs extracted from *E. coli* incubated with DpppGpp/SNT. Cells and supernatant were separated prior to lysis.

##### 4.4.7 Wildtype (BW25113) + $^{15}$N-DpppGpp/SNT + cell separation (Figure 5E)

Cells (5.55 mL, $OD_{600}$=0.54) were incubated with 150 µL DpppGpp/SNT in MOPS lowP medium. After washing with MOPSr medium, the pellet was resuspended in 6 mL MOPSr medium. Prior to lysis, samples were centrifuged (5 min 5 kG), supernatant was transferred to a tube with ice cold formic acid (38 µL) and the pellet was lysed in 1 M ice cold formic acid. The samples were directly frozen in liquid nitrogen.

#### 4.5 Fluorescence wash solution (Figure S9)

*E. coli* cell cultures (BW25113, *ΔacrA*, *ΔacrB*, *ΔtolC*) were grown in LB until $OD_{600}$ 0.5−0.6. To adjust to the cell concentration, we took one sample from each strain equal to $\frac{0.5}{OD_{600}}$ mL. Cells were separated by centrifugation (5 min, 5000 g) and washed once with MOPS lowP medium (0.5 mL). Each strain was incubated in DpppGpp/SNT (40 µM, 25 µL) for 30 min and incubation solution were separated from the cells by centrifugation (3 min, 2400 g). The incaution solution was diluted to 200 µL in a black 96 well plate. Then the samples were washed four times (200 µL MOPS, 5 min, 2400 g) and the washing solution were added to the 96 well plate. The cell pellets were resuspended in MOPS (200 µL) and also added to the plates. DpppGpp concentration was determined by fluorescence using a calibration curve constructed from five standards.

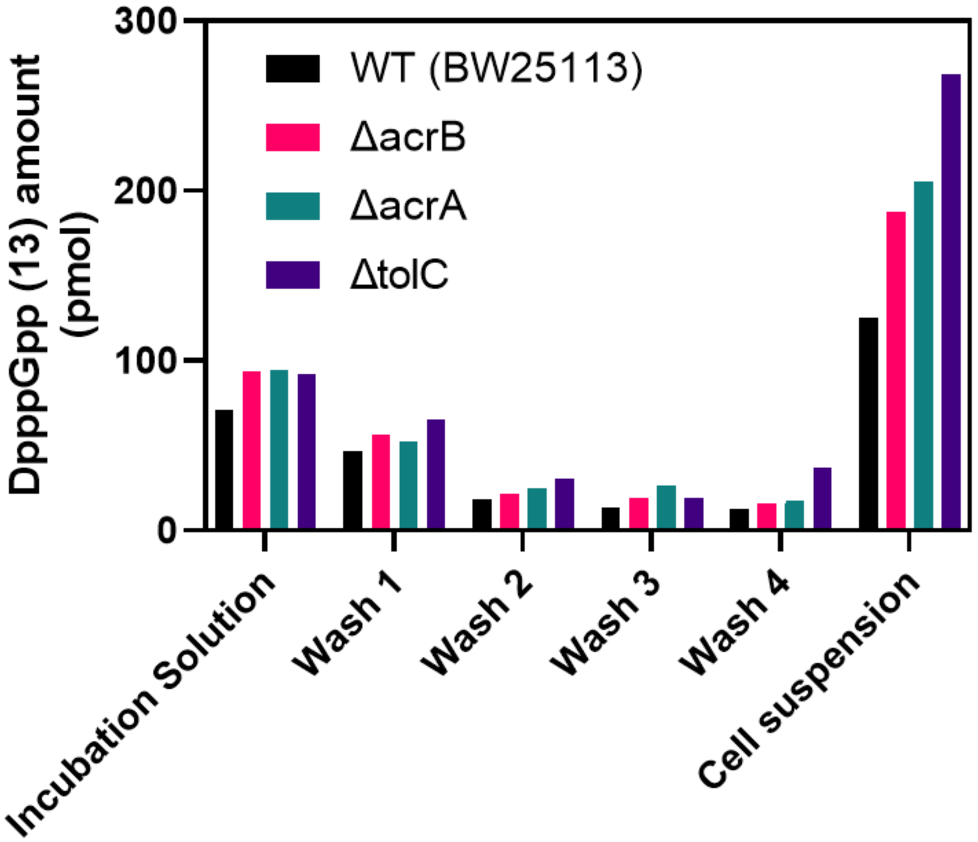

**Figure S9.** DpppGpp amount in incubation and washing solutions and in the remaining cells of *E. coli* wildtype (BW25113), , *ΔacrA*, *ΔacrB* and *ΔtolC* strains incubated with DpppGpp/SNT (40 µM, 25 µL, 1000 pmol).

#### 4.6 Growth Assay (Figure 6)

*E. coli* (p)ppGpp$^0$ cell culture was grown in LB until $OD_{600}$ 0.5−0.6. To adjust to the cell concentration, we took four samples equal to $\frac{0.5}{OD_{600}}$ mL. Cells were separated by centrifugation (5 min, 5000 g) and washed once with MOPS lowP medium. Samples were incubated either in the following conditions (25 µL, 40 µM in MOPS lowP):

- DpppGpp/SNT
- D-ATP/SNT
- SNT
- Only MOPS lowP

Or in these conditions:

- DppGpp/SNT
- D-ATP/SNT
- Only MOPS lowP

Incubation solution was removed by centrifugation (3 min, 2400 g). Then the samples were washed two times (0.2 mL MOPS+CAA, 5 min, 2400 g) and resuspended in 1 mL MOPS+CAA medium. Each sample were split in two 500 µL fractions. One fraction was kept in the dark, while the other one was irradiated for 30 s with the Darkbeam 400 nm LED. Three time 150 µL of each fraction were added to a 96 well plate and shaken at 37 °C for 18 h. Growth was measured through absorption at 600 nm.
Both experiments were conducted four times.

### 5 Bibliography

[1] G. R. Fulmer, A. J. M. Miller, N. H. Sherden, H. E. Gottlieb, A. Nudelman, B. M. Stoltz, J. E. Bercaw, K. I. Goldberg, "NMR chemical shifts of trace impurities: Common laboratory solvents, organics, and gases in deuterated solvents relevant to the organometallic chemist" *Organometallics* **2010**, *29*, 2176–2179.

[2] R. Zscherp, J. Coetzee, J. Vornweg, J. Grunenberg, J. Herrmann, R. Müller, P. Klahn, "Biomimetic enterobactin analogue mediates iron-uptake and cargo transport into E. coli and P. aeruginosa" *Chem Sci* **2021**, *12*, 10179–10190.

[3] P. Seyfried, L. Eiden, N. Grebenovsky, G. Mayer, A. Heckel, "Photo-Tethers for the (Multi-)Cyclic, Conformational Caging of Long Oligonucleotides" *Angew Chem Int Ed* **2017**, *56*, 359–363.

[4] X. Wang, University of Freiburg, PhD thesis, University of Freiburg, **2021**.

[5] A. Hofer, G. S. Cremosnik, A. C. Müller, R. Giambruno, C. Trefzer, G. Superti-Furga, K. L. Bennett, H. J. Jessen, "A Modular Synthesis of Modified Phosphoanhydrides" *Chem Eur J* **2015**, *21*, 10116–10122.

[6] K. Pahnke, C. Meier, "Synthesis of a Bioreversibly Masked Lipophilic Adenosine Diphosphate Ribose Derivative" *ChemBioChem* **2017**, *18*, 1616–1626.

[7] T. M. Haas, P. Ebensperger, V. B. Eisenbeis, C. Nopper, T. Dürr, N. Jork, N. Steck, C. Jessen-Trefzer, H. J. Jessen, "Magic spot nucleotides: tunable target-specific chemoenzymatic synthesis" *Chem Commun* **2019**, *55*, 5339–5342.

[8] T. M. Haas, P. Ebensperger, V. B. Eisenbeis, C. Nopper, T. Dürr, N. Jork, N. Steck, C. Jessen-Trefzer, H. J. Jessen, "Magic spot nucleotides: tunable target-specific chemoenzymatic synthesis" *Chem Commun* **2019**, *55*, 5339–5342.

[9] D. Qiu, E. Lange, T. M. Haas, I. Prucker, S. Masuda, Y. L. Wang, G. Felix, G. Schaaf, H. J. Jessen, "Bacterial Pathogen Infection Triggers Magic Spot Nucleotide Signaling in Arabidopsis thaliana Chloroplasts through Specific RelA/SpoT Homologues" *J Am Chem Soc* **2023**, *145*, 16081–16089.

[10] J. Ma, A. Ripp, D. Wassy, T. Dürr, D. Qiu, M. Häner, T. Haas, C. Popp, D. Bezold, S. Richert, B. Esser, H. J. Jessen, "Thiocoumarin Caged Nucleotides: Synthetic Access and Their Photophysical Properties" *Molecules* **2020**, *25*, 5325.

[11] K. A. Datsenko, B. L. Wanner, "One-step inactivation of chromosomal genes in Escherichia coli K-12 using PCR products" *Proc Natl Acad Sci U S A* **2000**, *97*, 6640–6645.

[12] F. C. Neidhardt, P. L. Bloch, D. F. Smith, "Culture Medium for Enterobacteria" *J Bacteriol* **1974**, *119*, 736–747.

[13] J. Schindelin, I. Arganda-Carreras, E. Frise, V. Kaynig, M. Longair, T. Pietzsch, S. Preibisch, C. Rueden, S. Saalfeld, B. Schmid, J.-Y. Tinevez, D. J. White, V. Hartenstein, K. Eliceiri, P. Tomancak, A. Cardona, "Fiji: an open-source platform for biological-image analysis" *Nat Methods* **2012**, *9*, 676–682.

[14] J. Bartoli, S. Citerne, G. Mouille, E. Bouveret, B. Field, "Quantification of guanosine triphosphate and tetraphosphate in plants and algae using stable isotope-labelled internal standards" *Talanta* **2020**, *219*, 121261.

### 6 Appendix

### 1H-NMR spectrum of 10

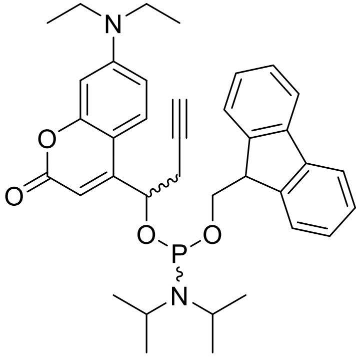

10

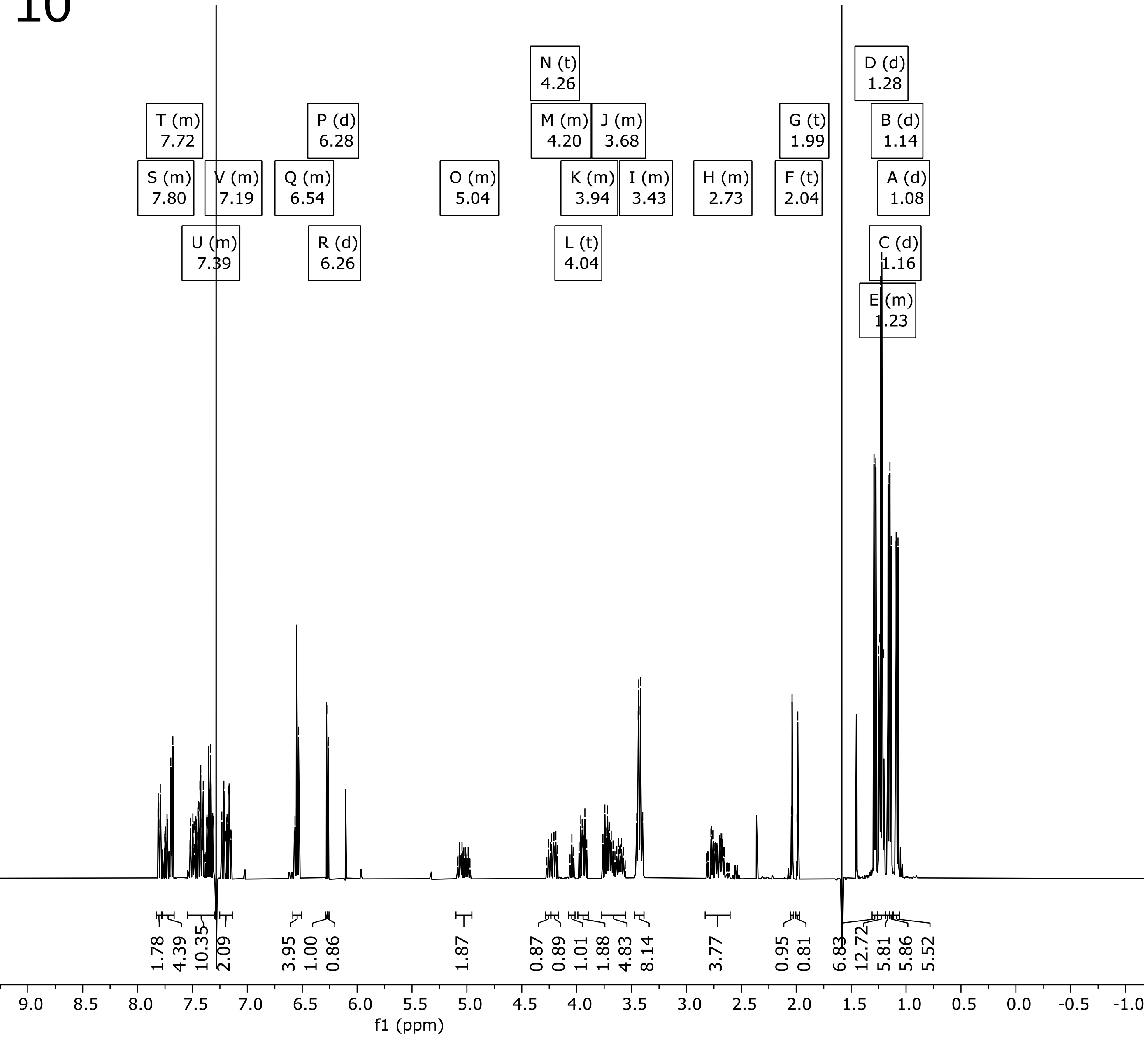

### $^{31}P\{^{1}H\}$-NMR spectrum of 10

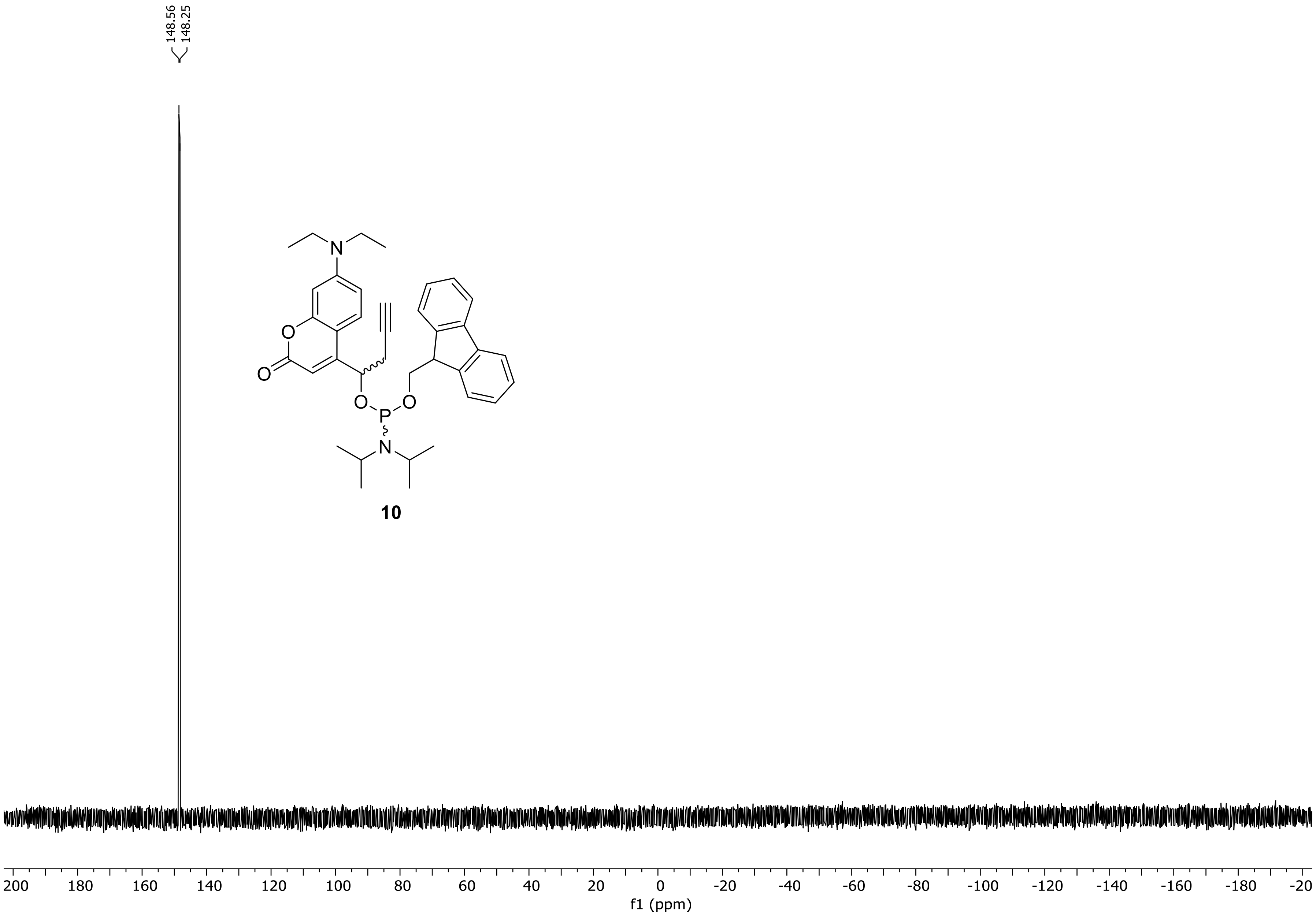

### $^{13}C$ {$^{1}H$}-NMR spectrum of 10

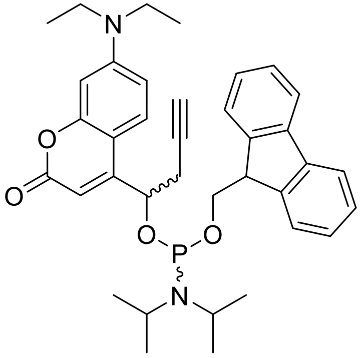

**10**

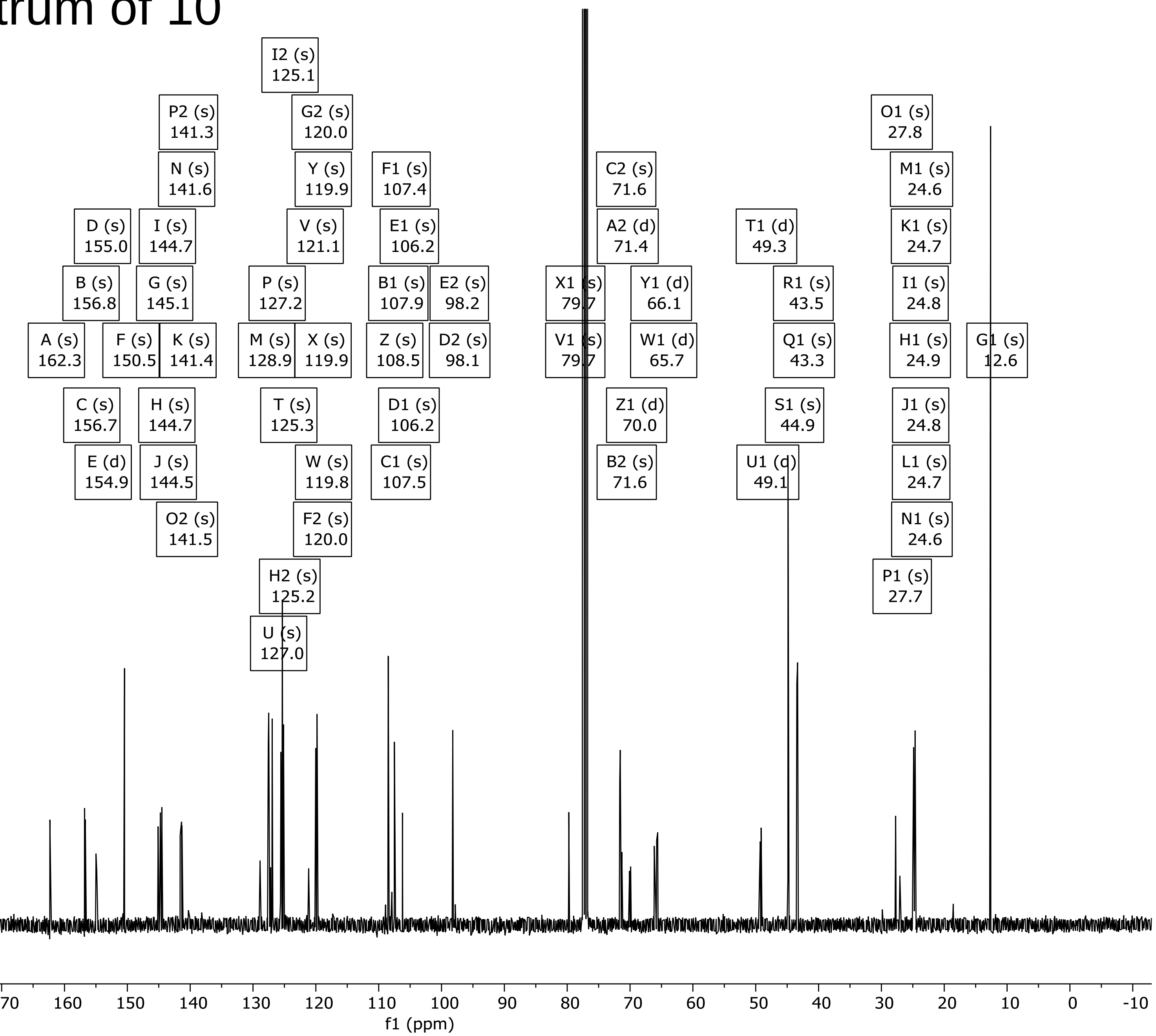

### $^1$H-NMR spectrum of 11

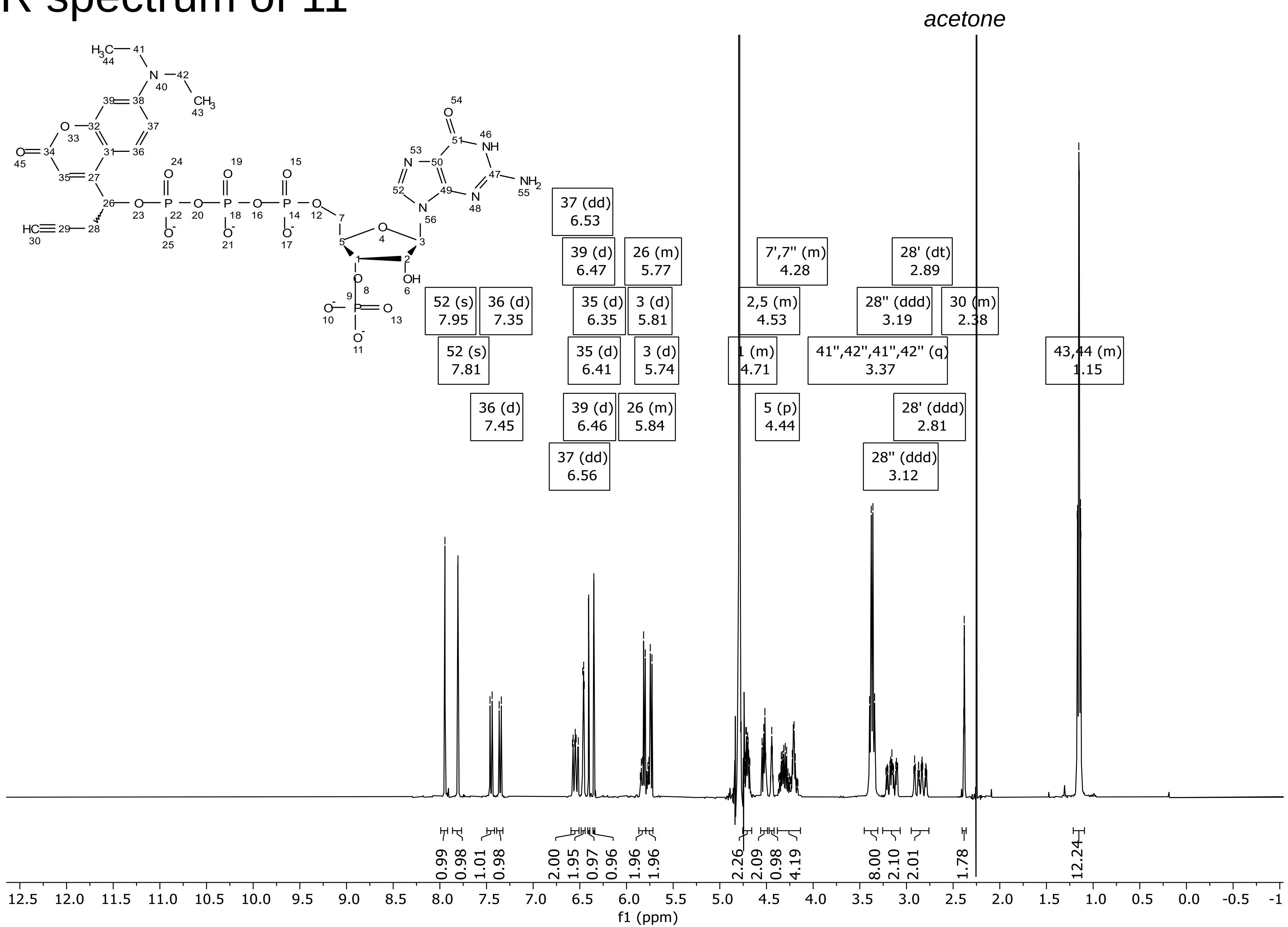

### $^{31}P\{^{1}H\}$ NMR spectrum of 11

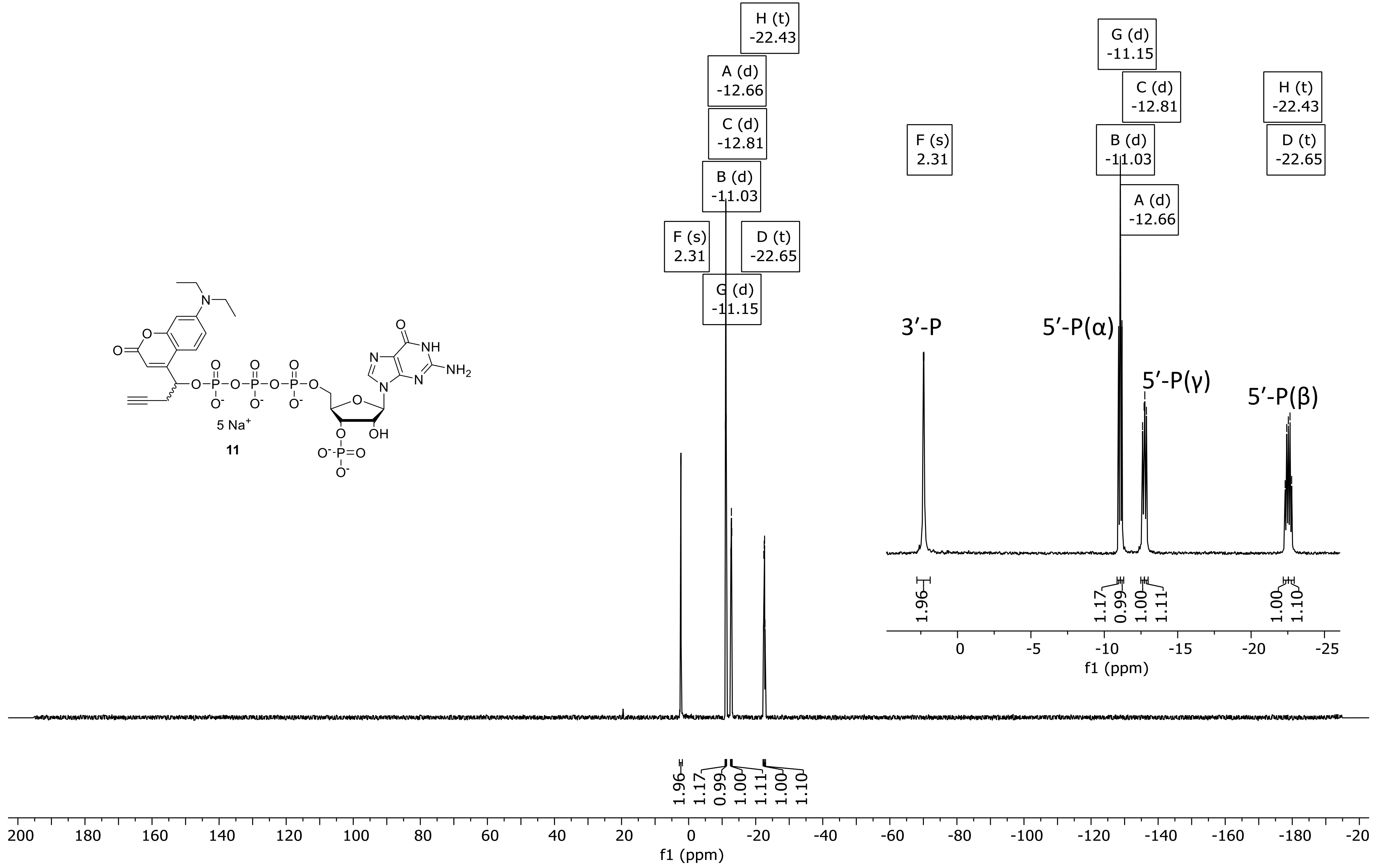

### $^{13}C$ {$^{1}H$} -NMR spectrum of 11

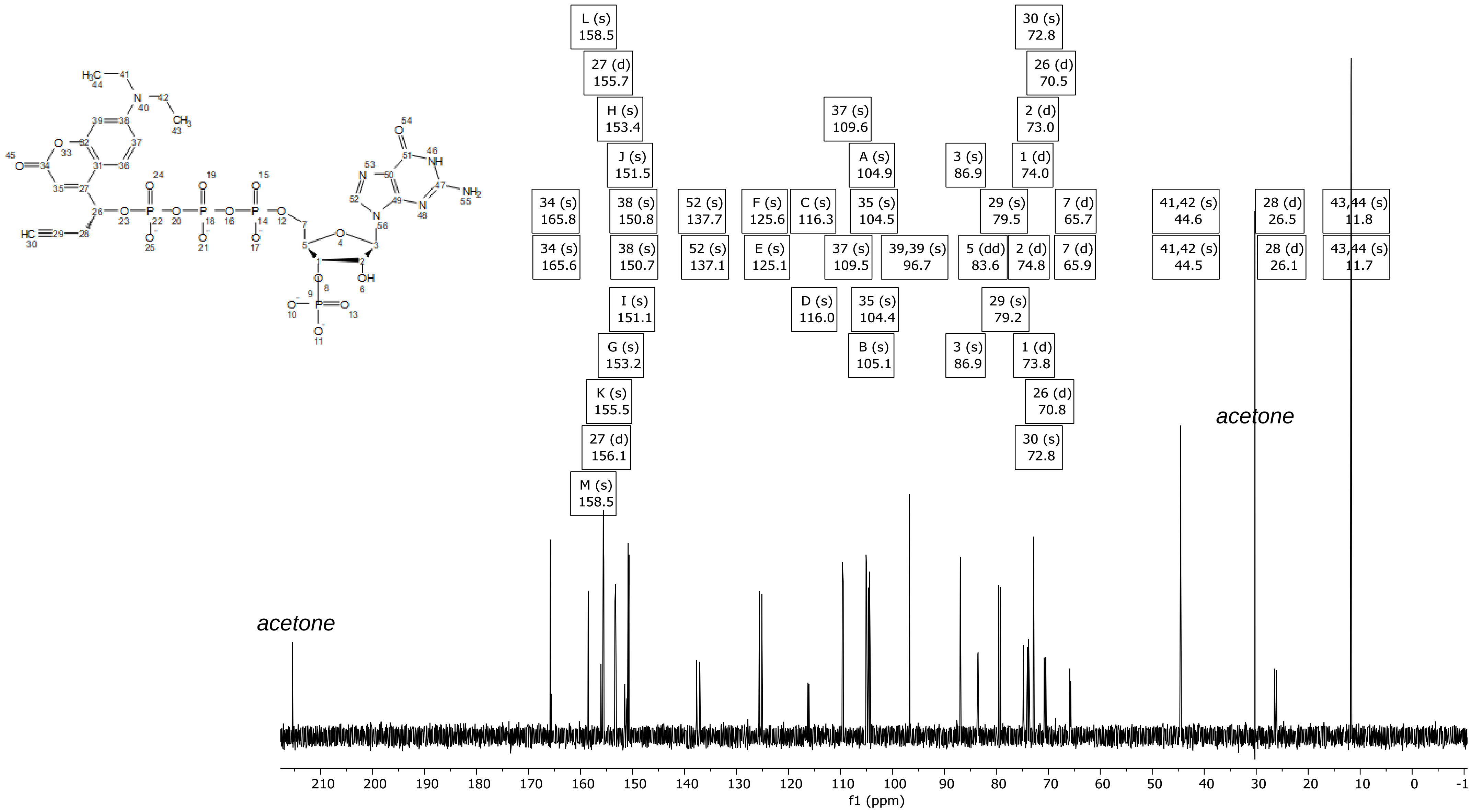

### DQF-COSY of 11

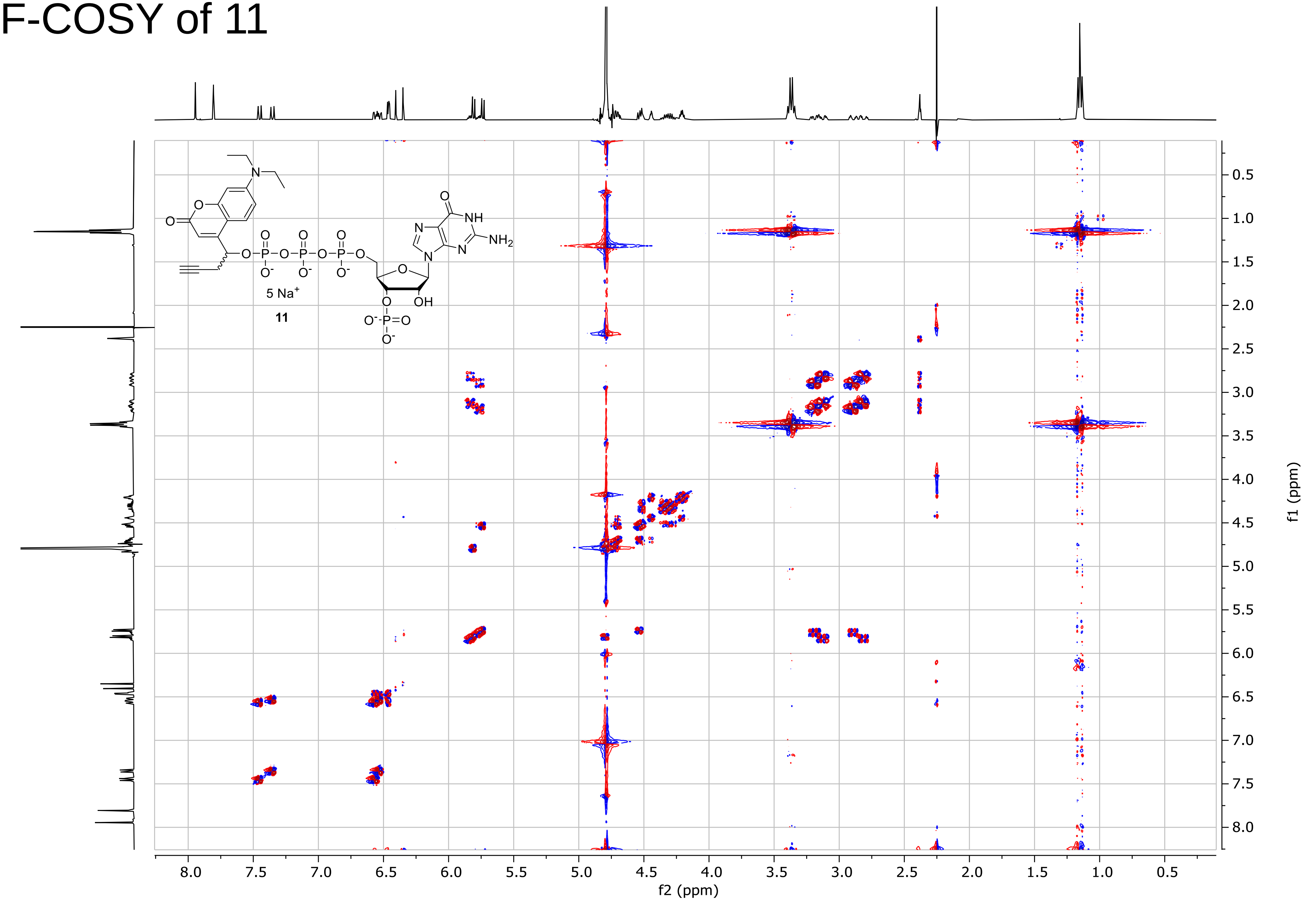

### ed-HSQC of 11

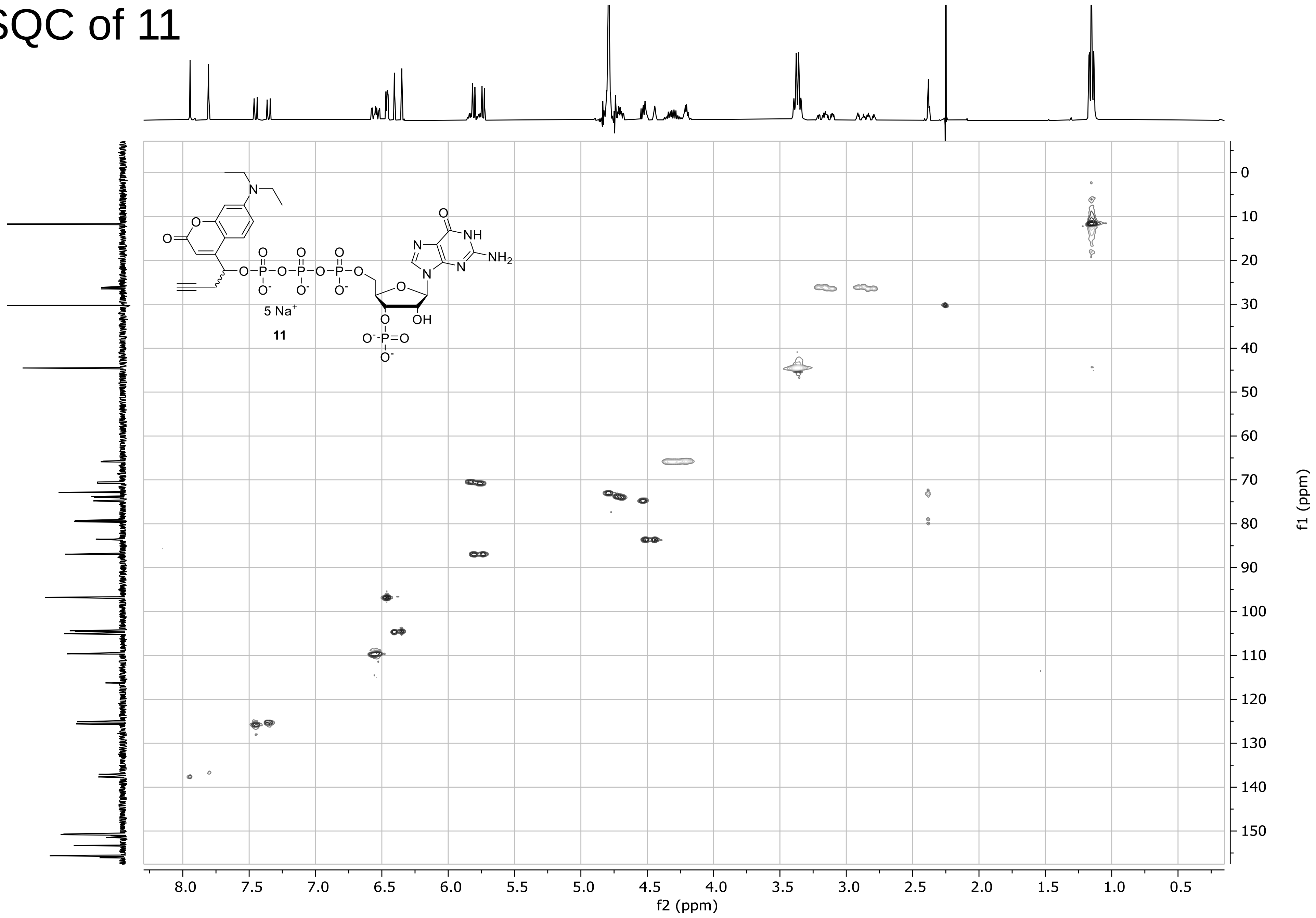

### HMBC of 11

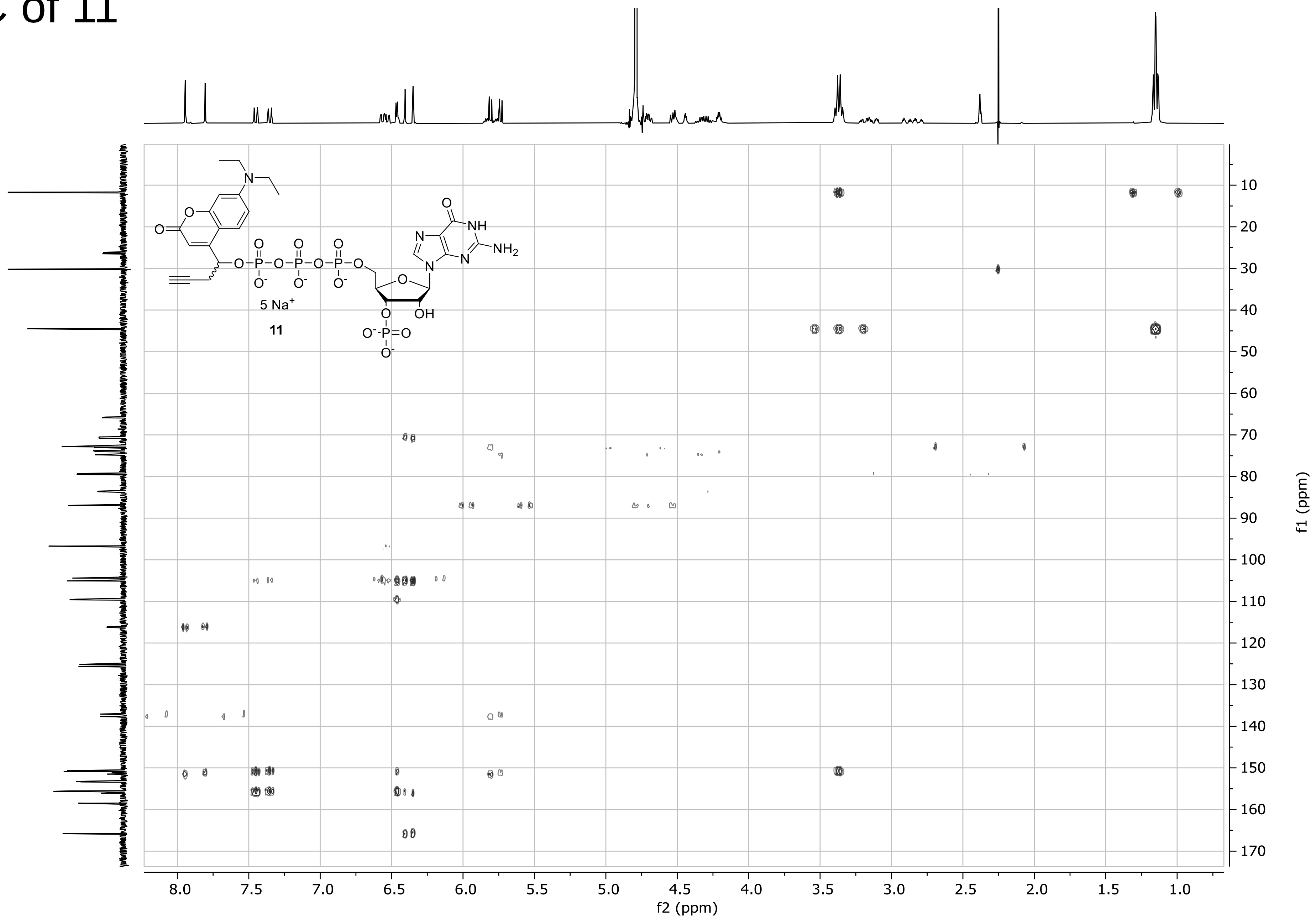

### $^{31}$P-HMBC of 11

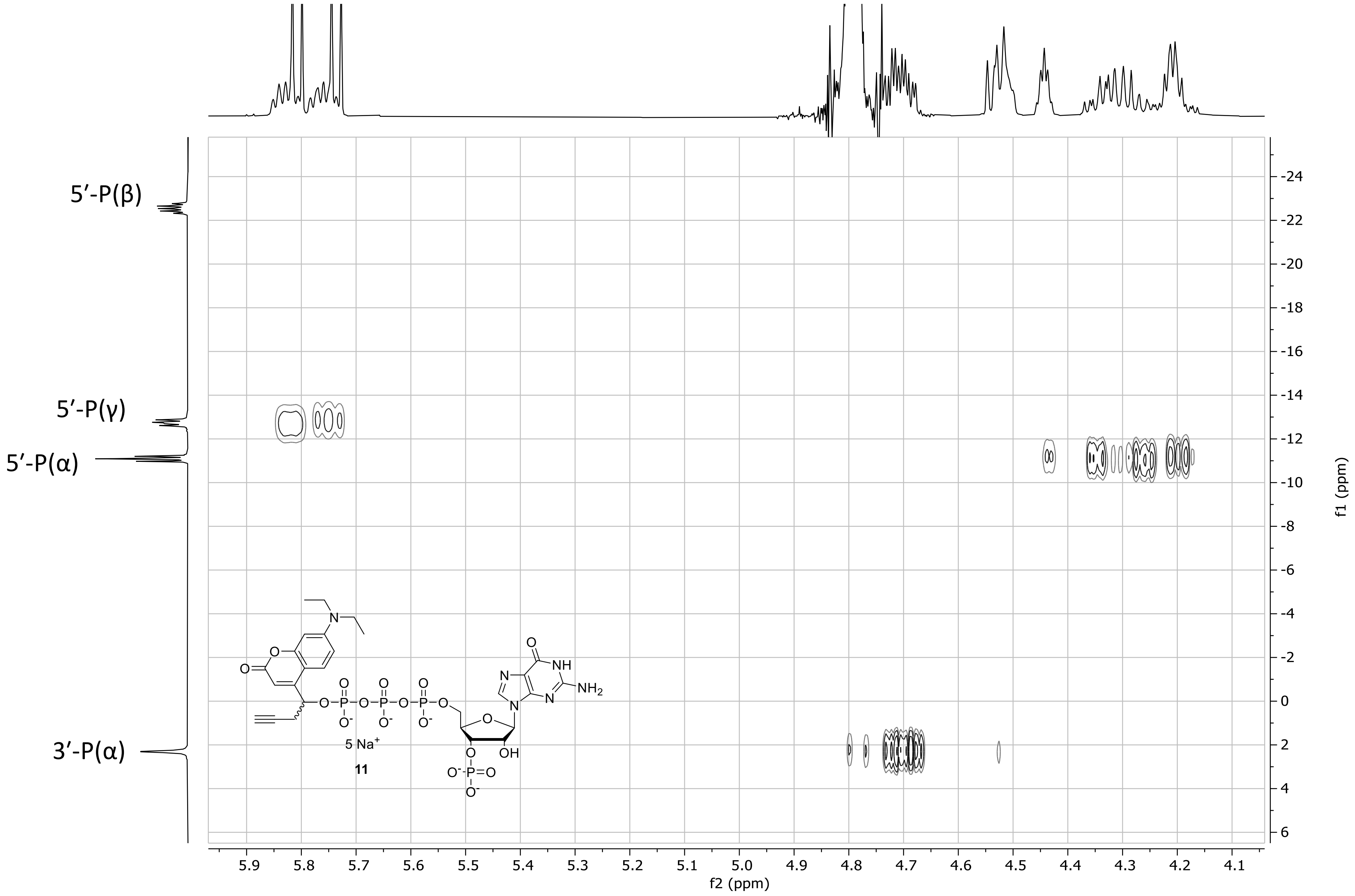

### $^1$H-NMR spectrum of 12

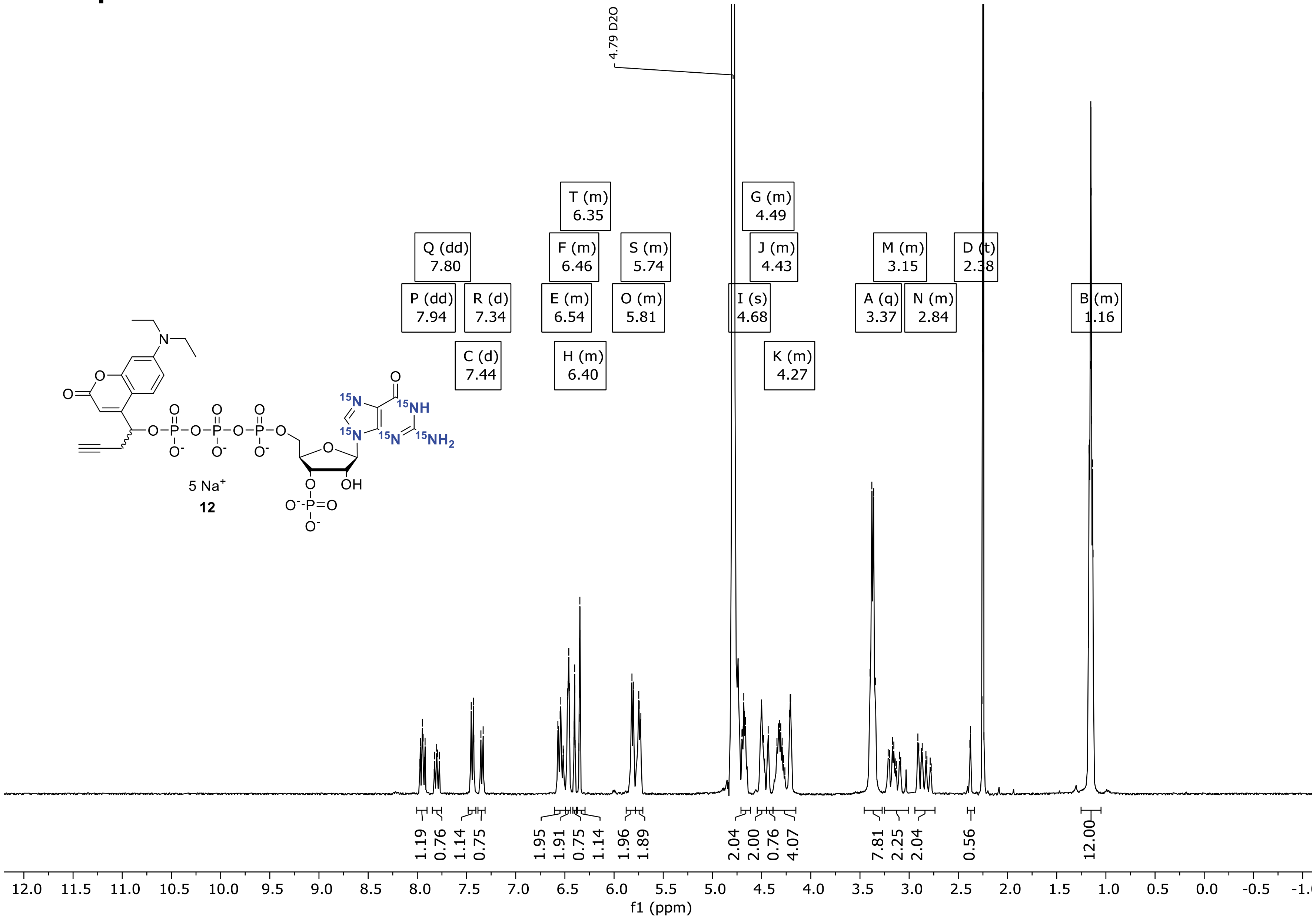

### $^{31}$P-NMR spectrum of 12

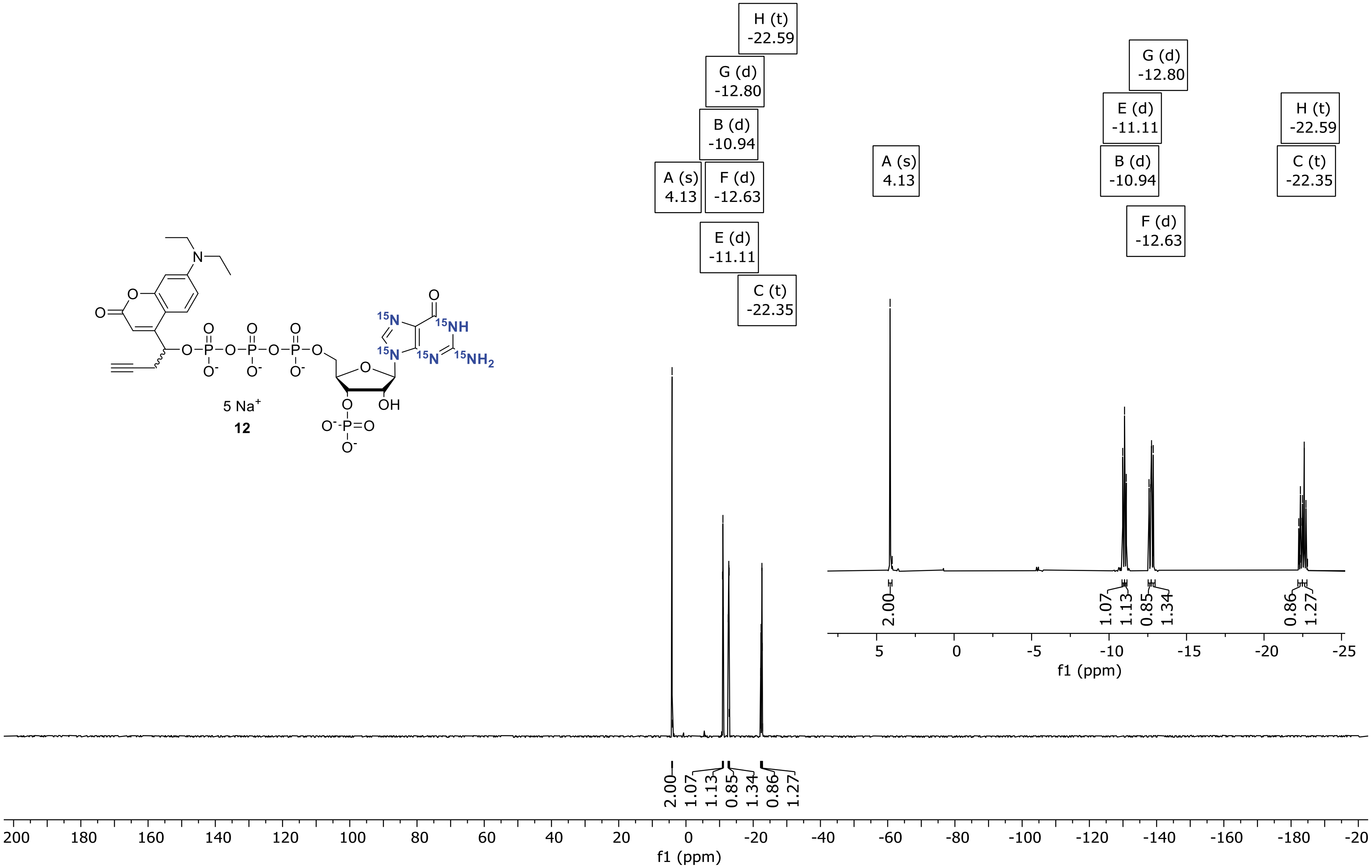

### $^{13}C$ {$^{1}H$}-NMR spectrum of 12

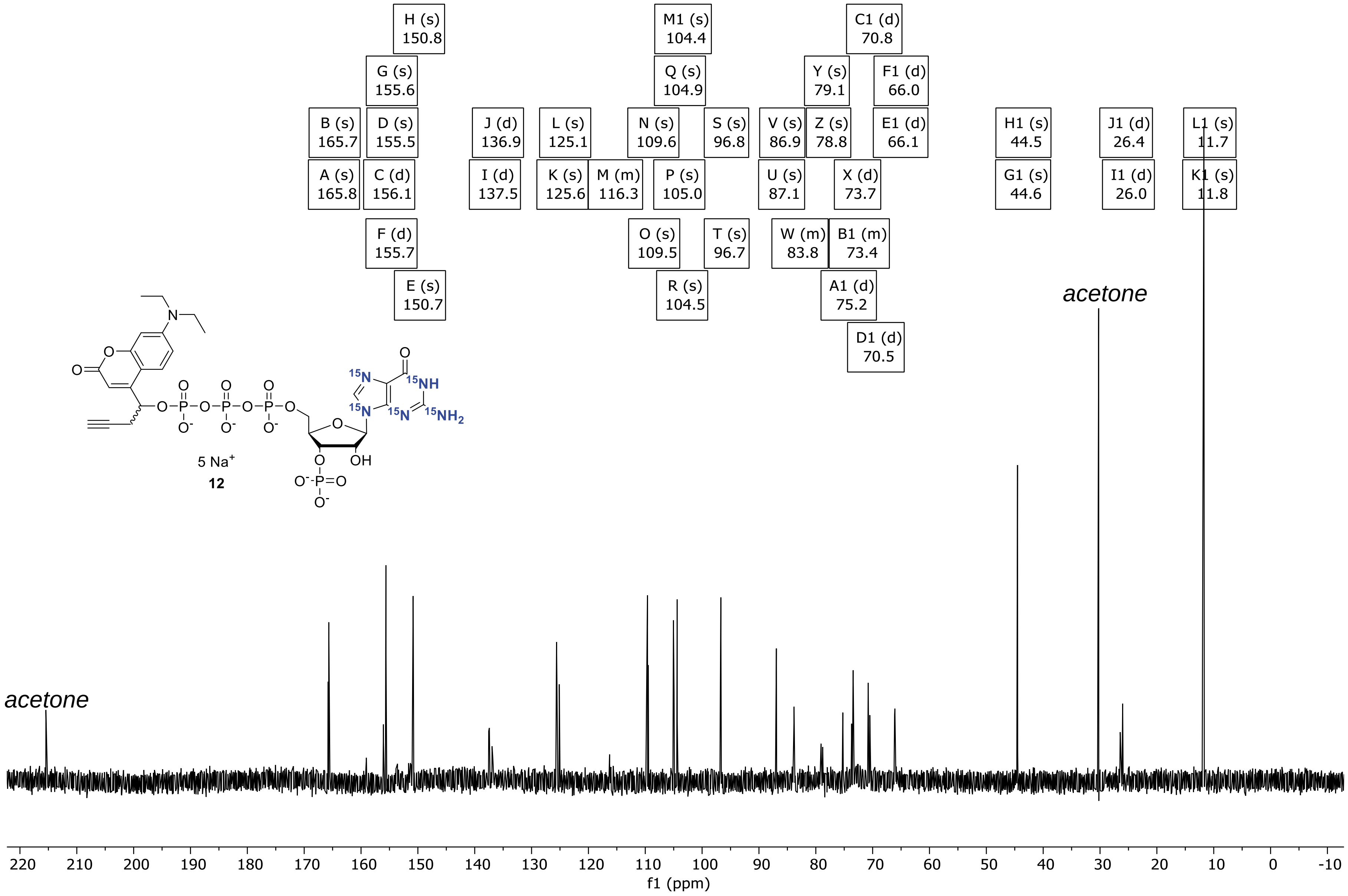

### $^{1}$H-NMR spectrum of 13

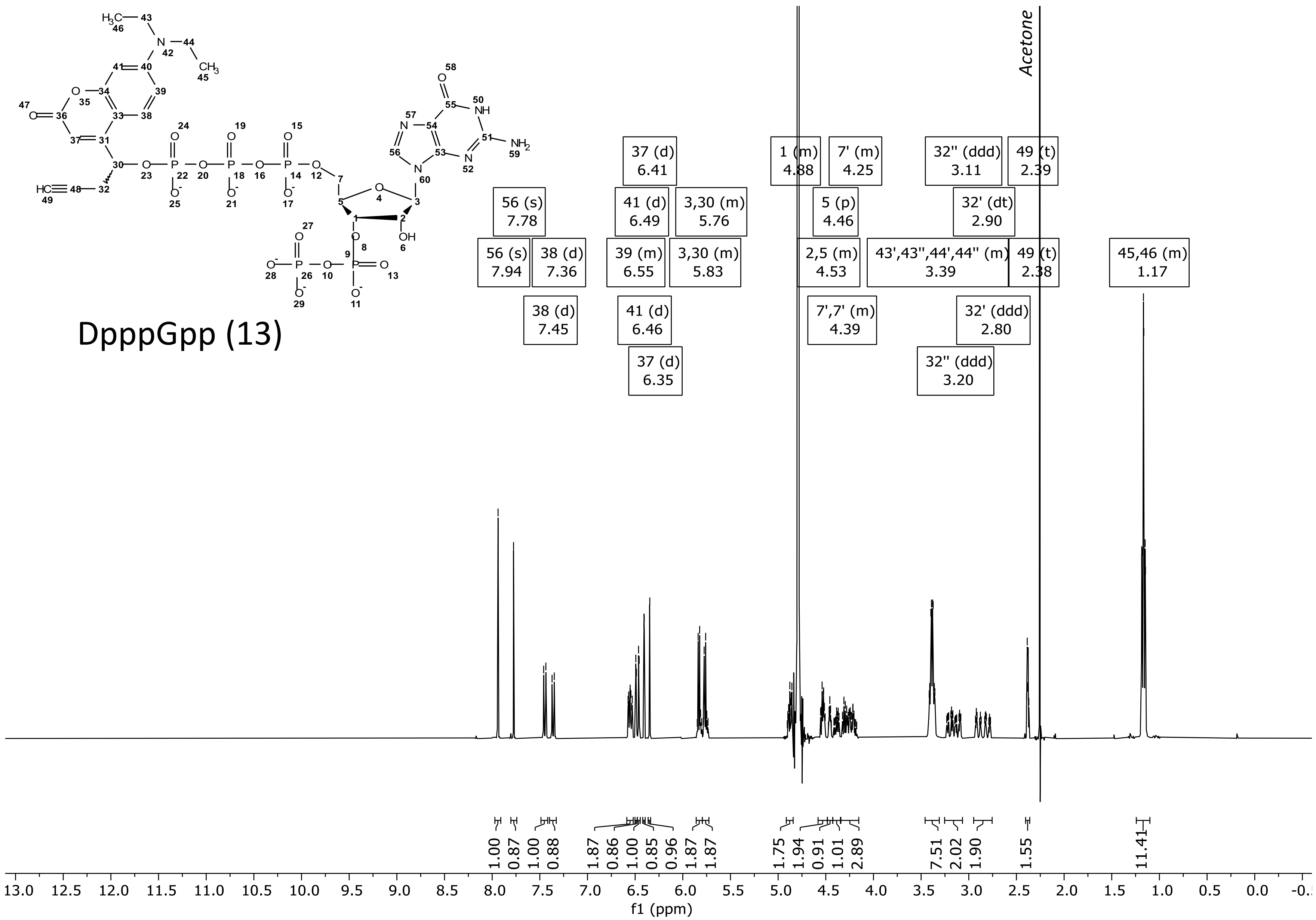

### $^{31}$P-NMR spectrum of 13

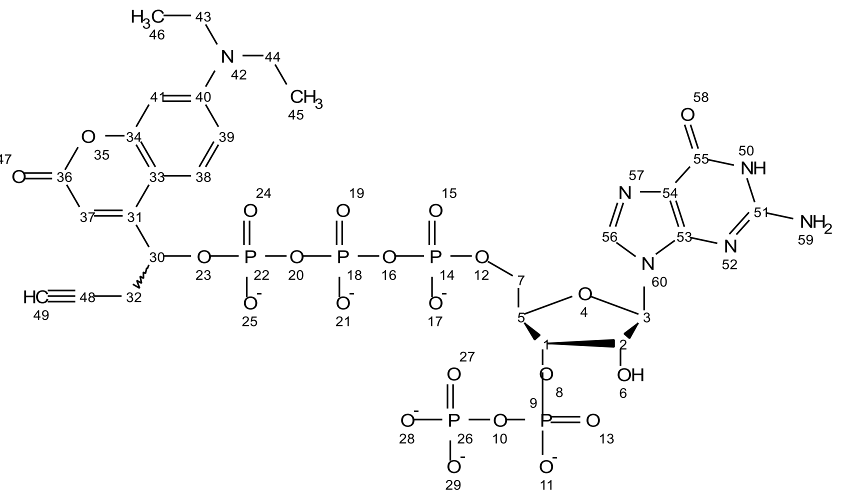

DpppGpp (13)

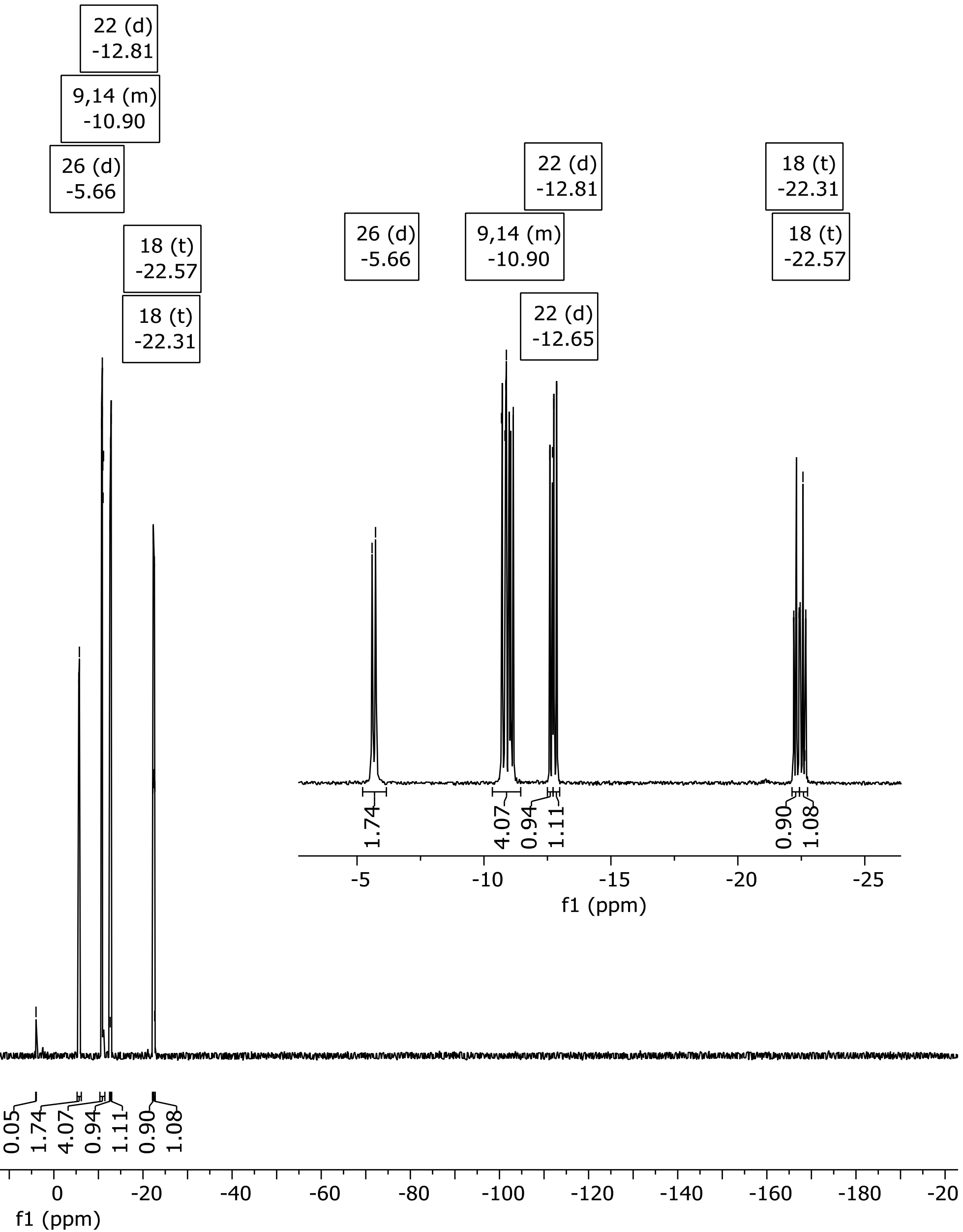

### $^{13}$C-NMR spectrum of 13

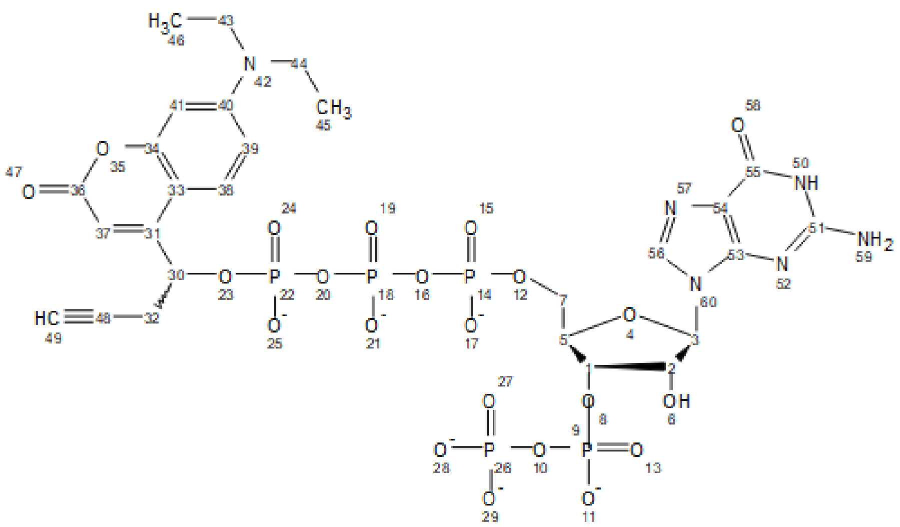

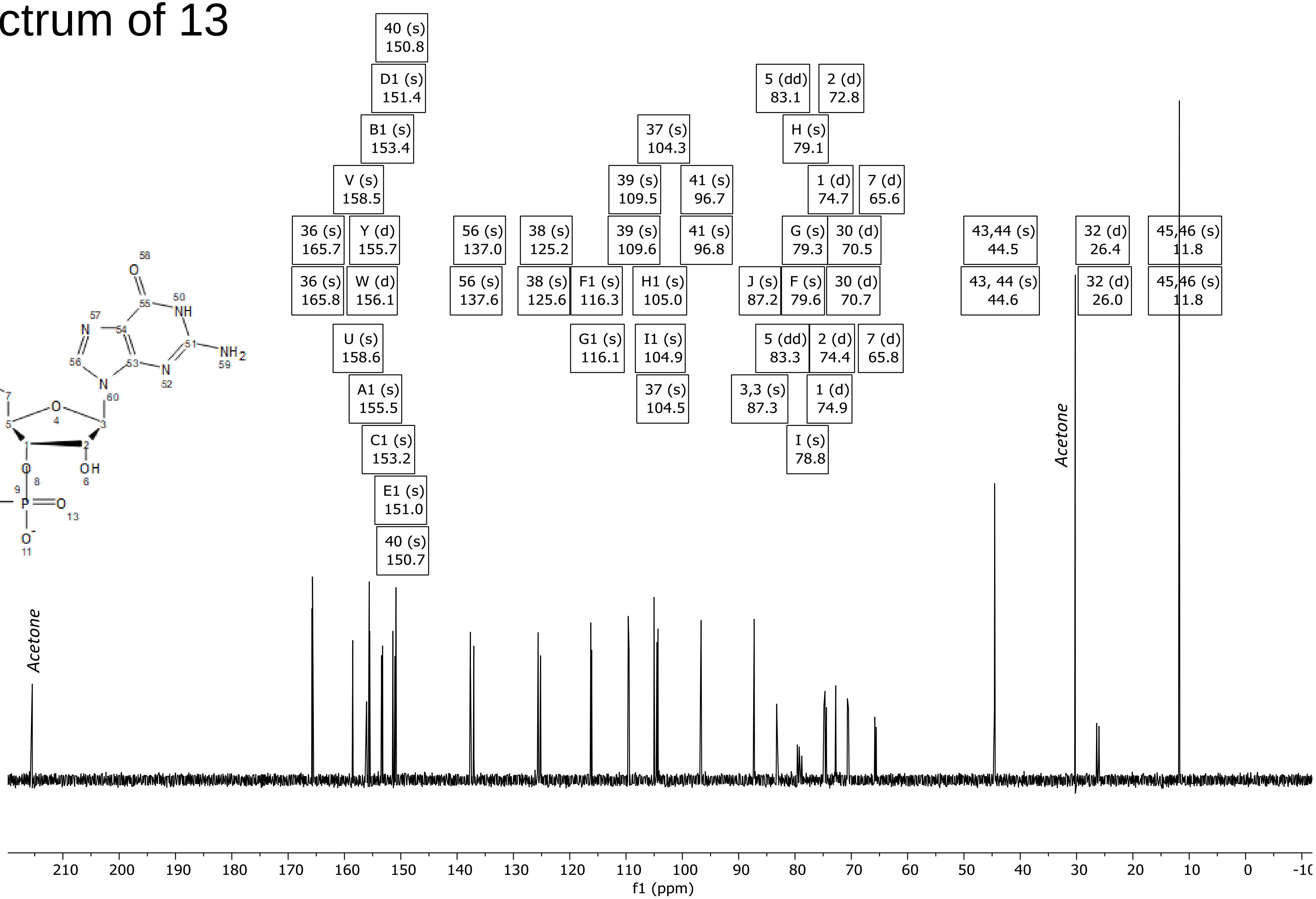

### DQF-COSY of 13

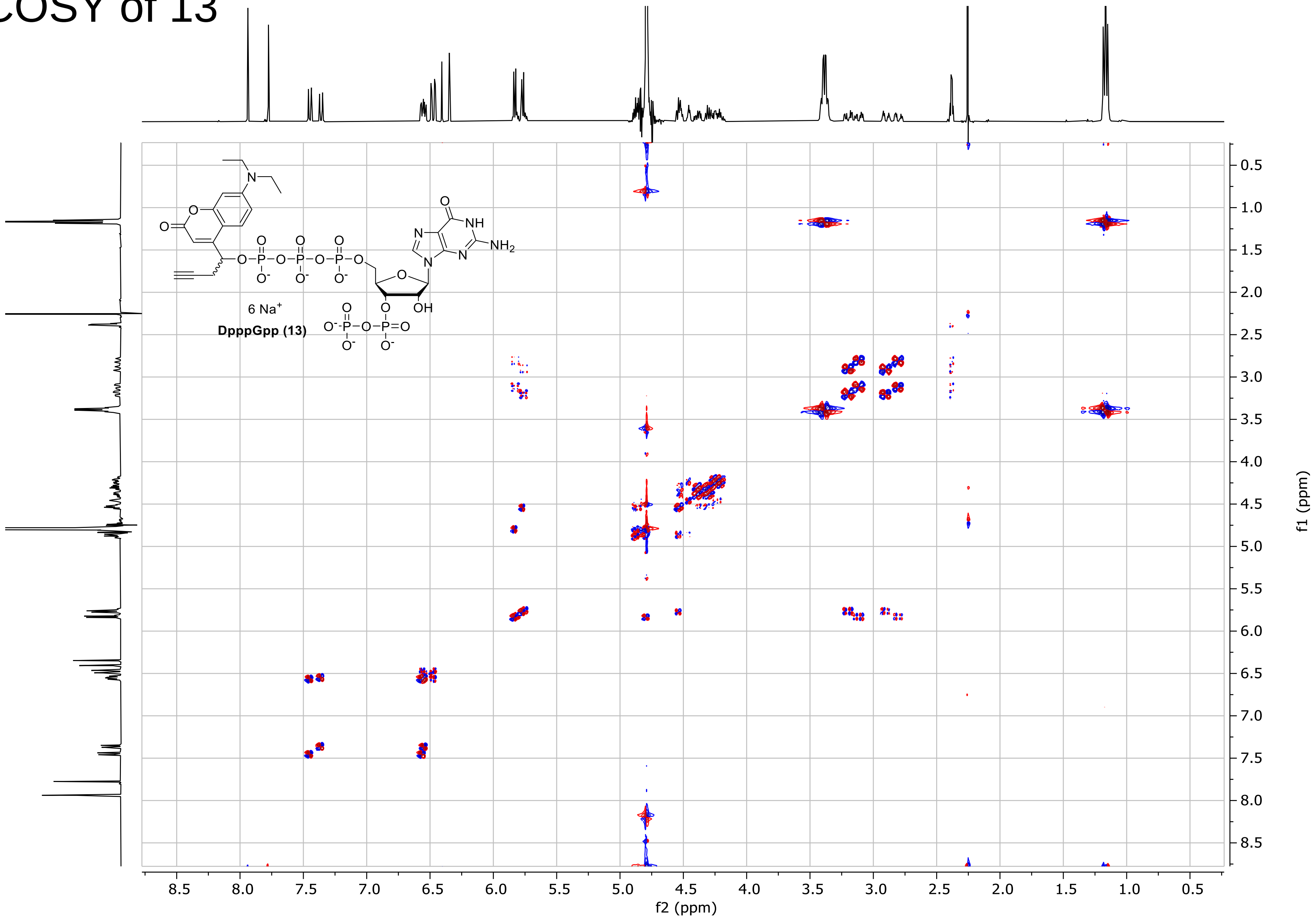

### ed-HSQC of 13

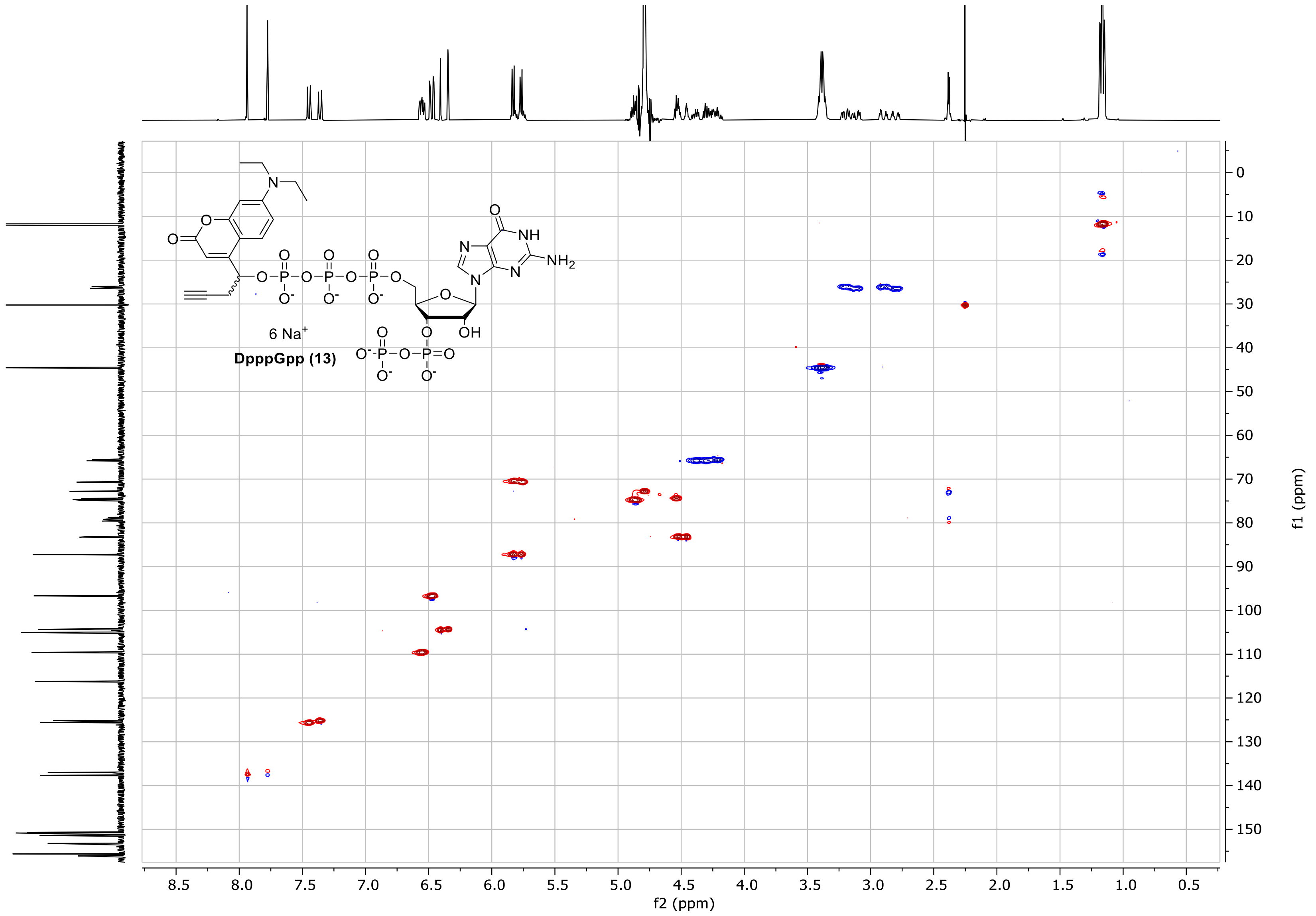

### HMBC of 13

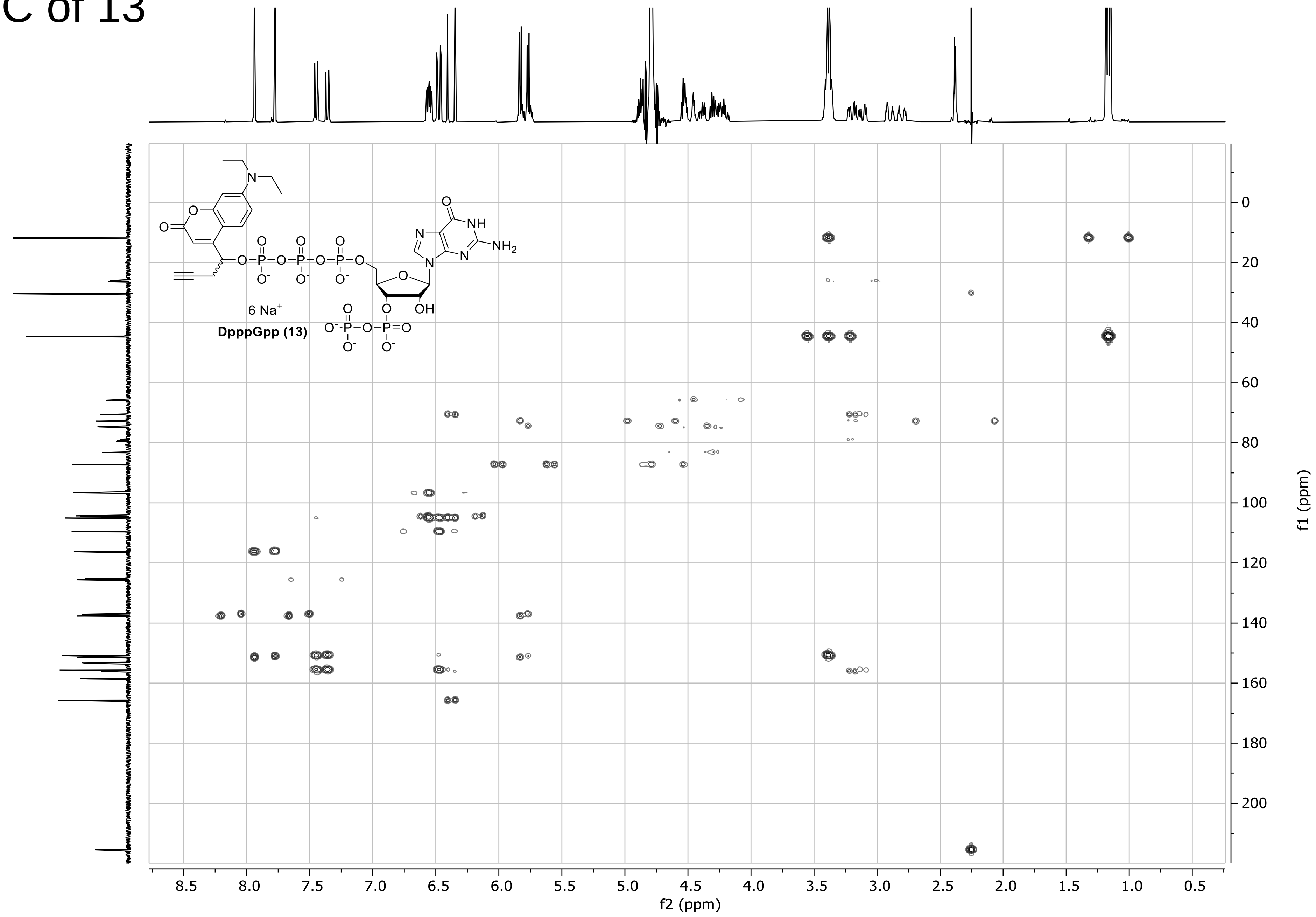

### $^{31}$P-HMBC of 13

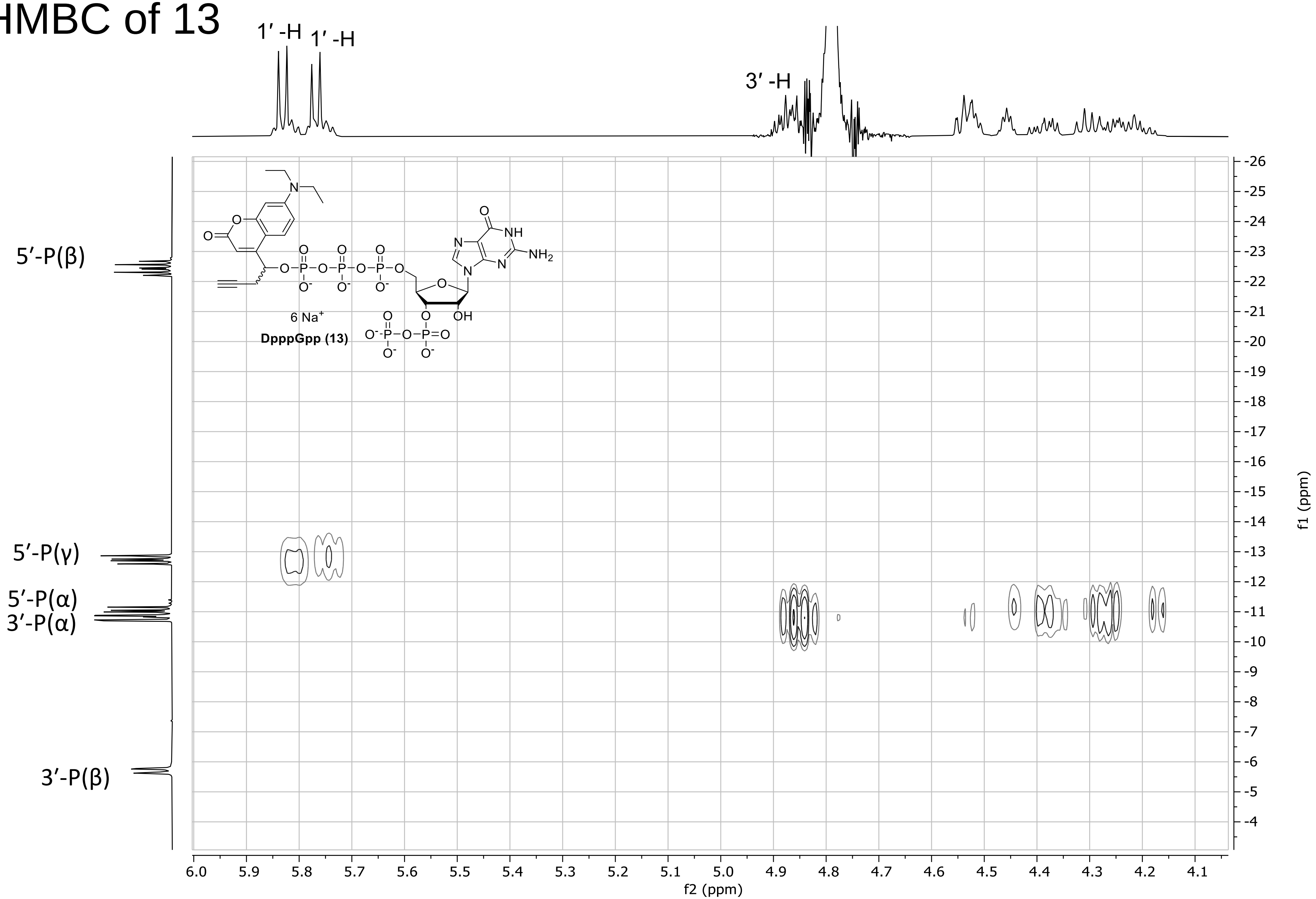

### $^{31}$P-COSY of 13

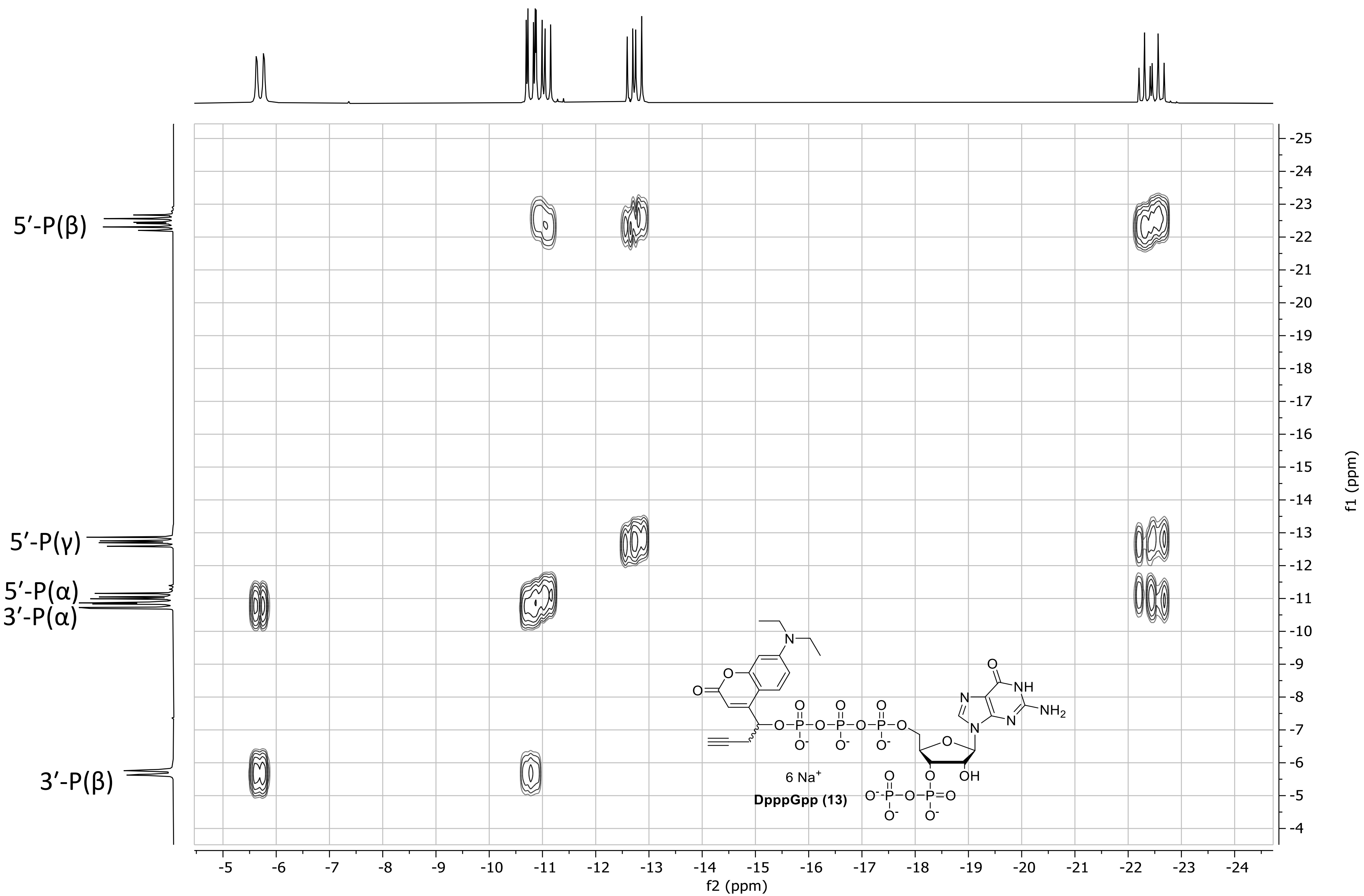

### $^{1}$H- NMR spectrum of 14

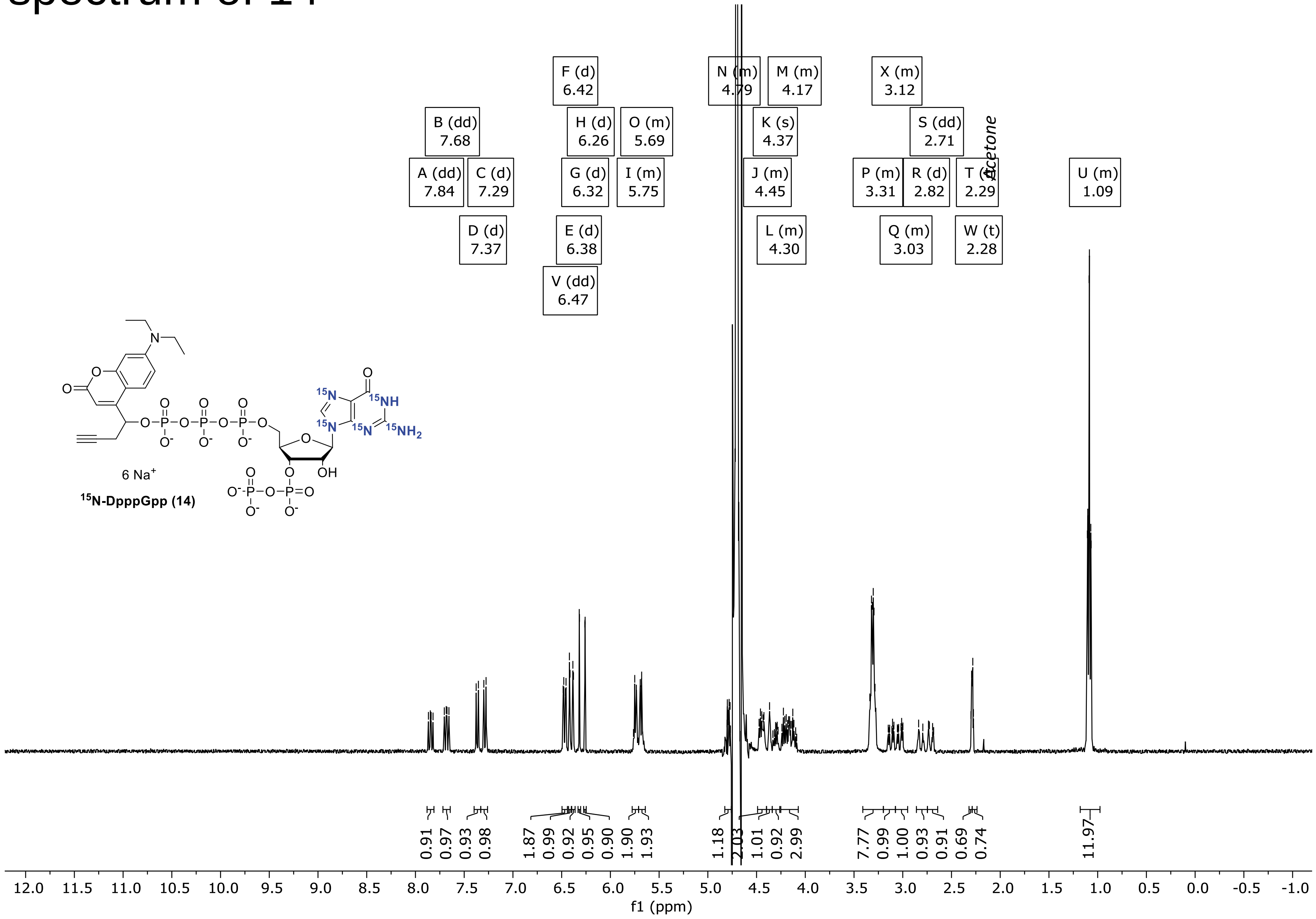

### $^{31}P\{^1H\}$-NMR spectrum of 14

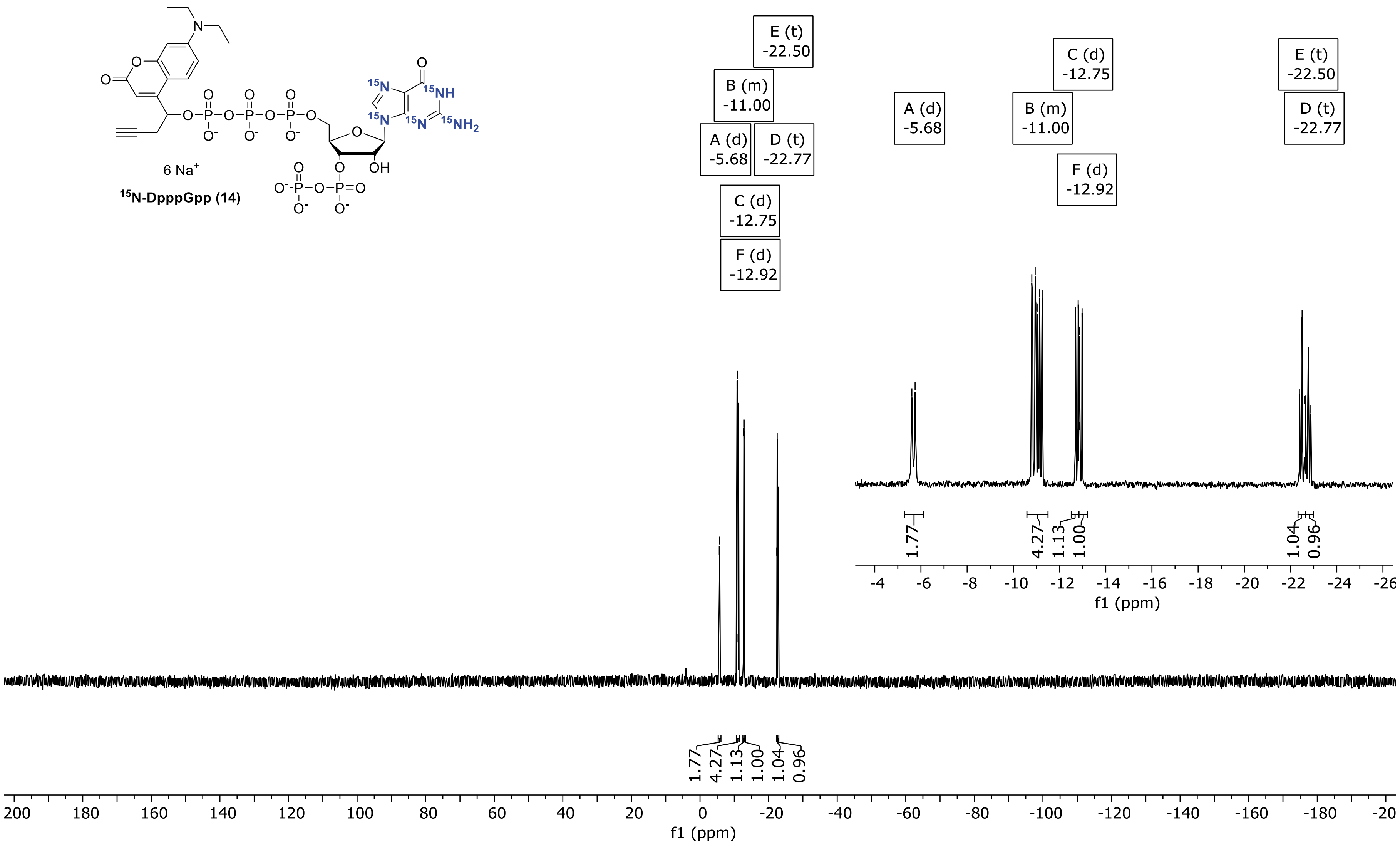

### $^{13}C$ {$^{1}H$}-NMR spectrum of 14

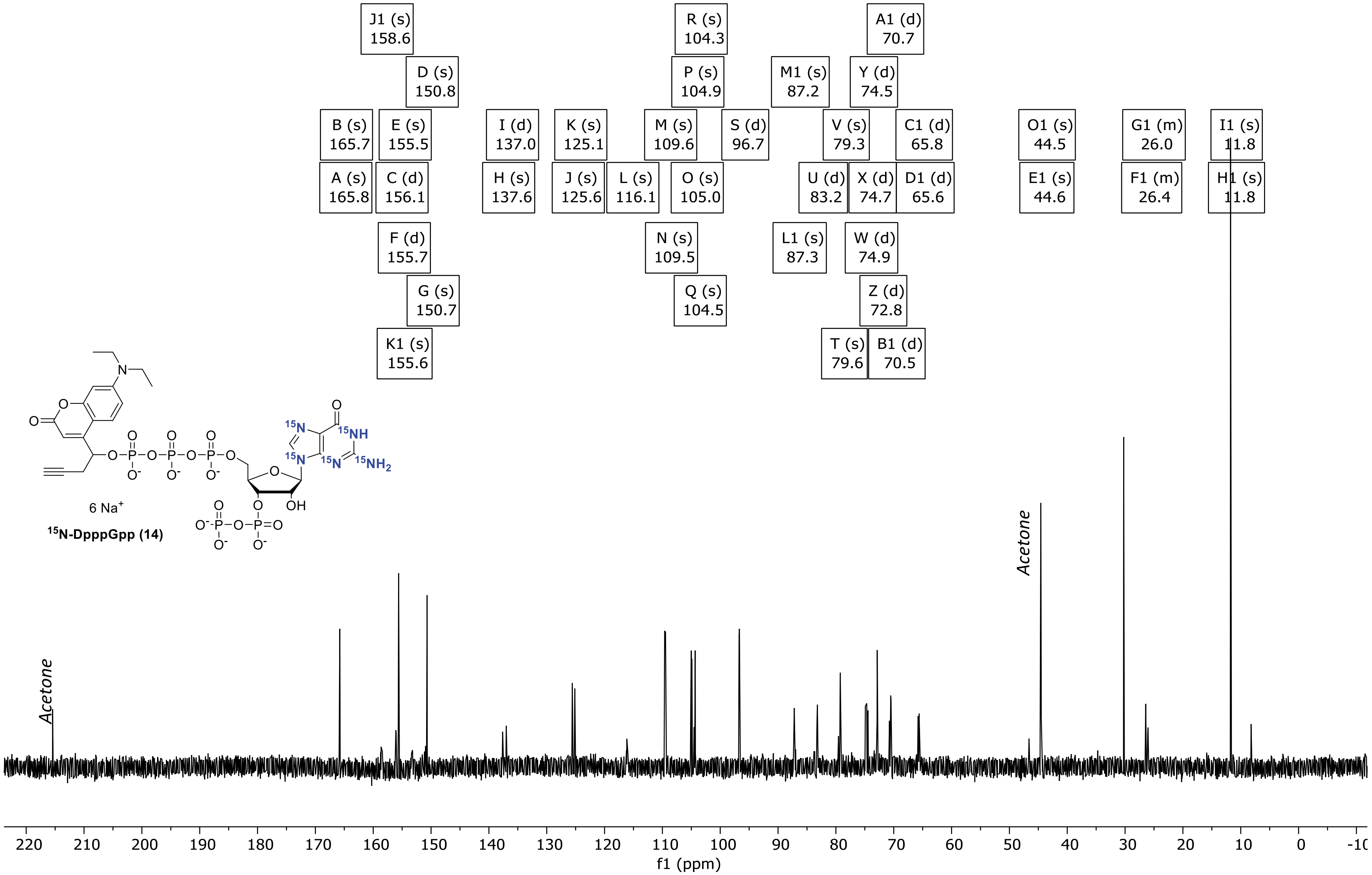

### $^1$H-NMR spectrum of 15

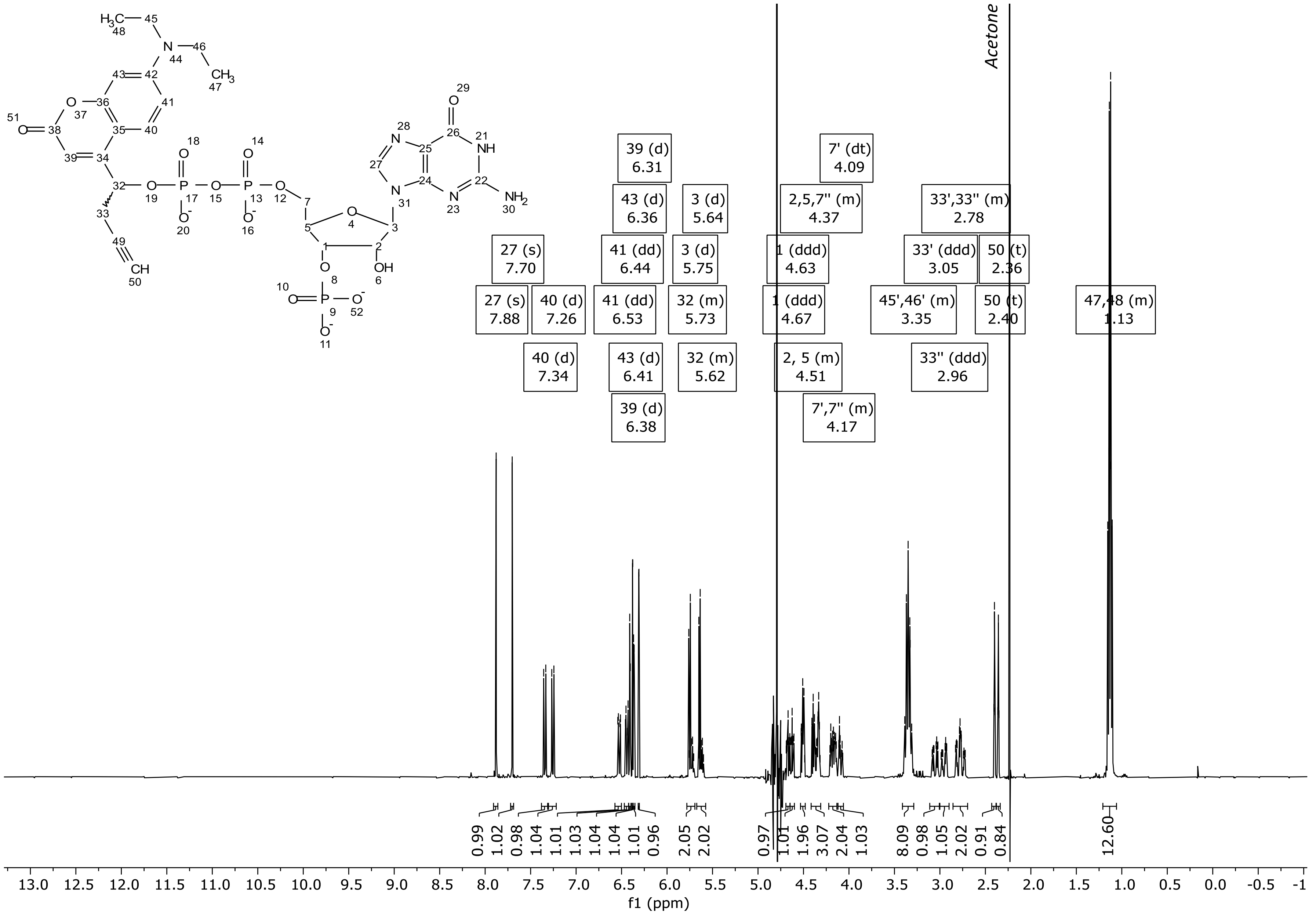

### $^{31}P\{^{1}H\}$ NMR spectrum of 15

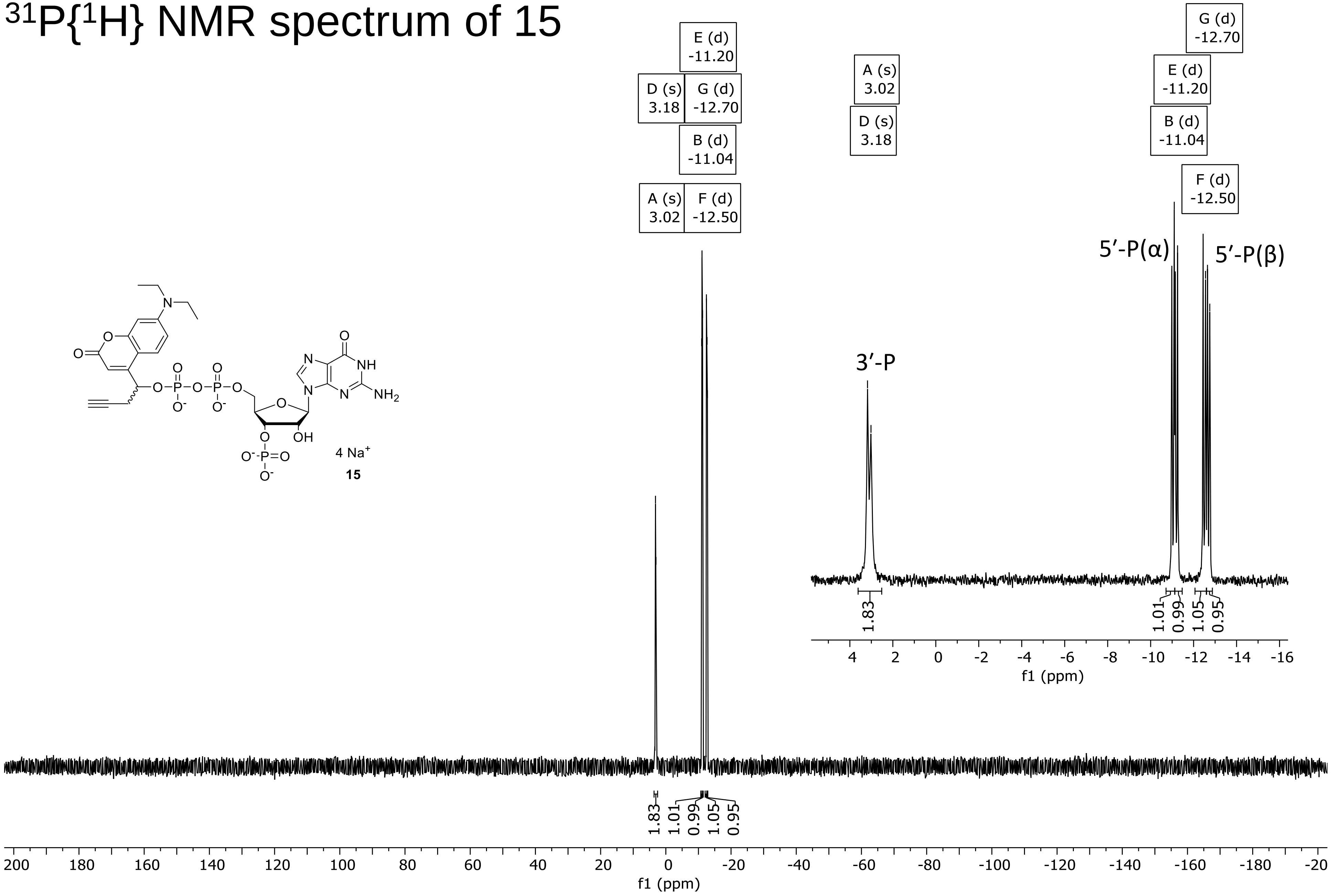

### $^{13}C\ \{^{1}H\}$-NMR spectrum of 15

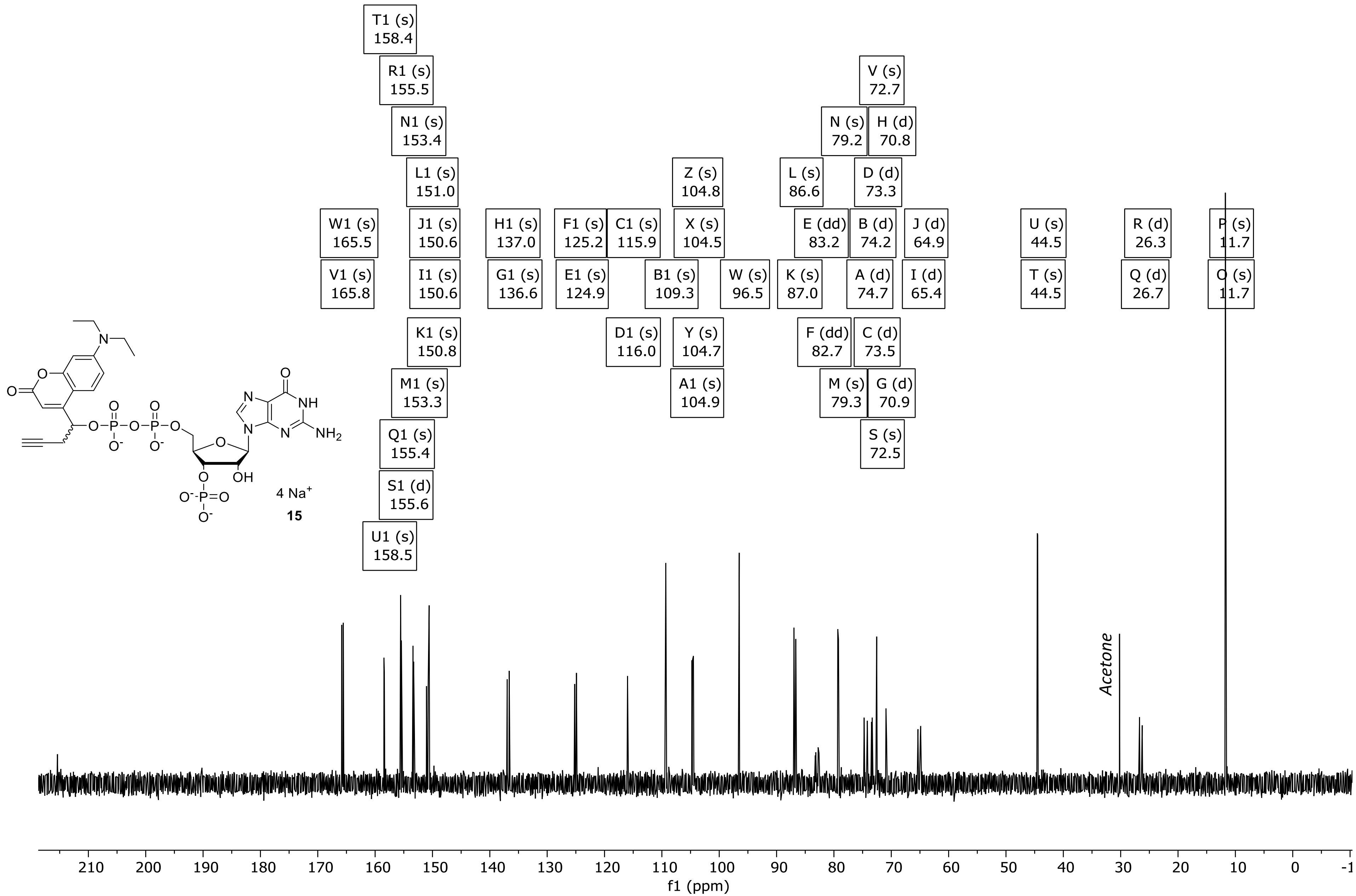

**15**

### DQF-COSY of 15

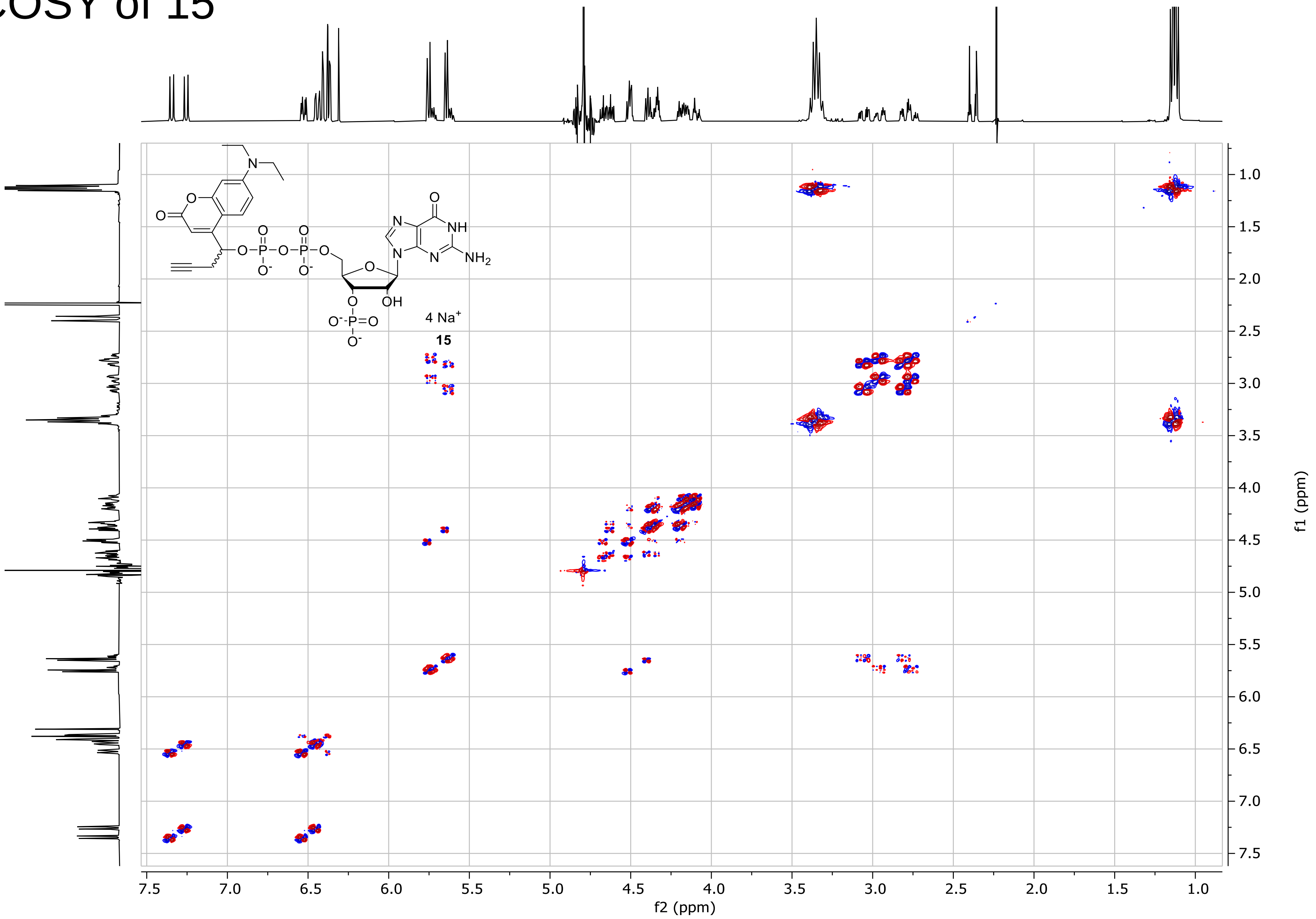

### DQF-COSY of 15

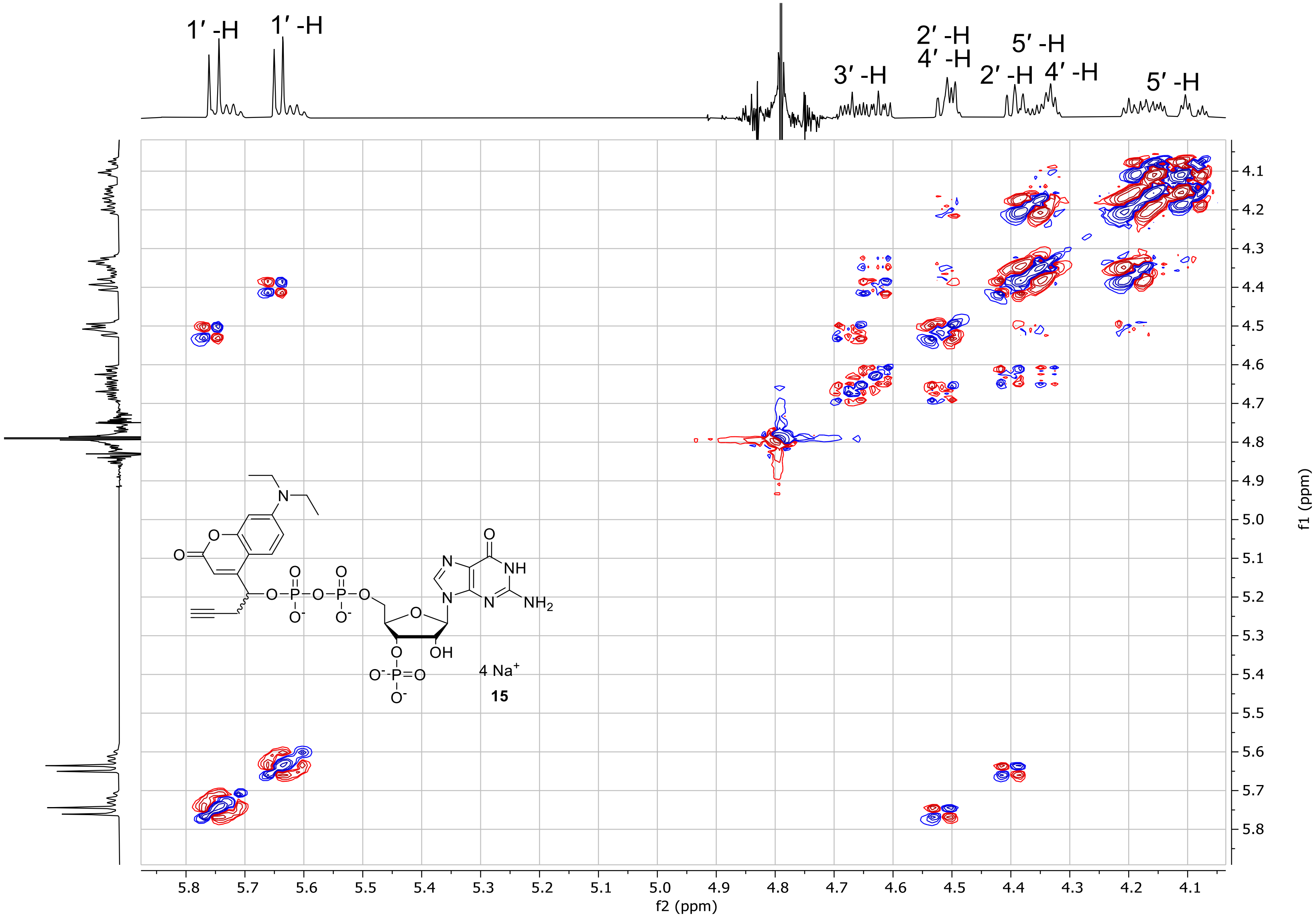

### HSQC of 15

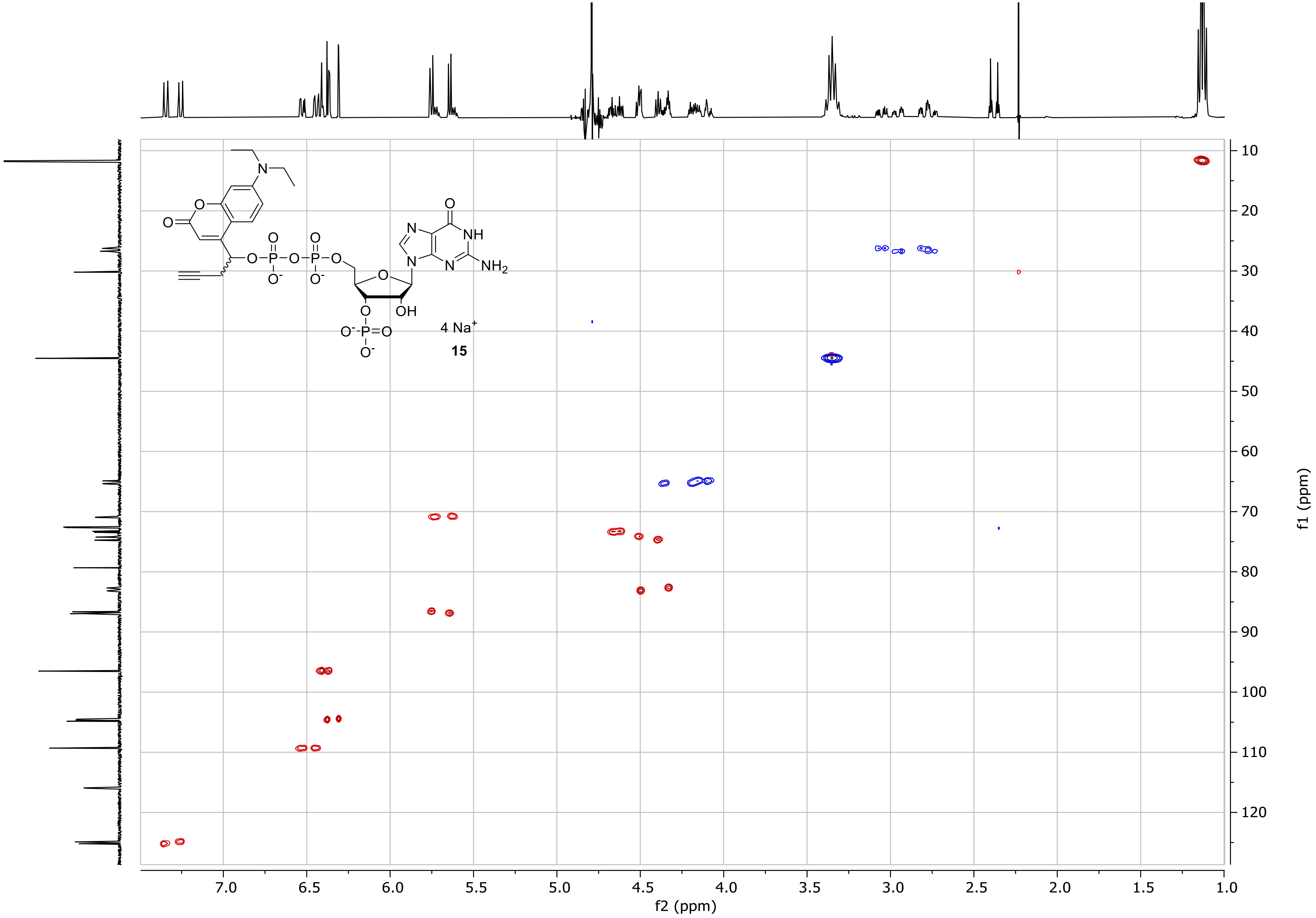

### HMBC of 15

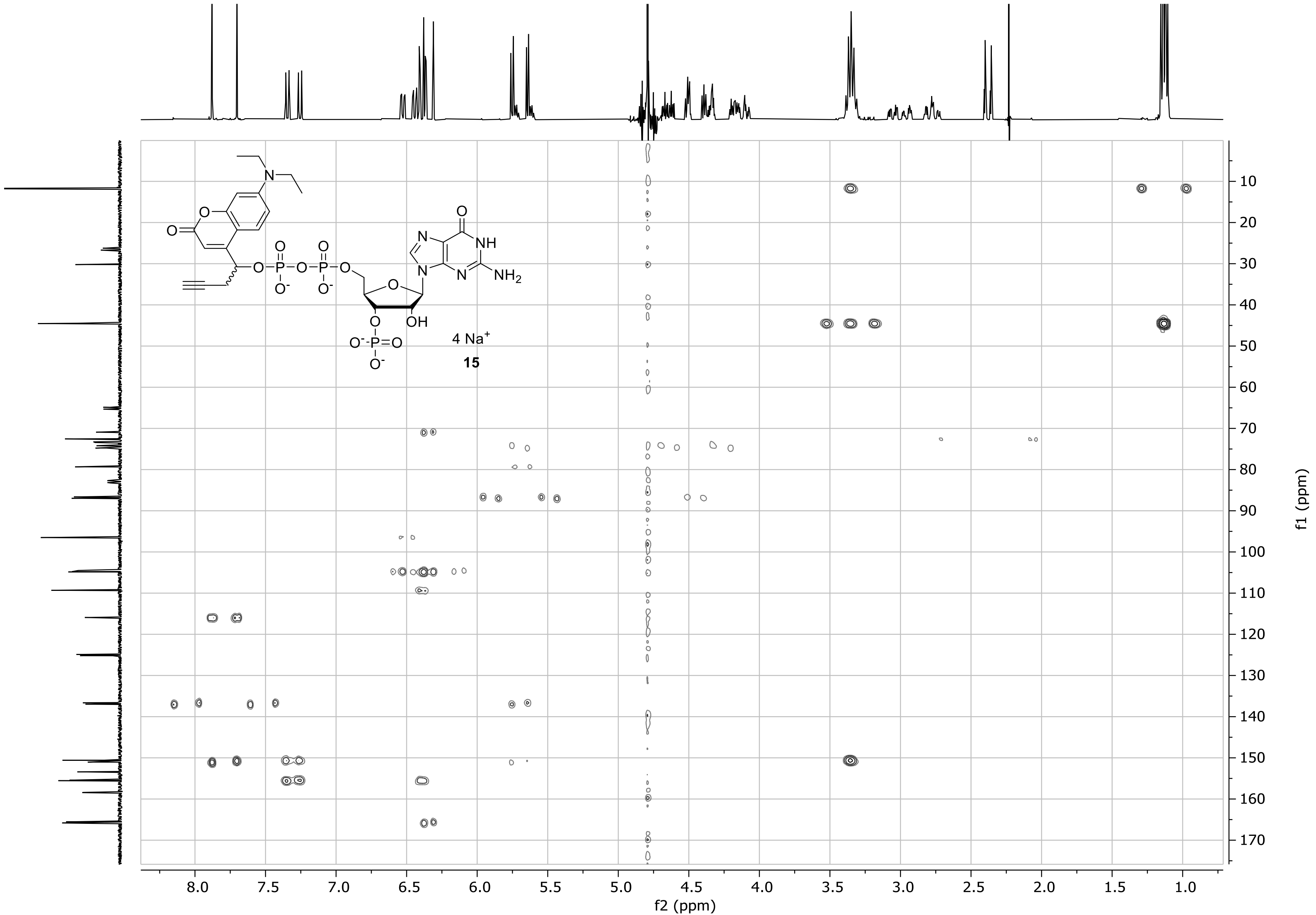

### $^{31}$P-HMBC of 15

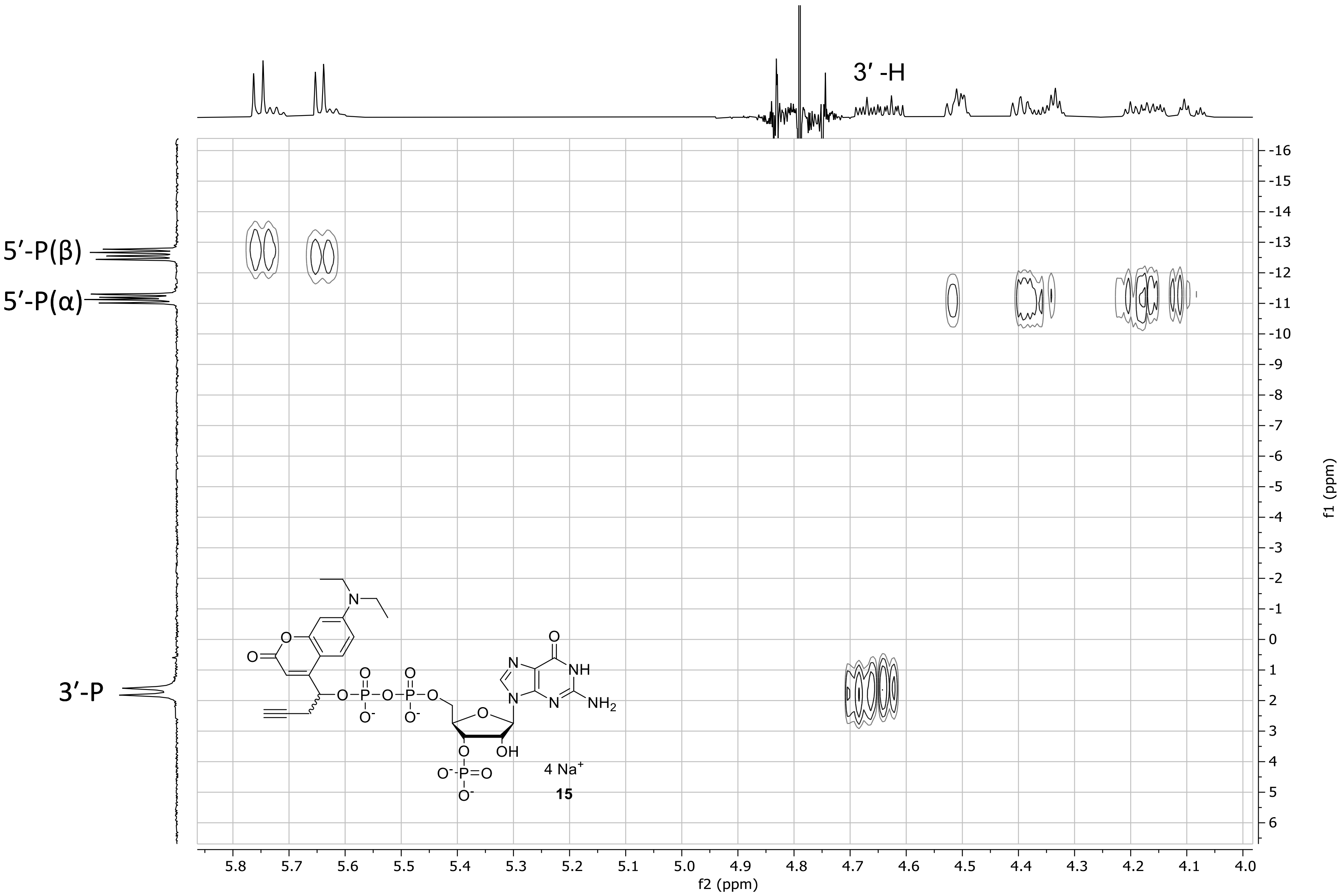

### $^{1}$H-NMR spectrum of 16

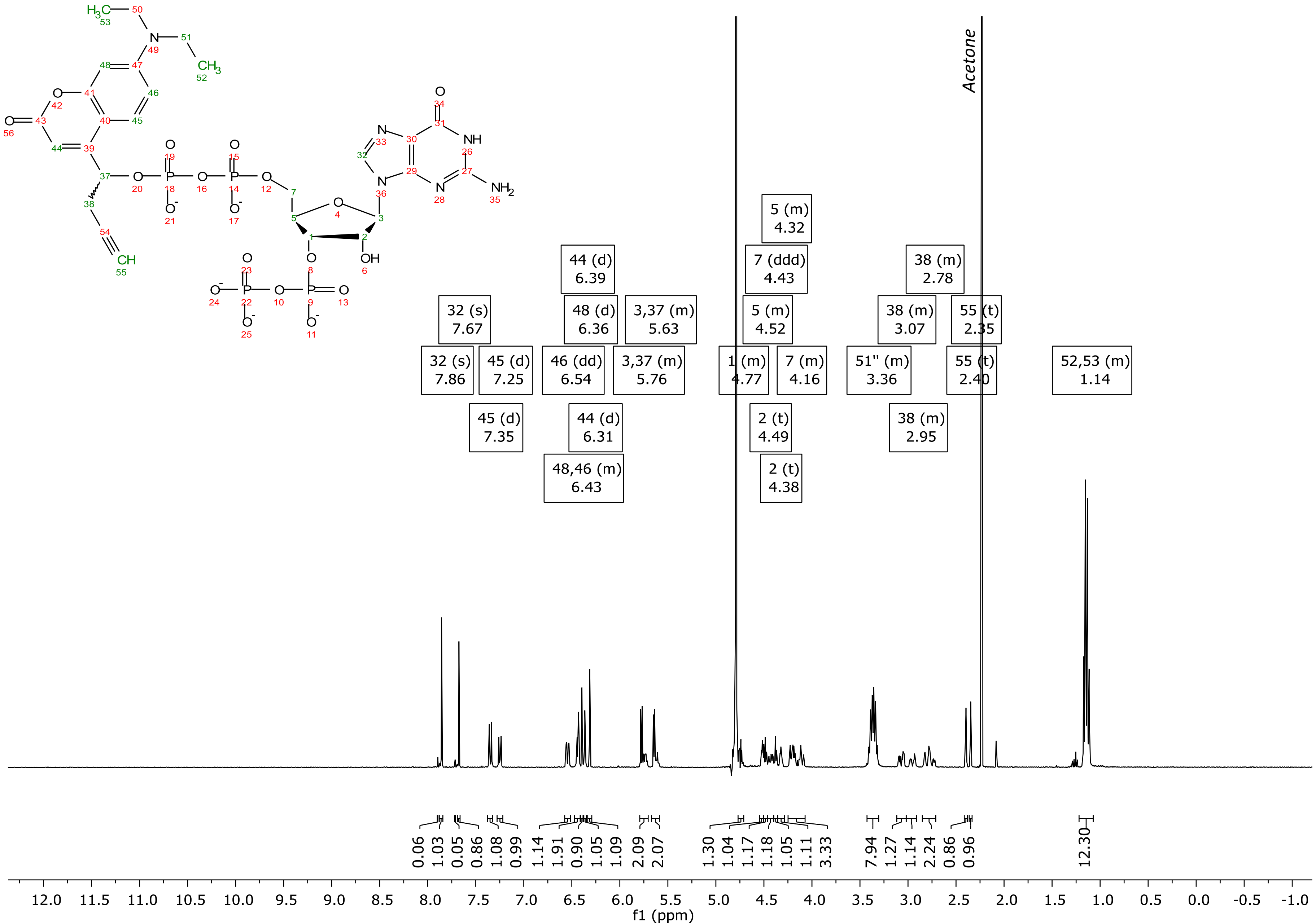

### $^{31}P\{^{1}H\}$ spectrum NMR of 16

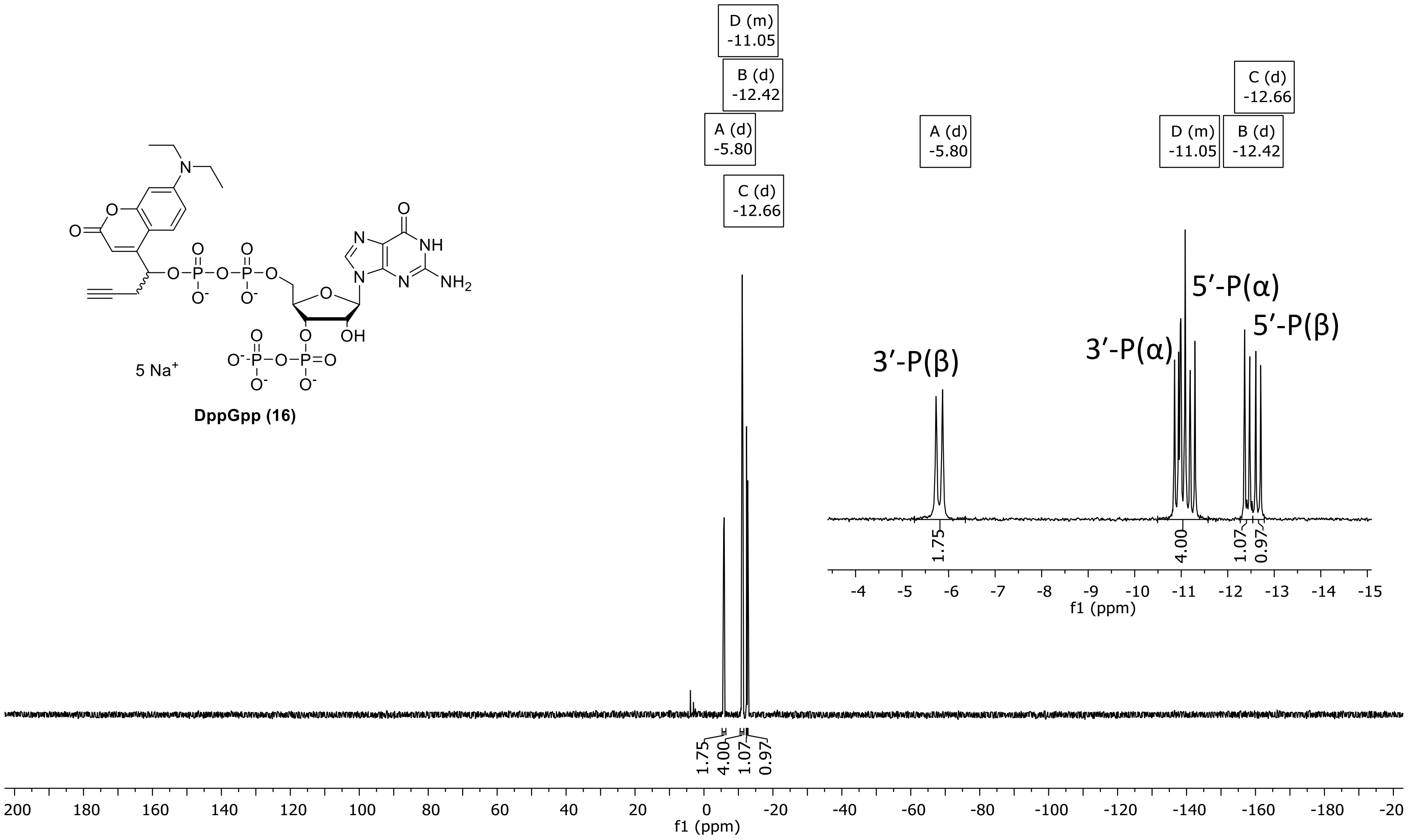

### $^{13}C$ {$^{1}H$} -NMR spectrum of 16

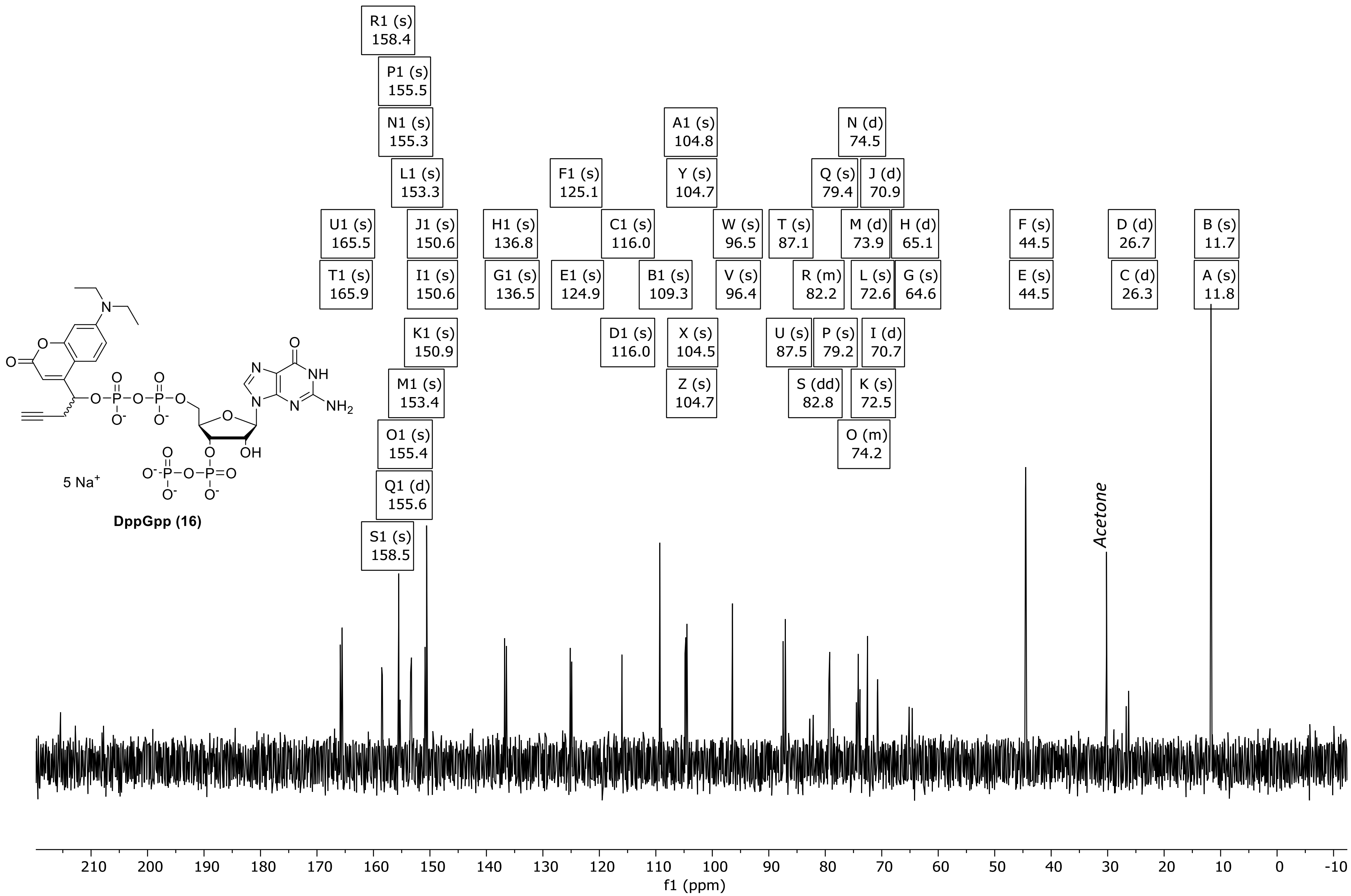

### DQF-COSY of 16

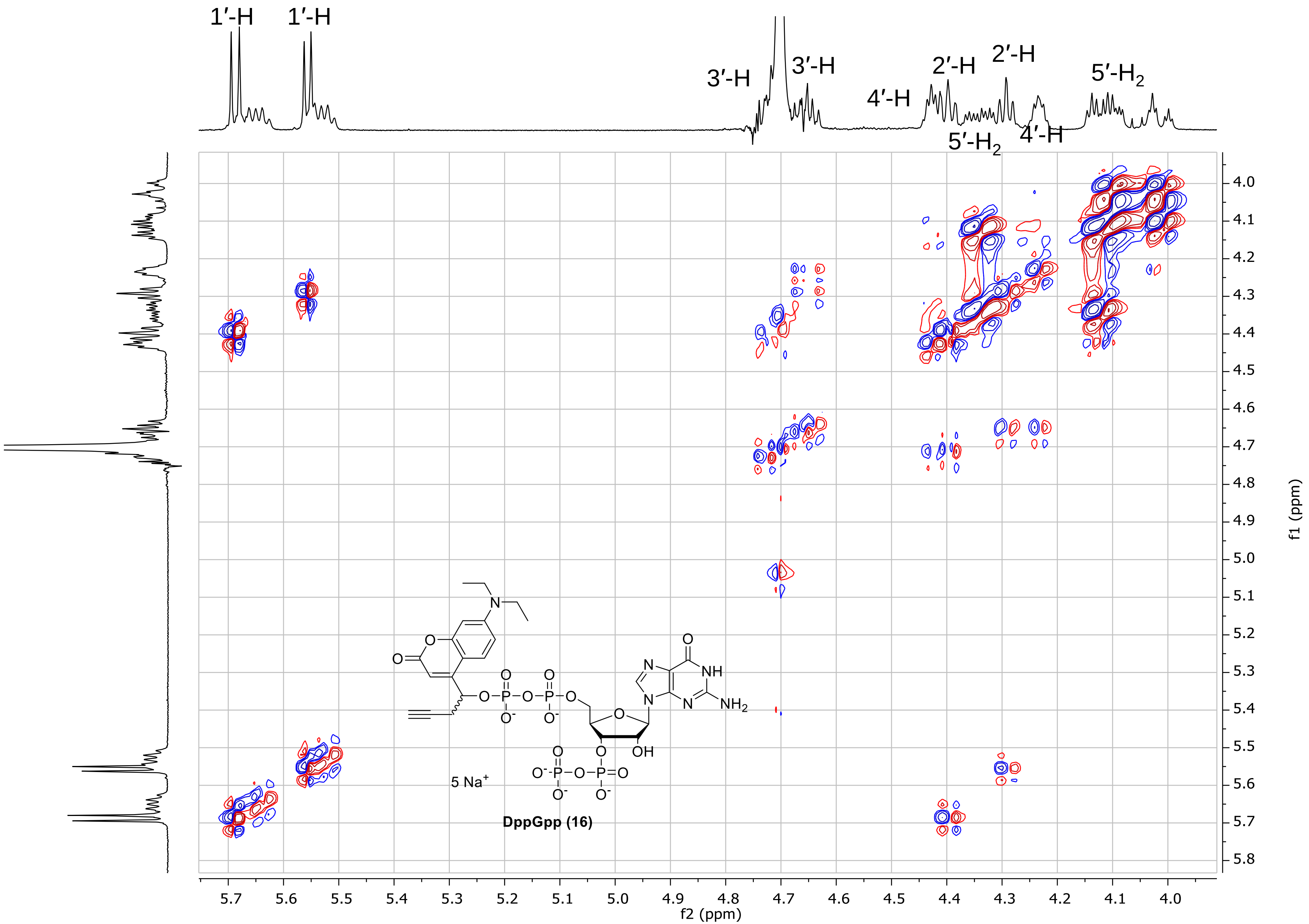

### HSQC of 16

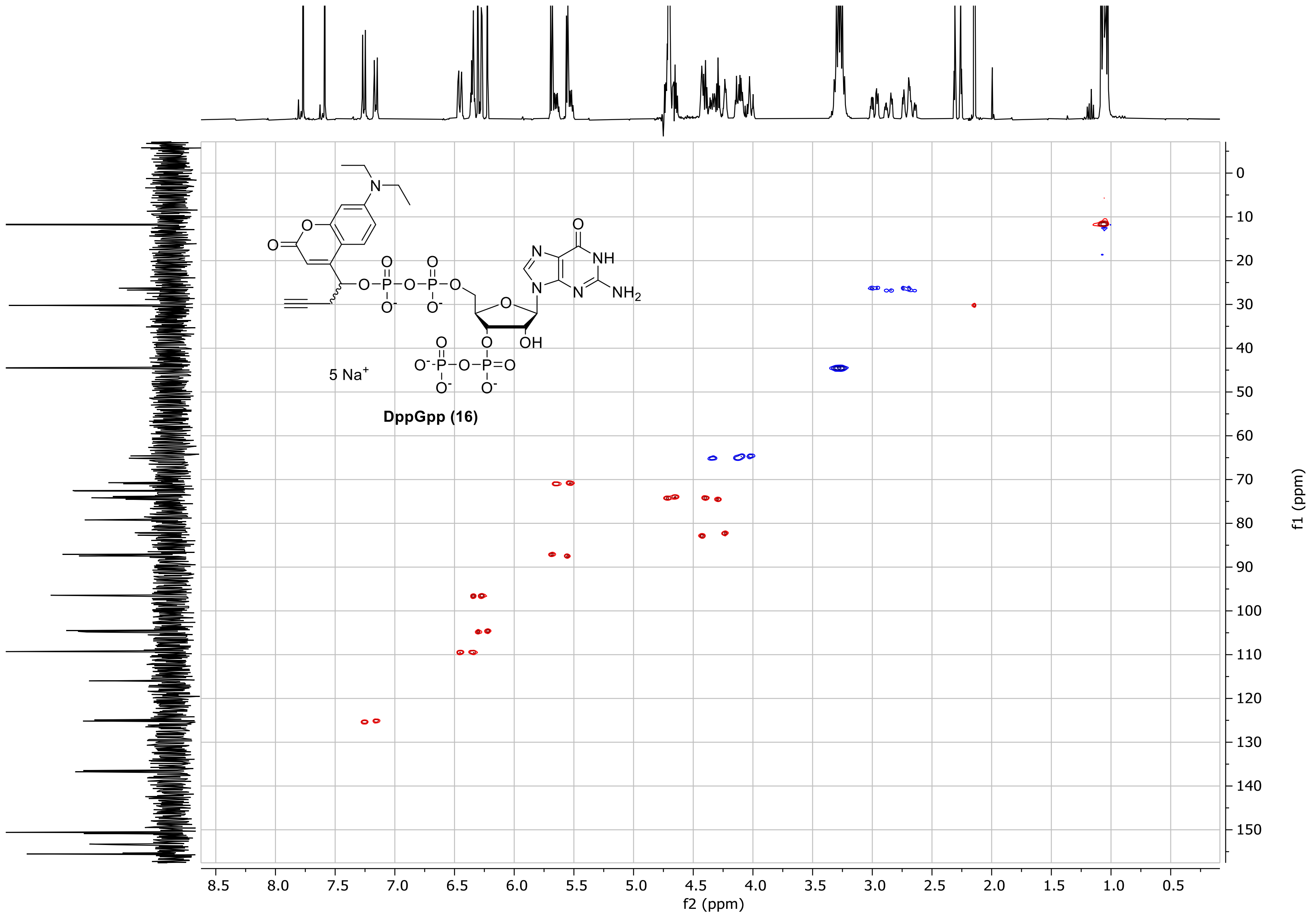

### HSQC of 16

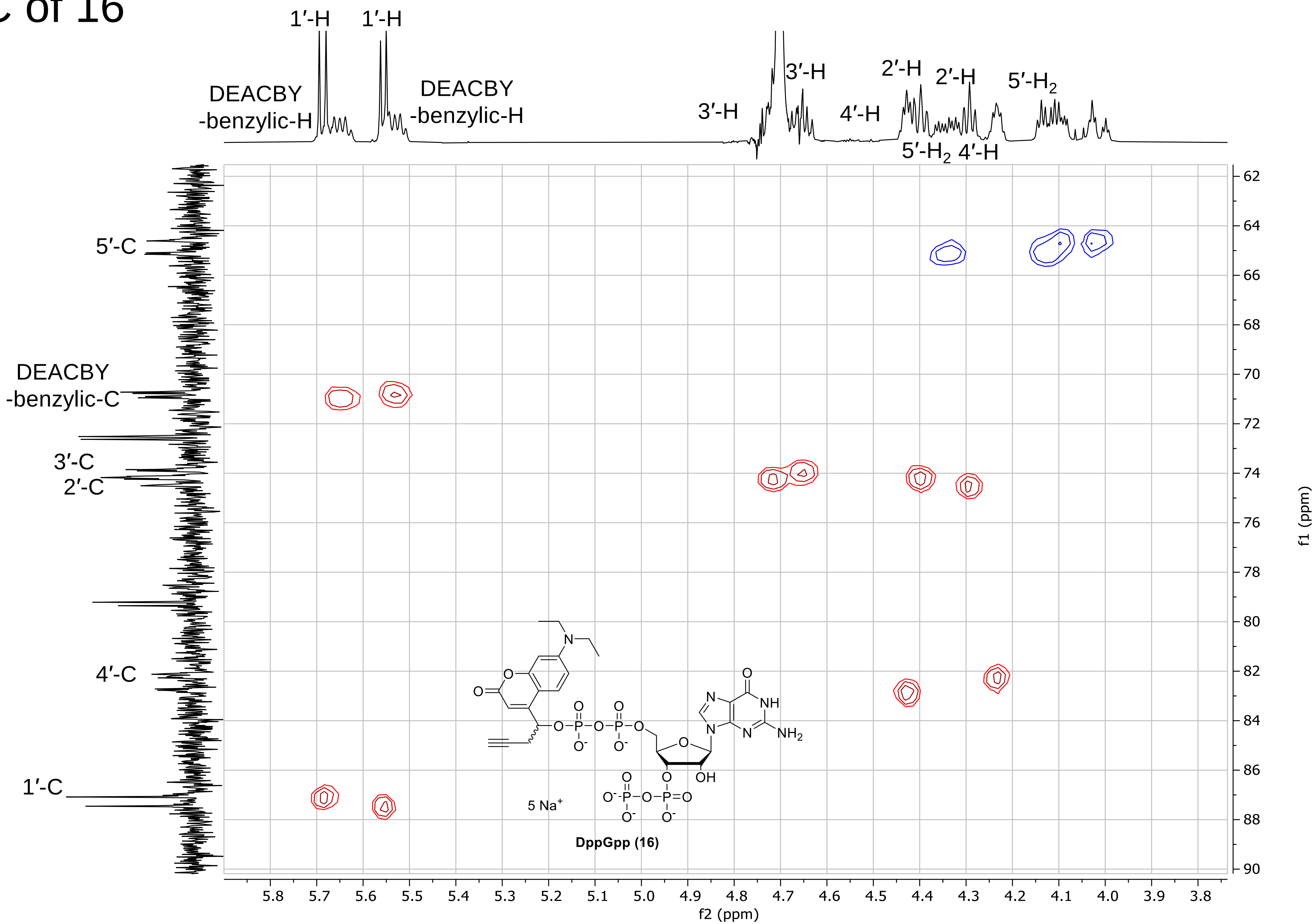
1′-H
1′-H
DEACBY
-benzylic-H
DEACBY
-benzylic-H
3′-H
3′-H
4′-H
2′-H
2′-H
5′-H2
5′-H2
4′-H
5′-C
DEACBY
-benzylic-C
3′-C
2′-C
4′-C
1′-C
5 Na+
DppGpp (16)
f1 (ppm)
f2 (ppm)

### $^{31}$P-HMBC of 16

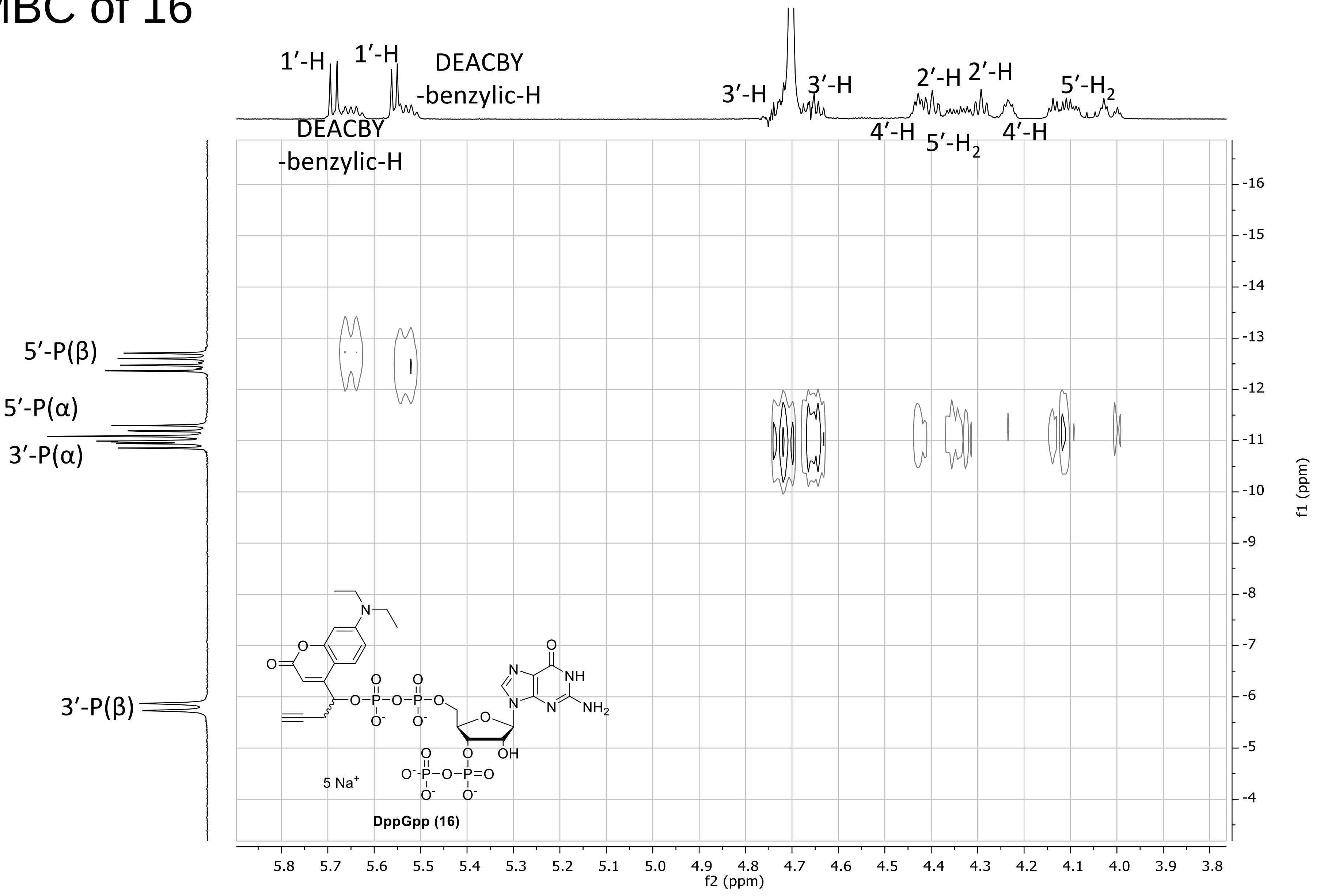

### HMBC of 16

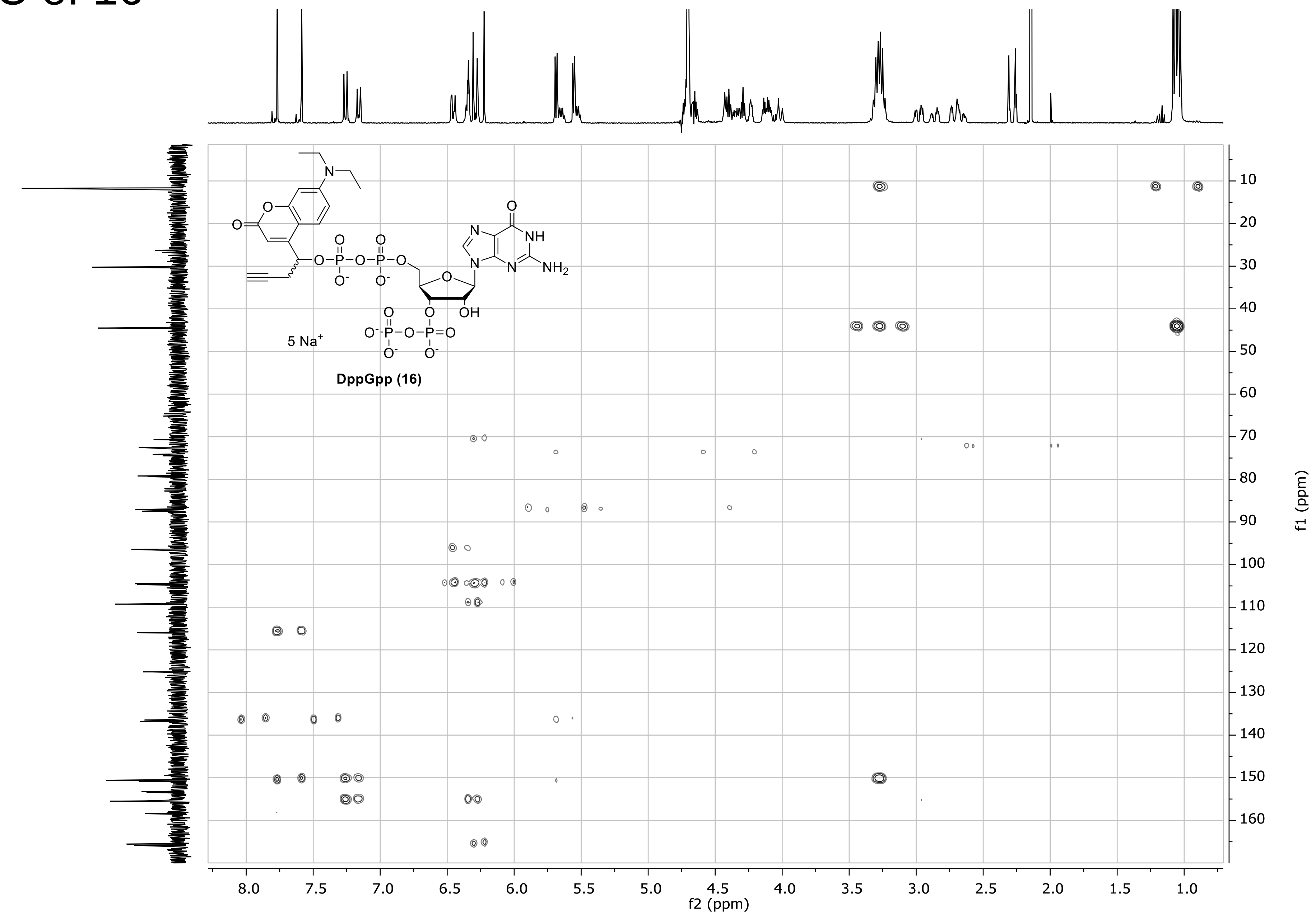

### $^{1}$H-NMR spectrum of S1

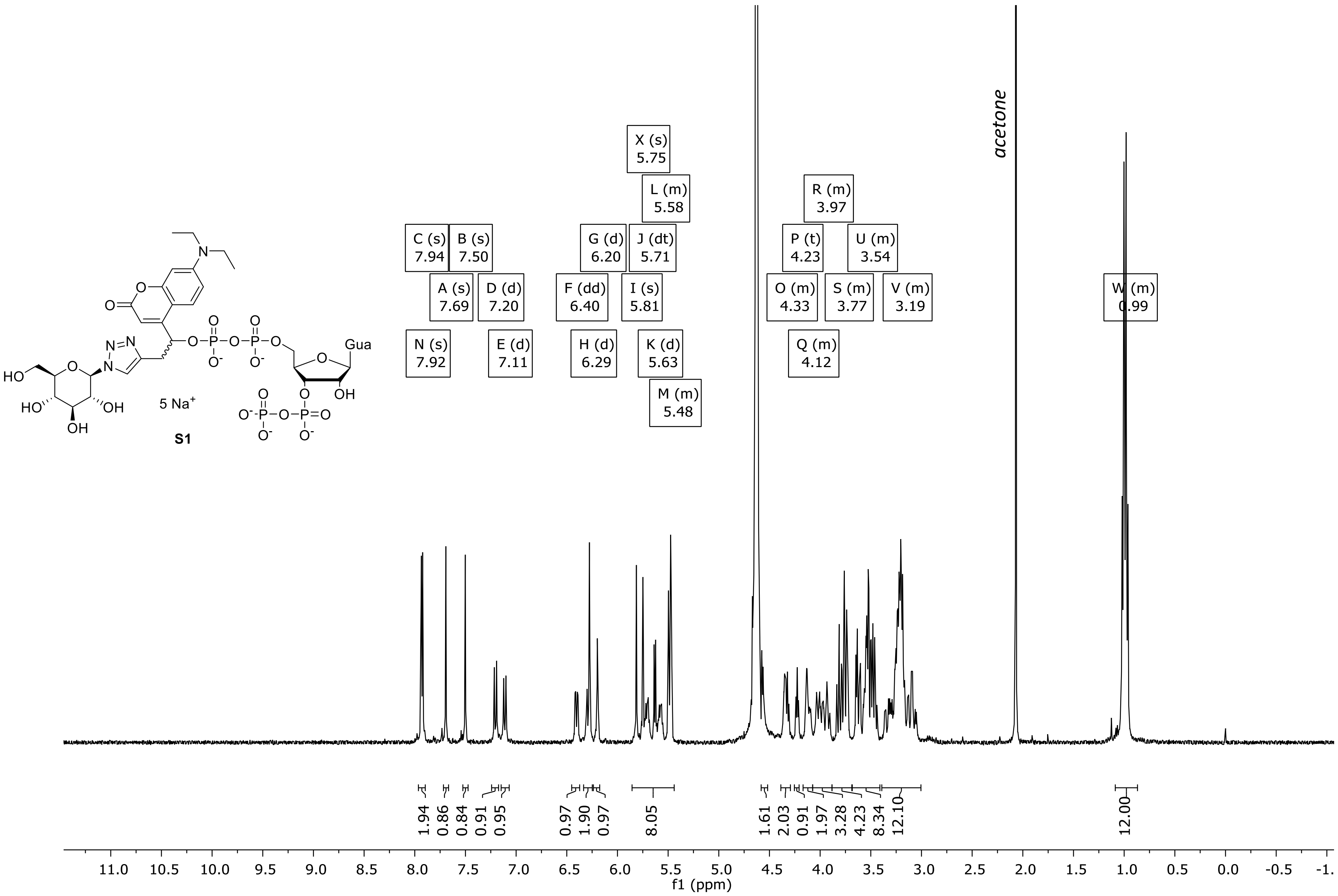

### $^{31}$P-NMR spectrum of S1

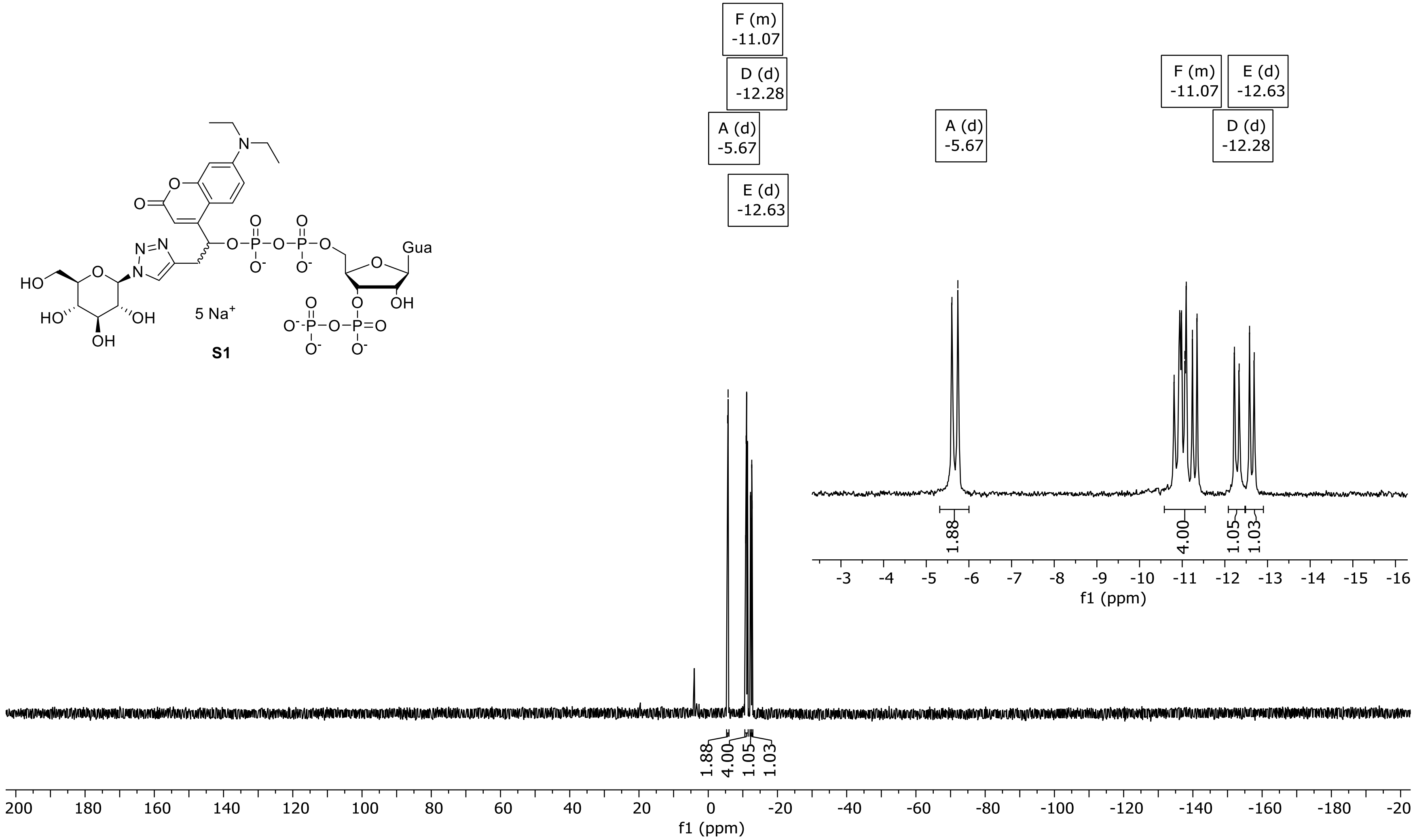

### DQF-COSY of S1

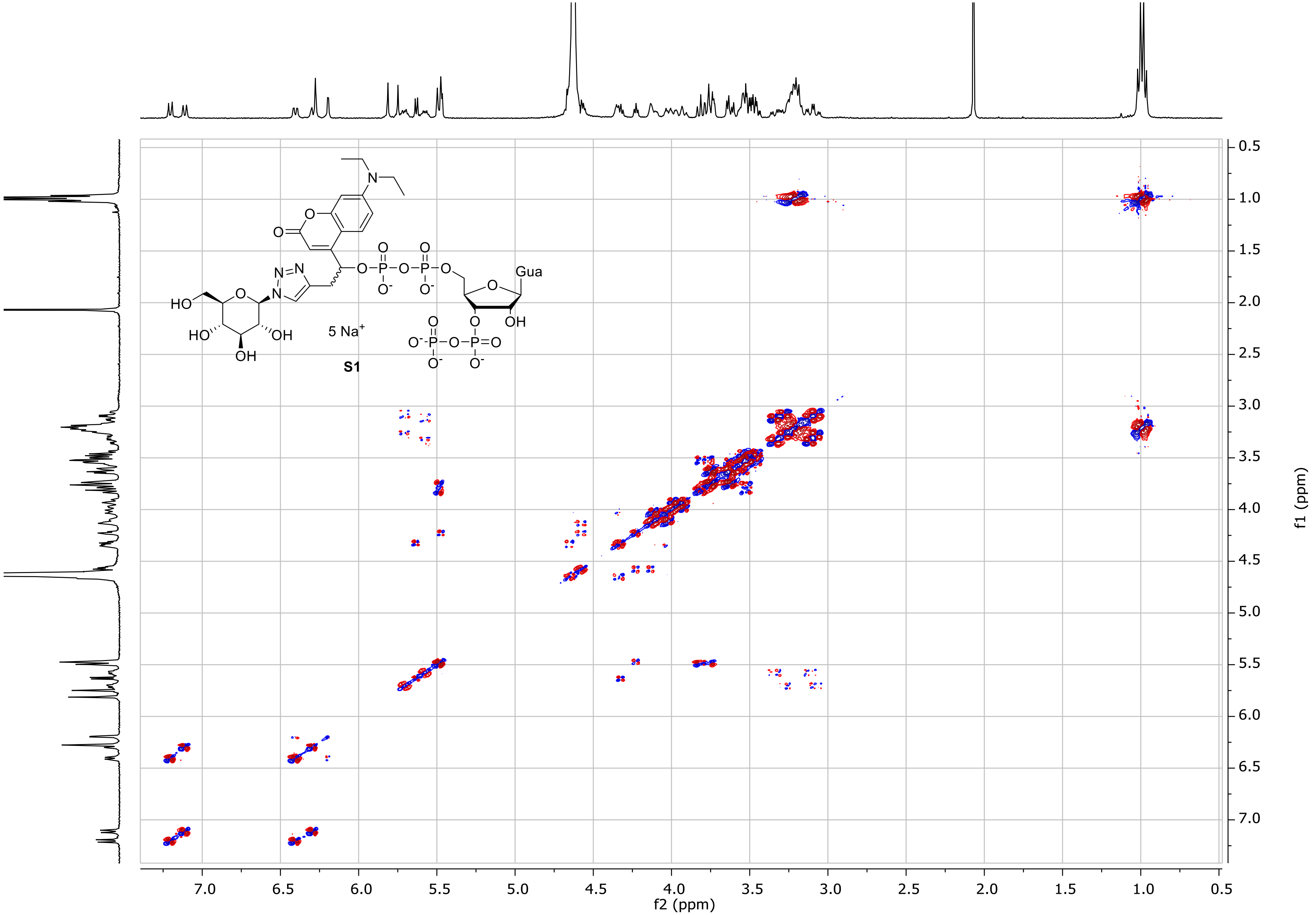

### $^{1}$H-NMR spectrum of S2

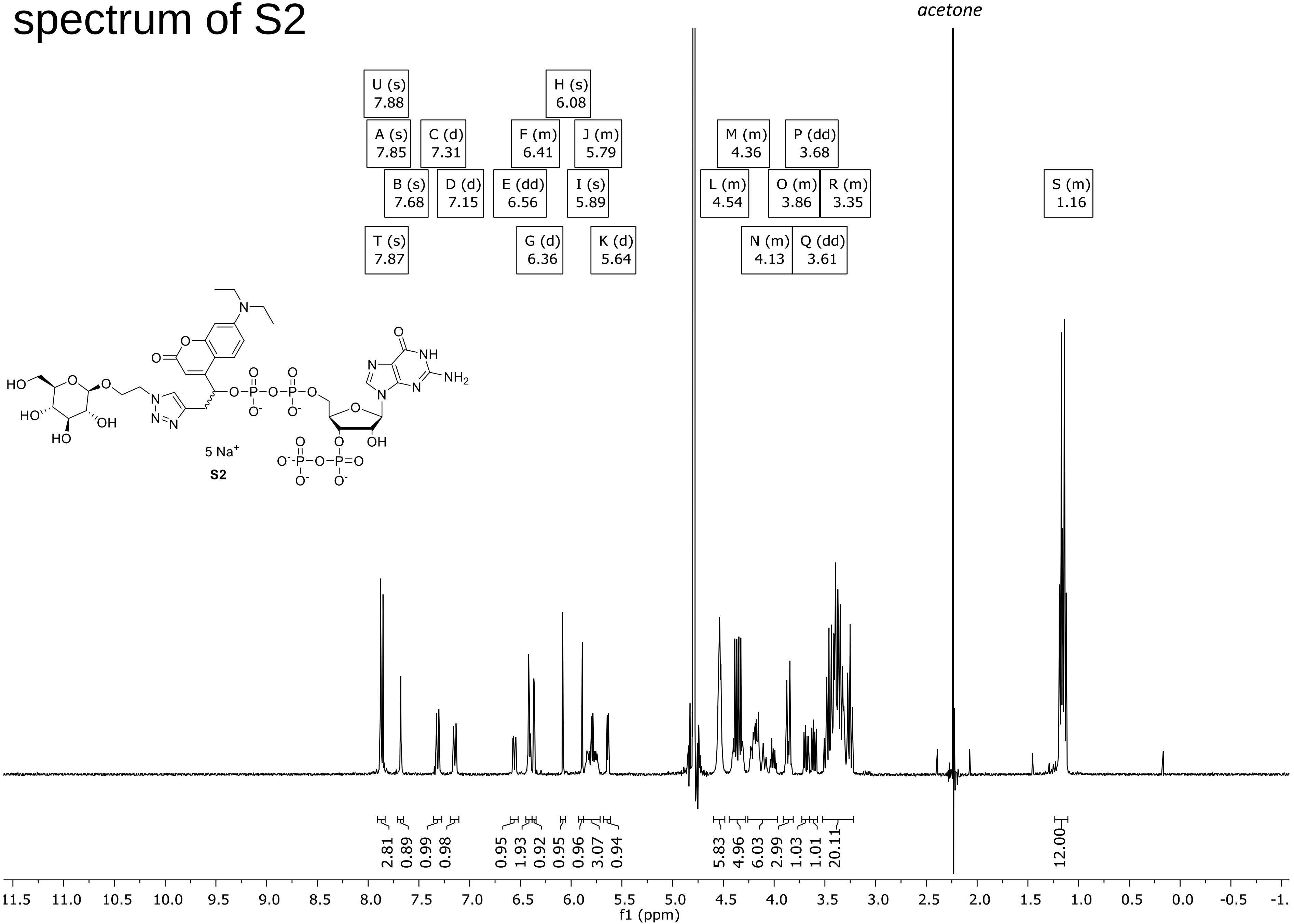

### $^{31}$P-NMR spectrum of S2

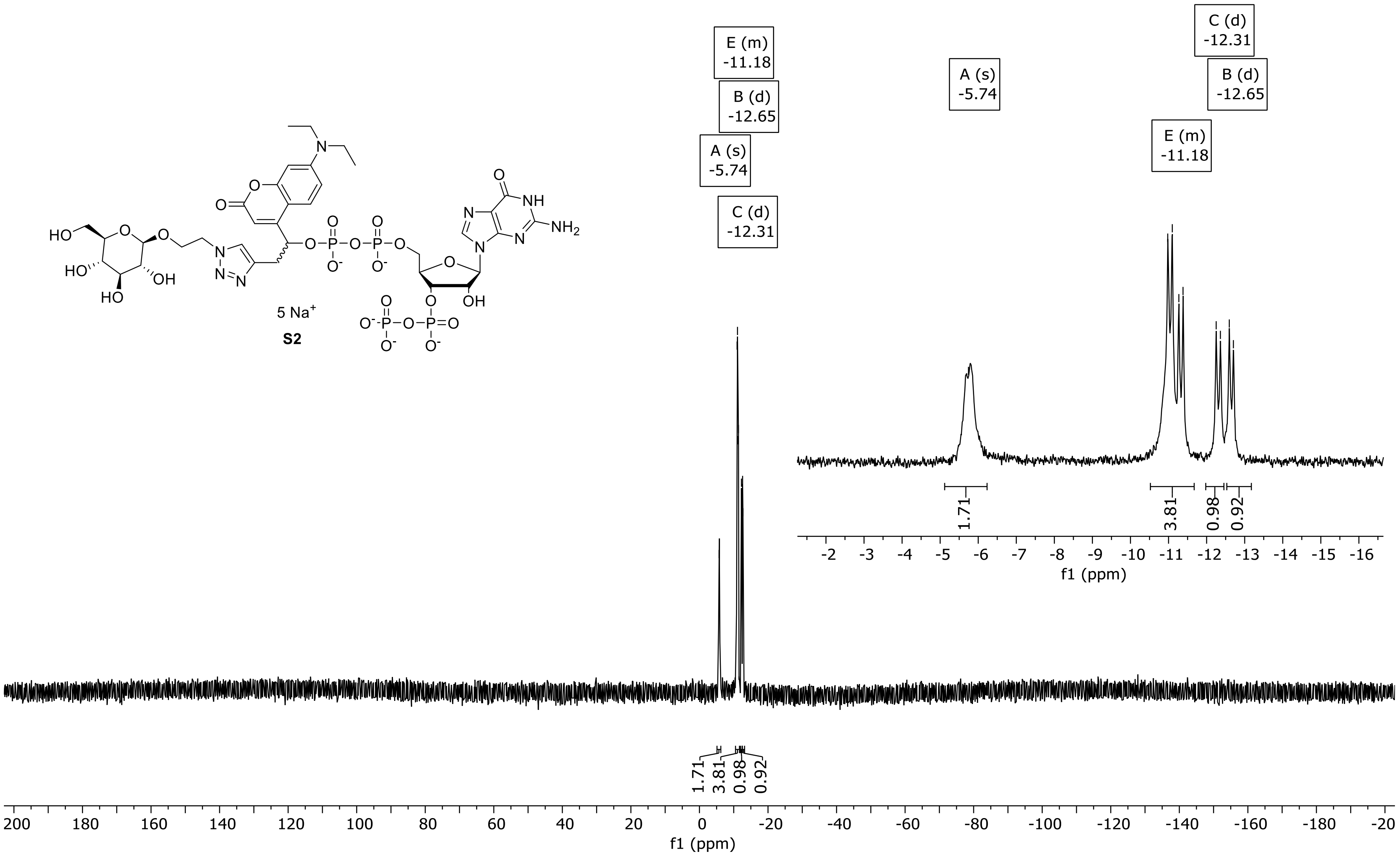

### [1]H-NMR spectrum of S3

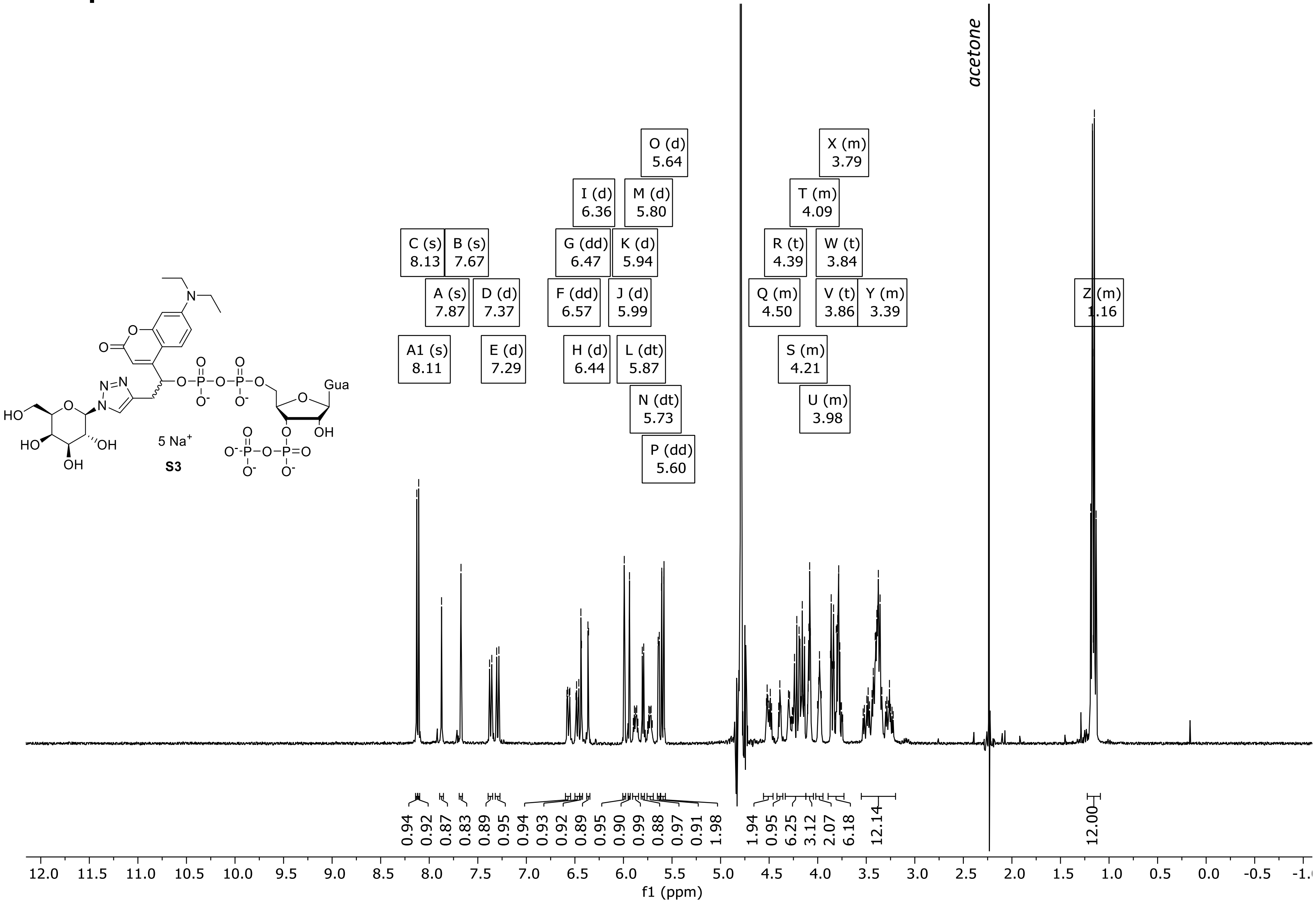

### $^{31}$P-NMR spectrum of S3

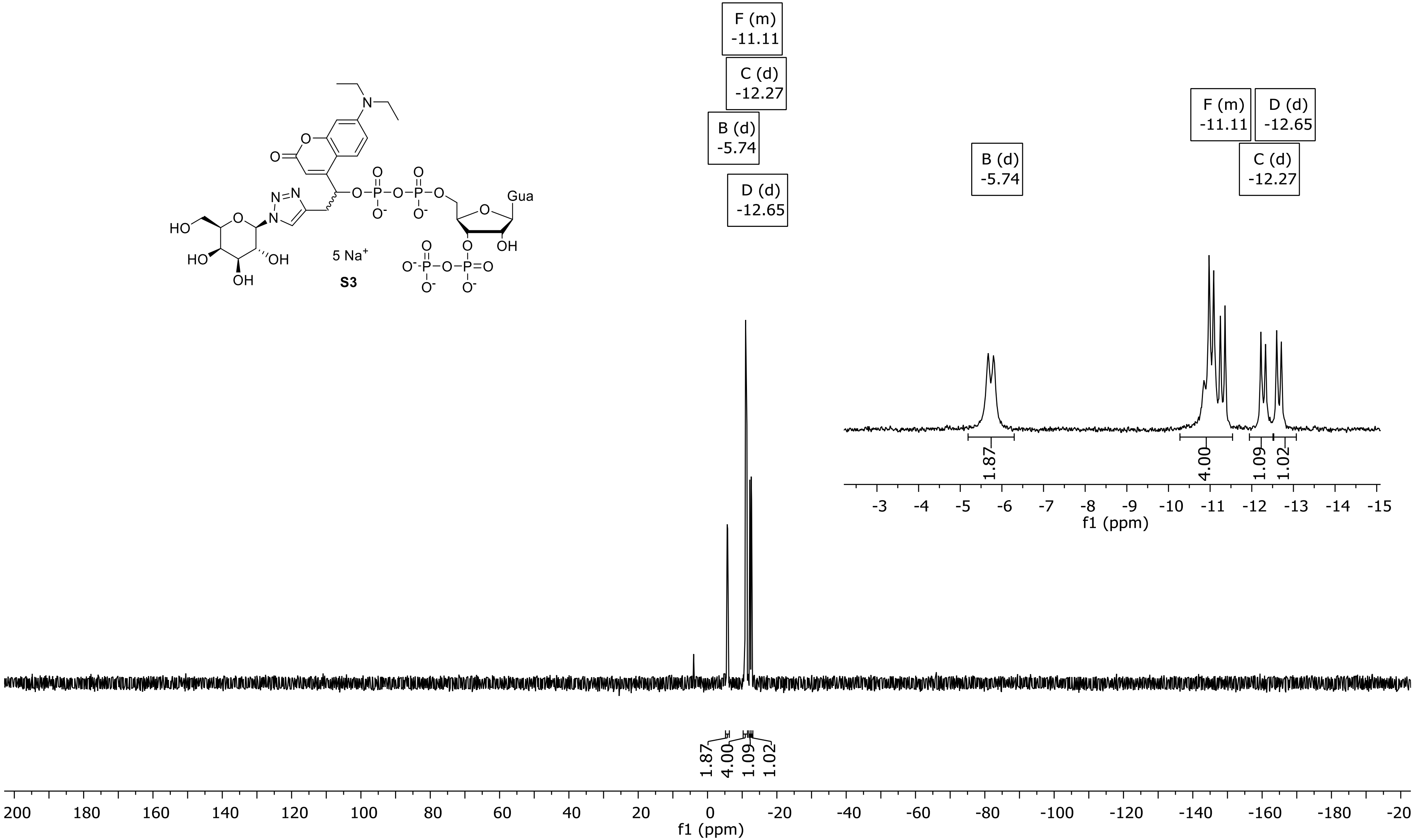

### $^1$H-NMR spectrum of S4

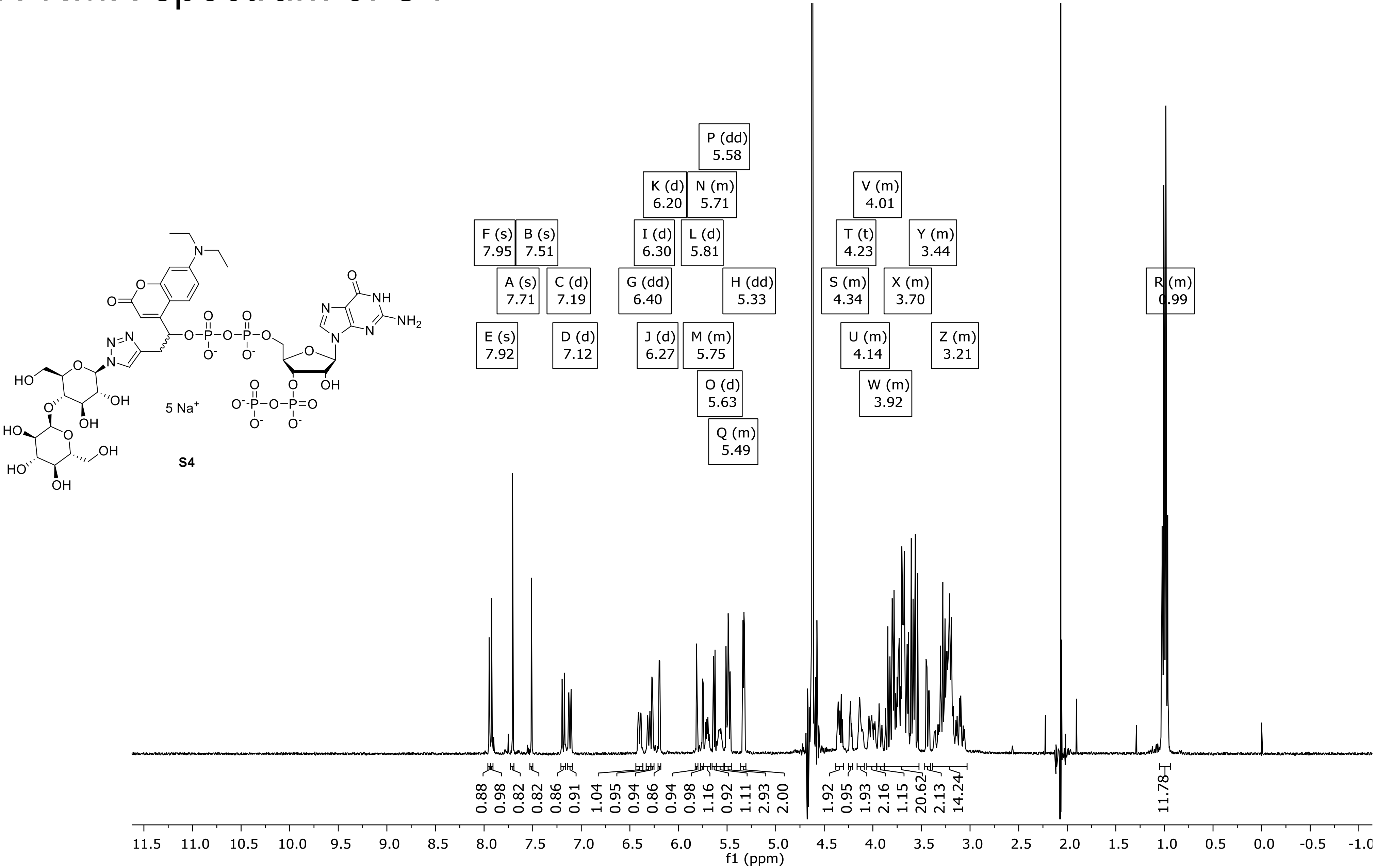

### $^{31}$P-NMR spectrum of S4

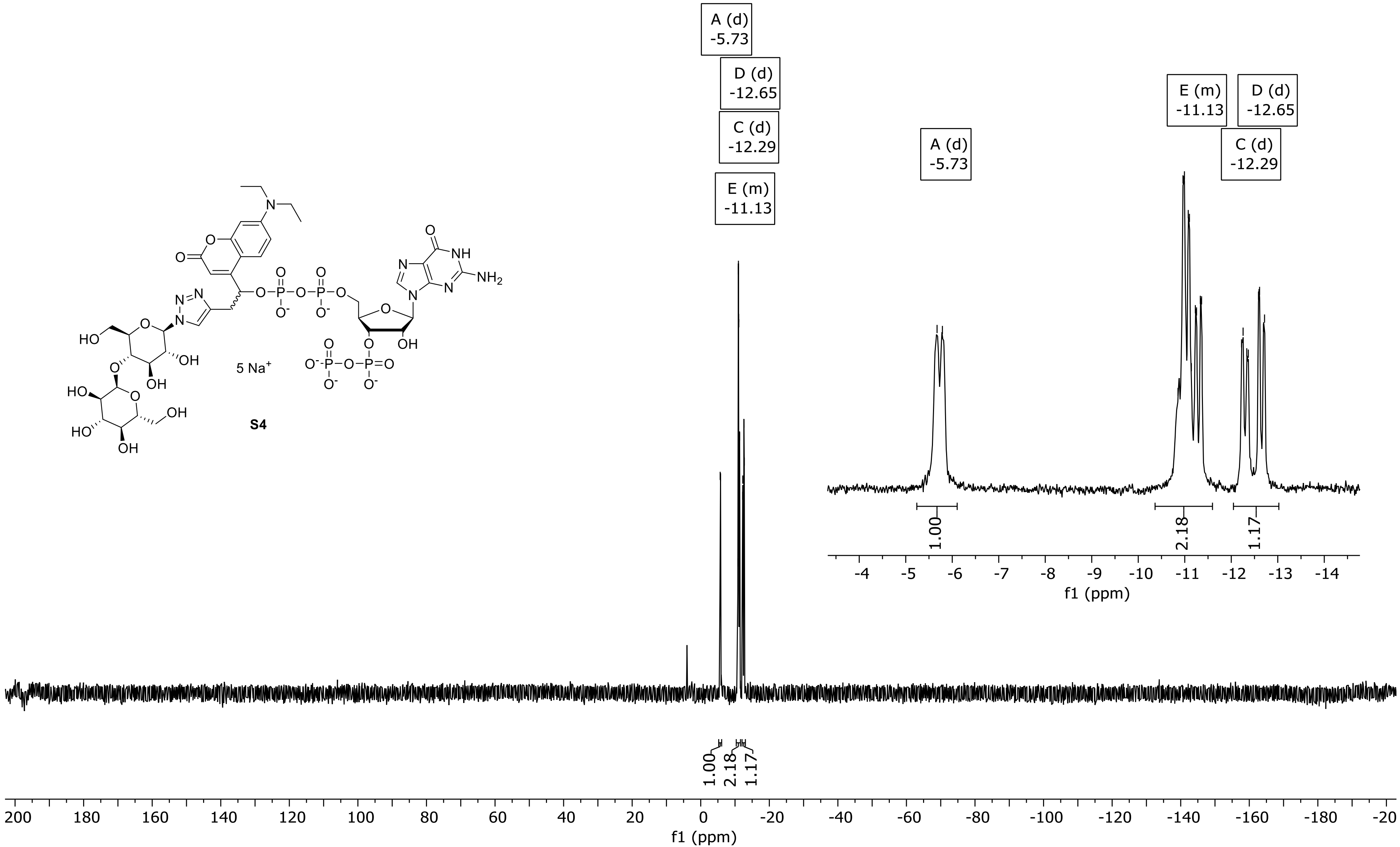

### DQF-COSY in $D_2O$ of S4

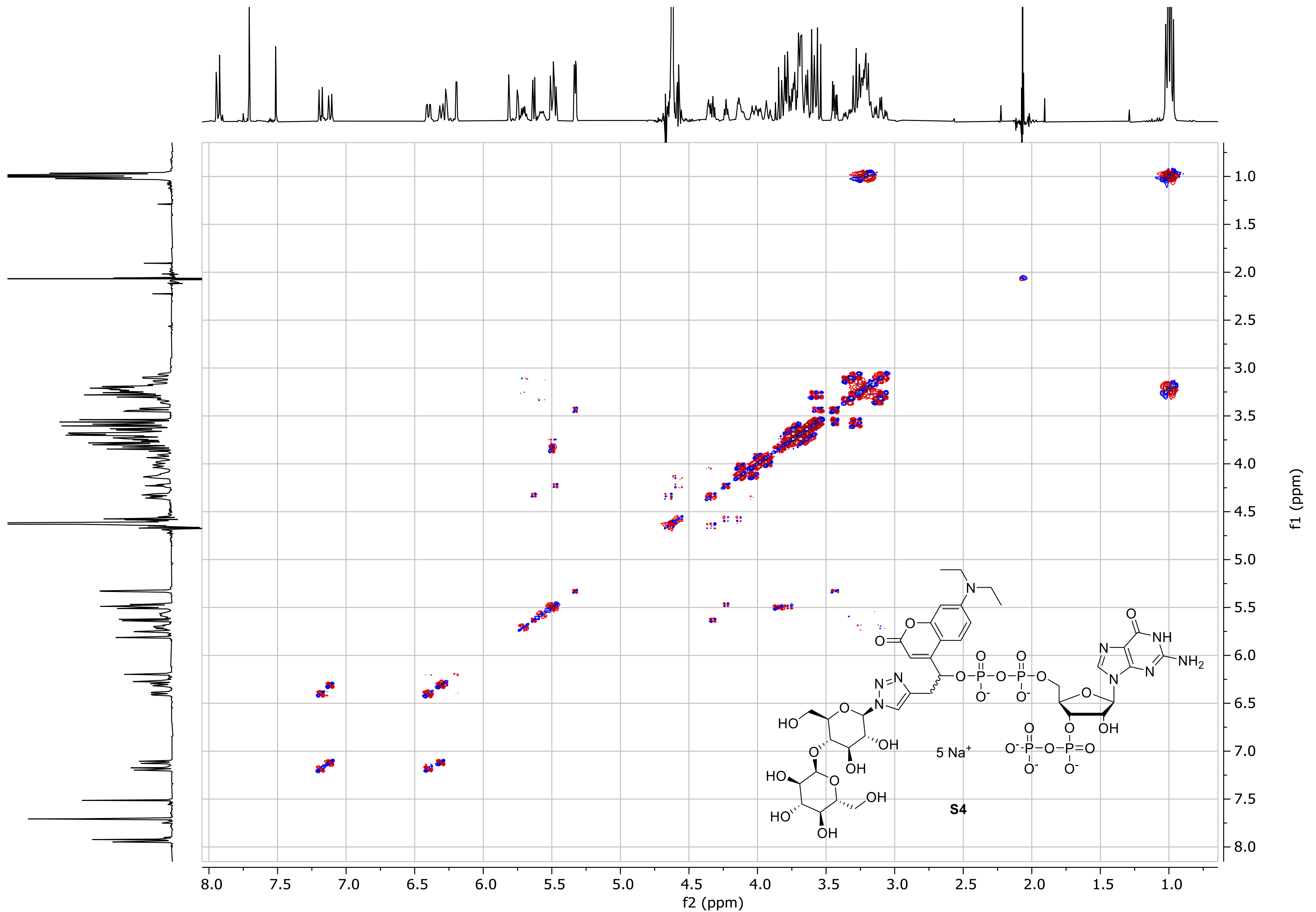

### 1H-NMR spectrum of S5

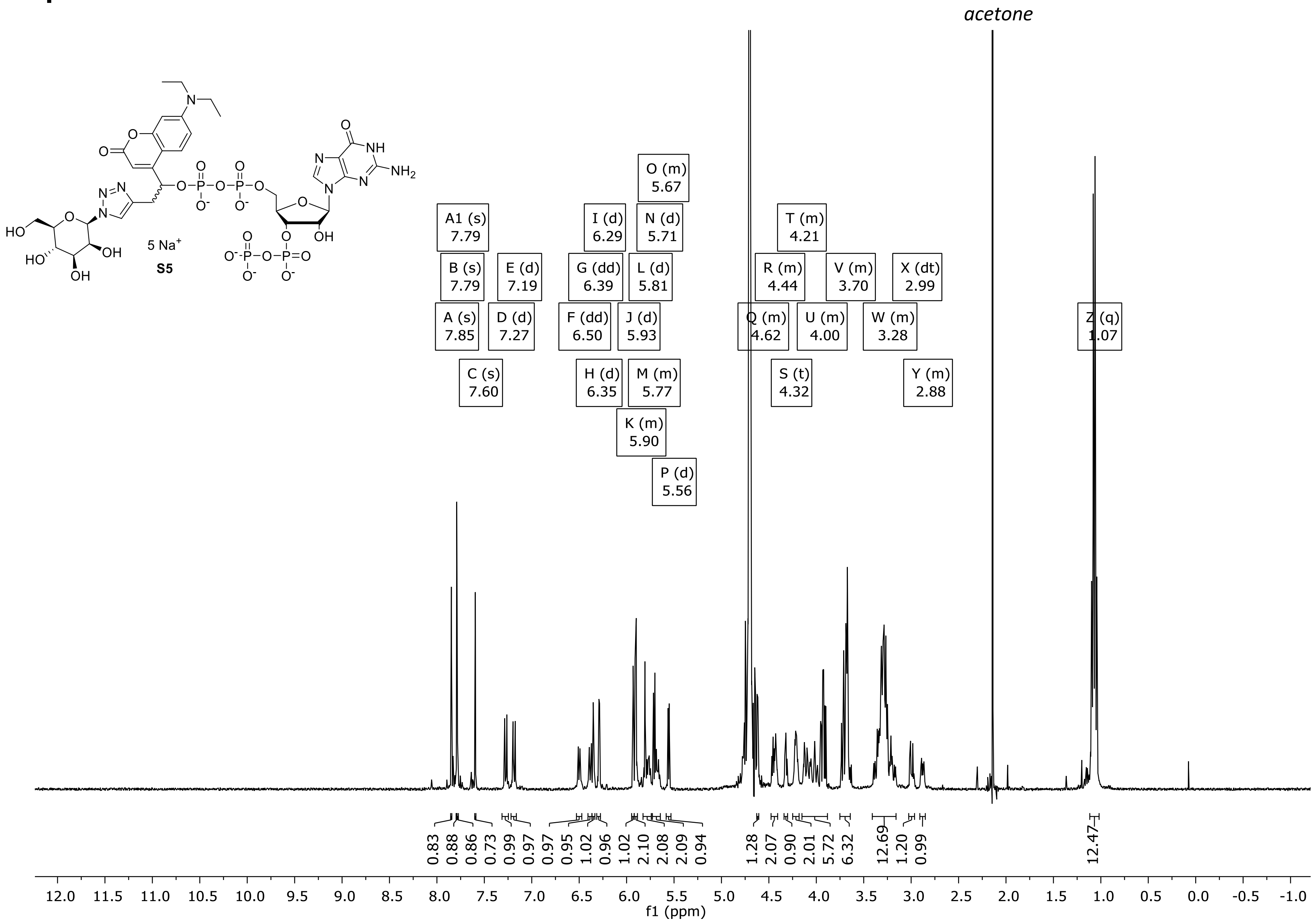

### $^{31}$P-NMR spectrum of S5

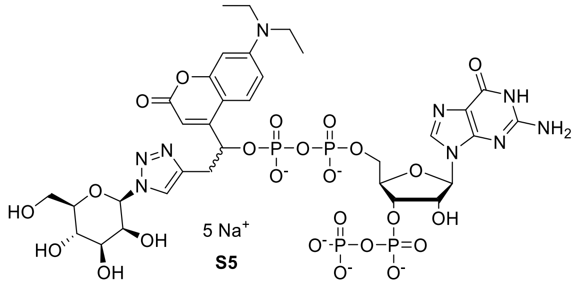

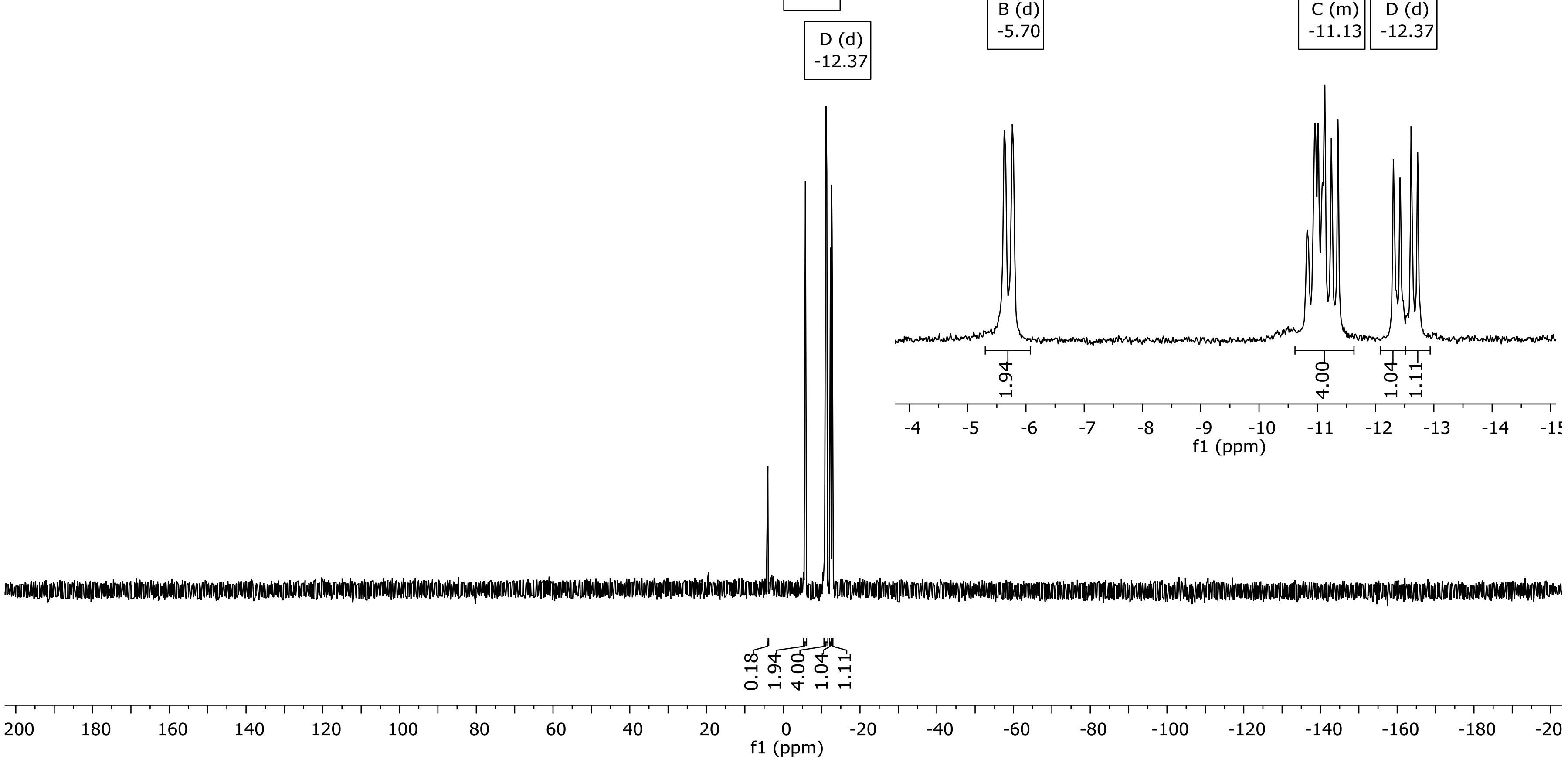

### $^{1}$H-NMR spectrum of S6

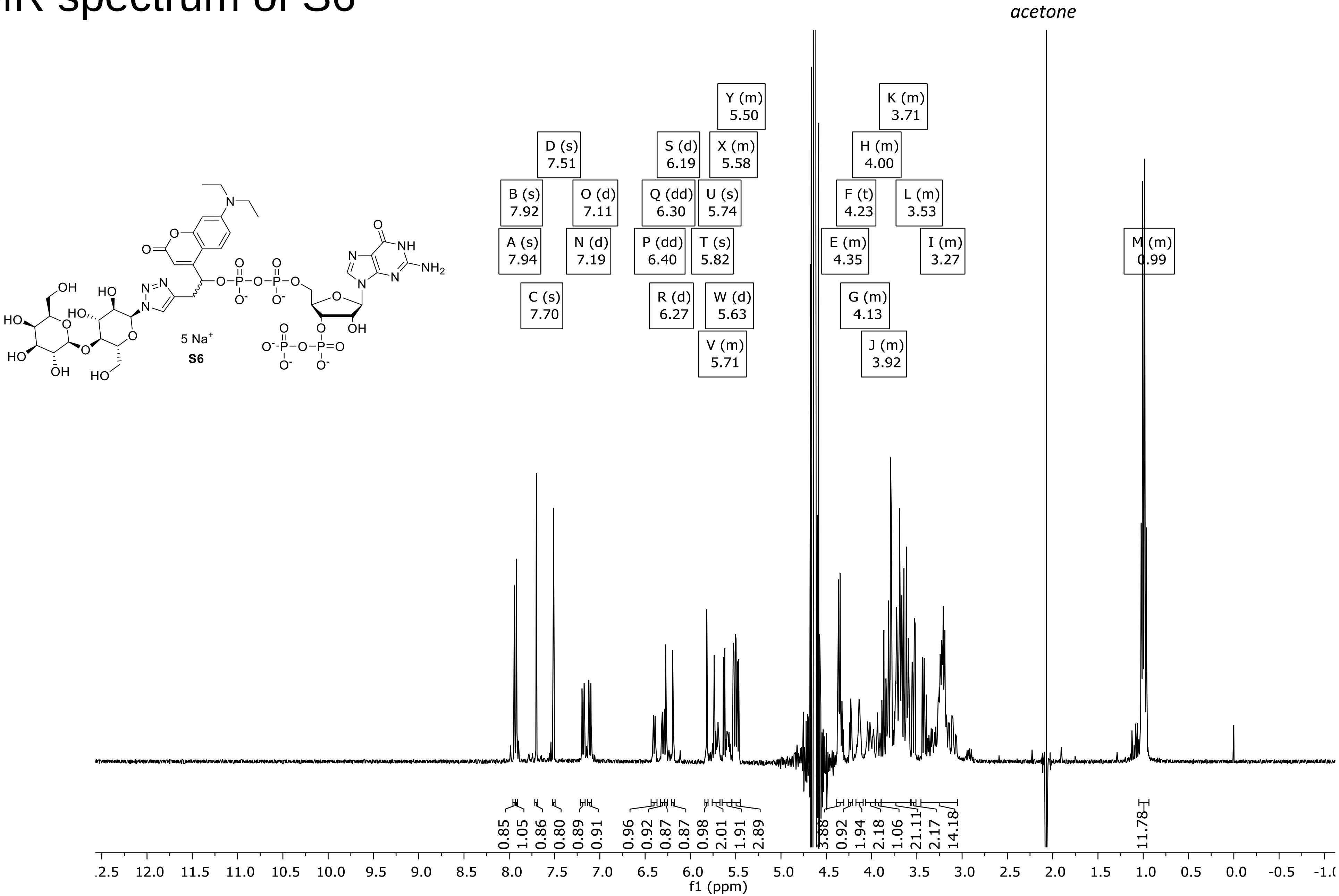

### $^{31}$P-NMR spectrum of S6

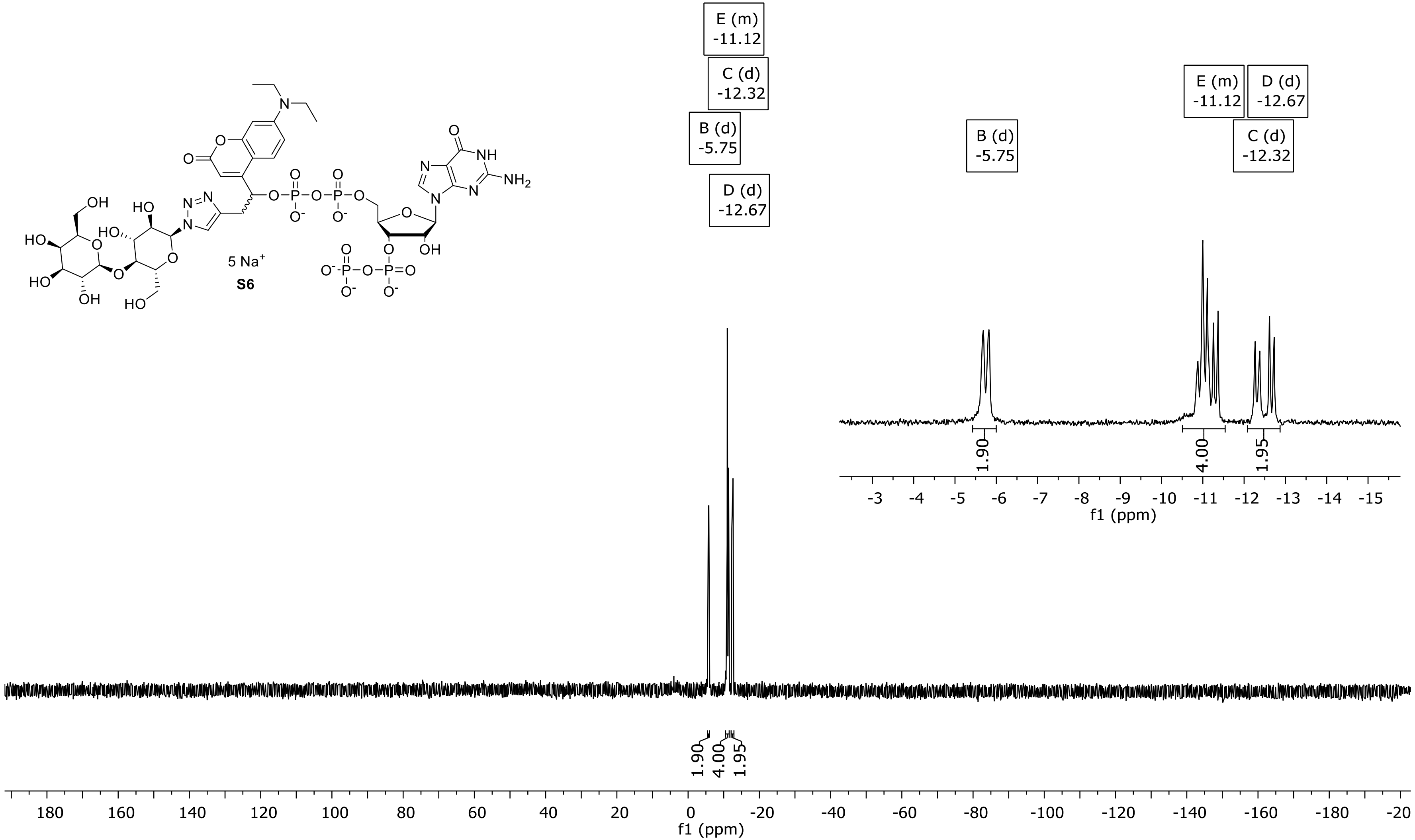

### DQF-COSY in $D_2O$ of S6

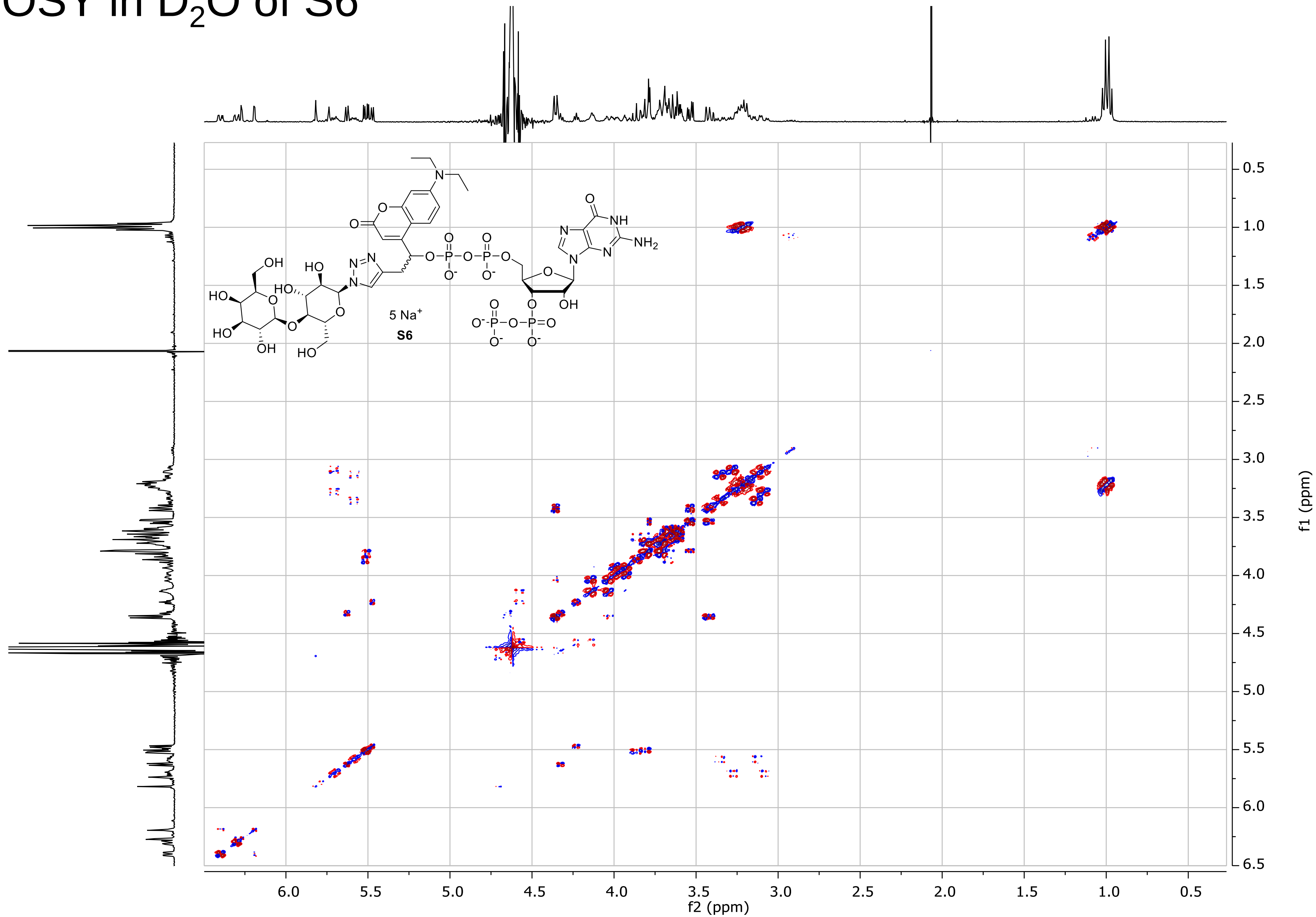

### 1H-NMR spectrum of S7

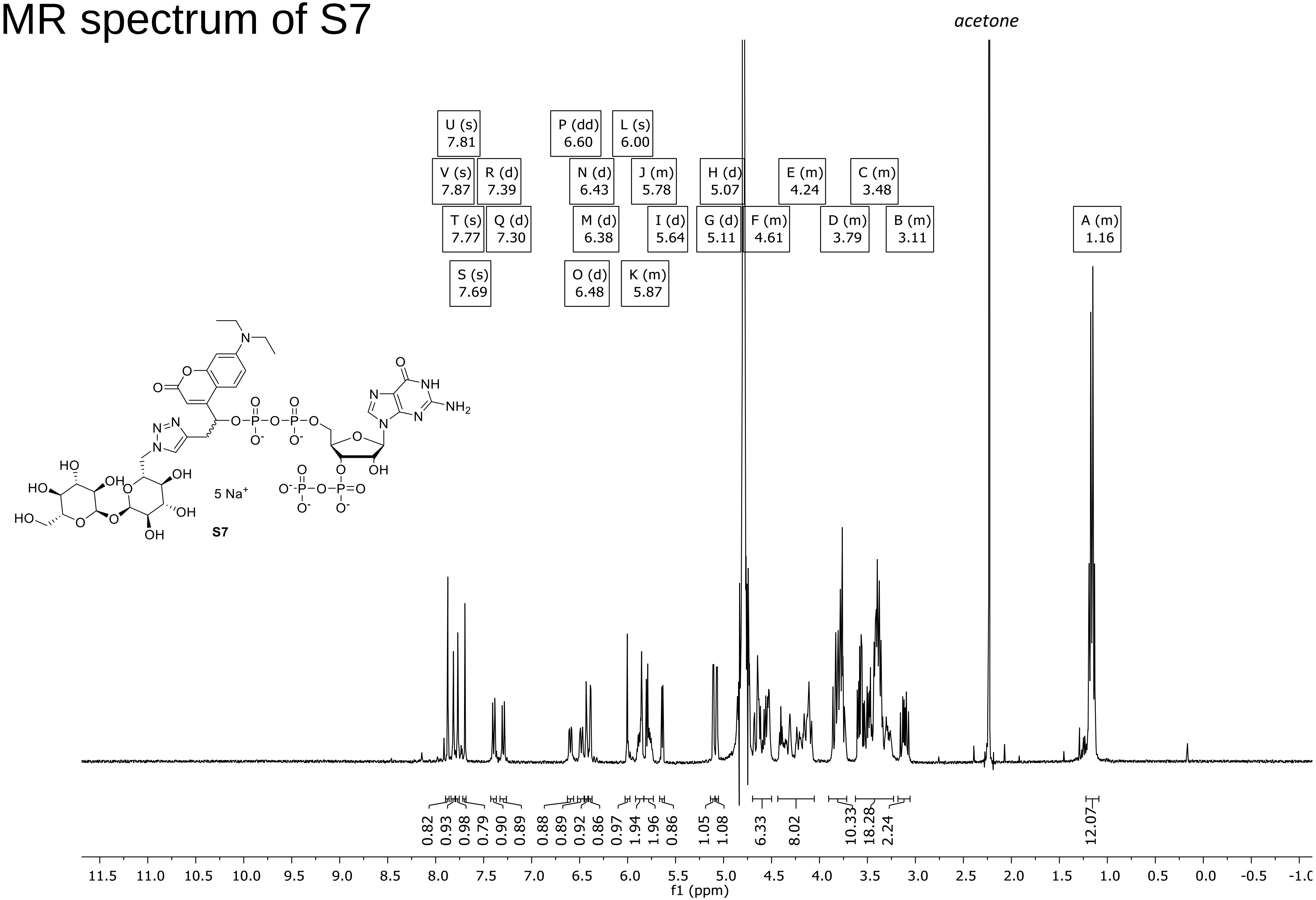

### $^{31}$P-NMR spectrum of S7

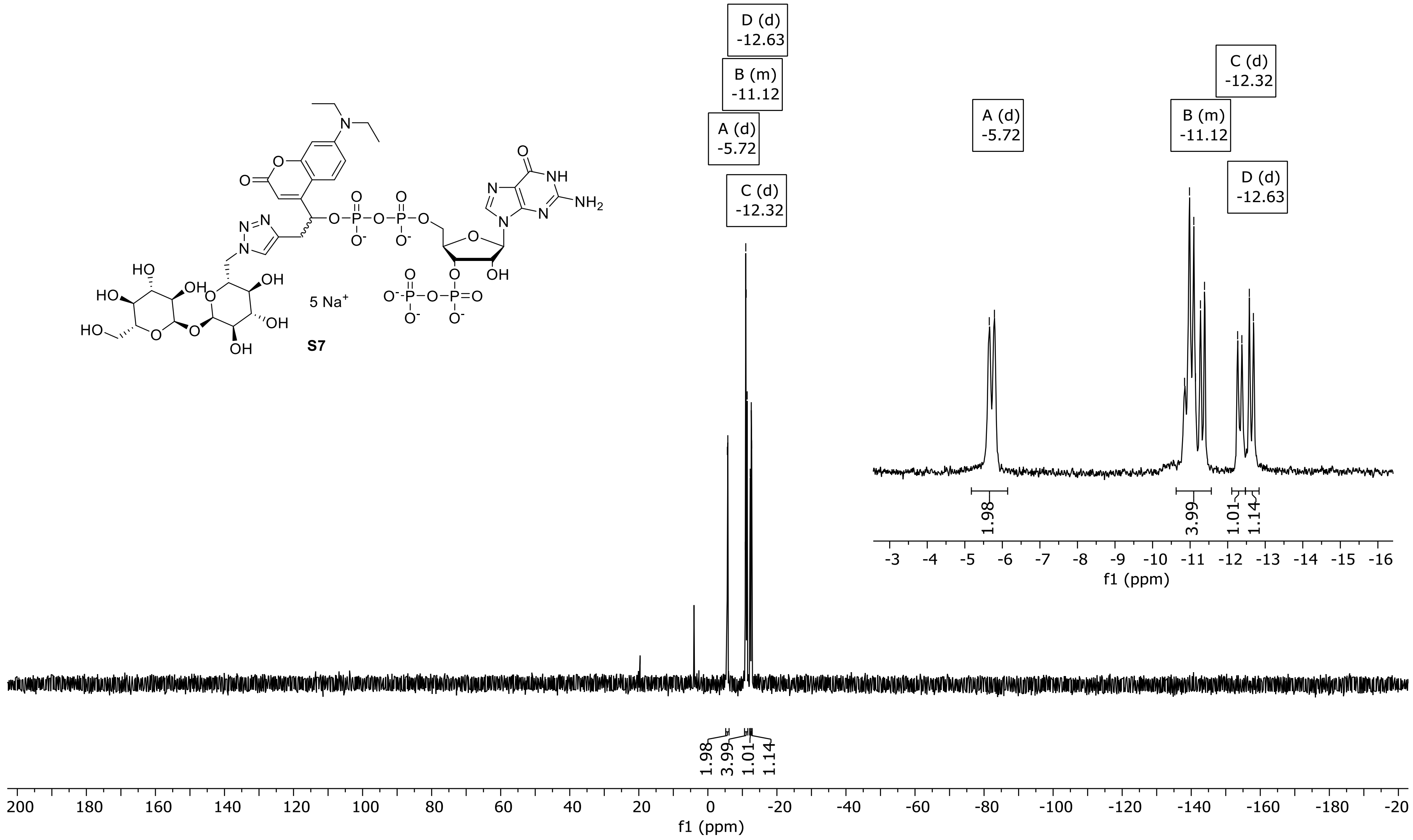

### $^1$H-NMR spectrum of S9

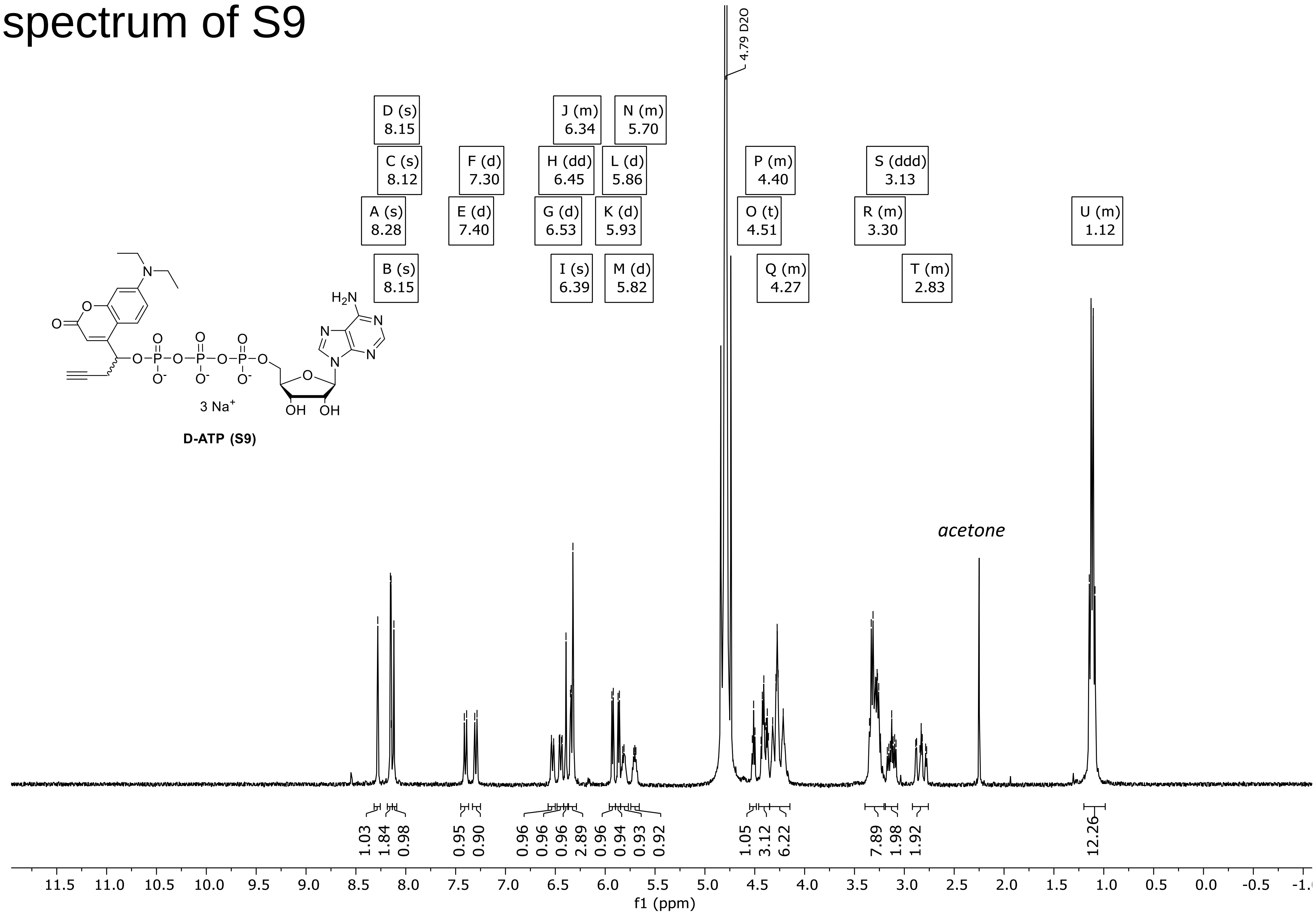

### $^{31}$P-NMR spectrum of S9

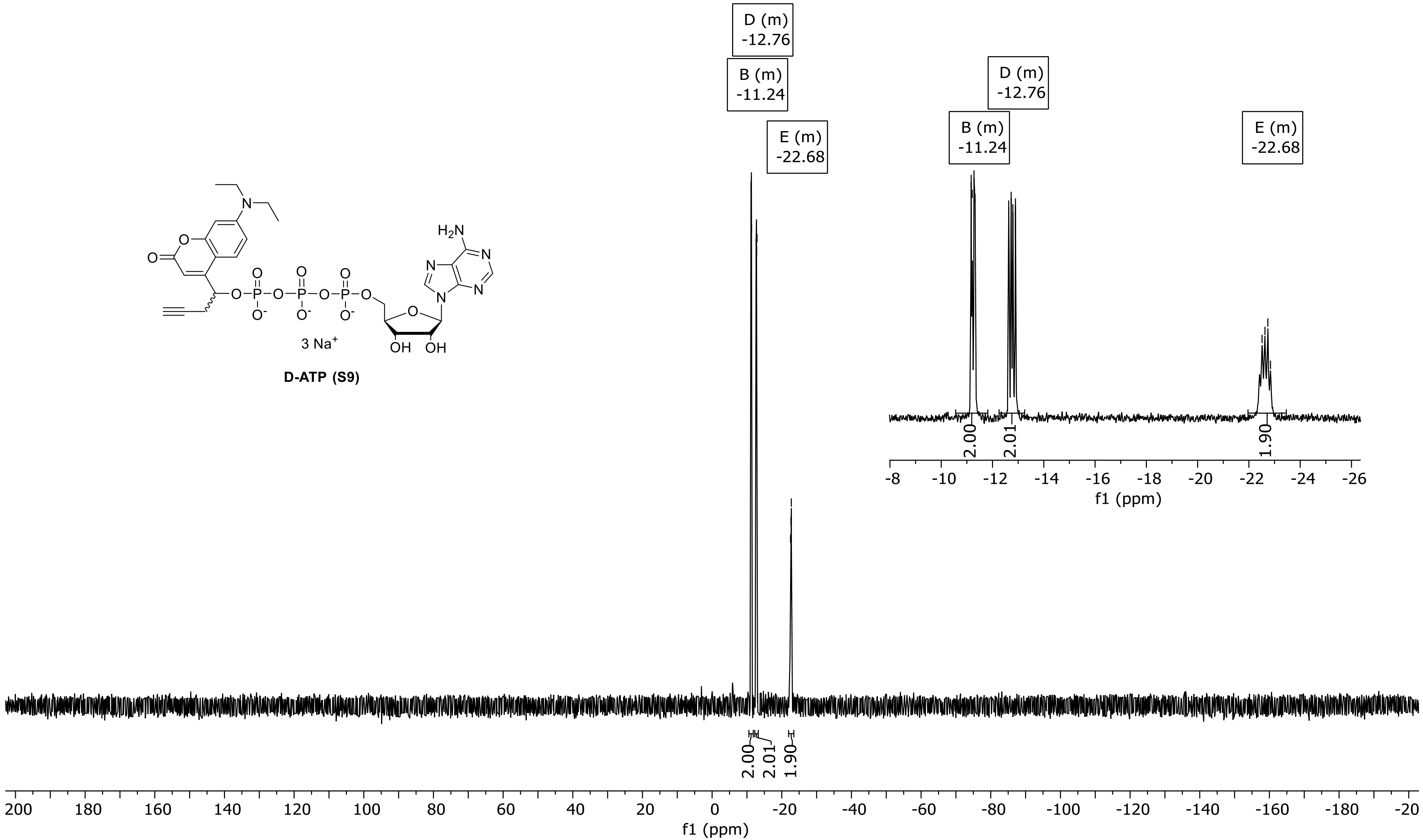

### $^{13}C\ \{^{1}H\}$ -NMR of S9

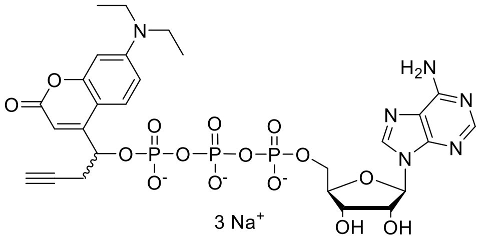

**D-ATP (S9)**

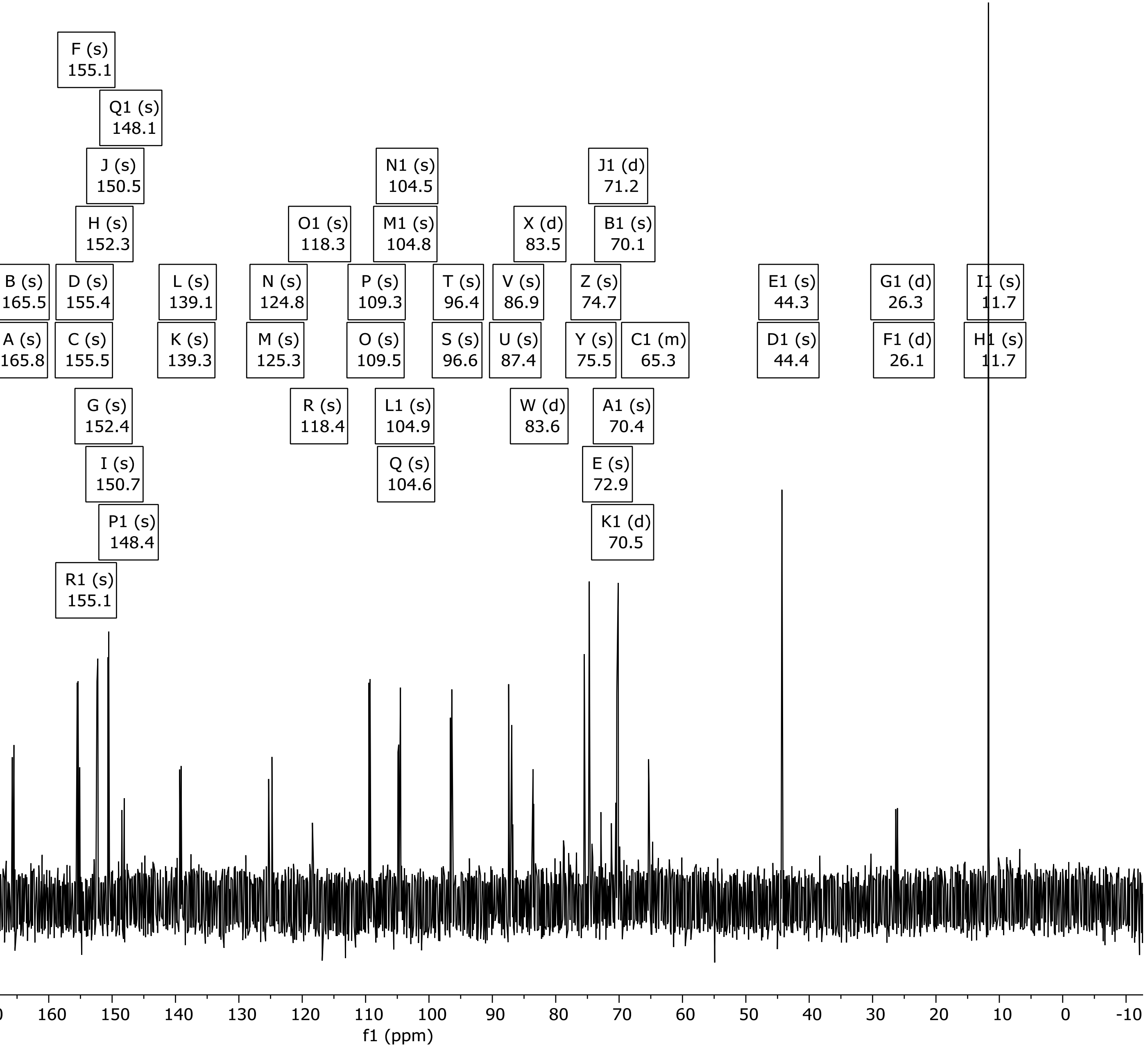

### DQF-COSY in $D_2O$ of S9

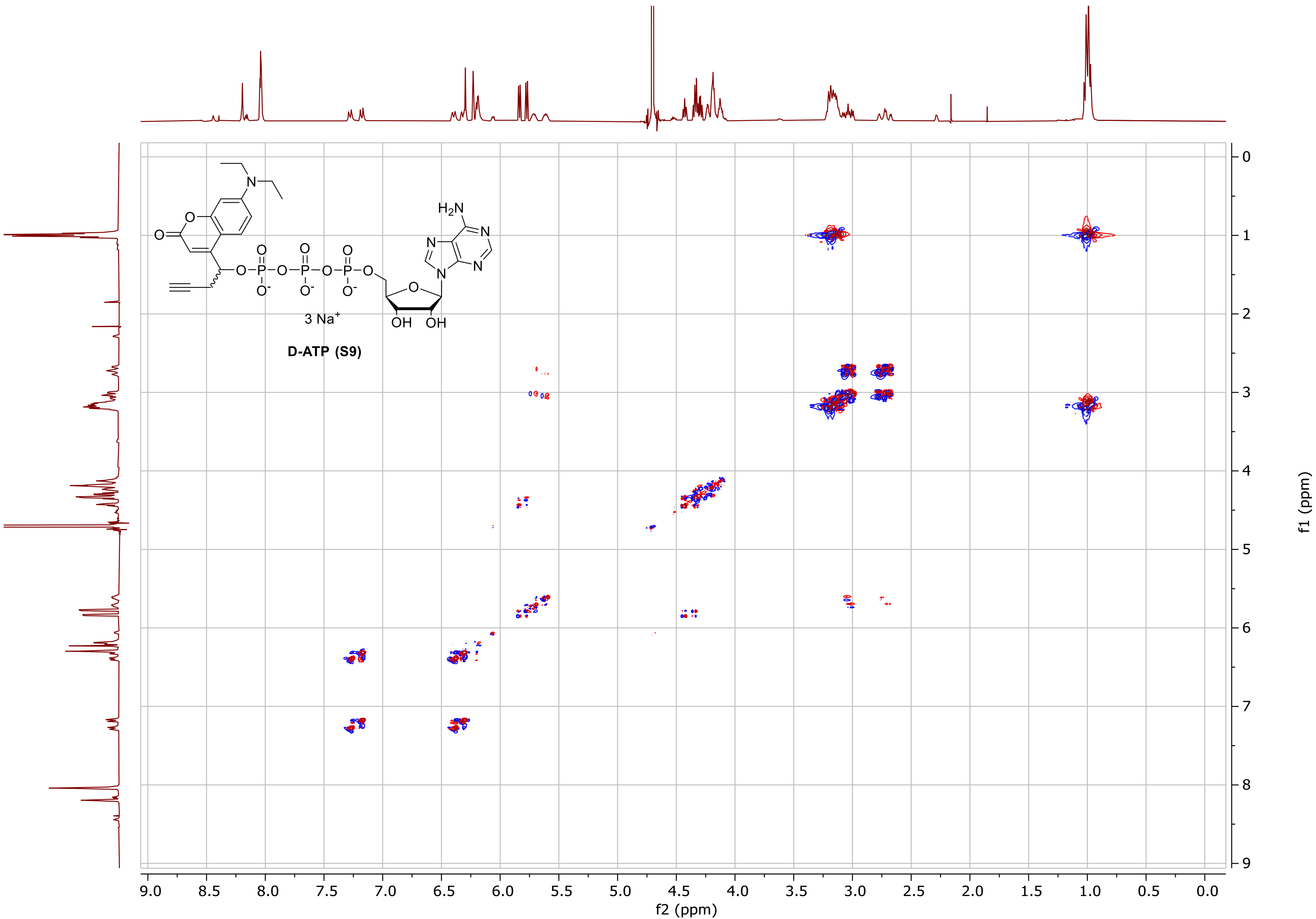

### HSQC in $D_2O$ of S9

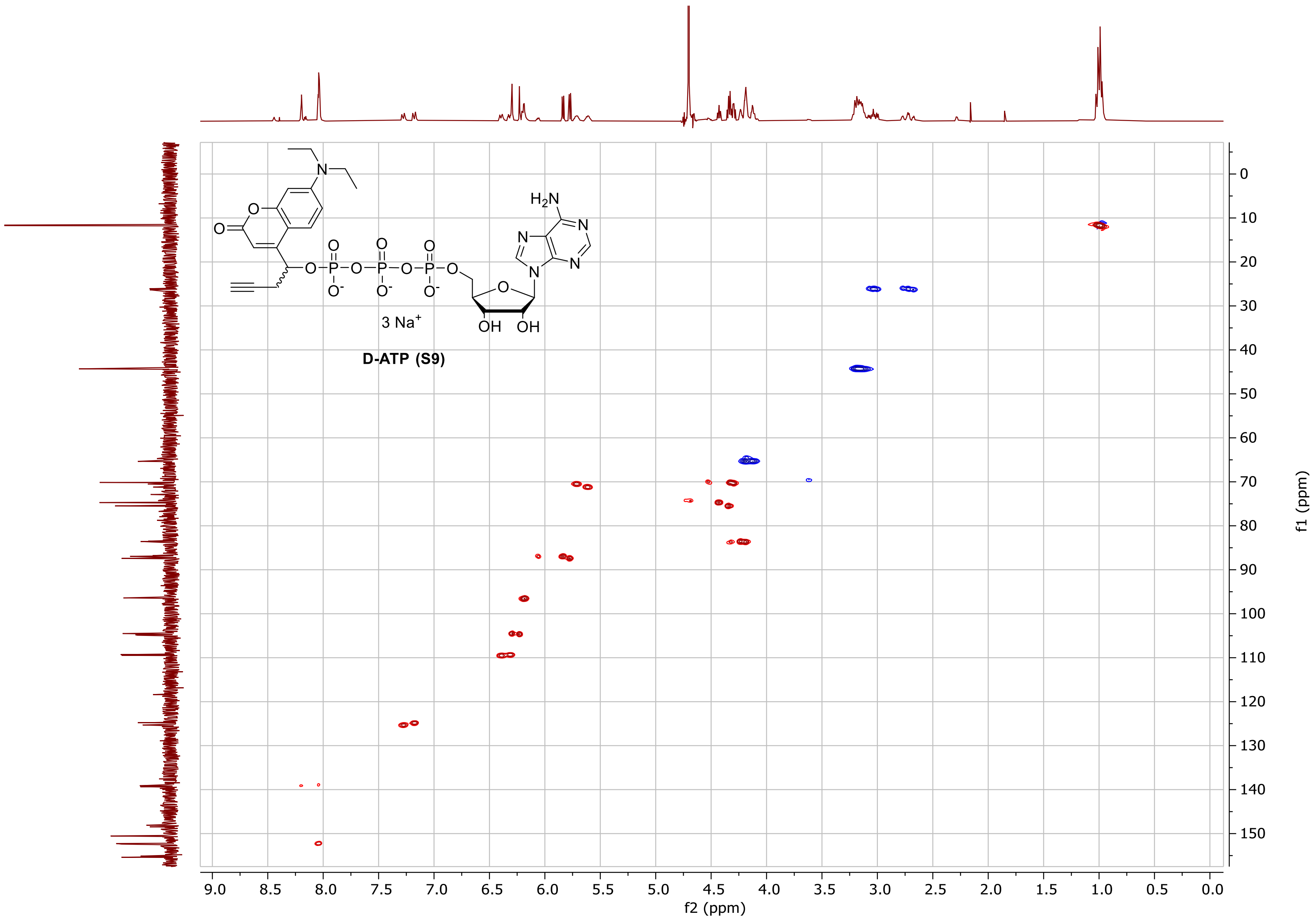

### [1]H-NMR spectrum of S10, peaks of diastereomer 1

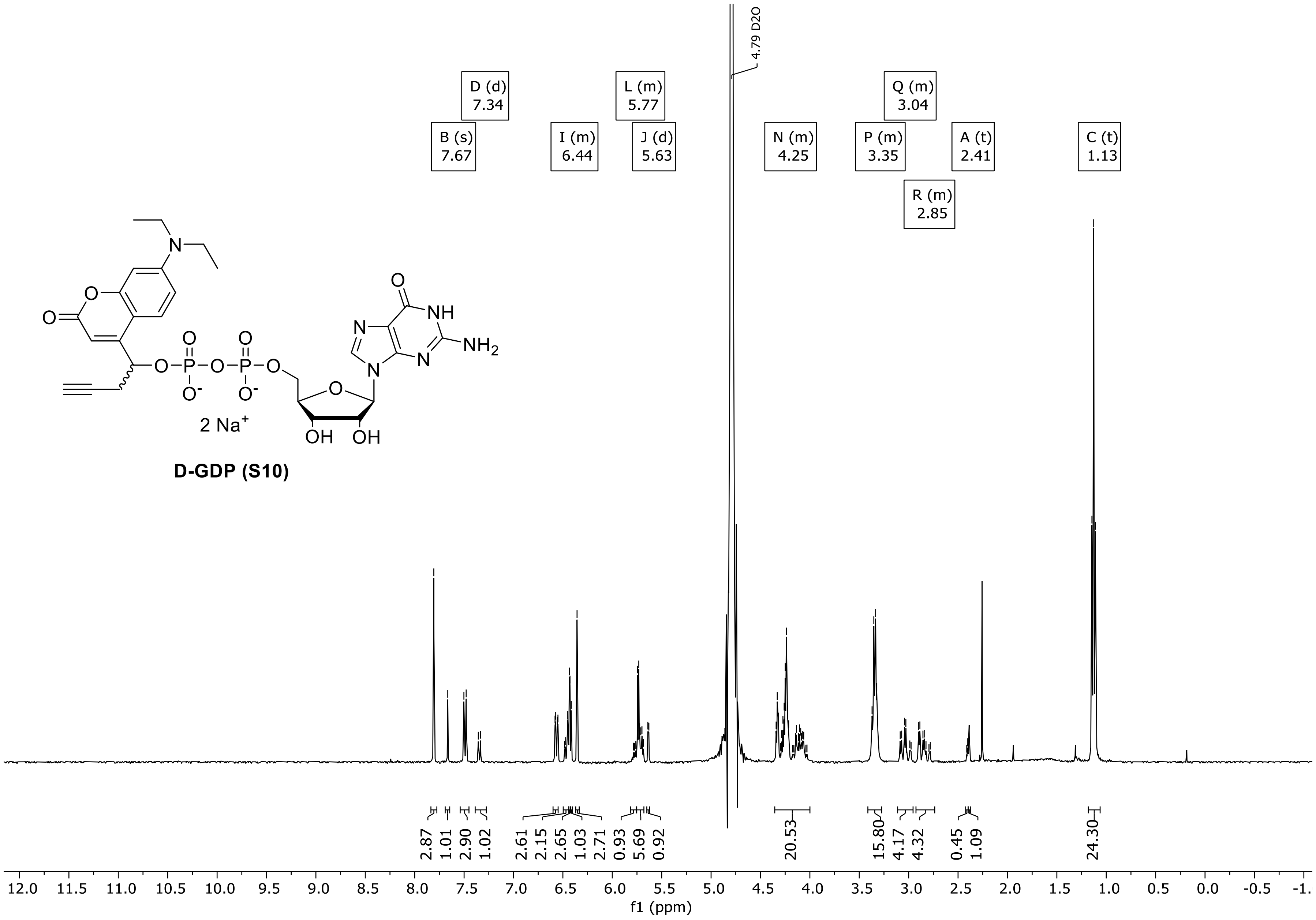

D-GDP (S10)

### $^{1}$H-NMR spectrum of S10, peaks of diastereomer 2

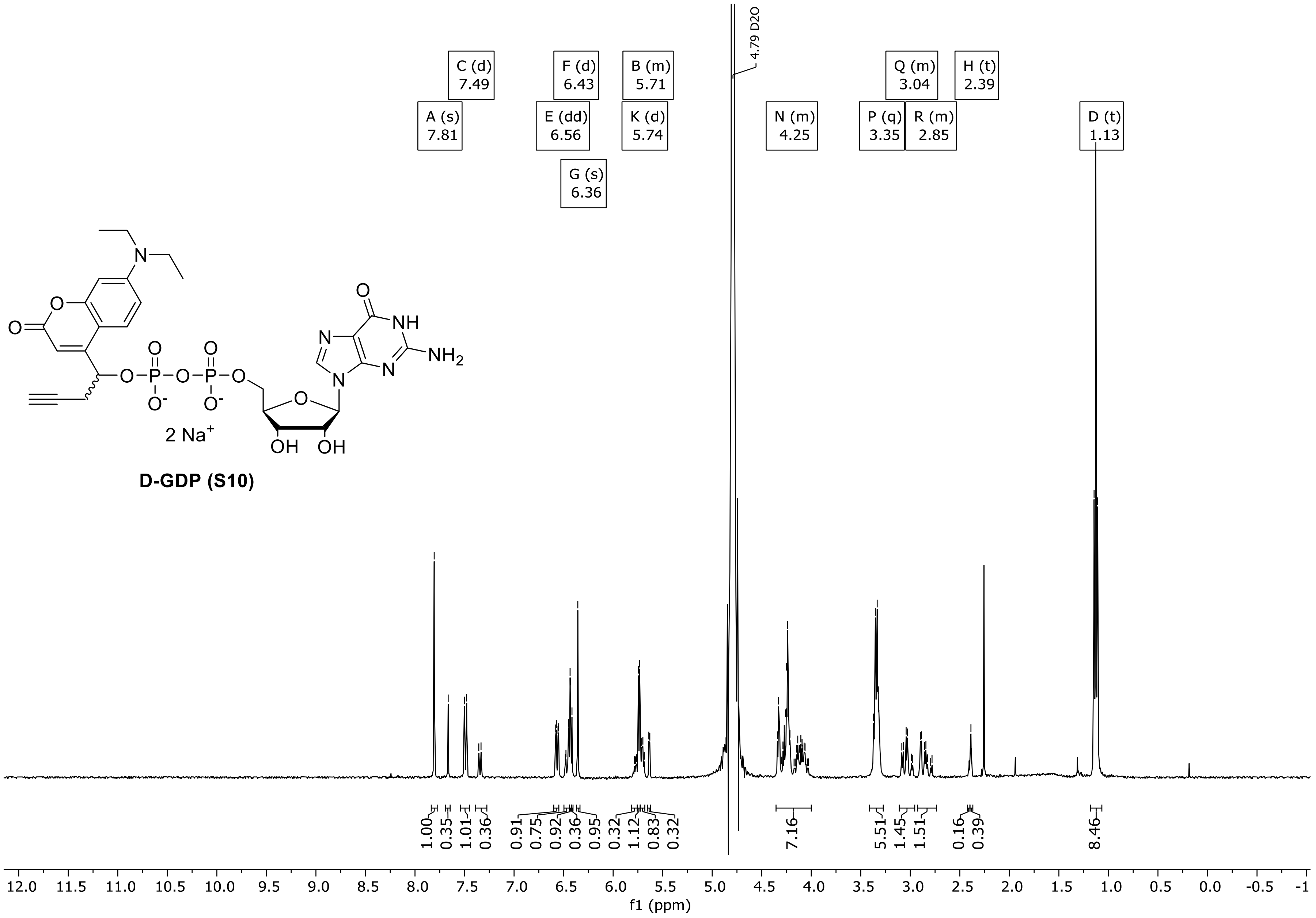

D-GDP (S10)

### $^{31}$P-NMR spectrum of S10, peaks of diastereomere 1

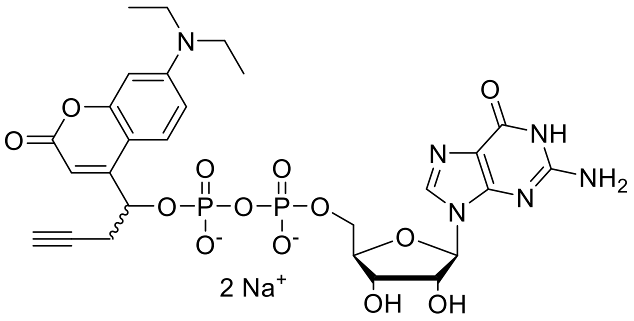

**D-GDP (S10)**

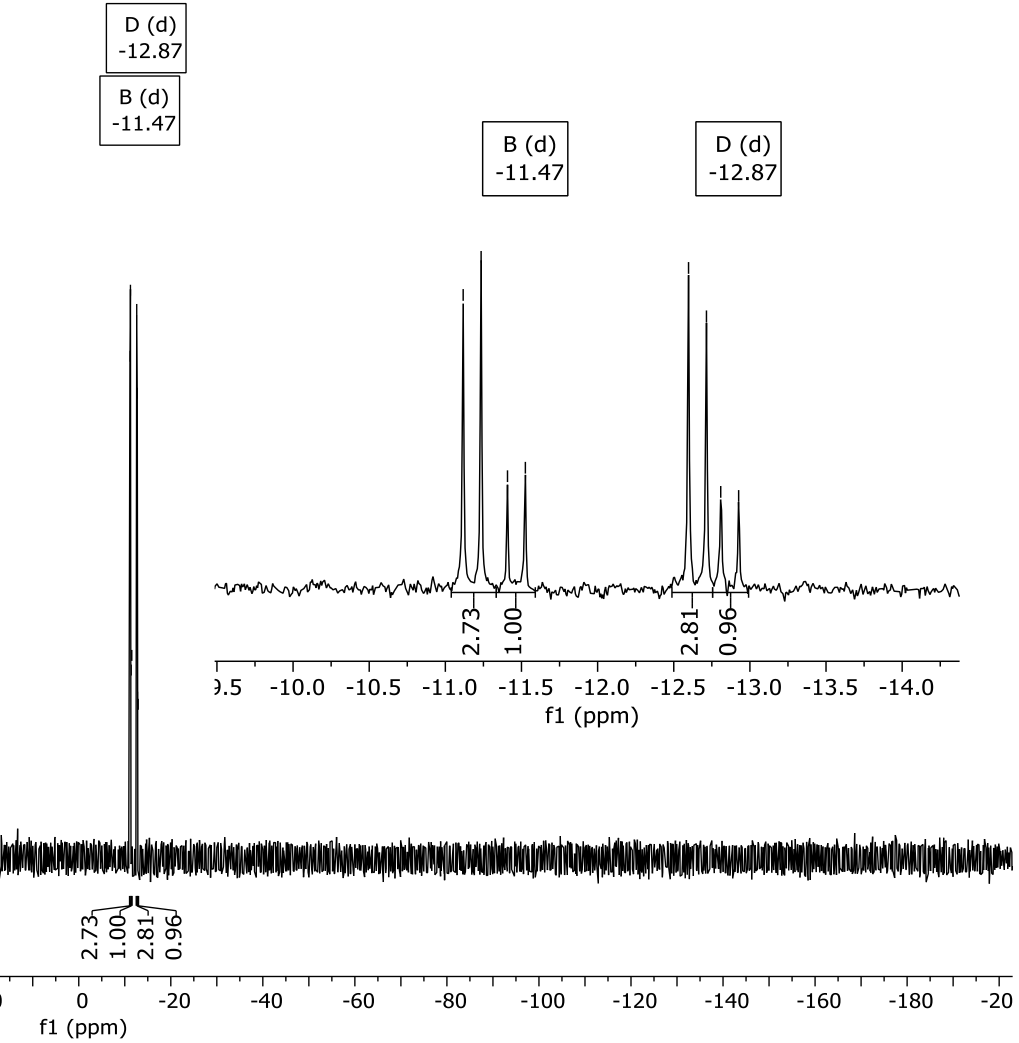

### $^{31}$P-NMR spectrum of S10, peaks of diastereomer 2

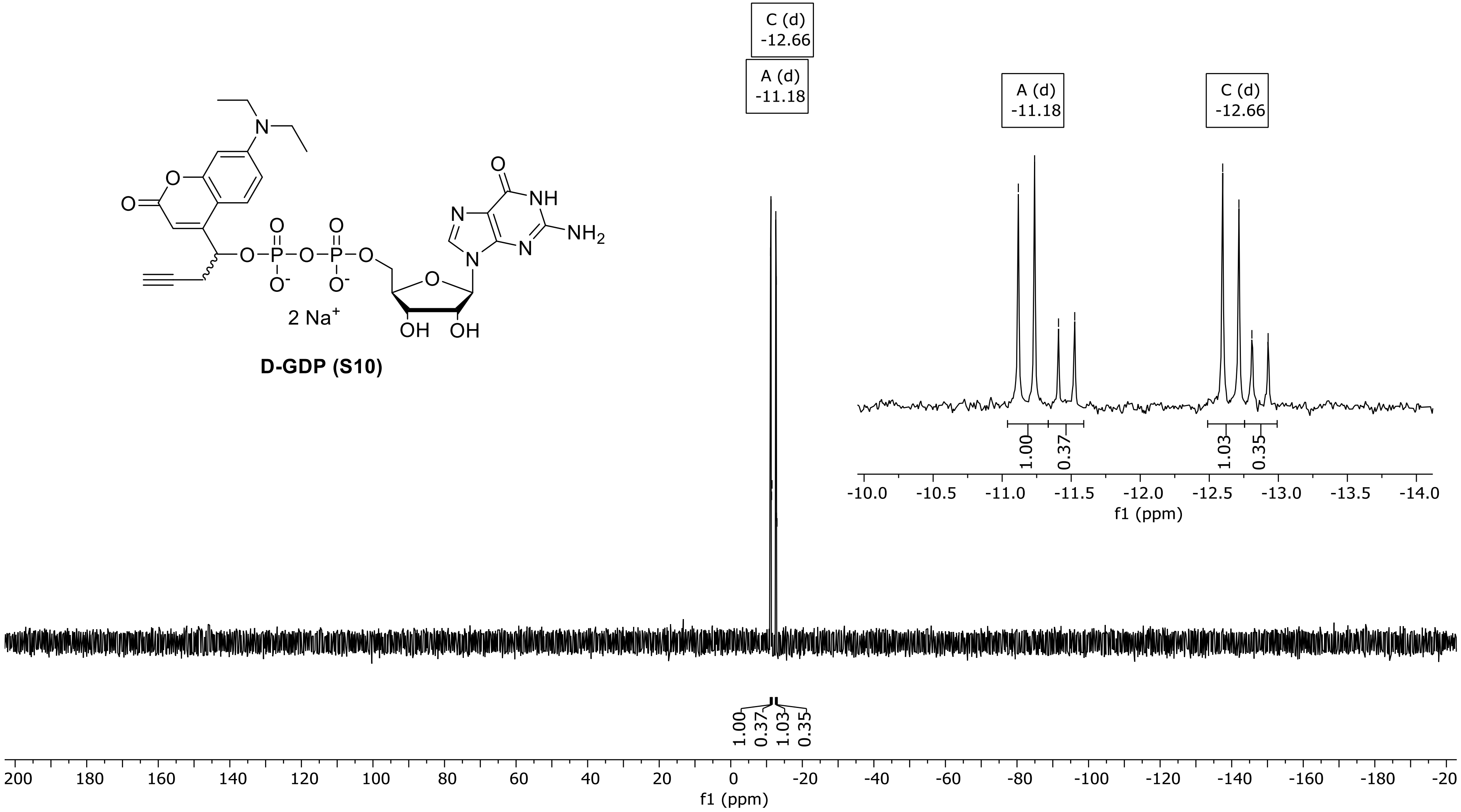

### $^{13}C$ {$^{1}H$} -NMR of S10

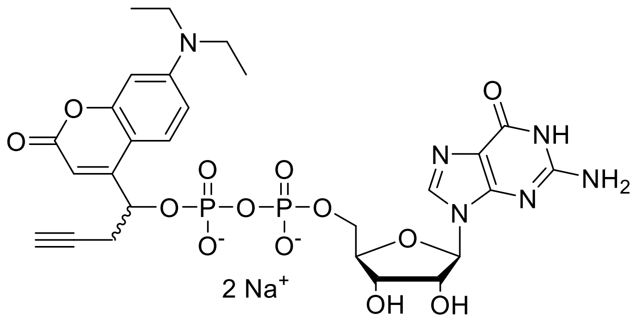

**D-GDP (S10)**

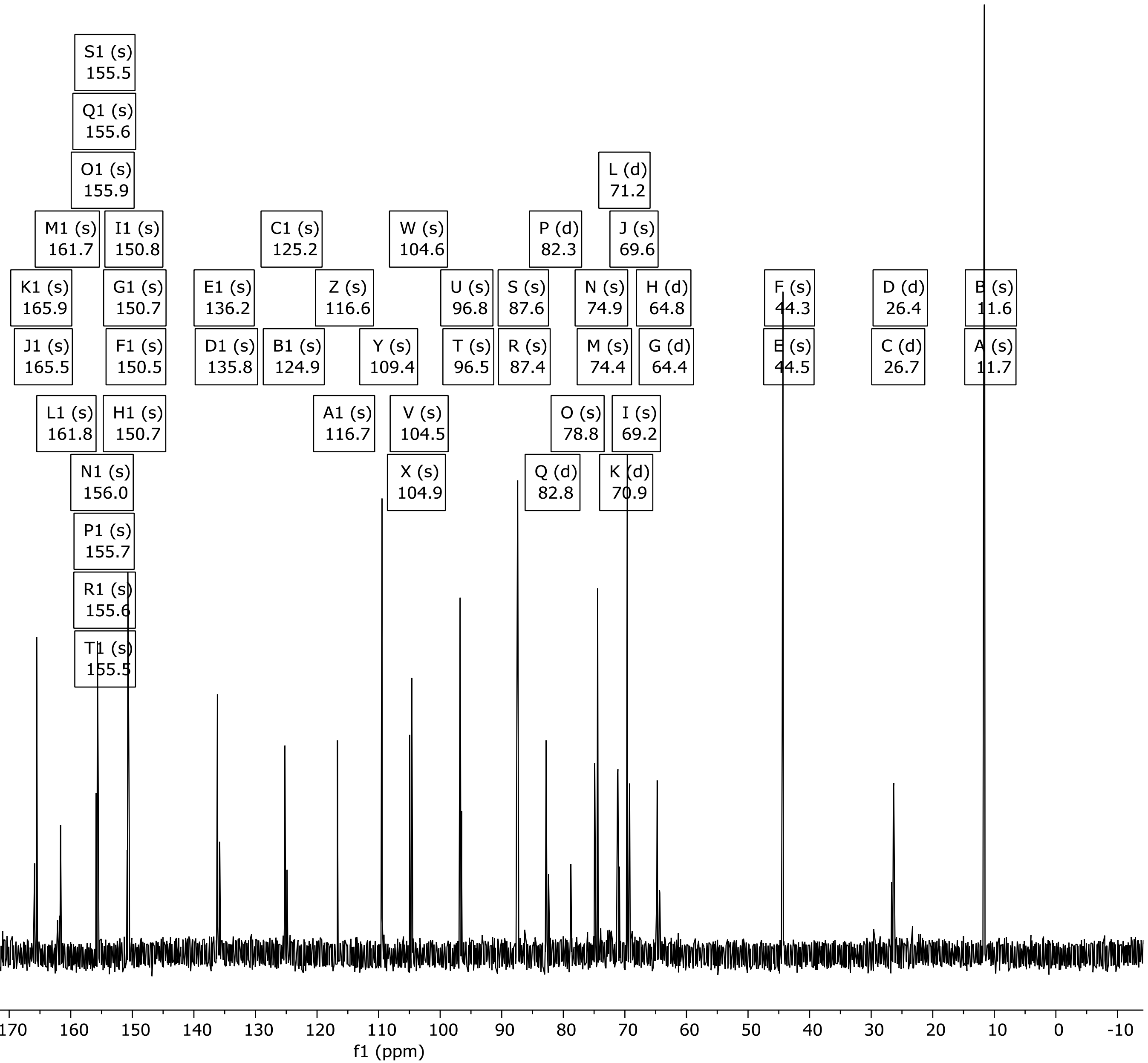

### 1H-NMR spectrum of S12

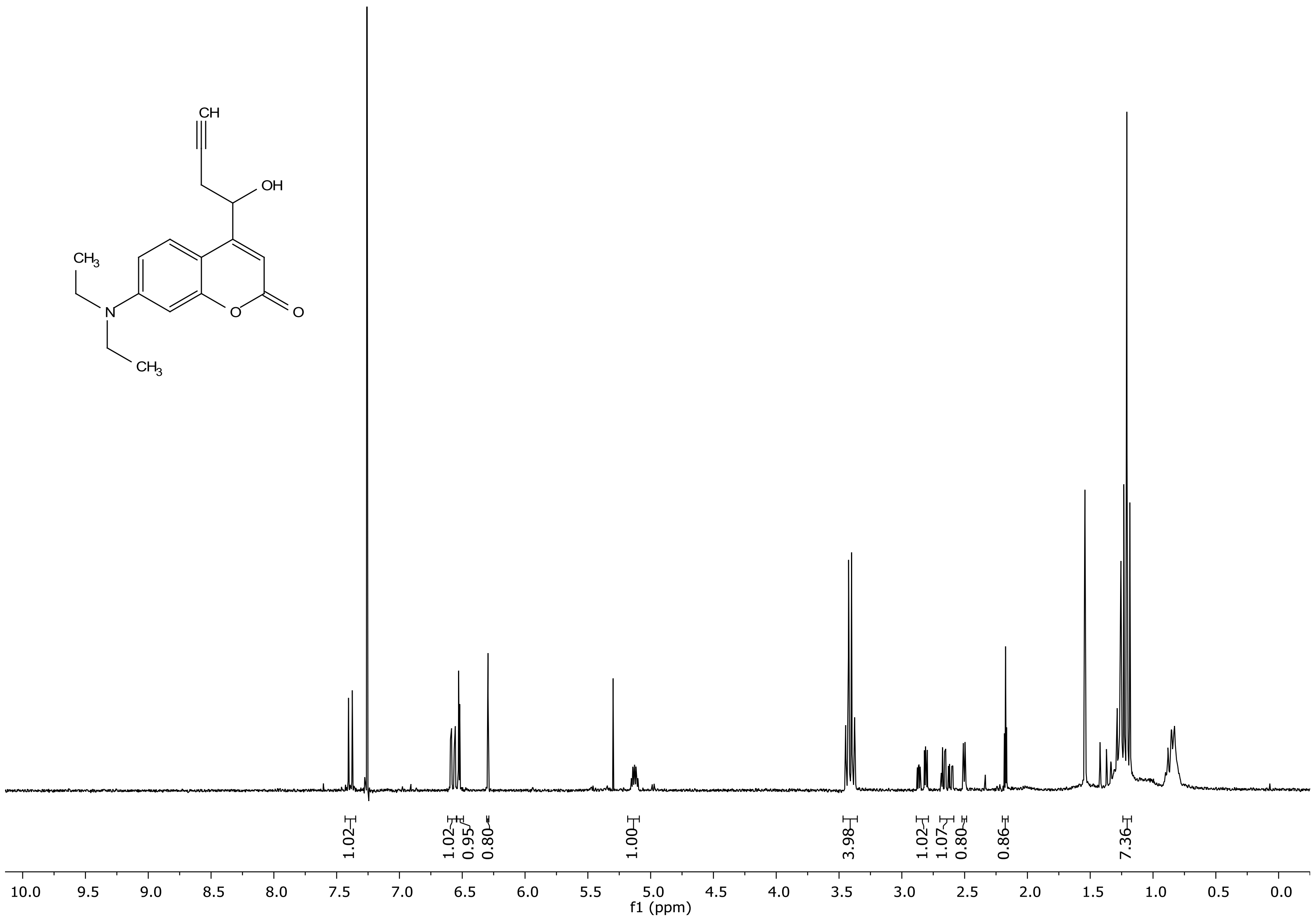

### $^{13}C$ {$^{1}H$}-NMR spectrum of S12

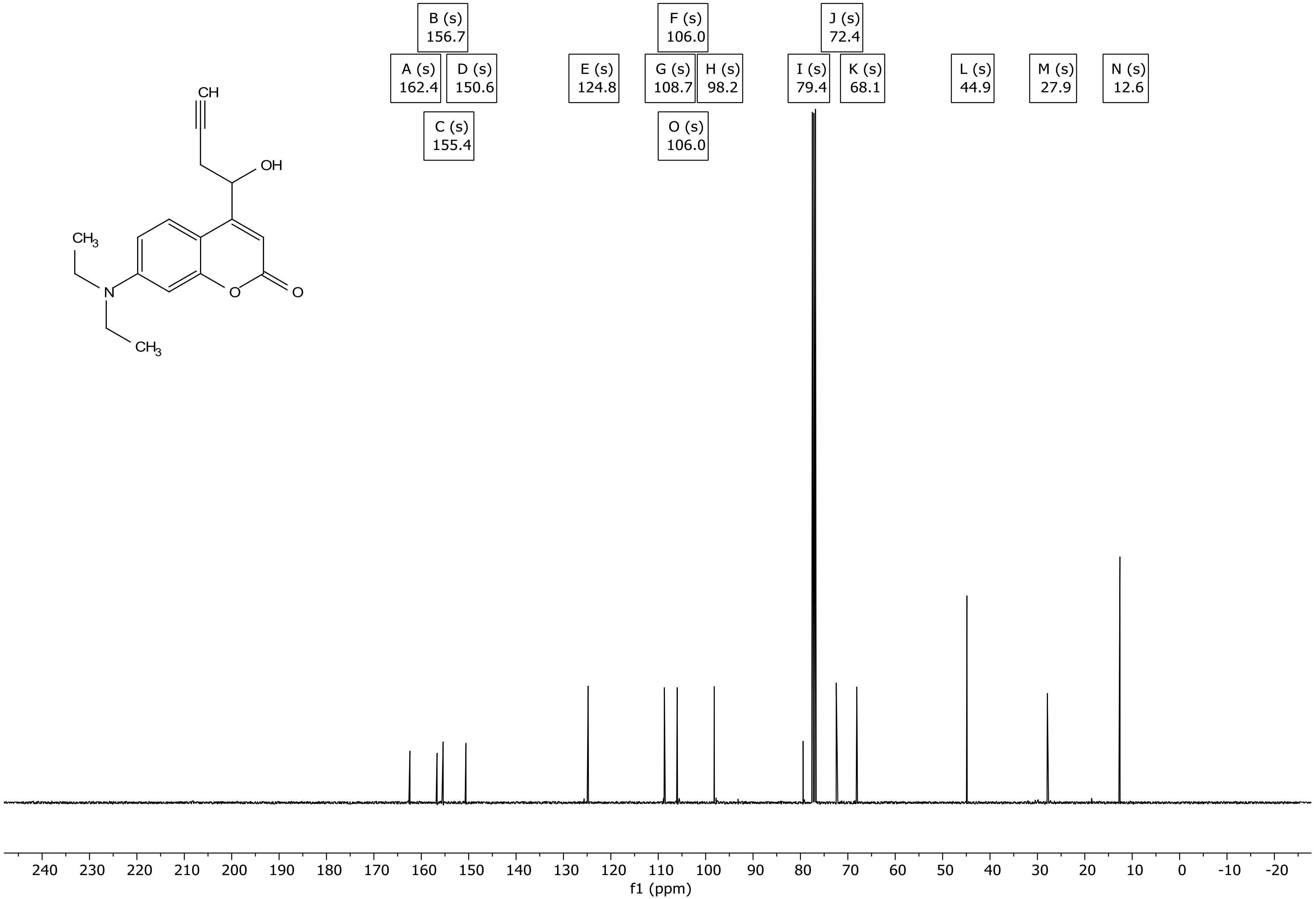

### HRMS (ESI$^+$) of 10

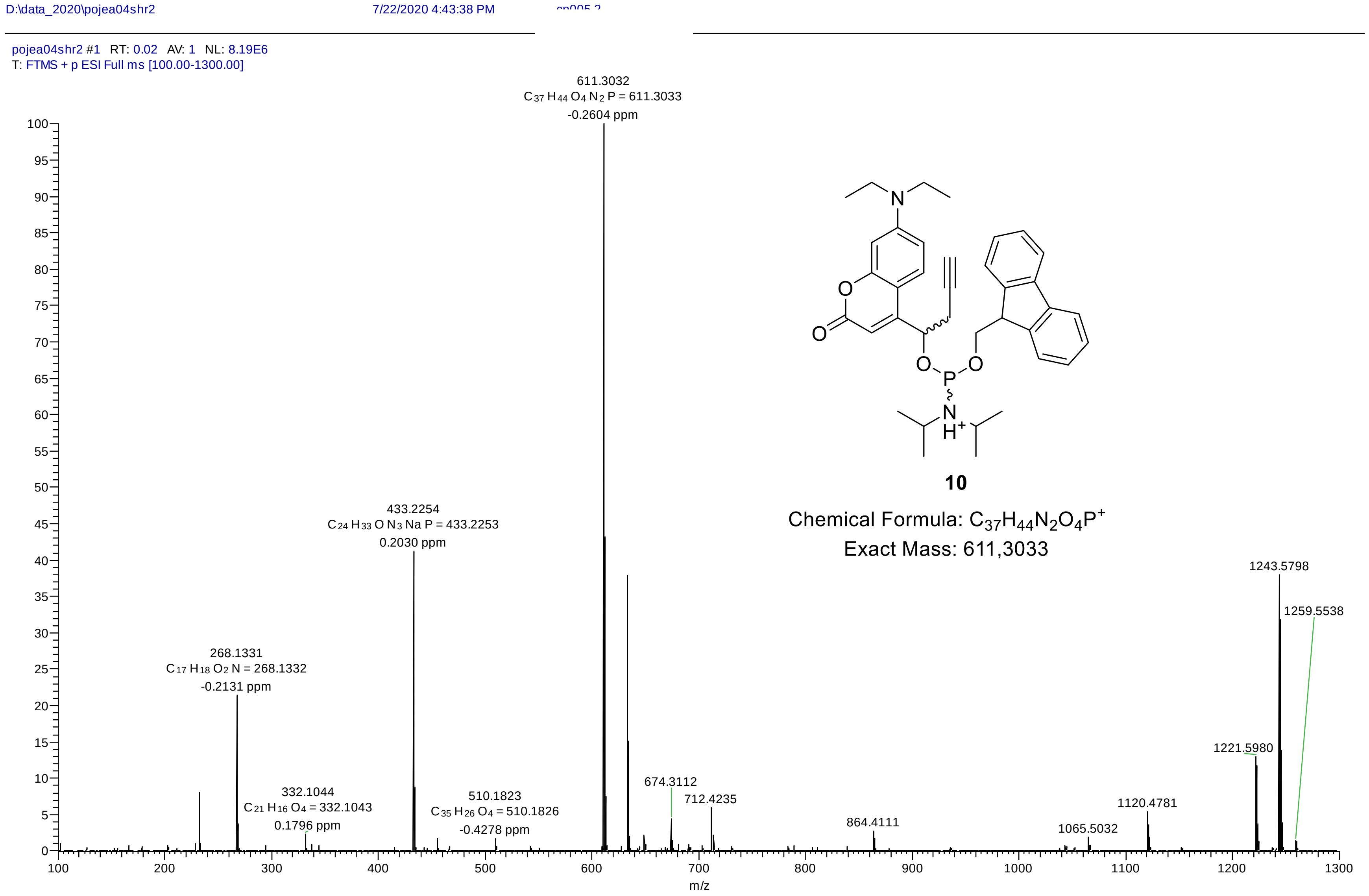

### HRMS (ESI$^-$) of 11

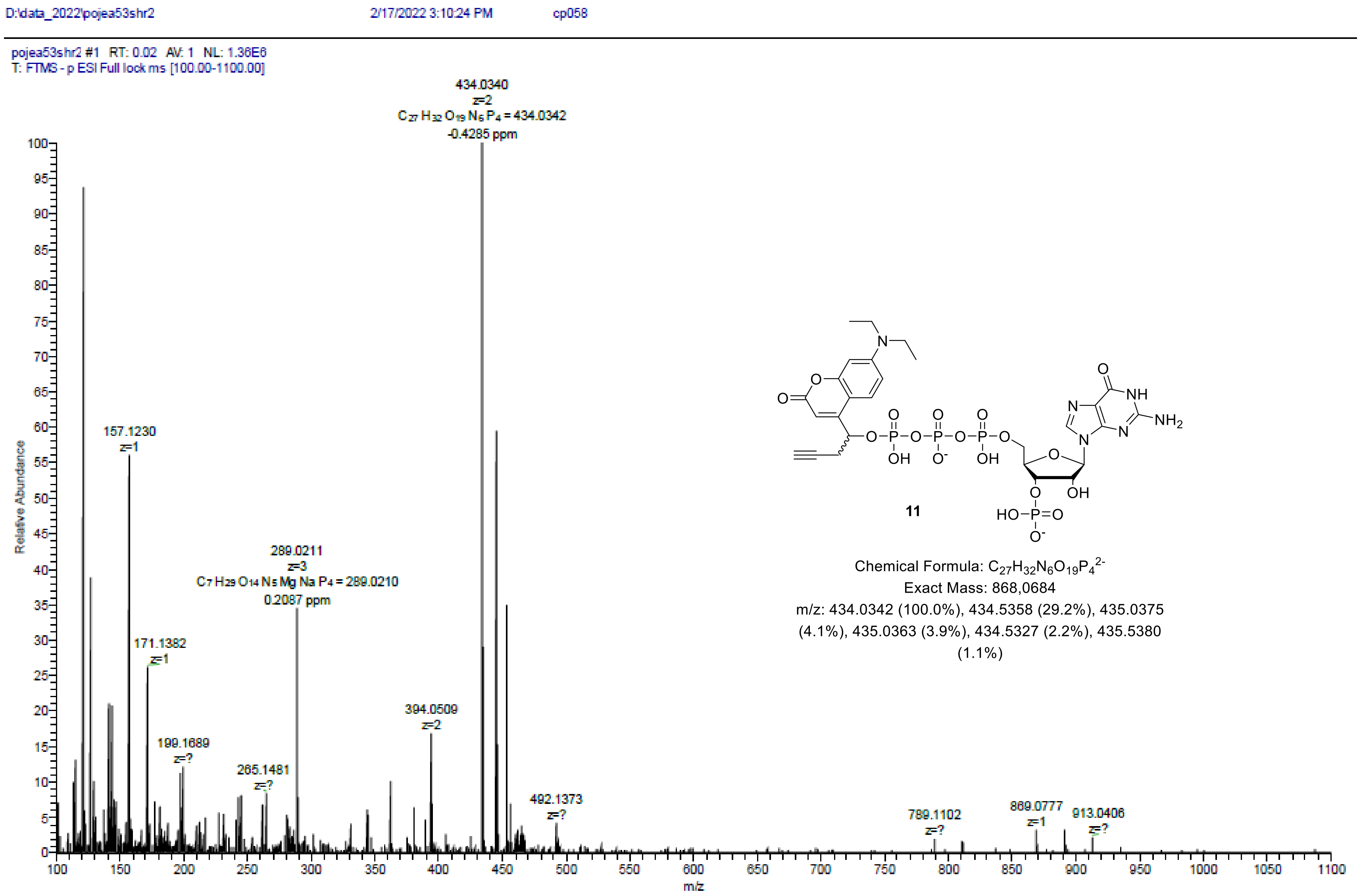

11

Chemical Formula: $C_{27}H_{32}N_6O_{19}P_4^{2-}$
Exact Mass: 868,0684
m/z: 434.0342 (100.0%), 434.5358 (29.2%), 435.0375 (4.1%), 435.0363 (3.9%), 434.5327 (2.2%), 435.5380 (1.1%)

### HRMS (ESI$^-$) of 12

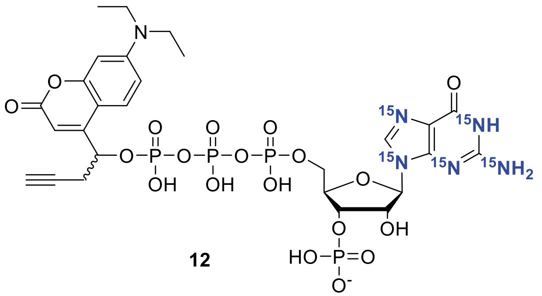

Chemical Formula: $C_{27}H_{33}N^{15}N_5O_{19}P_4^-$
Exact Mass: 874,0608
m/z: 874.0608 (100.0%), 875.0641 (29.2%), 876.0675 (4.1%), 876.0650 (3.9%), 877.0684 (1.1%)

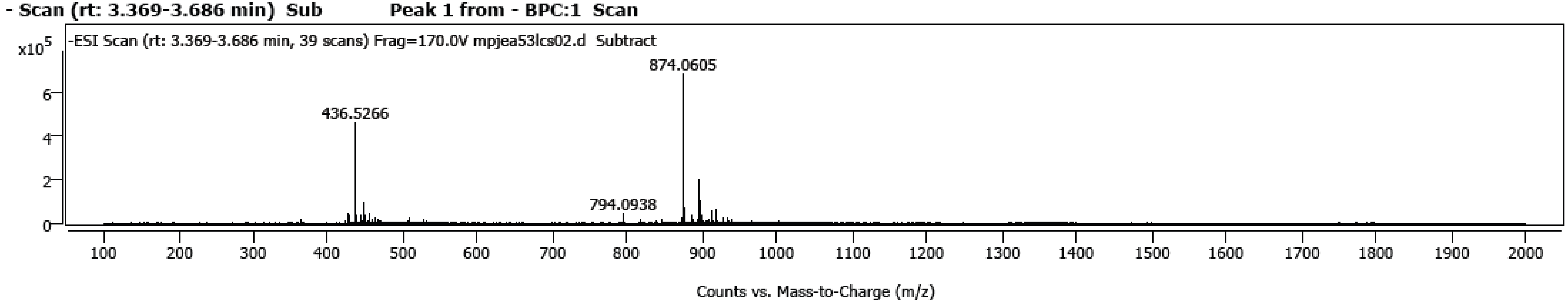

*Spectrum Peaks*

| m/z | Z | Abund | Abund % | m/z (Calc) | Diff (ppm) | Ion Species | Formula | Ion Type |
|---|---|---|---|---|---|---|---|---|
| 426.0509 | 1 | 40857 | 5.99 | 426.0510 | -0.22 | (M-H)- | C21 H16 O8 P | |
| 436.5266 | 2 | 459228 | 67.32 | 436.5268 | -0.32 | (M-2H)-2 | C27 H34 N [15N]5 O19 P4 | |
| 437.0290 | 2 | 122523 | 17.96 | 437.0284 | 1.35 | (M-2H)-2 | C27 H34 N [15N]5 O19 P4 | |
| 794.0938 | 1 | 43225 | 6.34 | 794.0950 | -1.42 | (M-H)- | C27 H23 N6 [15N]4 O19 | |
| 874.0605 | 1 | 682115 | 100.00 | 874.0608 | -0.36 | (M-H)- | C27 H34 N [15N]5 O19 P4 | |
| 875.0637 | 1 | 225772 | 33.10 | 875.0642 | -0.53 | (M-H)- | C27 H34 N [15N]5 O19 P4 | |
| 876.0657 | 1 | 67749 | 9.93 | 876.0664 | -0.76 | (M-H)- | C27 H34 N [15N]5 O19 P4 | |
| 896.0419 | 1 | 195995 | 28.73 | 896.0416 | 0.34 | (M-H)- | C27 H30 N5 [15N]3 O19 P4 | |
| 897.0435 | 1 | 68909 | 10.10 | 897.0446 | -1.33 | (M-H)- | C27 H30 N5 [15N]3 O19 P4 | |
| 898.0186 | 1 | 100212 | 14.69 | 898.0188 | -0.20 | (M-H)- | C27 H28 Cl N6 [15N] O20 P3 | |
| 447.5177 | | 70268 | 10.30 | | | | | |
| 448.5065 | | 94845 | 13.90 | | | | | |
| 455.5009 | | 44988 | 6.60 | | | | | |
| 912.0078 | | 55400 | 8.12 | | | | | |
| 918.0235 | | 60242 | 8.83 | | | | | |

### HRMS ($ESI^-$) of 13

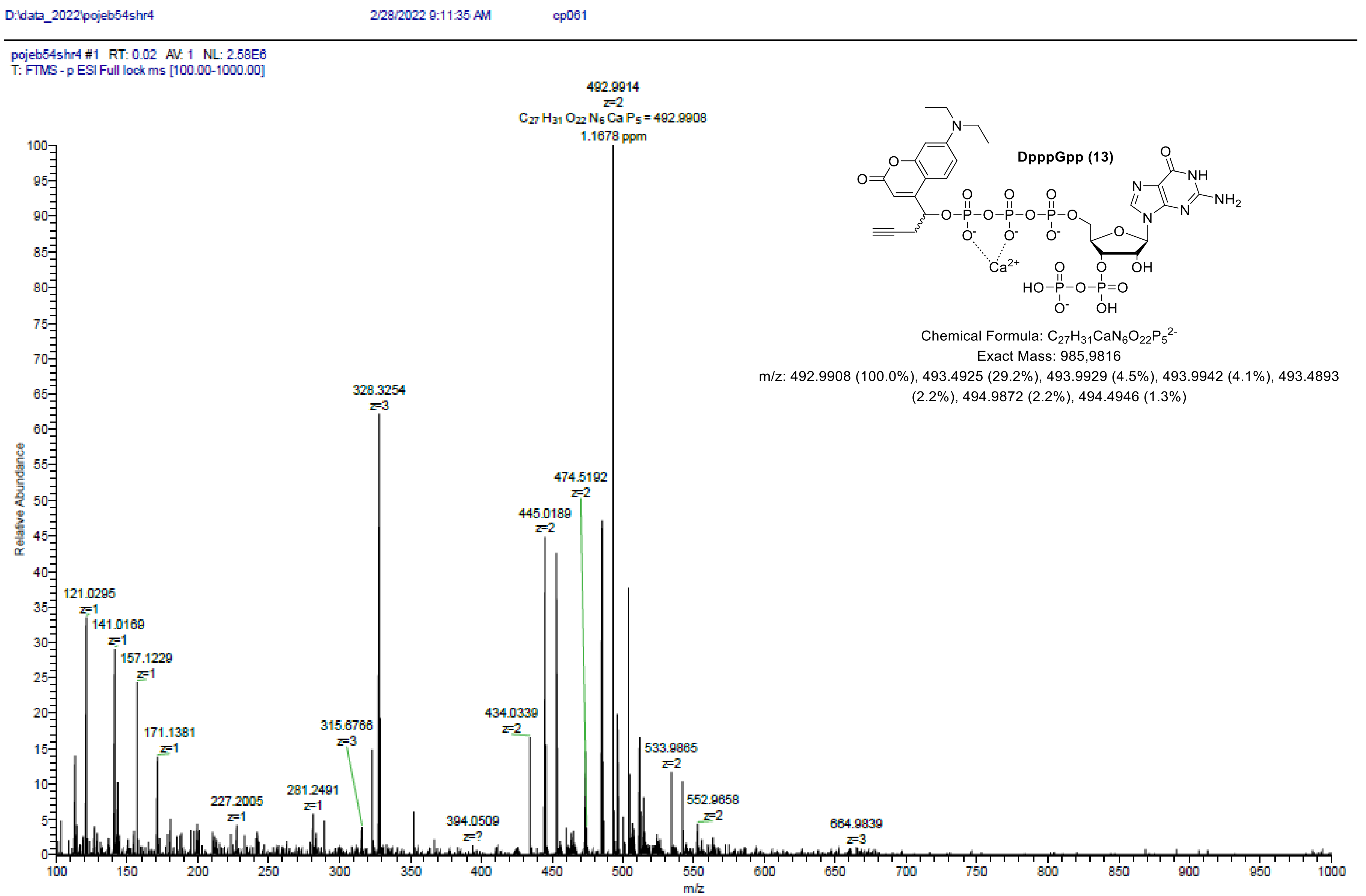

### HRMS (ESI$^-$) of 14

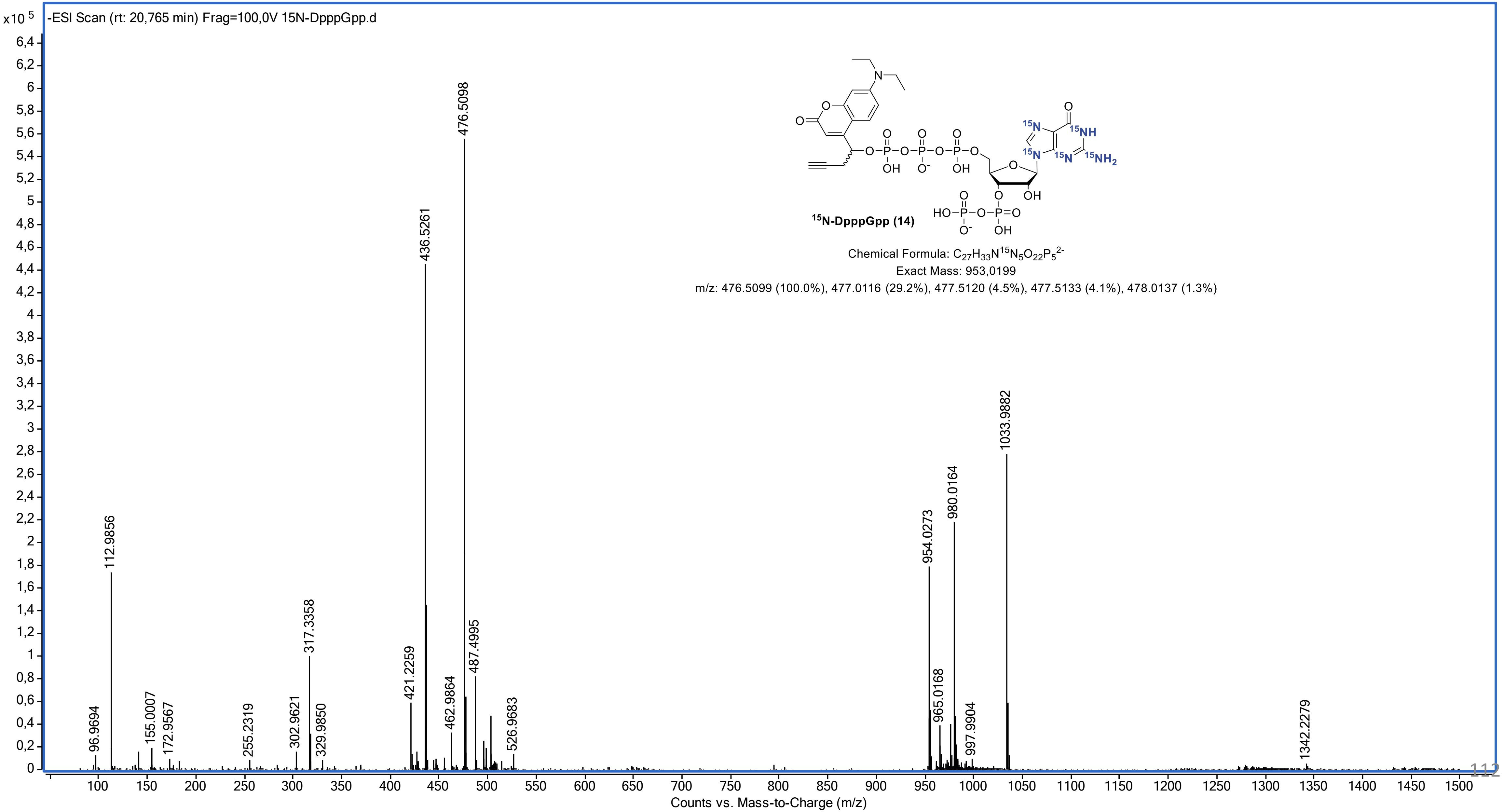

### HRMS (ESI$^-$) of 15

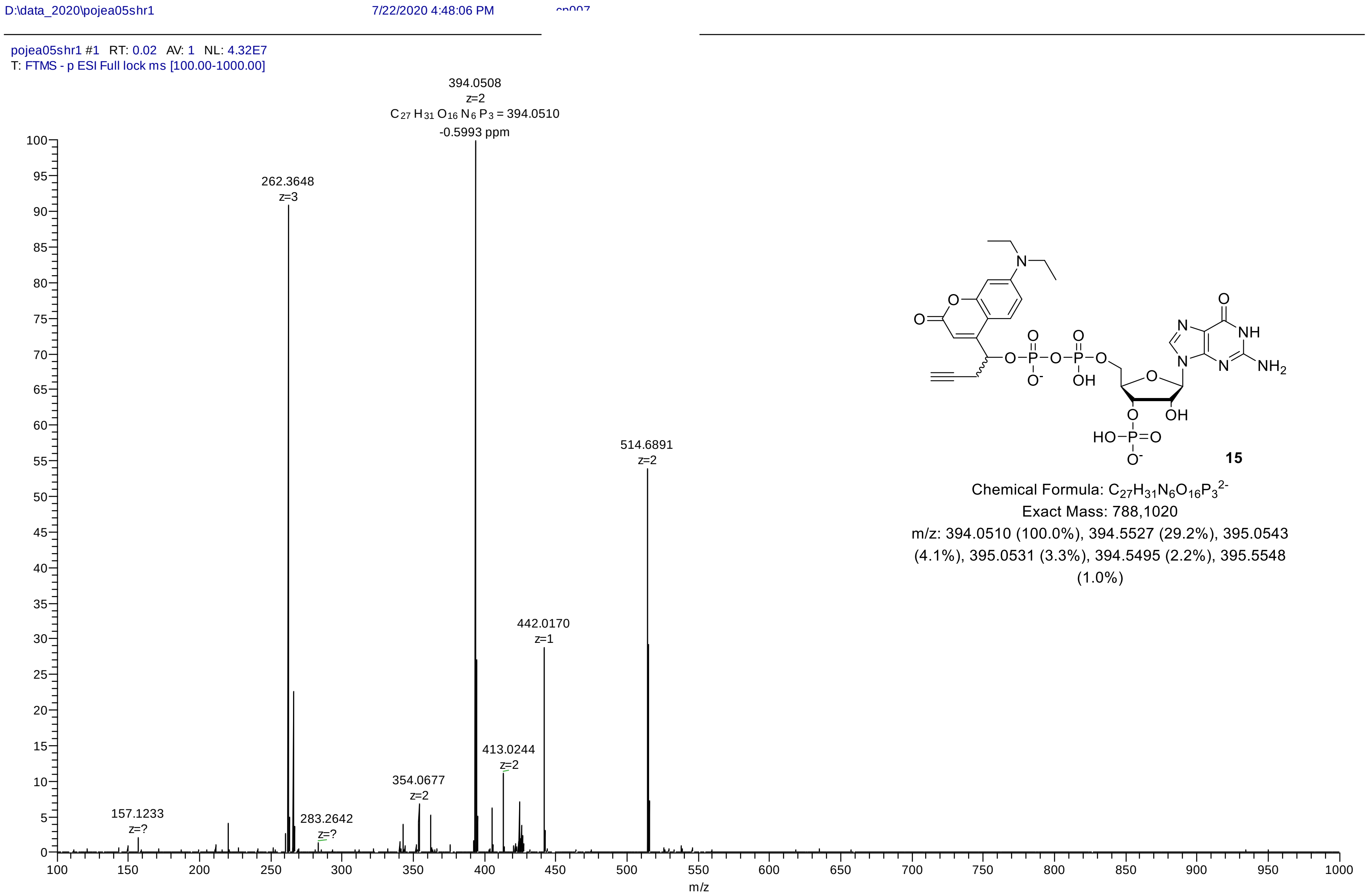

**15**

Chemical Formula: $C_{27}H_{31}N_6O_{16}P_3^{2-}$
Exact Mass: 788,1020
m/z: 394.0510 (100.0%), 394.5527 (29.2%), 395.0543 (4.1%), 395.0531 (3.3%), 394.5495 (2.2%), 395.5548 (1.0%)

### HRMS (ESI⁻) of 16

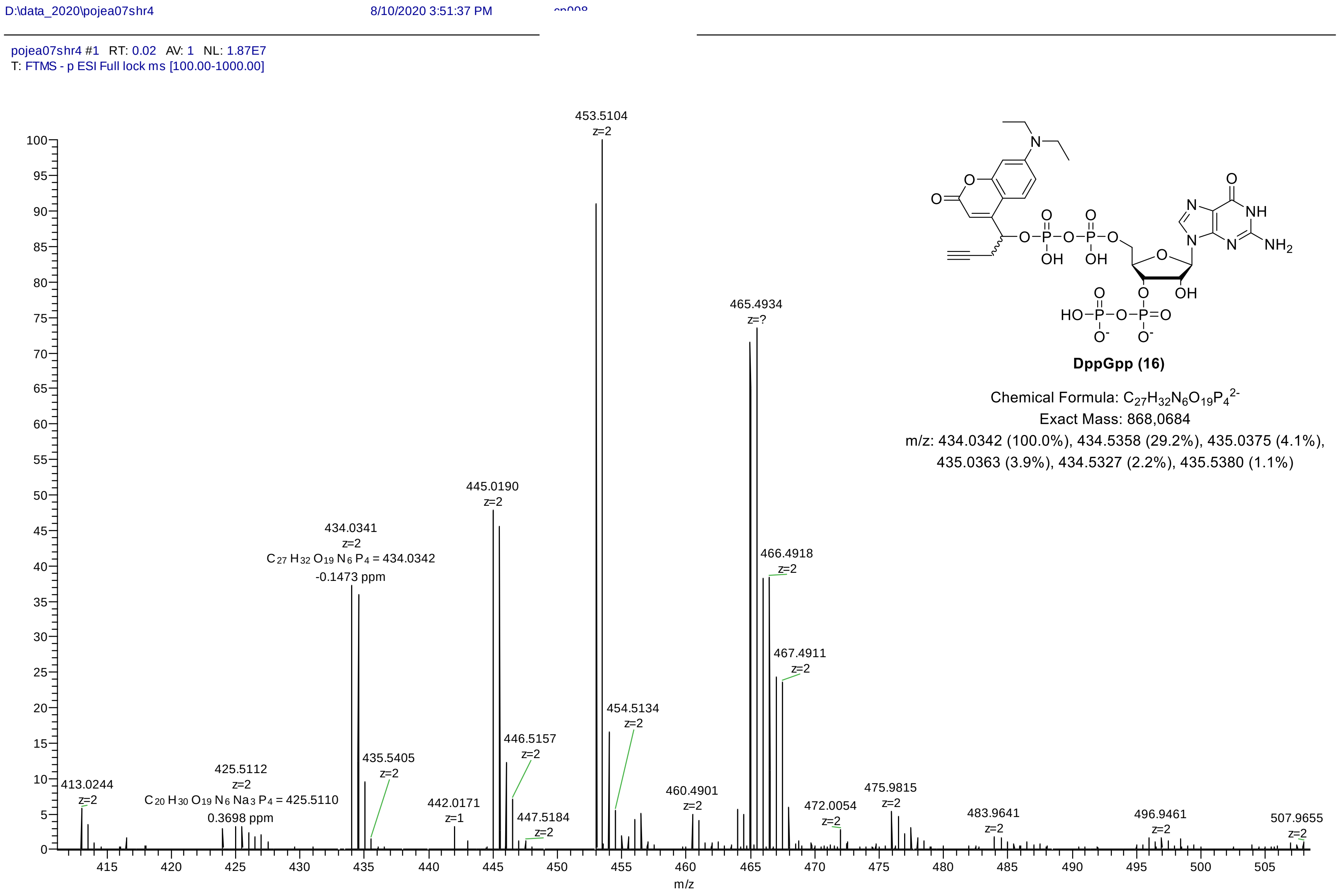

**DppGpp (16)**

Chemical Formula: $C_{27}H_{32}N_6O_{19}P_4^{2-}$

Exact Mass: 868,0684

m/z: 434.0342 (100.0%), 434.5358 (29.2%), 435.0375 (4.1%), 435.0363 (3.9%), 434.5327 (2.2%), 435.5380 (1.1%)

### HRMS (ESI$^-$) of S1

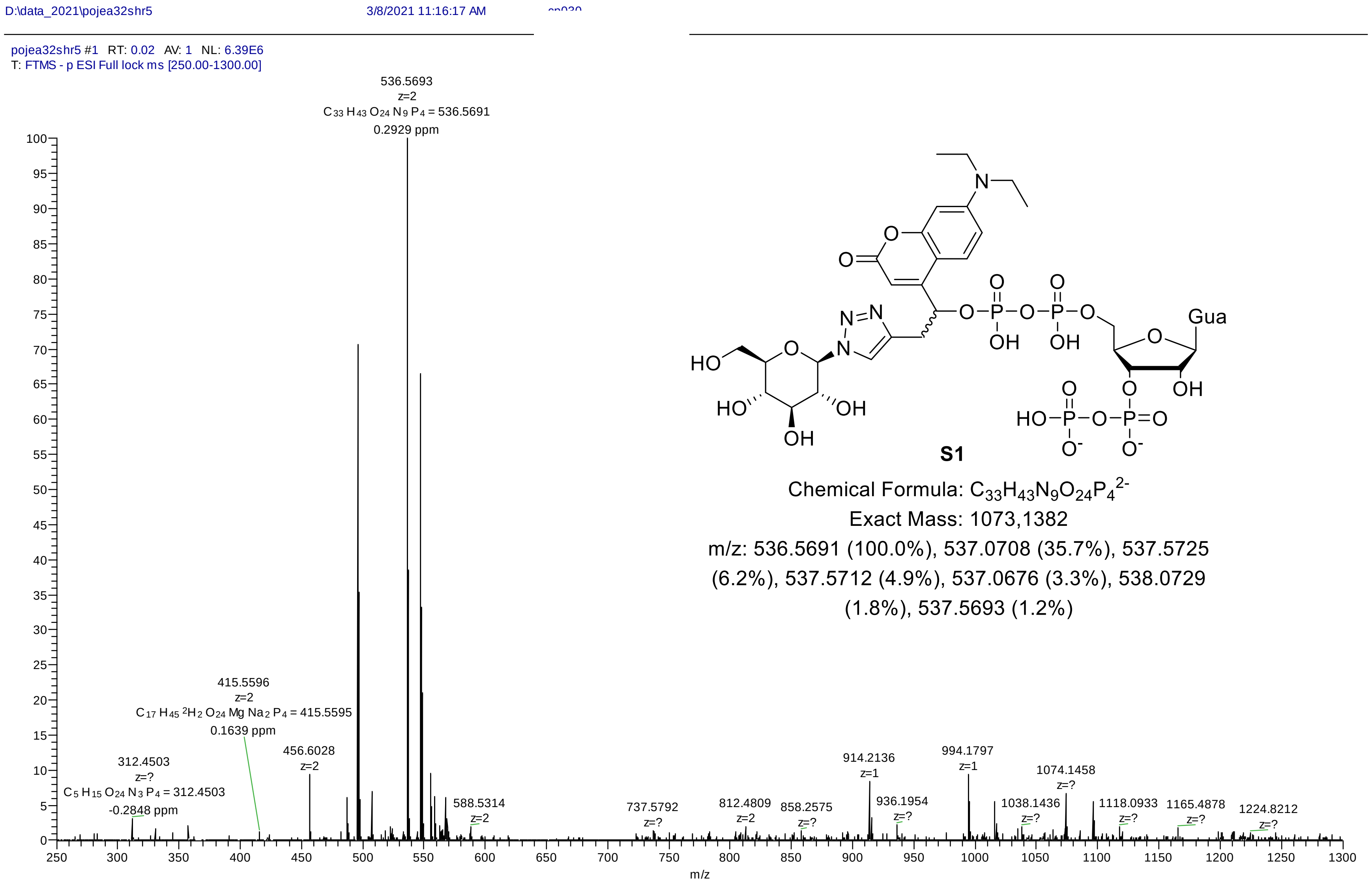

**S1**

Chemical Formula: $C_{33}H_{43}N_9O_{24}P_4^{2-}$

Exact Mass: 1073,1382

m/z: 536.5691 (100.0%), 537.0708 (35.7%), 537.5725 (6.2%), 537.5712 (4.9%), 537.0676 (3.3%), 538.0729 (1.8%), 537.5693 (1.2%)

### HRMS (ESI$^-$) of S2

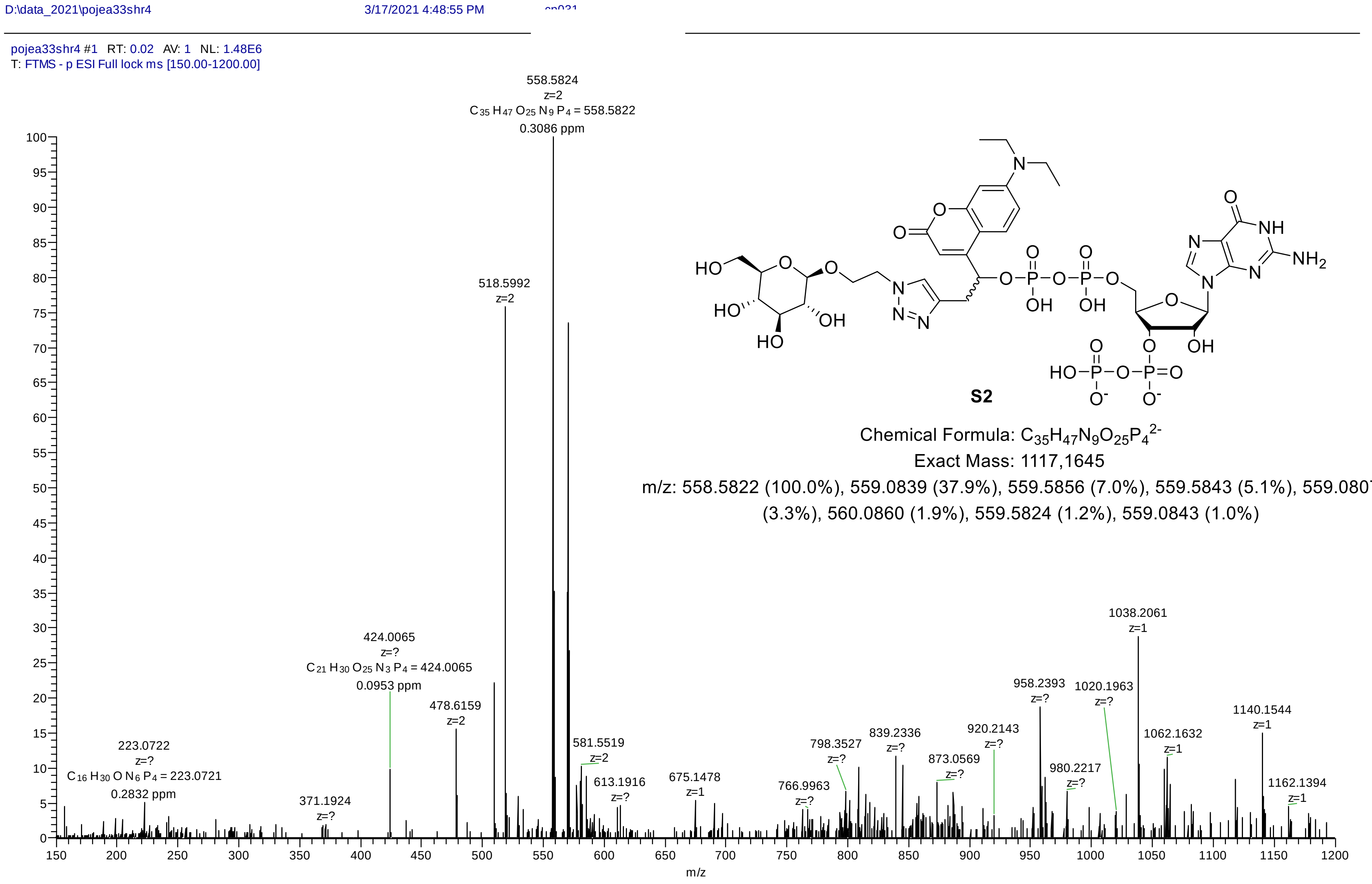

**S2**

Chemical Formula: $C_{35}H_{47}N_9O_{25}P_4^{2-}$

Exact Mass: 1117,1645

m/z: 558.5822 (100.0%), 559.0839 (37.9%), 559.5856 (7.0%), 559.5843 (5.1%), 559.0807 (3.3%), 560.0860 (1.9%), 559.5824 (1.2%), 559.0843 (1.0%)

### HRMS (ESI$^-$) of S3

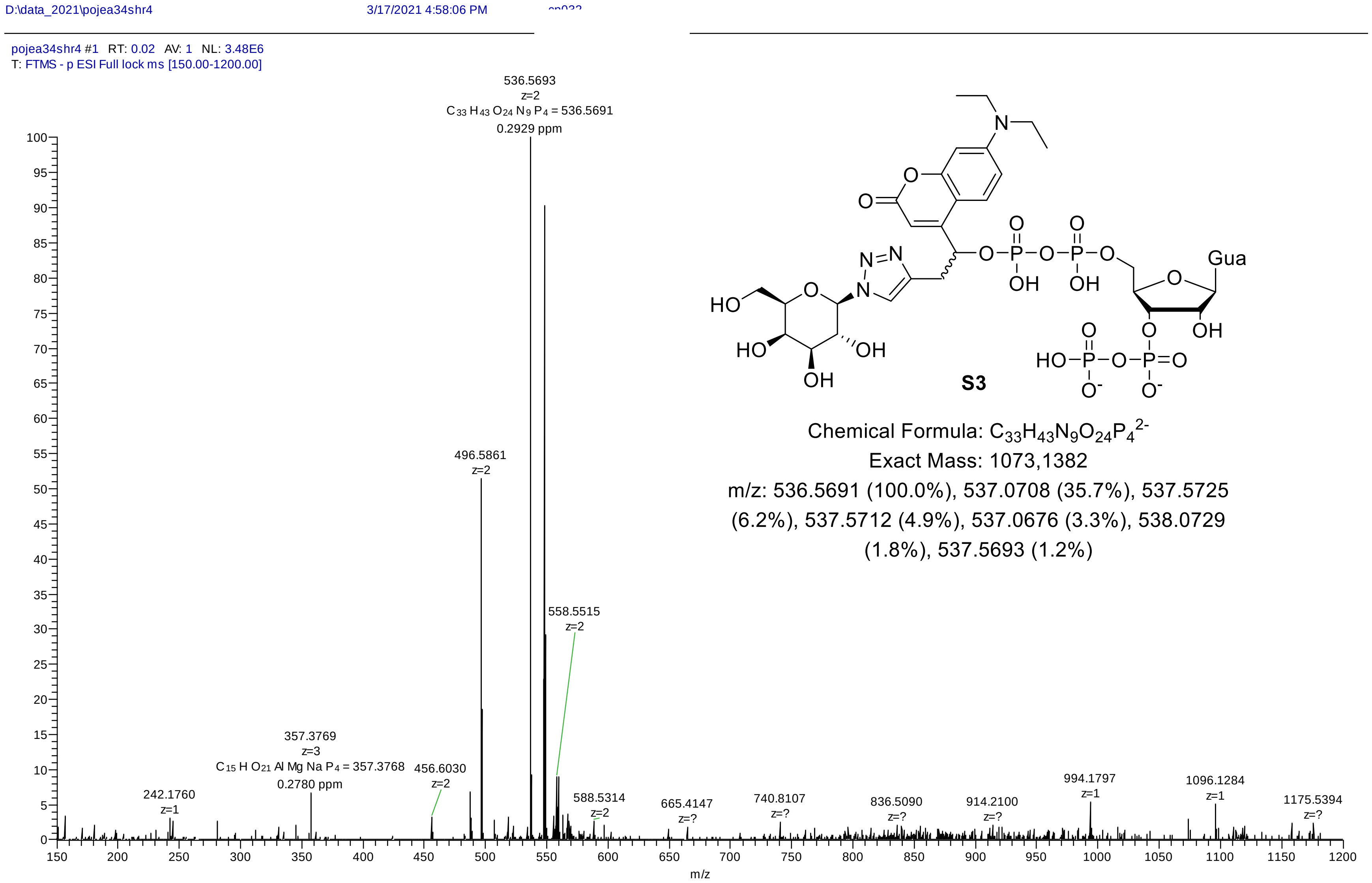

**S3**

Chemical Formula: $C_{33}H_{43}N_9O_{24}P_4^{2-}$

Exact Mass: 1073,1382

m/z: 536.5691 (100.0%), 537.0708 (35.7%), 537.5725 (6.2%), 537.5712 (4.9%), 537.0676 (3.3%), 538.0729 (1.8%), 537.5693 (1.2%)

### HRMS (ESI$^-$) of S4

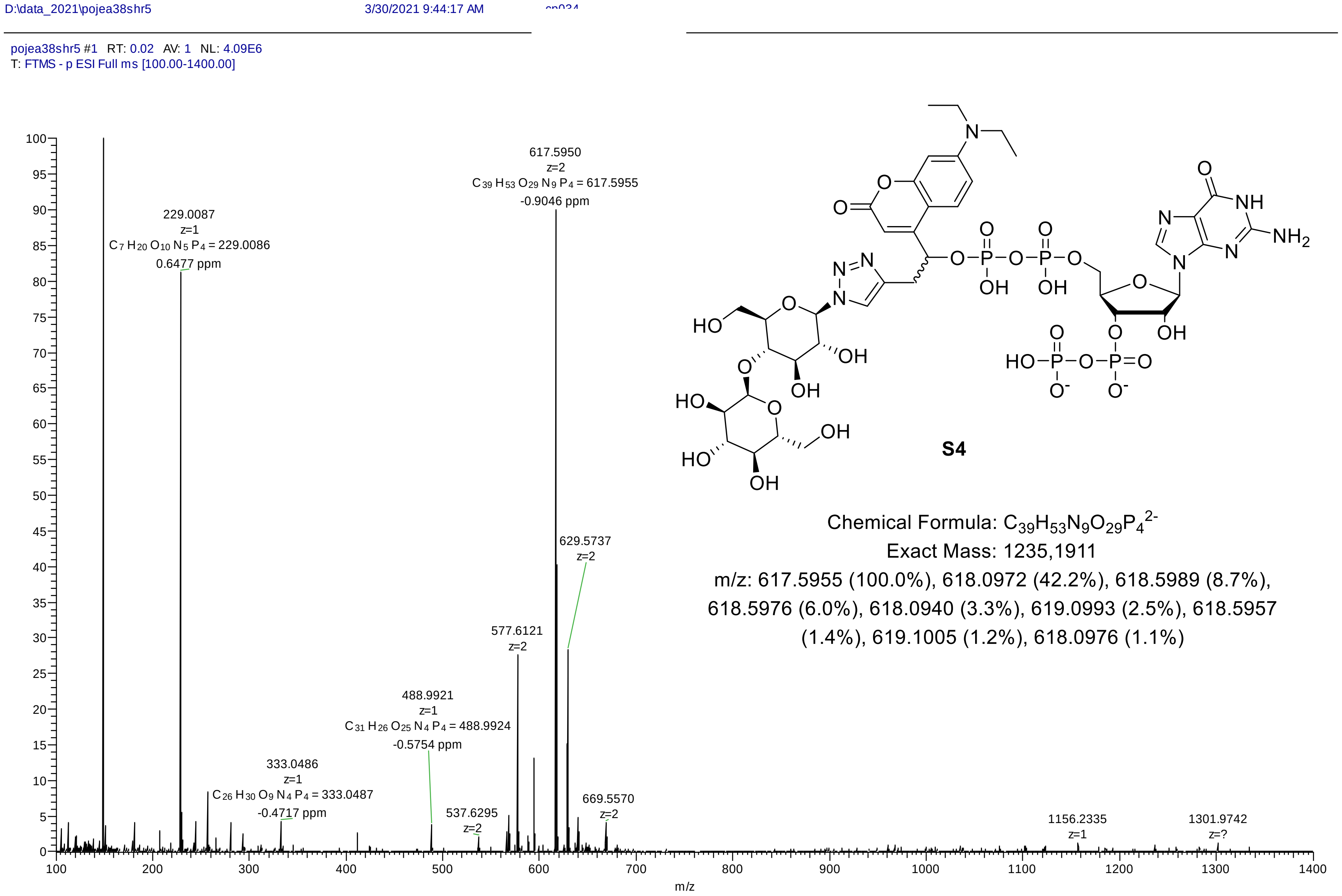

**S4**

Chemical Formula: $C_{39}H_{53}N_9O_{29}P_4^{2-}$

Exact Mass: 1235,1911

m/z: 617.5955 (100.0%), 618.0972 (42.2%), 618.5989 (8.7%), 618.5976 (6.0%), 618.0940 (3.3%), 619.0993 (2.5%), 618.5957 (1.4%), 619.1005 (1.2%), 618.0976 (1.1%)

### HRMS (ESI$^-$) of S5

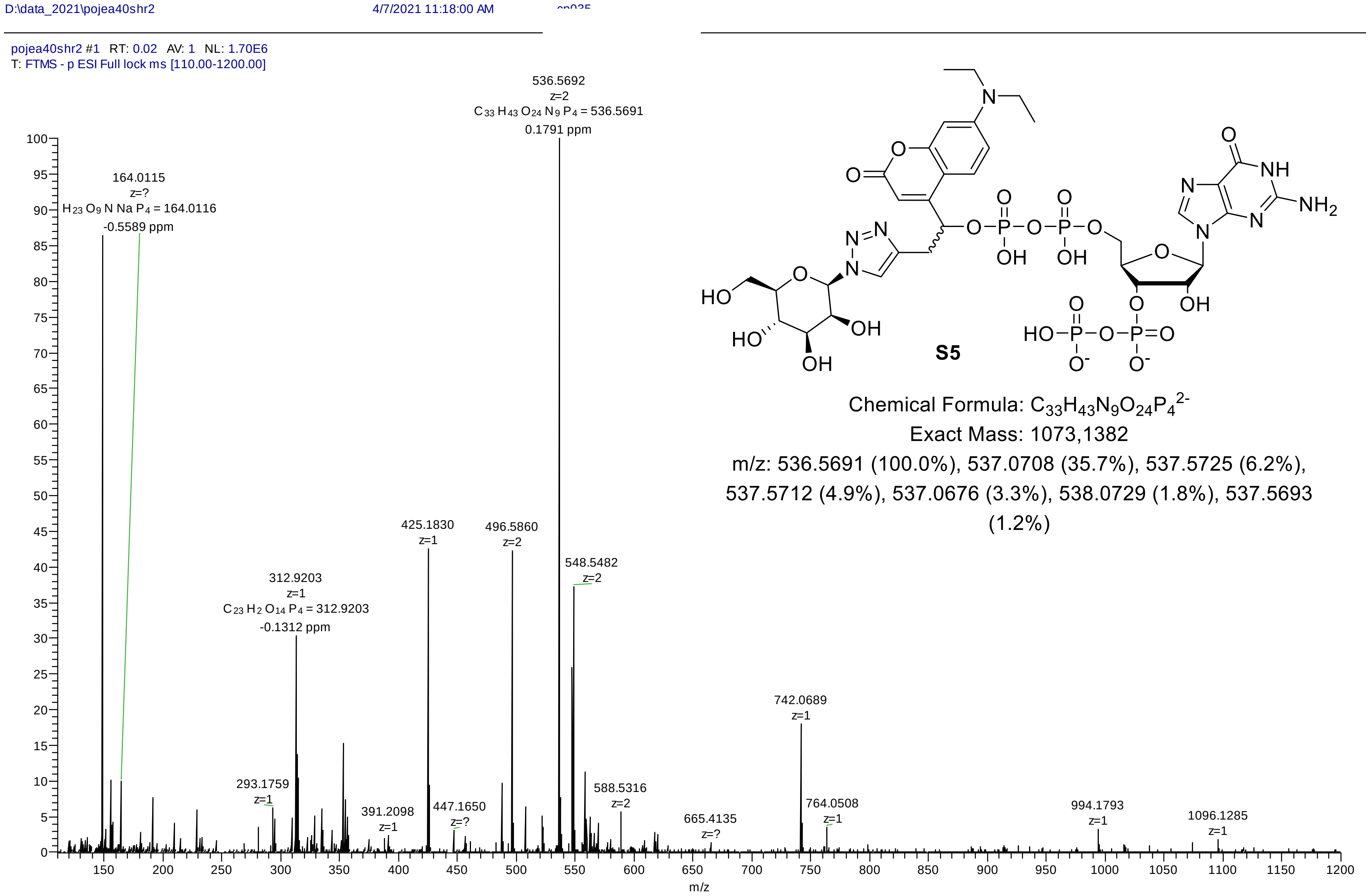

S5

Chemical Formula: $C_{33}H_{43}N_9O_{24}P_4^{2-}$

Exact Mass: 1073,1382

m/z: 536.5691 (100.0%), 537.0708 (35.7%), 537.5725 (6.2%), 537.5712 (4.9%), 537.0676 (3.3%), 538.0729 (1.8%), 537.5693 (1.2%)

### HRMS (ESI$^-$) of S6

D:\data_2021\pojea39shr2 3/30/2021 9:54:00 AM

pojea39shr2 #1 RT: 0.02 AV: 1 NL: 3.97E6
T: FTMS - p ESI Full ms [100.00-1400.00]

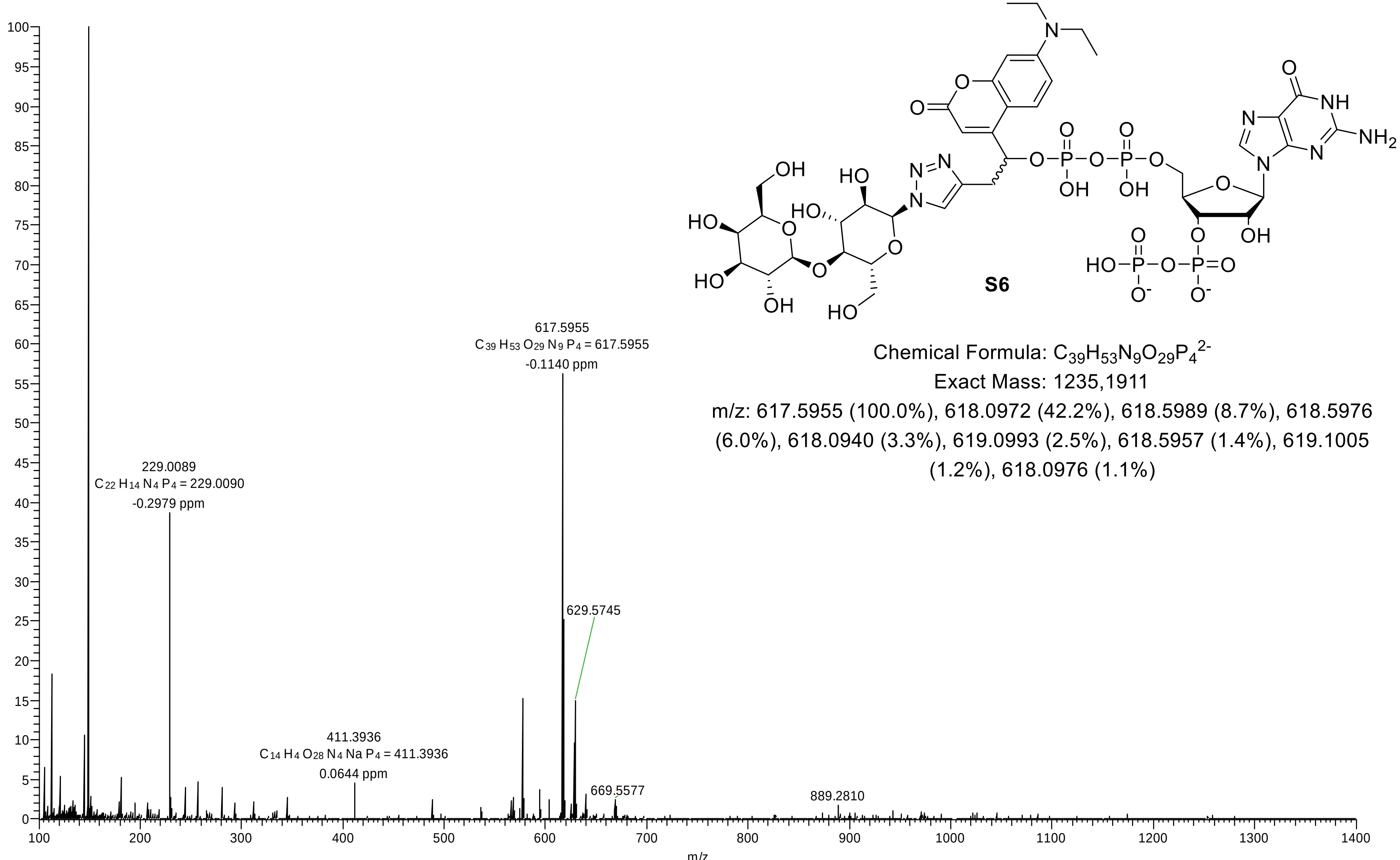

Chemical Formula: $C_{39}H_{53}N_9O_{29}P_4^{2-}$

Exact Mass: 1235,1911

m/z: 617.5955 (100.0%), 618.0972 (42.2%), 618.5989 (8.7%), 618.5976 (6.0%), 618.0940 (3.3%), 619.0993 (2.5%), 618.5957 (1.4%), 619.1005 (1.2%), 618.0976 (1.1%)

### HRMS (ESI$^-$) of S7

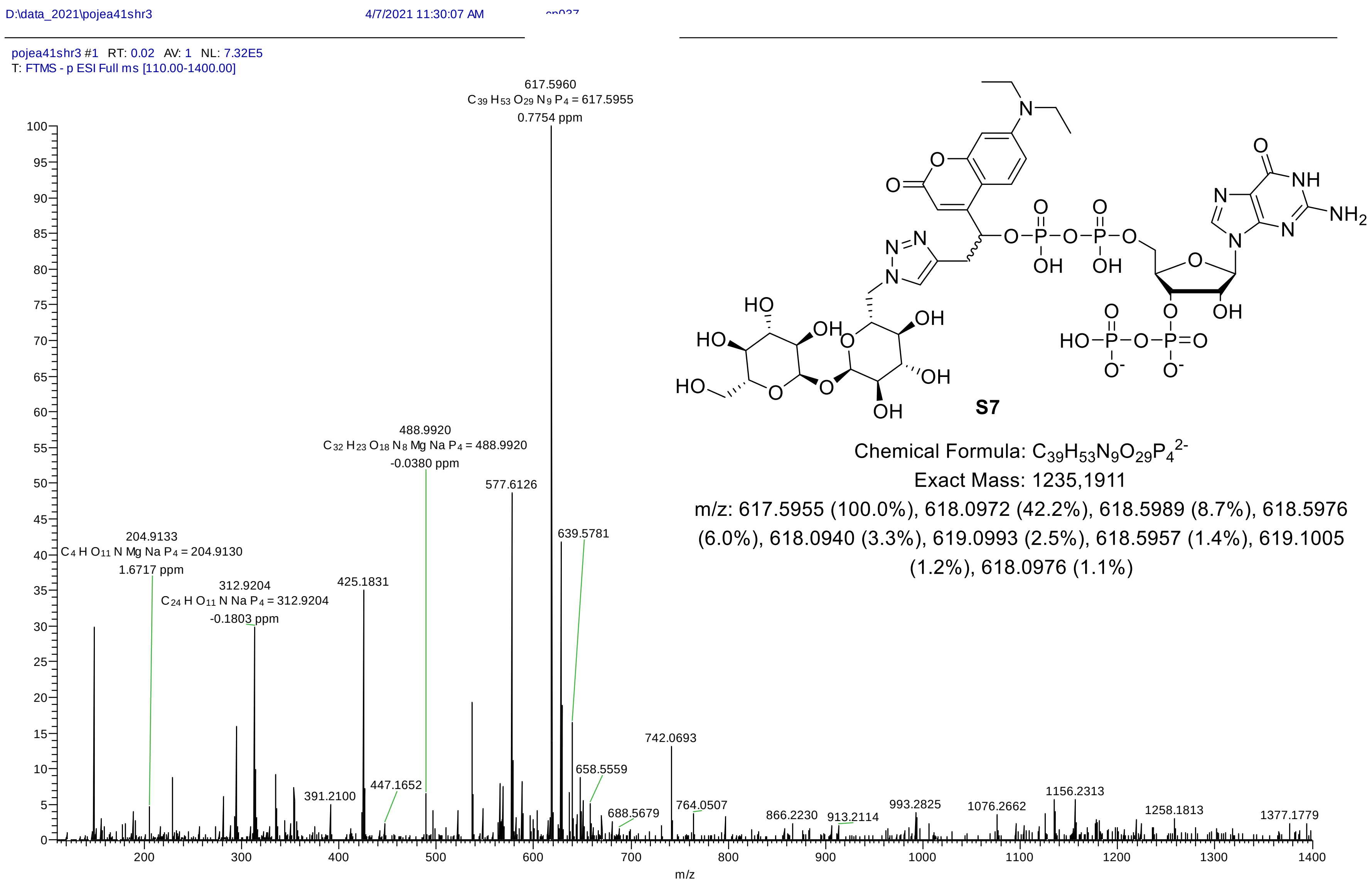

Chemical Formula: $C_{39}H_{53}N_9O_{29}P_4^{2-}$
Exact Mass: 1235,1911
m/z: 617.5955 (100.0%), 618.0972 (42.2%), 618.5989 (8.7%), 618.5976 (6.0%), 618.0940 (3.3%), 619.0993 (2.5%), 618.5957 (1.4%), 619.1005 (1.2%), 618.0976 (1.1%)

### HRMS (ESI$^-$) of S8

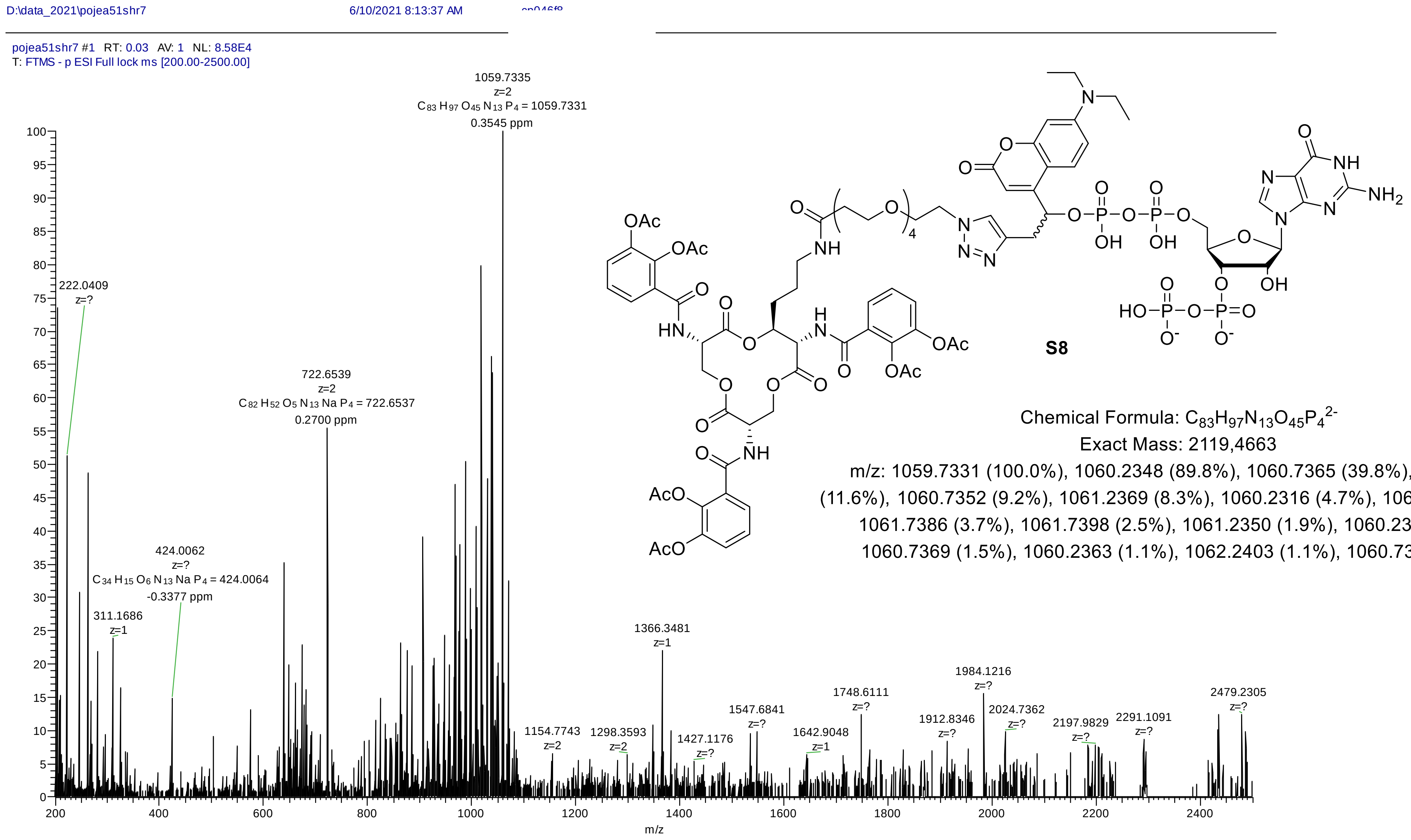

Chemical Formula: $C_{83}H_{97}N_{13}O_{45}P_4^{2-}$

Exact Mass: 2119,4663

m/z: 1059.7331 (100.0%), 1060.2348 (89.8%), 1060.7365 (39.8%), 1061.2382 (11.6%), 1060.7352 (9.2%), 1061.2369 (8.3%), 1060.2316 (4.7%), 1060.7333 (4.3%), 1061.7386 (3.7%), 1061.7398 (2.5%), 1061.2350 (1.9%), 1060.2352 (1.7%), 1060.7369 (1.5%), 1060.2363 (1.1%), 1062.2403 (1.1%), 1060.7379 (1.0%)

### HRMS (ESI$^-$) of S9

**Sample Spectra**

**- Scan (rt: 0.005-0.089 min) Sub**

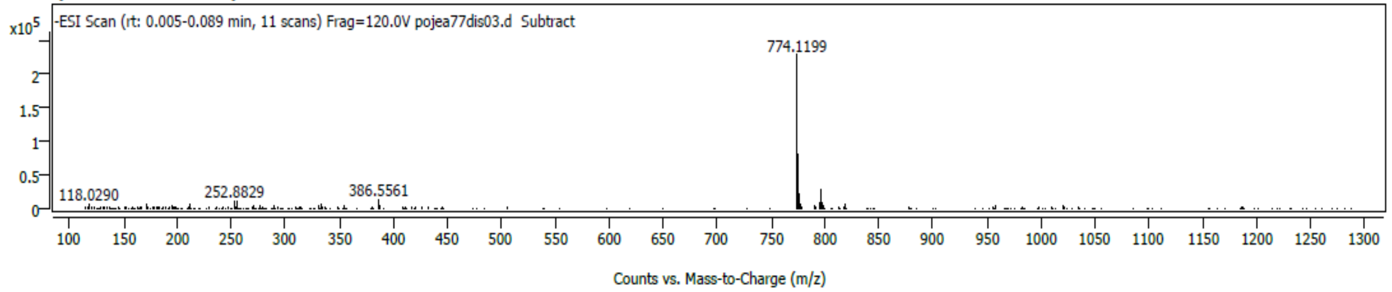

*Spectrum Peaks*

| m/z | Z | Abund | Abund % | m/z (Calc) | Diff (ppm) | Ion Species | Formula | Ion Type |
|---|---|---|---|---|---|---|---|---|
| 386.5561 | 2 | 12075 | 5.32 | 386.5567 | -1.45 | (M-2H)-2 | C27 H32 D N6 O15 P3 | |
| 773.1136 | | 69725 | 30.72 | 773.1128 | 0.91 | (M-H)- | C27 H31 D N6 O15 P3 | |
| 774.1199 | 1 | 226966 | 100.00 | 774.1207 | -1.04 | (M-H)- | C27 H32 D N6 O15 P3 | |
| 775.1226 | 1 | 80754 | 35.58 | 775.1236 | -1.42 | (M-H)- | C27 H32 D N6 O15 P3 | |
| 776.1248 | 1 | 21441 | 9.45 | 776.1260 | -1.55 | (M-H)- | C27 H32 D N6 O15 P3 | |
| 777.1281 | 1 | 5582 | 2.46 | 777.1285 | -0.44 | (M-H)- | C27 H32 D N6 O15 P3 | |
| 118.0290 | | 5989 | 2.64 | | | | | |
| 171.0654 | | 5151 | 2.27 | | | | | |
| 194.9046 | | 5001 | 2.20 | | | | | |
| 210.9714 | | 5978 | 2.63 | | | | | |
| 252.8829 | | 10770 | 4.75 | | | | | |
| 254.8809 | | 9886 | 4.36 | | | | | |
| 270.9377 | | 4492 | 1.98 | | | | | |
| 330.8599 | | 4822 | 2.12 | | | | | |
| 332.8575 | | 6534 | 2.88 | | | | | |
| 774.1802 | | 9782 | 4.31 | | | | | |
| 795.0957 | | 7954 | 3.50 | | | | | |
| 796.1011 | 1 | 27022 | 11.91 | | | | | |
| 797.1043 | 1 | 8538 | 3.76 | | | | | |
| 818.0846 | | 5339 | 2.35 | | | | | |

*Spectrum Identification Table*

| Best ID | Source | Name | Formula | Species | m/z | Diff (ppm) | CAS | Score | Score (Lib) | Score (DB) | Score (MFG) | Lib/DB |
|---|---|---|---|---|---|---|---|---|---|---|---|---|
| No | MFG | | C27 H32 D N6 O15 P3 | (M-H)- | 774.1199 | -1.15 | | 97.60 | | | 97.60 | |
| No | MFG | | C27 H32 D N6 O15 P3 | (M-2H)-2 | 386.5561 | -0.75 | | 95.19 | | | 95.19 | |
| No | MFG | | C27 H31 D N6 O15 P3 | (M-H)- | 773.1136 | 0.91 | | 47.04 | | | 47.04 | |
| No | MFG | | C27 H33 N6 O15 P3 | (M-H)- | 773.1136 | -1.09 | | 46.78 | | | 46.78 | |

Chemical Formula: $C_{27}H_{32}N_6O_{15}P_3^-$

Exact Mass: 773,1144

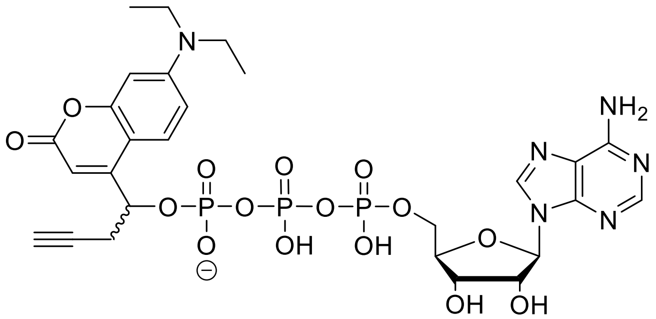

**D-ATP (S9)**

774.1199 enriched due to previous storage in deuterated solvent.

### HRMS (ESI$^-$) of S10

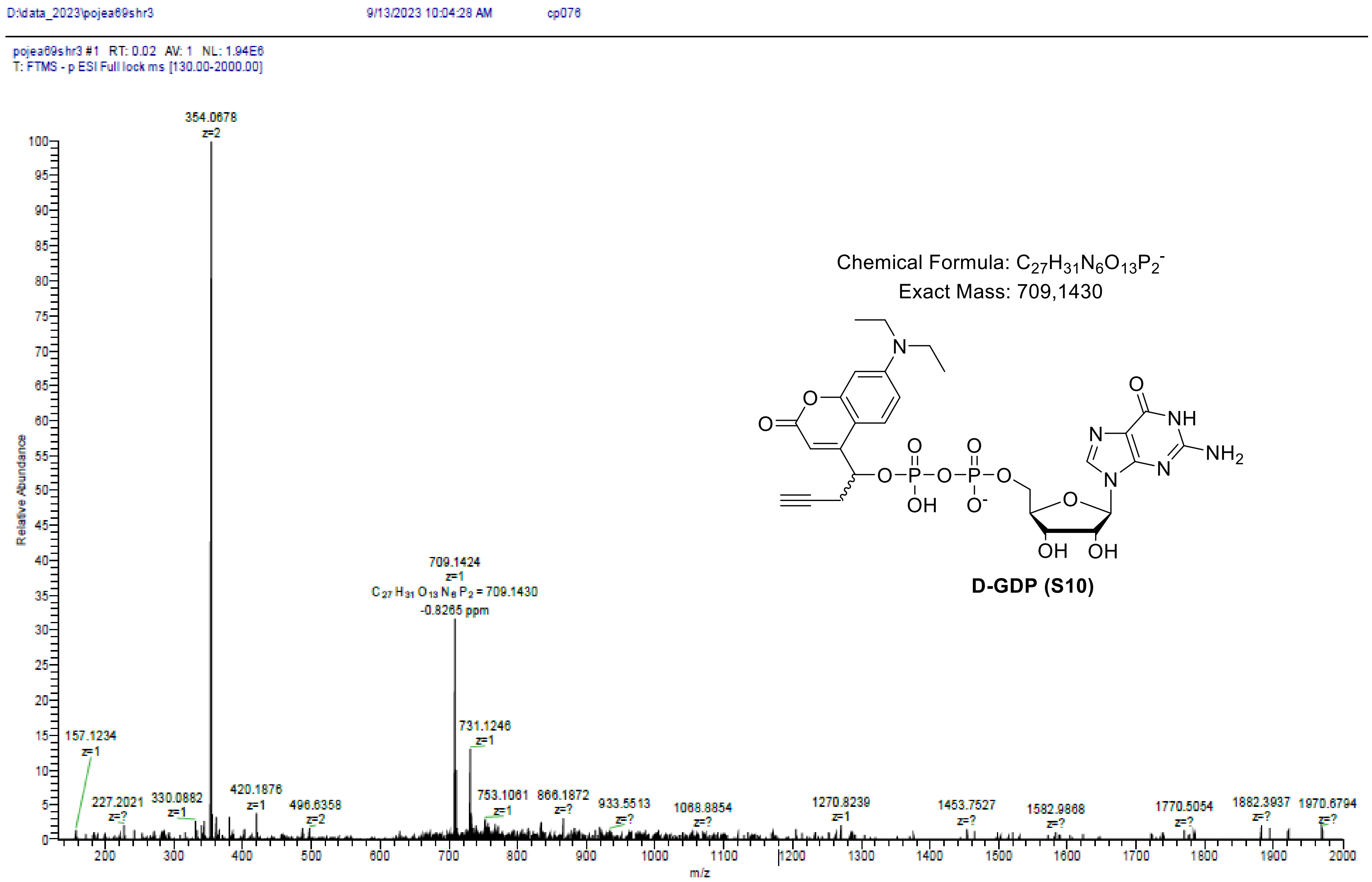

### HRMS (ESI$^+$) of S12

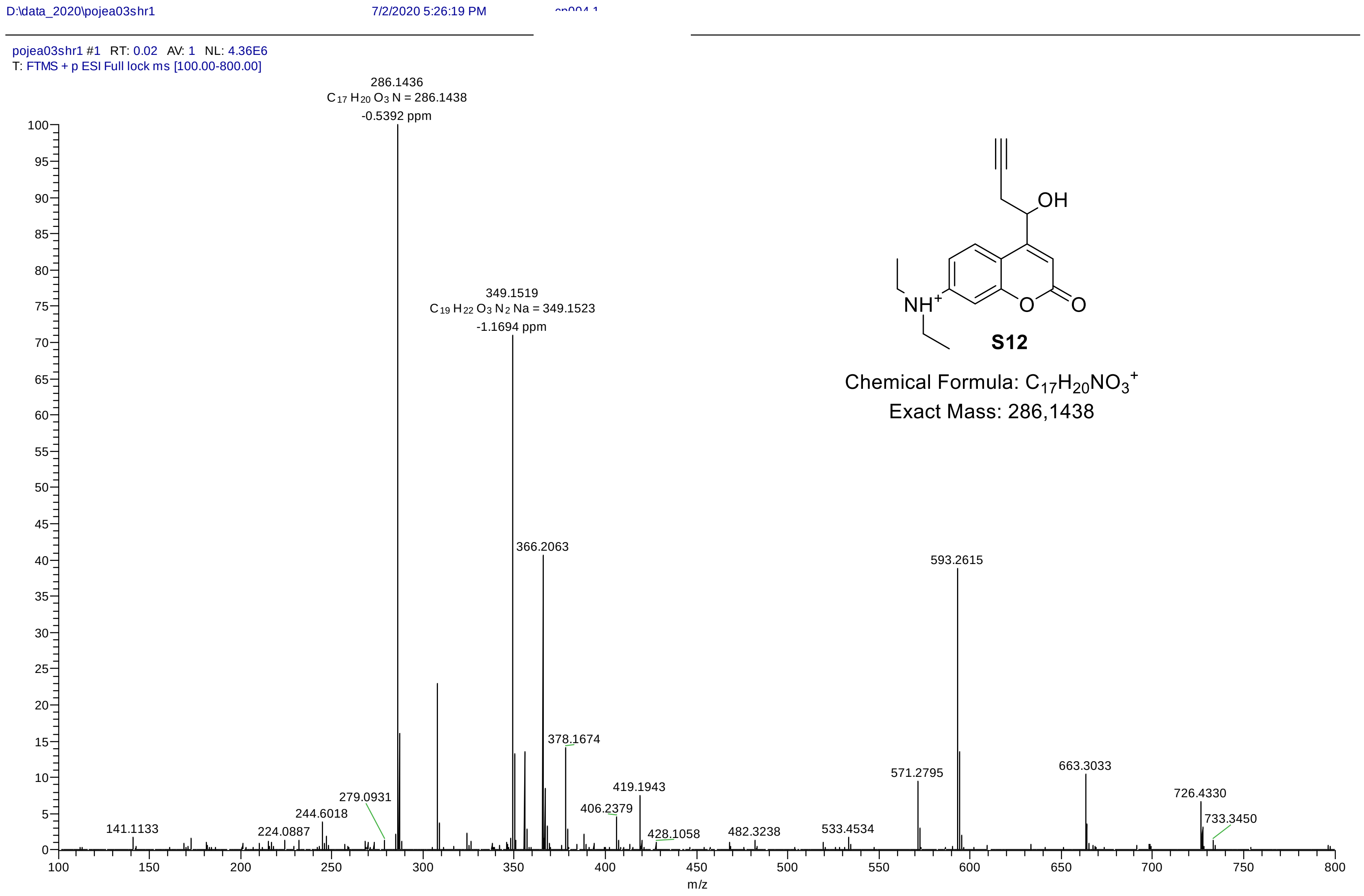

**S12**

Chemical Formula: $C_{17}H_{20}NO_3^+$
Exact Mass: 286,1438
